# Marker-assisted mapping enables effective forward genetic analysis in the arboviral vector *Aedes aegypti*, a species with vast recombination deserts

**DOI:** 10.1101/2021.04.29.442065

**Authors:** Chujia Chen, Austin Compton, Katerina Nikolouli, Aihua Wang, Azadeh Aryan, Atashi Sharma, Yumin Qi, Camden Delinger, Melanie Hempel, Antonios Augustinos, David W. Severson, Kostas Bourtzis, Zhijian Tu

## Abstract

*Aedes aegypti* is a major vector of arboviruses that cause dengue, chikungunya, yellow fever and Zika. Although recent success in reverse genetics has facilitated rapid progress in basic and applied research, integration of forward genetics with modern technologies remains challenging in this important species, as up-to-47% of its chromosome is refractory to genetic mapping due to extremely low rate of recombination. Here we report the development of a marker-assisted-mapping (MAM) strategy to readily screen for and genotype only the rare but informative recombinants, drastically increasing both the resolution and signal-to-noise ratio. Using MAM, we mapped a transgene that was inserted in a >100 Mb recombination desert and a sex-linked spontaneous red-eye (*re*) mutation just outside the region. We subsequently determined, by CRISPR/Cas9-mediated knockout, that *cardinal* is the causal gene of *re*, which is the first forward genetic identification of a causal gene in *Ae. aegypti*. This study provides the molecular foundation for using gene-editing to develop versatile and stable genetic sexing methods by improving upon the current *re-*based genetic sexing strains. MAM does not require densely populated markers and can be readily applied throughout the genome to facilitate the mapping of genes responsible for insecticide- and viral-resistance. By enabling effective forward genetic analysis, MAM bridges a significant gap in establishing *Ae. aegypti* as a model system for research in vector biology. As large regions of suppressed recombination are also common in other plant and animal species including those of economic significance, MAM will have broad applications beyond vector biology.

## Introduction

*Aedes aegypti* is a major vector of a number of arboviruses and filarial worms. Effective control of this one species could help reduce or prevent a number of vector-borne infectious diseases including dengue, chikungunya, yellow fever and Zika. Current strategies to reduce the incidence and burden of these diseases depend heavily on effective vector control, which is hindered by increasing insecticide-resistance. Novel control strategies, informed by improved understanding of mosquito biology, are urgently needed. Recent years witnessed rapid accumulation of genomic resources (e.g., (Matthews et al. 2018)) and effective applications of CRISPR/Cas9-mediated reverse genetic analysis of gene function in *Ae. aegypti* (e.g., (Hall et al. 2015); (Li et al. 2017); (Aryan et al. 2020)). However, successful integration of forward genetics with modern technologies to identify causal genes of important traits or phenotypes, as championed by many plant and animal geneticists (e.g. (Schneeberger 2014); (Navarro-Escalante et al. 2020); (Feng et al. 2021)), still awaits in this important vector species.

Recombination is fundamental to forward genetic screening and quantitative trait loci (QTL) mapping. Mapping a locus that contains the causal gene is possible when recombination separates the causal gene locus from its neighboring sequences, resulting in an enrichment of markers tightly linked to the causal gene in individuals manifesting the phenotype (Figure 1). However, *Ae. aegypti* has a lower per megabase recombination rate (~0.3 cM/Mb) than *Anopheles gambiae* (0.8 cM/Mb) and *Drosophila melanogaster* (1.6 cM/Mb) (Wilfert, Gadau, and Schmid-Hempel 2007). In addition, up to 47% of its chromosome surrounding the centromere belongs to so-called low recombination regions (LRRs), with rates much lower than the 0.3 cM/Mb average ((Juneja et al. 2014); (Fontaine et al. 2017); (Dudchenko et al. 2017)). Although it is not unique to *Ae. aegypti* to have LRR near centromeres, having nearly half of the genome in LRR further complicates genetic efforts to map genes that determine particular traits. Thus, the low recombination rate in *Ae. aegypti* limits both the resolution and efficiency of these genetic mapping studies. We are not aware of any report of the successful determination of a causal gene using forward genetics in *Ae. aegypti*, despite the intense interest in identifying genes that underly insecticide- and pathogen-resistance or genes that could be used as selectable markers for genetic sexing (Koskinioti et al. 2021; Ward et al. 2021). The presence of extensive regions with very low recombination rates, or recombination deserts, has also been reported in many plant and animal species (Stapley et al. 2017a, 2017b). Therefore, forward genetic mapping in these recombination deserts is a significant and broadly important challenge.

**Figure 1.**
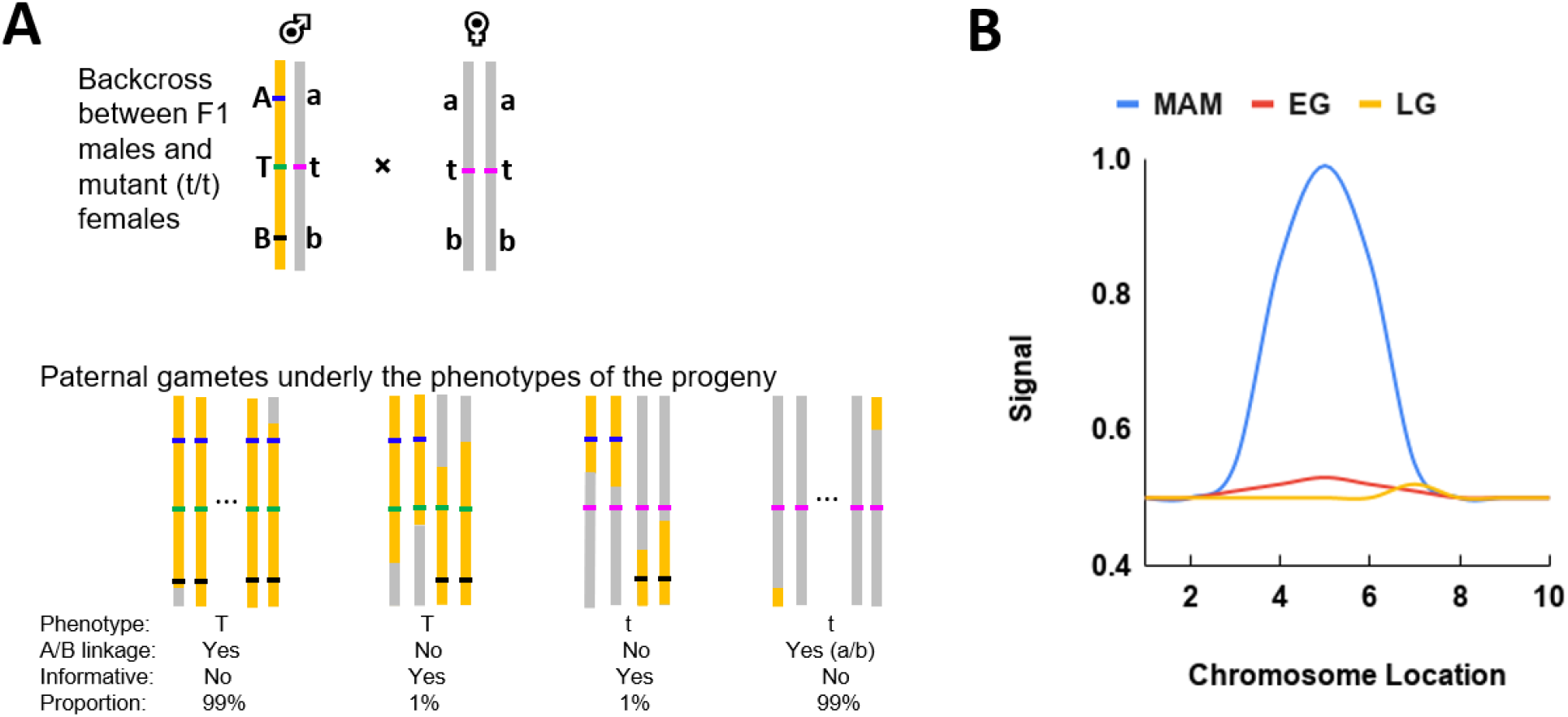
The marker-assisted mapping (MAM) strategy. **A)** The progeny of a backcross between F1 males and mutant (t/t) females can be categorized according to the mutant phenotype (T/t versus t/t) and the linkage between two easy-to-screen flanking markers (A and B). Only paternal gamete is shown. In the example shown here, the genetic distance between A and B is 1 cM. Thus, 99% of the progeny that retain the linkage between the A and B markers are not informative and can be ignored. **B).** Expected signal strengths using either marker-assisted mapping (MAM) when only the informative progeny (T/t and t/t progeny showing breakage between markers A and B), extreme genotyping (EG) when hundreds of T/t and t/t progeny are analyzed, or limited genotyping (LG) when a limited sample of the T/t and t/t progeny are genotyped. In the current example, genotyping 100 T/t and 100 t/t progeny will likely only include 1 informative T/t and 1 informative t/t recombinant between the flanking markers, severely limiting the signal to noise ratio and the resolution of the mapping result. The position of the “peak” in LG could also be misleading due to limited sampling.

Here we report the development of a marker-assisted-mapping (MAM) strategy and demonstrate its success in rapidly mapping a transgene insertion in a region of suppressed recombination near the sex locus in *Ae. aegypti*. Using the same method in conjunction with introgression analysis, we also identified the causal gene for a spontaneous red-eye (*re*) mutation in *Ae. aegypti,* which was first reported nearly 60 years ago (McClelland 1962; McClelland 1966). The *re* mutation is currently used for developing genetic sexing strains (GSSs) due to its sex-linkage and the robustness of the *re* strain (Koskinioti et al. 2021). This study provides the molecular foundation for future improvements and expansion of GSSs and enables effective forward genetic analysis in *Aedes aegypti*. As large regions of suppressed recombination are also common in other plant and animal species including those of economic significance, MAM will have broad applications beyond vector biology.

## RESULTS

### The marker-assisted mapping (MAM) strategy

The general strategy of MAM and its advantages are depicted in Figure 1A and 1B, respectively. Rare and informative recombinants are selected for genotyping analysis based on the breakage of linkage of the selectable markers (A and B) flanking the target locus (T or t), thus greatly increasing both the resolution and efficiency of genetic mapping. In the example shown in Figure 1, 99% of the non-informative progeny are ignored, thus both the resolution and signal-to-noise-ratio are increased by 100 fold. If informative positive samples and informative negative samples are genotyped in bulk, MAM represents a novel application of the bulk-segregant analysis (Schneeberger 2014) that can identify causal genes in previously inaccessible regions of suppressed recombination.

### The chromosomal region under investigation and the markers

The low recombination regions (LRRs) in *Ae. aegypti* (Dudchenko et al. 2017), which encompass up to 47% of a chromosome, show little or no recombination at the resolution of the available genetic crosses (Juneja et al. 2014). The same pattern of vast regions of suppressed recombination is reproduced (Supplemental Figure S1) using the most recent PacBio-based assembly (Matthews et al. 2018). We recently obtained a sex-linked transgenic insertion line named P10 (Supplemental Figure S2, Table 1), which expresses a polyUb-driven GFP. The transgene is inserted at position 224.7 Mb of the q arm of chromosome 1, as shown by Oxford Nanopore sequencing (Supplemental Figure S3). The genetic distance between P10 and the sex locus (M locus, 151.68-152.95 Mb on chromosome 1) is approximately 1.02 to 1.55 cM (Table 1). Thus, the per megabase recombination rate in the ~72 Mb (224.68-152.95=71.73 Mb) region is 0.014-0.022 cM/Mbp, approximately 73 to 114 fold lower than that of the *D. melanogaster* (1.6 cM/Mb) (Wilfert, Gadau, and Schmid-Hempel 2007). A previously reported recessive red-eye (*re*) locus is also on the q arm of chromosome 1, and it is 2-3 cM away from the M-locus, with an unknown chromosomal location (Augustinos et al. 2020; Koskinioti et al. 2021). We first seek to establish the marker-assisted mapping (MAM) strategy by mapping the location of the P10 insertion using the maleness (M locus) and wildtype black-eye (+/+ or +/re) as flanking markers (Figure 2).

**Table 1.**
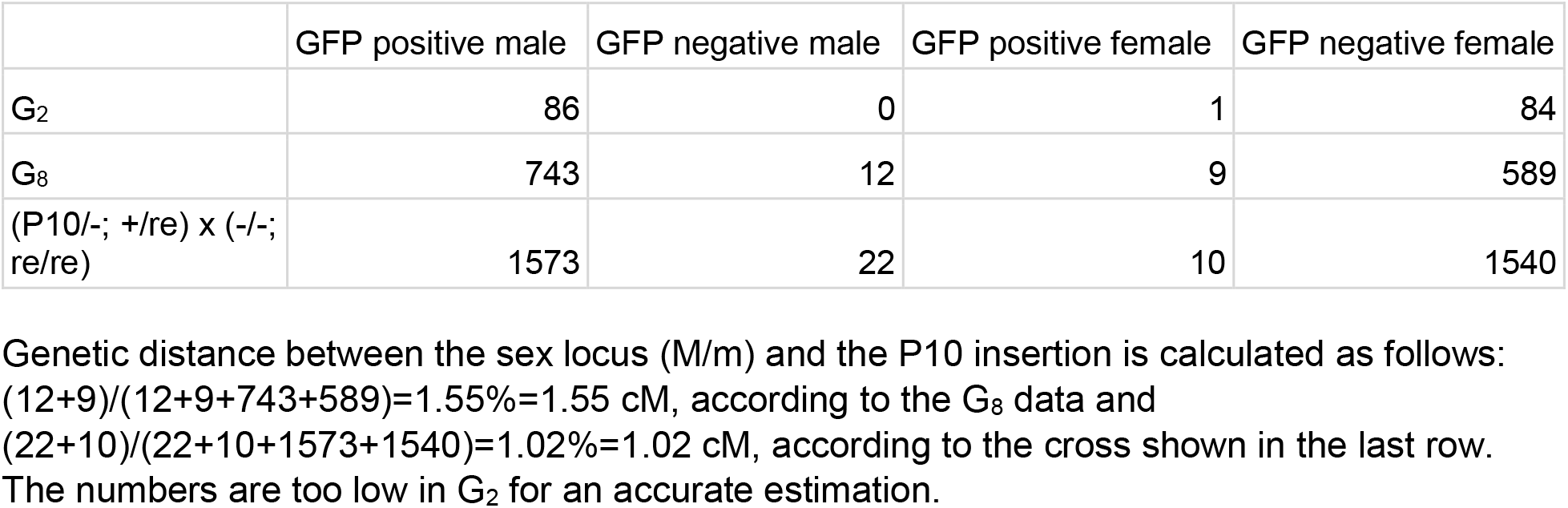
Sex-linkage of the P10 (GFP) transgene insertion.

**Figure 2.**
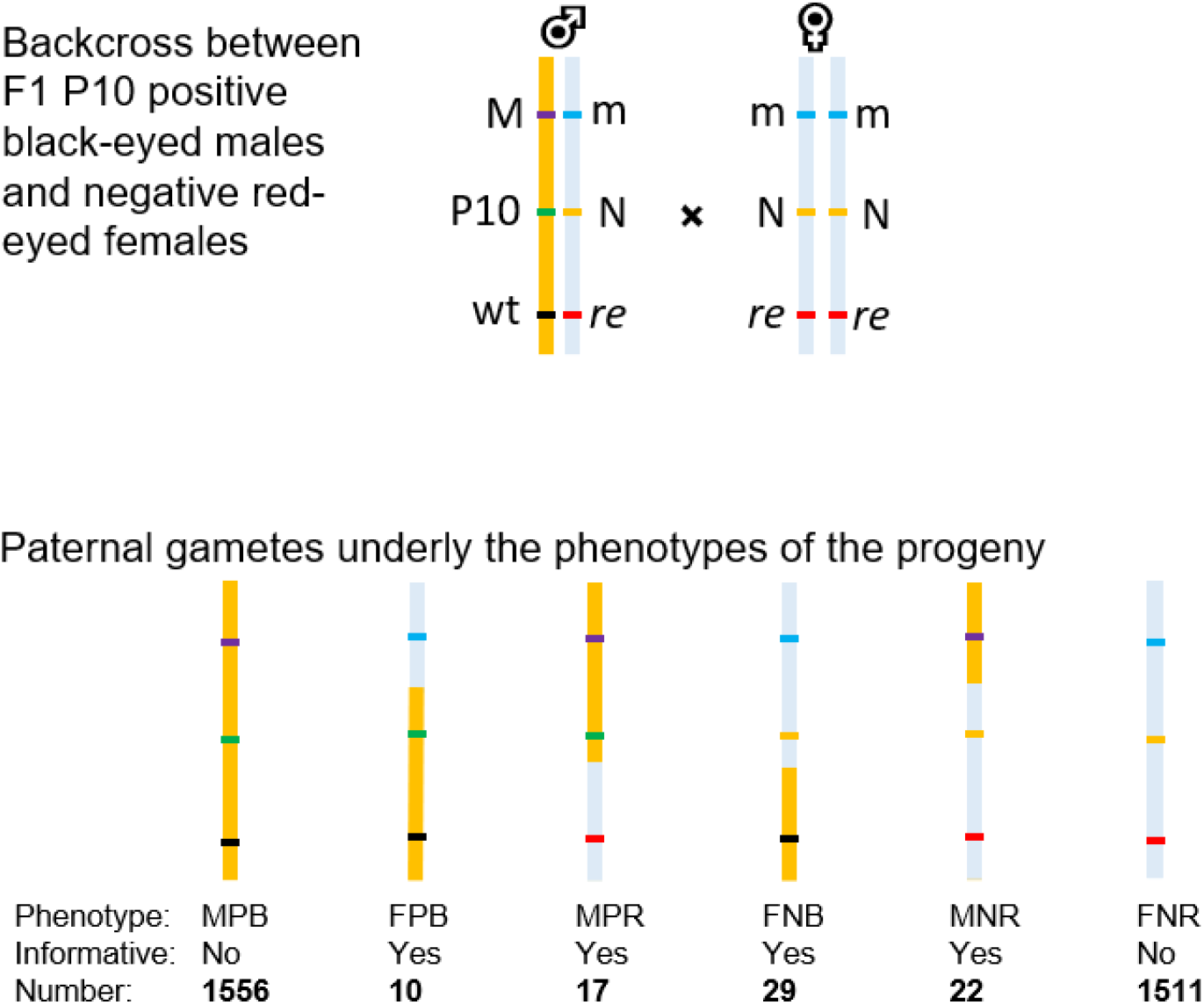
Marker-assisted mapping of the P10 transgene insertion. An F1 backcross and the resulting F2 progeny. Double crossovers, which are negligible in this case, are not shown. M: the dominant male-determining locus; M/m are at ~152 Mb of chromosome 1. P10: positive for the P10 GFP transgene insertion; N: negative or no transgene insertion; *re*: the recessive red-eye allele; wt: wildtype black-eye allele. To denote the phenotype, M is male while F is female; P is P10 transgene positive while N is negative; B is black-eyed while R is red-eyed. For example, MPB is a male P10 positive black-eyed individual. While crossover could potentially happen anywhere between markers, only one example is provided. The answer regarding to informative or not is in the context of mapping P10. The genetic distance between P10 and the M/m loci is (10+22)/(1556+10+17+29+22+1511)=32/3145=1.02%, or 1.02 cM. The genetic distance between P10 and *re* is (17+29)/3145=1.46%, or 1.46 cM. The genetic distance between *re* and the M/m locus is (10+22+17+29)/3145=2.48%, or 2.48 cM.

### Rapid and effective mapping of the P10 transgene using marker-assisted mapping

To map P10 which expresses GFP, rare and informative recombinants are selected for genotyping analysis based on the breakage of linkage between the two flanking markers, maleness and black-eye. A backcross (Figure 2) produced dozens of informative recombinants among thousands of non-informative progeny. We grouped the informative recombinants into four categories for Illumina sequencing (Supplemental Table S1) as we are also interested in mapping *re* in a later analysis (see below). The four groups include male positive red-eye (MPR); female positive black-eye (FPB); female negative black-eye (FNB); male negative red-eye (MNR). As shown in Figure 3, when MPR and FPB (all informative P10-positive recombinants) are compared with FNB and MNR (all informative P10-negative recombinants), MAM successfully mapped the P10 location to be between 223.3 and 226.1 Mb with 95% confidence. The midpoint of the peak is 224.7 Mb, precisely the location of the P10 insertion.

**Figure 3.**
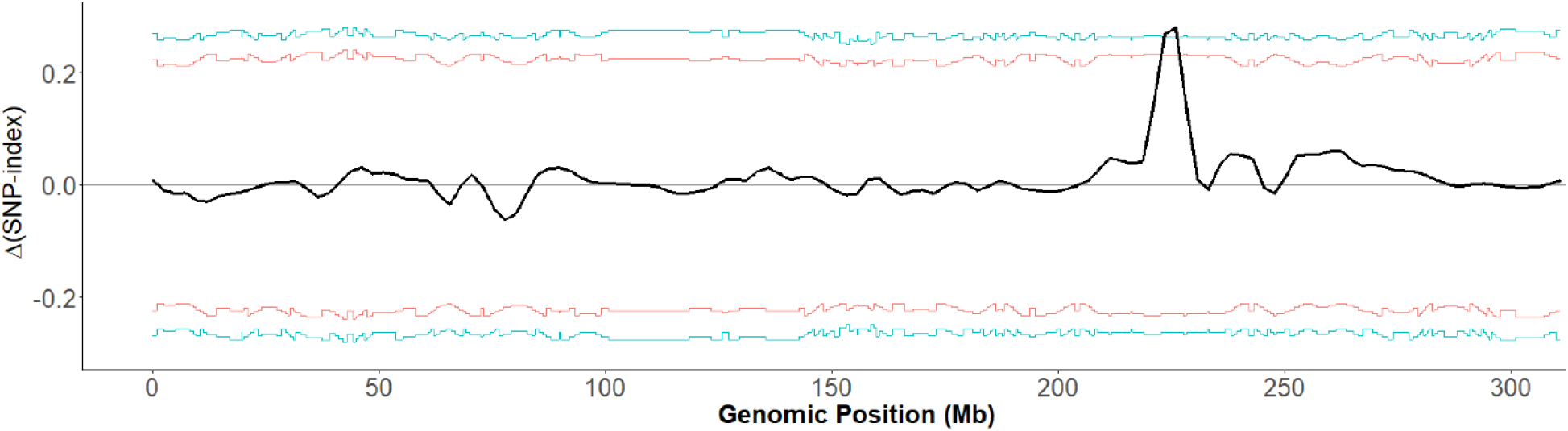
Marker-assisted mapping (MAM) identifies the location of the P10 transgene insertion. The tricube‐smoothed Δ(SNP‐index) is shown in 1 Mb sliding windows. The Δ(SNP‐index) value is calculated by comparing all informative P10 positive recombinants (MPR and FPB) with all informative P10 negative recombinants (FNB and MNR). The corresponding two-sided confidence intervals are shown as red (90%) or blue (95%) lines. X-axis is the genomic position on the homomorphic sex chromosome 1. The confidence intervals were estimated using 10,000 replicate simulations for all SNP positions with the given read depths. P10 is located at position 224,678,447 on chromosome 1, according to ONT sequencing (Supplemental Figure S3). Positions from 223288153 to 226114389 exceeds 95% confidence and the left and right inflection points of this peak are 223412798 – 225833455, respectively.

### Marker-assisted mapping of *re*

The above-mentioned sequencing data are also used to map *re*, as we can compare the recombinant male red-eye and the recombinant female black-eye samples. A potential caveat is the lack of resolution from the telomeric side of *re*, as the cross was originally intended for mapping P10 and a marker to the telomeric side of *re* was not used. However, given the interest in developing *re* as a marker for GSS (Koskinioti et al. 2021), the fact that *re* is one of the earliest reported spontaneous mutations isolated in *Ae. aegypti* (McClelland 1966; McClelland 1962), and the lack of any successful examples of forward genetic identification of causal genes in *Ae. aegypti*, we attempted to map *re* using the existing data. As shown in Figure 4A, the *re* locus is mapped to the region between 271 Mbp and 278 Mbp with 95% confidence.

**Figure 4.**
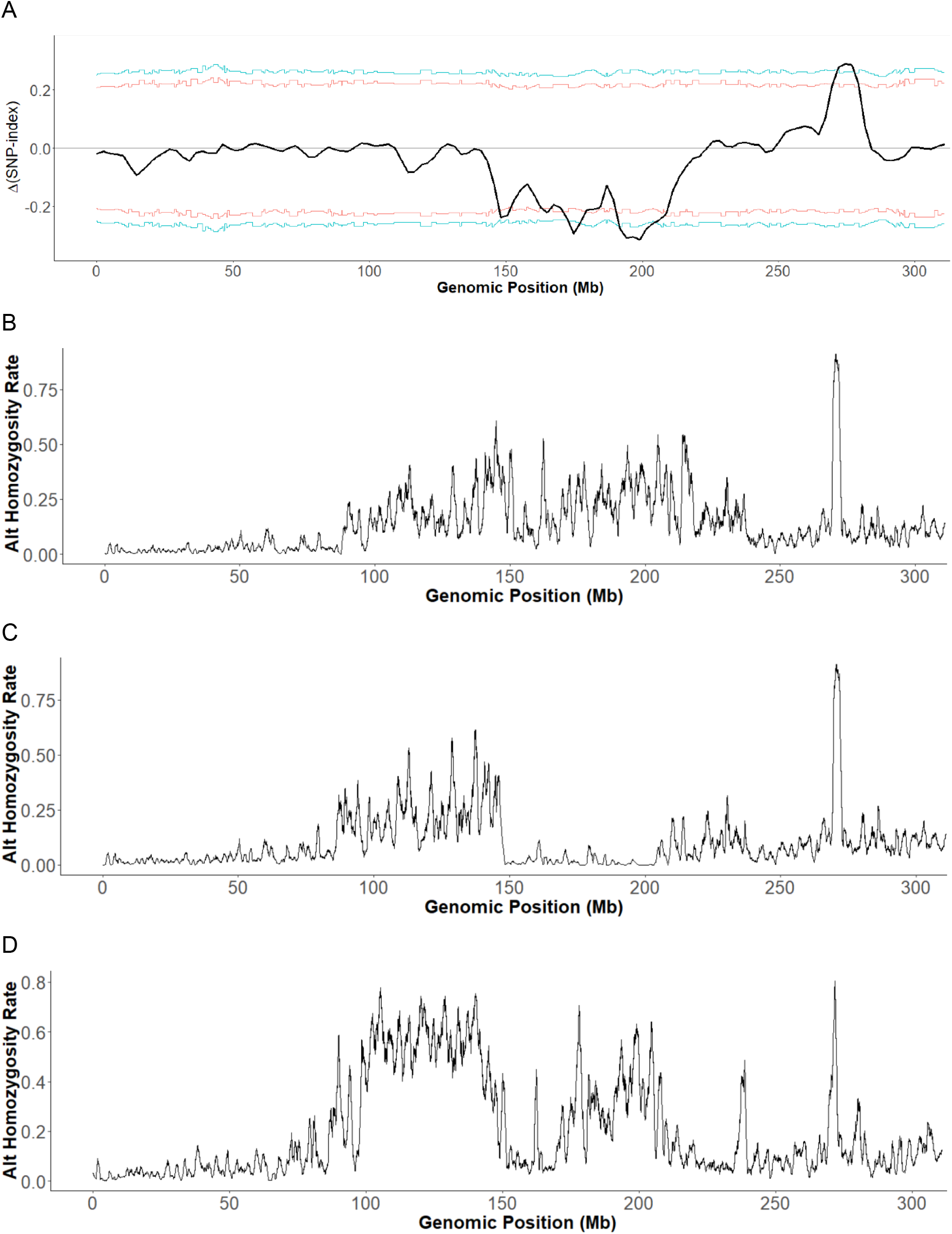
Mapping of *re*. **A)** Marker-assisted mapping (MAM) of *re* by plotting the Δ(SNP‐index) values in 1 Mb sliding windows. The corresponding two-sided confidence intervals are shown as red (90%) or blue (95%) lines. X-axis is the genomic position on the homomorphic sex chromosome 1. The confidence intervals were estimated using 10,000 replicate simulations for all the SNP positions with the given read depths. The broad depression between 150-210 Mb likely reflects the sex-linked differences while the peak around 275 Mb is likely the location of *re*. Positions from 271,120,117 to 277,843,549 exceeds 95% confidence and the left and right inflection points of this peak are 271,970,355 and 276,823,052, respectively. **B)-D)**. Homozygosity plots of red-eyed hybrids from F8ThaiF9Mex_F (B), F8ThaiF9Mex_M (C) and F7PAKF3BRA_F (D). X-axis is the genomic location on chromosome 1. Y-axis is the homozygosity of the alternative or non-reference alleles, which is calculated as (the number of unique alternative SNPs showing 100% frequency)/(the number of all unique alternative SNPs) in a 1 Mb window. We focused on unique alternative SNPs as we are only interested in SNPs showing 100% frequency. The peak for F8ThaiF9Mex_F (B), F8ThaiF9Mex_M (C) and F7PAKF3BRA_F (D) are 270-271 Mb, 270-271 Mb, and 271-272 Mb, respectively.

### Mapping *re* using the introgression lines

The above mapping result for *re* is unexpected as the previously published data indicate that the genetic distance between the sex locus and the 271-278 Mb region is at least 10 cM (Dudchenko et al. 2017), much higher than the 2.48 cM observed between the sex locus and *re* (Figure 2 legend). We sought independent evidence for the location of *re*. As a part of an ongoing effort to develop GSS, the Rexville red-eye strain (Koskinioti et al. 2021) was introgressed with *Ae. aegypti* from various regions for a minimum of 11 generations to establish GSSs of various local genetic backgrounds (Supplemental methods). The *re*-containing strain used in the above-mentioned MAM experiment is the RED strain, which is homozygous for the *re* locus on chromosome 1, the *s* (*spot abdomen*) locus on chromosome 2, and the *b/t* (*black tarsus*) on chromosome 3 (Craig and Hickey 1967). Genetic complementation crosses confirmed that the *re* in RED and the red-eye locus in the Rexville-derived introgression lines share the same mutant gene (Supplemental Figure S4). These introgressed GSSs contain the *re* mutation in divergent genetic backgrounds, which could be used to narrow down the location of *re*, by identifying regions of enriched homozygosity. To increase the cost-effectiveness, we performed Illumina sequencing of red-eyed hybrids (Supplemental Table S1) from crosses between different GSSs to further highlight the enrichment of homozygosity near the *re* locus. As shown in Figure 4B-D, homozygosity is the highest in a region around 271-272 Mb. We note that the homozygosity peak of the three red-eyed introgression samples falls on the centromeric side of the MAM peak (271-278 Mb), which is consistent with expectation as the P10 marker is to the centromeric side of *re* and there is no marker to the telomeric side of *re*.

### Identification of a candidate gene *cardinal*

To further narrow down the candidate region, we focused the homozygosity analysis on the 265-280 Mb region using combined SNP data from all three hybrid red-eyed samples. As shown in Figure 5, the region with the highest homozygosity is around 271.5 Mb. Interestingly, a gene encoding a chorion peroxidase, which is an ortholog of the *Drosophila* eye pigment gene *cardinal*, is located between 271.574 and 271.583 Mb and this gene is expressed throughout development after 20hr post egg deposition and its transcripts are enriched in the brain of both sexes in *Ae. aegypti* (Matthews et al. 2018). Thus, *cardinal* is the top candidate for the causal gene for the spontaneous *re* mutation. Indeed, the *cardinal* gene sequences in red-eyed samples show a number of non-synonymous mutations leading to changes of amino acid residues that are conserved across divergent mosquitoes (Supplemental Figure S5).

**Figure 5.**
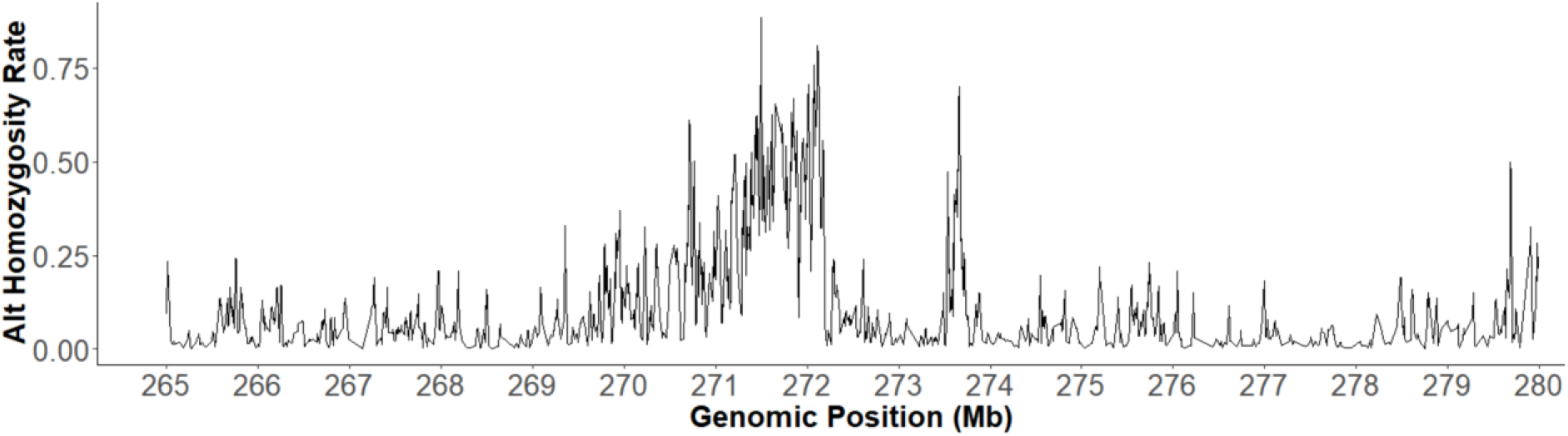
Homozygosity analysis between 265 Mbp and 280 Mbp on chromosome 1 by pooling all three hybrid red-eyed samples. Y-axis is the homozygosity of the alternative or non-reference alleles, which is calculated as (the number of unique alternative SNPs showing 100% frequency)/(the number of all unique alternative SNPs) in a 20 kb window. The highest level of homozygosity is at position 271,505,401.

### CRISPR/Cas9-mediated knockout of *cardinal* produced somatic knockouts showing mosaic red-eyed G_0_ individuals

To increase the likelihood to observe a somatic *cardinal* knockout phenotype in the G_0_, we first performed CRISPR/Cas9-mediated knockout in heterozygous +/*re* individuals. The sgRNAs used to target *cardinal* are shown in Figure 6A and provided in Supplemental Table S2. Mosaic red-eyes were indeed observed in +/*re* G_0_ individuals (Figure 6B; Supplemental Figure S6). Such somatic mosaicism may reflect knockout of the wildtype *cardinal* allele (cd-) in some cells, resulting in the red-eye phenotype in the cd-/*re* genotyped cells. Thus, the result is consistent with *cardinal* being the causal gene of the *re* mutation. However, we cannot rule out the possibility that the mosaic phenotype may have nothing to do with *re* and it simply resulted from somatic knockout of both copies of the *cardinal* gene. Regardless, this result confirmed that the sgRNAs are effective and the *cardinal* gene is indeed involved in eye-color in *Ae. aegypti*, as recently demonstrated in *Culex quinquefasciatus* (Feng et al. 2020).

**Figure 6.**
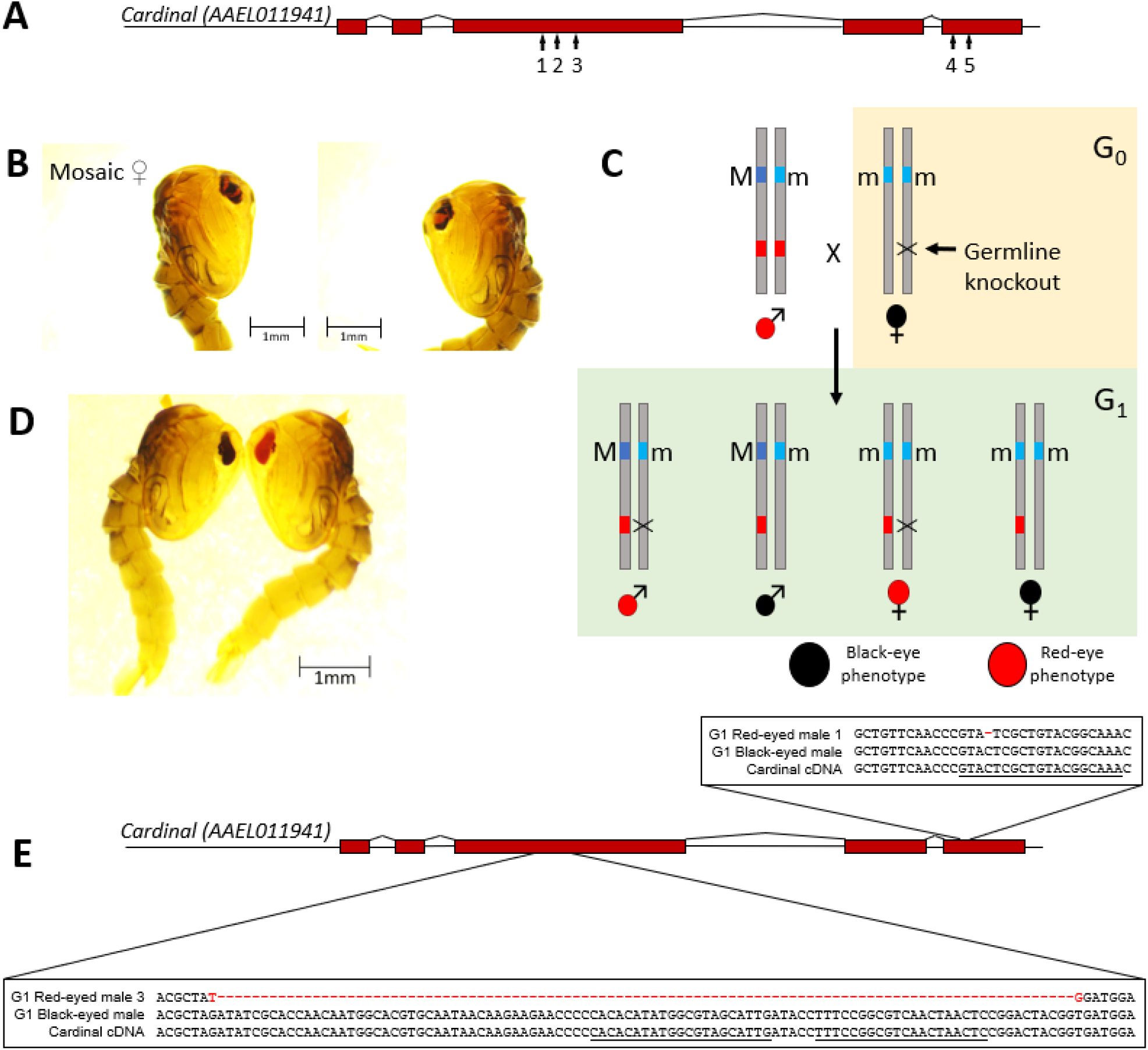
A) Illustration of the *cardinal* gene and the relative positions of the 5 sgRNAs. B) Somatic knockout of *cardinal* in a heterozygous (+/re) female results in a mosaic eye. Cas9 mRNA and *cardinal* sgRNAs were injected into eggs that were collected from red-eyed females (re/re) mated with wildtype Liverpool strain males (+/+). The two images are the right and left eyes of the same female. Images for the control black-eyed siblings and a similar mosaic-eyed male individual are shown in Supplemental Figure S6. C) Design for germline knockout of *cardinal*. D) G_1_ black-eye (left) and red-eye (right) phenotype. Four red-eyed G_1_ females and seven red-eyed G_1_ males were identified among approximately 500 black-eyed G_1_ progeny. E) Sequences show a 1bp deletion (in red) in red-eyed G_1_ male #1 and a 99bp deletion (in red) in red-eyed G_1_ male #3. The 1 bp deletion was found in the other sequenced red-eyed G_1_ individuals (male #1, 5, 6 and 7). None of the deletions was found in black-eyed G_1_ individuals. The sgRNA sequences near the deletion sites are underlined.

### Germline knockout of *cardinal* showed that it is the causal gene of the *re* mutant

To isolate germline knockout of *cardinal* (Figure 6C), sgRNAs and Cas9 mRNAs were injected in embryos collected from wildtype Lvp females (+/+) mated with wildtype Lvp males (+/+). The resulting G_0_ females were black-eyed and selected to mate with red-eyed males (*re*/*re*). Four female and seven male red-eyed G_1_ individuals were identified among approximately 500 black-eyed G_1_ individuals (Supplemental Table S3; Figure 6C-D). PCR and subsequent Sanger sequencing of 5 red-eyed G_1_ males using primers flanking the sgRNAs (Supplemental Table S4) identified two types of mutations. A single base deletion that results in a frameshift was found in 4 of the G_1_ males (#1, 5, 6, and 7) and a 99 base deletion in the other (G_1_ male #3; Figure 6E). However, none of these deletions were found in the black-eyed G_1_ individuals (Figure 6E). These red-eyed G_1_ individuals can only have one *re* allele from the father and one cd- allele from the mother. Therefore, *cardinal* knockouts (cd-) resulting from either 1- or 99-bp deletion fail to complement *re*, suggesting that *cardinal* is indeed the causal gene of the *re* mutant.

### Establishing and characterizing a homozygous *cardinal* knockout

To obtain homozygous knockout individuals, black-eye G_2_ females (+/cd-) were backcrossed to their father, red-eye G_1_ male#3 (also +/cd-; Figure 7A). Their G_3_ offspring were evaluated and four genotypes were expected and four phenotypes were observed: red-eyed males, black-eyed males, yellow-eyed females, and black-eyed females (Figure 7B, Supplemental Table S5). The yellow-eyed females are homozygous for the 99 bp *cardinal* deletion. The yellow-eyed phenotype is observed in the pre-pupal compound eye (Figure 7B and Supplemental Figure S7). However, the yellow-eyes gradually turned into red-eyes over the course of pupae development and adult emergence (Figure 7C-D). Therefore, the yellow-eye phenotype is in fact *yellow-turn-red*. No yellow-eyed L2/L3 instars were found and the ocelli of the yellow-eyed pre-pupa are red, indicating that the *yellow-turn-red* phenotype is associated with the remodeling of the compound eye during metamorphosis. The sibling red-eyed males inherited one copy of the *re* spontaneous mutation and one copy of the *cardinal* knockout (Figure 7A), again indicating that *cd-* does not complement *re*. As expected, when the G_2_ females (+/cd-) were crossed to RED males (*re*/*re*), approximately half of the offspring are red-eyed in each sex (Supplemental Figure S8 and Table S6).

**Figure 7.**
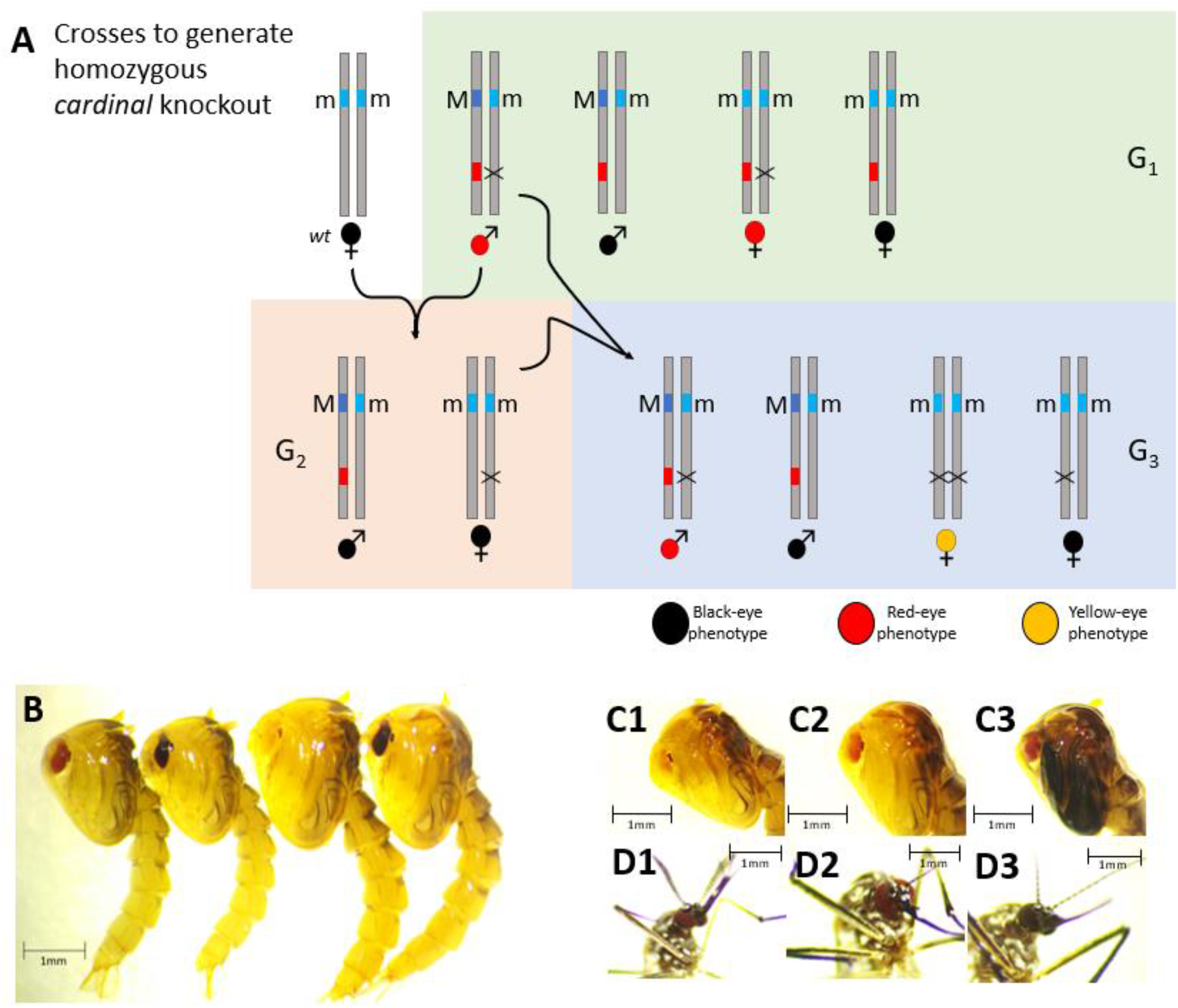
A) Crossing scheme to generate homozygous *cardinal* knockout. B) Four phenotypes observed in the G_3_ progeny from a cross shown in panel A. From left-to-right: red-eyed male, black-eyed male, colorless-eyed female, black-eyed female. The G_1_ red-eye male has the 99 bp deletion (G1 red-eye male#3). C) The eyes in the colorless-eyed female turn red over the course of pupal development. C1-3: 1, 2, and 3 day old pupae. To highlight the eye phenotype, the contrast was adjusted to 20% and the brightness was adjusted to 20% (for C1-2) and 40% (C3). D) 1-day-old G_3_ adult eye phenotypes of the red-eyed male (D1), yellow-turn-red-eyed female (D2), and black-eyed female (D3). To highlight the eye phenotypes, the contrast of images was adjusted to 25% and the brightness to 25% (D1-2) and 50% (D3). All unmodified photos can be found in the Supplemental images file.

## DISCUSSION

### Marker-assisted mapping (MAM) enables effective forward genetic analysis in and outside the recombination desert in *Ae. aegypti*

There has not been a report of a successful forward genetic discovery of the causal gene of any mutant phenotype in *Ae. aegypti*, despite the significant interest in the fundamental biology and vectorial capacity of this key arboviral vector. The overall low recombination rate (0.3 cM/Mb, (Wilfert, Gadau, and Schmid-Hempel 2007)) and the existence of vast recombination deserts known as the LRRs ((Juneja et al. 2014); (Fontaine et al. 2017); (Dudchenko et al. 2017); Supplemental Figure S1) represent a key bottleneck for forward genetics in *Ae. aegypti*. We designed the marker-assisted mapping strategy to overcome this key bottleneck by selecting and genotyping only the rare but informative recombinants between easy-to-screen markers flanking the target locus, thus, drastically improving the resolution, signal-to-noise ratio, and cost-effectiveness for genetic mapping (Figure 1). Using MAM, we successfully mapped P10, a dominant transgene insertion that shows an extremely low rate of recombination relative to the sex locus (0.014-0.022 cM/Mbp) and the red-eye (*re*) marker (1.46 cM/46 Mb=0.032 cM/Mb). The MAM peak spans from 223.3 Mb to 226.1Mb with the midpoint being 224.7 Mb, precisely the location of the P10 insertion. Therefore, we have demonstrated the power of MAM in making forward genetics accessible in the LRRs which were previously inaccessible or challenging.

The same set of MAM data was also used to map *re*, a recessive spontaneous mutant. Despite the lack of a marker to the telomeric side of *re*, we were able to map *re* to a broader peak between 271.1 Mb and 277.8 Mb at a 95% confidence interval. The location of the *re* causal gene *cardinal* is at 271.5 Mb, much closer to the centromeric side (271.1 Mb) of the peak than to the telomeric side. The lack of resolution at the telomeric side, as indicated by the 6.3 Mb distance between 271.5 Mb and 277.8 Mb, likely reflects the lack of a marker to “assist” in selecting informative recombinants. The resolution from the telomeric side could be even worse if the telomeric side of *re* were in the LRR. We also showed that homozygosity analysis using hybrids between repeatedly introgressed strains can also facilitate mapping studies. The introgressed GSSs are highly valuable but time consuming to create as a minimum of 11 generations were required, thus may not be available to other mapping studies. However, even in the absence of one of the two flanking markers and without the use of additional resources such as these introgression lines, MAM can still identify the candidate locus with the causal gene being proximal to the marker-side of the peak.

As mentioned earlier, the location of *re* is somewhat unexpected as the previously published data indicate that the genetic distance between the sex locus and the 271.5 Mb region is at least 10 cM (Dudchenko et al. 2017), much higher than the 2-3 cM between the sex locus and *re* (Augustinos et al. 2020; Koskinioti et al. 2021 and Figure 2 legend). Although such a discrepancy may result from different genetic backgrounds or other conditions that may affect the rate of recombination (Craig and Hickey 1967), it may also reflect a limitation of the previous mapping studies in which a small number of progenies were genotyped. The ease of screening for the P10 and *re* markers affords a high-resolution estimate of the recombination rate based on rapid screening of a large number of progenies. This is especially important when working with loci in the LRRs.

The identification of *cardinal* as the causal gene for the spontaneous *re* mutant is the first report of a successful identification of the causal gene using forward genetics in *Ae. aegypti*. The *cardinal* gene is outside the previously defined LRRs ((Juneja et al. 2014); (Fontaine et al. 2017); (Dudchenko et al. 2017)). Thus, we have shown that MAM enabled forward genetics both in and outside the LRRs. Our study focused on the 1q arm which is approximately 159 Mb long, spanning from the centromeric sex locus (151.68-152.95 Mb) to the telomere (located at 311 Mb). P10 (located at 224.7 Mb) and *re* (located at 271.5 Mb) are ~72 Mb and 119.5 Mb away from the sex locus, respectively. Therefore, it is not necessary to have densely populated markers for MAM to succeed. With the ease in obtaining transgene insertions and the availability of targeted knock-in (Kistler, Vosshall, and Matthews 2015), MAM can be readily established throughout the entire *Ae. aegypti* genome to greatly facilitate forward genetic studies. Knocking in a marker near a candidate gene can also help locate the precise location of the causal gene before embarking on functional analysis (Ding et al. 2016). Coupling MAM with targeted genotyping, in place of whole-genome sequencing, will further improve the efficiency and reduce the cost of genetic mapping efforts.

### Insights into the function of *cardinal*

We did not observe any phenotypic differences between *re/re* and *re*/cd- individuals. However, the homozygous progeny of a 99 bp deletion in *cardinal* showed a phenotype of eyes gradually turning from yellow to red. The yellow-eyed phenotype was first observed in the pre-pupal compound eye and gradually turned into red-eyes over the course of pupae development and adult emergence. Thus, the phenotype is associated with the remodeling of the compound eye during metamorphosis. The *cardinal* gene encodes a peroxidase which catalyzes the formation of the eye pigment xanthommatin from 3-hydroxykynurenine in the ommochrome pathway and is believed to also participate in the synthesis of ommin ((Howells, Summers, and Ryall 1977); (Harris et al. 2011); (Osanai-Futahashi et al. 2016); (Zhang et al. 2017); (Xu et al. 2020)). Indeed, knockout of *cardinal* near its C-terminus in *Culex quiquefasciatus* resulted in a red-eye mutation visible from the larval to the early adult (Feng et al. 2020), while an amorphic *cardinal* knockout in the diamond back moth resulted in yellow-eyed adults which turned into red-eyed as they aged (Xu et al. 2020). The gradual change from yellow to red in eye color was explained by the slow auto-oxidation of the yellow pigment 3-hydroxykynurenine leading to the eventual accumulation of the red pigment xanthommatin. Similarly, the 99 bp *cardinal* deletion in *Ae. aegypti* likely resulted in either an amorphic mutation or a more severe hypomorphic mutation than the spontaneous *re*.

### Implications to the development of GSSs

The development of GSSs has been a key challenge to SIT programs from the start. Indeed, it took more than 20 years to develop one of the most successful GSSs for the Mediterranean fruit fly, the *Ceratitis capitata* VIENNA 8 GSS (Franz et al. 2021). Only recently the causal gene of the *white-pupae* (*wp*) mutation used in the VIENNA 8 GSS was identified (Ward et al. 2021). The necessity to develop a mosquito GSS does not only stem from economic concerns but also from the behavioral traits of the species since only females feed on vertebrate hosts and transmit pathogens. Therefore, GSS that allows for effective sex-separation is a critical component of any genetic approaches to control mosquito-borne infectious diseases. Several mosquito GSSs were developed for *Anopheles* species since the 1970s (Kaiser P.E. 1978, Robinson AS 1986, Curtis CF 1978, Lines JD 1985, Yamada et al. 2012, 2015). However, the high genetic instability of these strains and in some cases the reliance on insecticide-resistance as a selection marker are problematic. Recent field releases of transgenic, irradiated and/or *Wobachia*-infected sterile males showed promising results for suppressing Aedes populations in areas of moderate sizes (reviewed in (Papathanos et al. 2018), Kittayapong et al. 2019, (Zheng et al. 2019), (Crawford et al. 2020), Balatsos et al. 2021). However, robust, precise, and economical sex-separation remains a significant bottleneck to large-scale implementations of genetic measures to control mosquito-borne infectious diseases.

The current red-eye GSS has shown excellent performance and genetic stability under laboratory conditions (Koskinioti et al. 2021) and an induced chromosomal inversion further increased the precision and effectiveness of these GSS lines (Augustinos et al. 2020). In this study we identified the causal gene of the red-eye phenotype. The identification of the red-eye causal gene will facilitate the development of GSS lines, not only for *Ae. aegypti* but also for other medically important mosquitoes such as *Ae. albopictus* and *Anopheles* species. For example, the red-eye phenotype can be created by knocking out *cardinal* in these species using CRISPR/Cas-mediated genome editing approaches. Tight sex-linkage could be achieved either by X-ray or CRISPR/Cas9-induced chromosomal translocation, or by CRISPR/Cas9-mediated knock-in of a wildtype *cardinal* in the male sex locus. This would significantly accelerate the process of the GSS development thus saving time and resources.

### Concluding remarks

Genomic, transcriptomic and population genomic resources have rapidly accumulated in the past few years for *Ae. aegypti* (e.g., (Matthews et al. 2018); (Mysore, Li, and Duman-Scheel 2018); (Rose et al. 2020)). Effective application of genome-editing and improved efficiency of transposon-mediated transformation ((Hall et al. 2015); (Li et al. 2017); (Aryan et al. 2020)) make *Ae. aegypti* increasingly amenable to genetic research. We anticipate that MAM will significantly improve forward genetic studies in *Ae. aegypti* and facilitate the mapping of genes responsible for insecticide- and viral-resistance, and/or other large-effect loci identified from QTL analyses. In addition to overcoming the challenges of mapping in the recombination deserts, MAM offers another advantage as time-consuming assays for insecticide- or viral-resistance will only need to be performed on informative recombinants. MAM fills a gap in establishing *Ae. aegypti* as a model system for basic and applied research in vector-borne infectious diseases. As large regions of suppressed recombination are also common in other plant and animal species including those of economic significance (Stapley et al. 2017a, 2017b), MAM will have broad applications beyond vector biology.

## Materials and Methods

### Mosquito rearing

The Liverpool (LVP) strain (obtained from www.BEI.org) and the RED strain (obtained from the Severson laboratory at the University of Notre Dame) of *Ae. aegypti* were maintained at Virginia Tech at 26-28°C and 60-70% relative humidity with a 14/10 hour day/night light cycle. Adult mosquitoes were fed 10% sucrose and fed blood using artificial membrane feeders and defibrinated sheep’s blood (Colorado Serum Company; Denver, CO).

### The P10 Transgenic line

A piggyBac donor plasmid (500ng/ul, Supplemental Figure S2) that contains an EGFP transformation marker driven by the *Ae. aegypti polyubiquitin* promoter and was co-injected with an *in vitro* transcribed piggyBac mRNA (300ng/ul) into less than 1 hr old embryos of the Liverpool strain of *Ae. aegypti* (Coates et al. 1998). The piggyBac-hsp70-transposase (Handler et al. 1998) was used as a template for *in vitro* transcription using the mMessage mMachine T7 Ultra kit (Thermofisher), followed by MEGAclear (Thermofisher) column purification. Approximately 1200 embryos were injected, resulting in 150 G_0_ females and 160 G_0_ males). Surviving G_0_ females were mated to Liverpool males in pools of 20-25. Each G_0_ male was mated individually with 5 Liverpool females in individual cages and mosquitoes from 15-20 of these cages were merged into one large pool. G_1_ larvae were screened for green fluorescence using a Leica M165 FC fluorescence microscope. Positive G_1_ individuals were out-crossed to Liverpool mosquitoes to ensure that all transgene cassettes were stably inherited to the G_2_ generation.

### Identification of the P10 insertion site

Oxford Nanopore sequencing was performed using genomic DNA isolated from four G_5_ P10-positive males. The Qiagen Genomic Tip DNA Isolation kit (Cat. No. 10,243 and 19,060, Qiagen, Hilden, Germany) was used. These four males were homogenized using a hand held pestle motor mixer (Cole Palmer) with a sterile disposable plastic pestle (USA Scientific) for approximately 10 seconds in the lysis buffer. The homogenate was incubated with 300 mAU Proteinase K (Qiagen, Hilden, Germany) overnight at 55°C, and then transferred into a 15 mL conical tube for centrifugation at 5000g for 15 minutes at 4°C to remove debris. The lysate was loaded on a 20-G column for purification following the protocol. The purity, approximate size, and concentration of the DNA were measured using a nanodrop spectrophotometer, 0.5% agarose gel electrophoresis, and Qubit dsDNA assay, respectively. Approximately 2000 ng gDNA was used to prepare the library using the 1D ligation sequencing kit (SQK-LSK108, Oxford Nanopore Technologies, UK). Adapter ligation was performed at room temperature for 45 minutes. Approximately 530 ng of the prepared library was loaded on the flow cell. Approximately 7.6 Gbases reads were obtained and submitted to NCBI ((PRJNA718905, sample SAMN18579040). Four long reads that contain the piggyBac insertion were identified and used to map the location of the P10 insertion (Supplemental methods 1; Supplemental Figure S3).

### Sample Collections for Marker assisted mapping

The crossing scheme is shown in Figure 2. GFP-positive, black-eyed P10 males (n=66) were mated with GFP-negative, red-eyed females (n=37) to generate the F1 progeny. The GFP-positive, black-eyed F1 males (n=26) were backcrossed with GFP-negative, red-eyed females (n=46) to generate F2 progeny. The F2 progenies were screened and counted. The following numbers of individuals were pooled for genomic DNA isolation and Illumina sequencing: male positive black-eyed (n=60), male positive red-eyed (n=12), female positive black-eyed (n=20), female negative red-eyed (n=60), male negative red-eyed (n=11) and female positive black-eyed (n=7). Illumina sequencing coverage and other statistics are provided in Supplemental Table S1.

### MAM data analysis

Adapters and low quality Illumina reads were trimmed by Trimmomatic v0.38 (Bolger, Lohse, and Usadel 2014) and quality control were performed using FastQC v0.11.8. [https://www.bioinformatics.babraham.ac.uk/projects/fastqc/]. Variants were called using GATK v4.1 (McKenna et al. 2010) following the “Best Practices workflow” ((DePristo et al. 2011); (Van der Auwera et al. 2013)). First, paired reads of each sample were aligned to the *Aedes aegypti* genome L5 from VectorBase [www.vectorbase.org] using BWA-MEM v0.7.17 (Li and Durbin 2009) and sam files were converted to bam files and sorted by samtools (Li et al. 2009). PCR duplicates were marked by GATK. SNPs of each sample were called by GATK using the Haplotype caller mode. GVCF files were combined by GATK CombineGVCFs. Combined GVCF files were converted to VCF files by GATK GenotypeGVCFs. Variants are selected from VCF to a table by GATK VariantsToTable for the subsequent Bulk Segregant Analysis by QTLseqr (Takagi et al. 2013; Mansfeld and Grumet, 2018). Variants were filtered using GATK. The number of SNPs, minimum read depth, and tricube-smoothed Δ(SNP-index) within a 1 Mb sliding window were calculated by QTLseqr. The slope of Δ(SNP-index) was used to calculate the inflection point. Simulations with 10000 bootstrap replicates were used to calculate the two-sided confidence intervals. The simulation was performed based on data derived from read depths and the type and size of the population. Alternative allele frequency was calculated per bulk and Δ(SNP-index) was calculated over multiple replications for each bulk under the null hypothesis that no QTLs exist. The percentile from the simulation was used to estimate the confidence intervals.

To map the P10 insertion, male negative red-eye (MNR) and female negative black-eye (FNB) were used as the control while the female positive black-eye (FPB) and male positive red-eye (MPR) were used as the experimental group. Variants were filtered based on 10 < total read depth < 400, 0.2 <= sum of reference allele frequency of two bulks <= 0.8, per sample read depth >= 5 and GATK GQ score >= 99. Approximately 2.7 million SNPs remained after variant filtering. To map the *re* locus, female positive black-eye (FPB) and female negative black-eye (FNB) were used as the control while male negative red-eye (MNR) and male positive red-eye (MPR) were used as the experimental group. Variants were filtered based on the same parameters as mentioned above. Approximately 2.5 million SNPs remained after variant filtering.

### *Ae. aegypti* strains used for homozygosity analysis

All strains described in this section were maintained in the insectary of the Insect Pest Control Laboratory (Joint FAO/IAEA Center of Nuclear Techniques in Food and Agriculture, Seibersdorf, Austria) at 27 ± 1 °C, 80% relative humidity and a 12/12 h day/night photoperiod. Adult mosquitoes were provided a 10% sucrose solution and females were fed with porcine blood twice per week. The blood used was collected in Vienna, Austria during routine slaughtering of pigs in a nationally authorized abattoir, conducted at the highest possible standards strictly following EU laws and regulations. Egg collections were initiated 72 h after the last blood feeding using moistened oviposition papers (white germination paper, Sartorius Stedium Biotech, Austria). The Rexvillle strain was provided by Dr. Margareth Capurro at the Department of Parasitology, University of Sao Paulo, Brazil. All Rexvillle individuals have red eye color which is evident throughout all developmental stages and darkens as adults age. The Rexvillle strain has been shown to complement the red-eye mutation (*re*) in the RED strain (Supplemental Figure 4). The Rexville red-eye phenotype was introgressed into the BRA, Mex, Thai, and Pak strains to create red-eye genetic sexing strains (GSSs) with different genomic backgrounds in a separate study (Augustinos et al. 2020; Koskinioti et al. 2021). The 11-generation introgression protocol is described in Supplemental Methods 2.

### Crosses and sample preparation for the homozygosity analysis

The above mentioned introgressed GSS contain the *re* mutation in divergent genetic backgrounds, which could be used to narrow down the location of *re*, by identifying regions of enriched homozygosity. Thirty F8 red-eye females from the THAI Red-Eye GSS and fifteen F9 red-eye males from the MEX Red-Eye GSS were mass-crossed. All progeny was red-eyed and a pool of 20 freshly emerged adult females (F8ThaiF9Mex_F) and a pool of 20 males (F8ThaiF9Mex_M) were used to extract DNA using the QIAGEN Genomic tip 20/G kit (Qiagen, Germany). In another sample, one BRA male was crossed with five virgin BRA females to create an isomale BRA line. Males from this isomale line were crossed with red-eyed Rexville females and the F1 were sib-mated. F2 red-eye females were then backcrossed with the wild-type black-eyed males from the isomale line. F3 black-eyed males were crossed with the red-eyed females originated from the F7 PAK Red-Eye GSS. Twenty red-eyed female progeny (F7PAKF3BRA_F) were used for HMW DNA extraction, as described above. Genomic DNA from F8ThaiF9Mex_F, F8ThaiF9Mex_M and F7PAKF3BRA_F were sequenced by Illumina (Supplemental Table S1).

### Homozygosity analysis

Mapping and variant calling were performed as described for MAM except that samples are the hybrid sequencing samples F8ThaiF9Mex_F, F8ThaiF9Mex_M and F7PAKF3BRA_F. As the coverage is lower than those in the MAM analysis, variant filtering was adjusted based on 5 < total read depth < 500, only one alternative in the variant and GATK GQ score >= 50. Within a given sliding window size (e.g., 1 Mb or 20 kb), the alternative homozygosity rate is calculated by

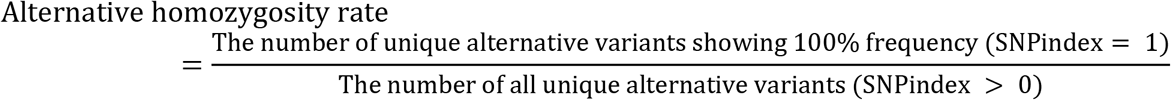

We focused on unique alternative SNPs as we are only interested in SNPs showing 100% frequency.

### Mosquito imaging

Imaging larvae, pupae, and adult specimens were performed using an AMSCOPE LED Trinocular Zoom Stereo Microscope 3.5X-180X and 10MP USB3 Camera (https://www.amscope.com/applications/veterinary-zoology/entomology/led-trinocular-zoom-stereo-microscope-3-5x-180x-and-10mp-usb3-camera.html).

### CRISPR/Cas9-mediated Knockout of cardinal

For somatic knockout of cardinal, LVP males were mated with RED female mosquitoes in a G-1 cross. The resulting G_0_ embryos were injected with a mixture containing 300 ng/uL Cas9 mRNA and 100 ng/uL each of sgRNAs 1, 2, and 3. Somatic knockout of *cardinal* was evaluated visually at the larval pupal stages. To produce a germline knockout of cardinal, an injection mixture containing 300 ng/uL Cas9 mRNA and 100ng/uL each of sgRNAs 1, 2, 4, and 5 was injected into LVP embryos. Male and female G_0_ adult survivors were separated and mated with RED adults of the opposite sex. Specific cardinal mutations were validated by sequencing PCR amplicons from each sgRNA target region. Briefly, genomic DNA extracted using the QiaAMP gDNA-micro kit (Qiagen) or Zymo Quick-gDNA (Zymo Research) was used as the template for PCR using primers listed in Table S4. PCR amplicons were purified using the Nucleospin PCR and gel clean-up kit (Machery-Nagel) and sequenced using Sanger (Genomics Sequencing Center, Virginia Tech). Trace files used to validate germline knockouts of *cardinal* are in the Supplemental File.

### sgRNA design and sgRNA synthesis

sgRNAs were designed to target two regions within the predicted coding sequence of the *cardinal* gene (Figure 6A). The first sgRNA target region is located in exon 3 at 664-737bp from the predicted start codon and the second in exon 5 at 1,698-1,776bp and is near the region targeted for the recent knockout of *cardinal* in *Culex quinquefaciatus* (Feng et al. 2020). All five sgRNAs (Table S2) were designed using CHOPCHOP ((Labun et al. 2016; Labun et al. 2019), (Montague et al. 2014)) and CRISPOR (Concordet and Haeussler 2018) web tools. sgRNA DNA templates were synthesized by PCR using previously described protocols (Bassett et al. 2013) using primers listed in Supplemental Table S4. sgRNA was synthesized by T7 in vitro transcription using the MEGAscript T7 kit, and purified using the MEGAclear kit (Thermo Fisher Scientific) then aliquoted for embryonic microinjections (Basu et al. 2015).

## Supporting information

Four ONT reads that contain P10

Sanger Trace Files

Original photo

## Acknowledgements

This work is supported by NIH grants R01AI123338 and 1R01AI157491 and the Virginia Agriculture Experimental Station. AC is supported by a fellowship from the Robert Wood Johnson Foundation. This study is also financially supported by the Joint FAO/IAEA Insect Pest Control Subprogramme of the Joint FAO/IAEA Centre of Nuclear Techniques in Food and Agriculture and the U.S. State Department in the frame of the ‘Surge Expansion of the Sterile Insect Technique (SIT) to Control Mosquito Populations that Transmit the Zika Virus’ project. This study benefitted from discussions during meetings in the frame of the IAEA Coordinated Research Project “Generic approach for the development of genetic sexing strains for SIT applications”. We thank Song Li for discussions on data analysis. We thank the Chloé Lahondère laboratory for use of the Amscope microscope imaging system and for advice on imaging. We thank Margareth Capurro, Jorge Aurelio Torres Monzon, Pat Kittayapong, and Muhammad Misbah-ul-Haq for kindly providing the BRA, Mex, Thai and Pak strains used in the present study.

## Data deposition

Illumina and Oxford Nanopore sequencing data are deposited and available at https://www.ncbi.nlm.nih.gov/sra/PRJNA718905.

