## Supplemental Methods, Tables and Figures for "Marker-assisted mapping enables effective forward genetic analysis in the arboviral vector *Aedes aegypti*, a species with vast recombination deserts"

**Supplemental information**

**Supplemental methods 1 and 2**

**Supplemental Figures S1-S7**

**Supplemental Tables**

**Supplemental data:**

**1) The four Oxford Nanopore reads that contain the P10 transgene insertion**

**2) Trace files of Sanger sequencing**

**3) Original image files**

**Supplemental method 1. Determine the location of the P10 insertion.** Sequencing was performed using genomic DNA isolated from a pool of four G_5_ GFP positive (P10) males. The fastq file is submitted to NCBI (PRJNA718905, sample SAMN18579040). Approximately 7.6 Gbases reads were obtained. BLASTN, using the piggyBAC (pBAC) transgene construct including the pBAC arms and the GFP sequence (Supplemental Figure S2) as the query, identified 4 reads that had a pBAC insertion. All four ONT reads were retrieved and used as query in a BLASTN search against the AaegyL5 genome assembly (Matthews et al. 2018). The tabular output of the BLASTN result is shown below and the four ONT reads are provided as supplemental data.

```
Query (ONT reads)		           Subject (chr)			%ID	Length	Qstart	Qend	Sstart	Send	E-value
e677388e-9ab7-49f3-bf3f-d42041a58300	NC_035107.1_224600001	93.91	23755	168	23221	78384	54959	0.0
a52f6291-2d70-49b2-9048-f0bd9b812ad0	NC_035107.1_224600001	91.28	5622	33	5480	72910	78378	0.0
fe307069-d616-41c4-8026-58873d6065ba	NC_035107.1_224600001	92.34	9609	6860	16166	78977	88375	0.0
fe307069-d616-41c4-8026-58873d6065ba	NC_035107.1_224700001	91.72	7850	27577	35213	1	7671	0.0
173ccd42-fdc8-4e1c-b262-0d6c006819e6	NC_035107.1_224600001	94.29	36773	5551	41340	78384	42123	0.0
```

As shown above, all four piggyBAC containing reads matched to chromosome 1 at position 224 Mb. The *Aedes aegypti* genome was fragmented into 100 kb pieces for Blastn to help reveal potential repeats. NC_035107.1_224600001 refers to the 100 kb fragment of chromosome 1 (NC_035107.1) that starts at 224600001. NC_035107.1_224700001 refers to the 100 kb fragment of chromosome 1 (NC_035107.1) that starts at 224700001. Note that fe307069-d616-41c4-8026-58873d6065ba encompasses two 100 kb fragments. Also note that single molecule sequencing (ONT) reads has >5% error rate, which is reflected in the 91-94% identity score in the table. An example of the alignment between a ONT read flanking the piggyBAC insertion and the AaegyL5 genome is also shown in Supplemental Figure S3, which identified the exact insertion site the hallmark target site duplication.

**Supplemental method 2. Introgression Protocol, using the Brazilian strain BRA as an example.** This protocol is designed to transfer the red-eye mutation into a Brazilian genomic background (BRA), aiming to create a red-eye genetic sexing strains (GSS) in the desired background. This is important to avoid regulatory problems in sterile insect applications. The same protocol is used to introgress the red-eye phenotype into other backgrounds including Thailand (Thai), Mexico (Mex) and Pakistan (PAK).

Definitions and prior knowledge:

- r: the mutated allele for the red eye colour
- r^+^: the wild type allele for the black eye colour
- r^+^ > r: the wild type r^+^ is dominant over the r. That means that r^+^/r^+^ and r^+^/r genotypes are wild type (black), while only r/r genotypes are red
- this locus is on chromosome 1 and very close to the sex determining M locus. Therefore:
- r m / r m = red eye females
- r^+^ m / r m = black eye females
- r m / r^+^ m = black eye females
- r^+^ m / r^+^ m = black eye females
- r M / r m = red eye males
- r^+^ M / r m = black eye males
- r M / r^+^ m = black eye males
- r^+^ M / r^+^ m = black eye males
- The recombination rate between the sex locus (M/m) and the eye-color locus are 2-3%

Experimental approach:

1. Cross rm/rm (red eye strain) x r^+^M/r^+^m (wild type BRA strain) (P). F1 progeny are all wild type and with 50% BRA genomic background.
2. Backcross F1 females with ‘BRA males. F2 progeny are all wild type and with 75% BRA genomic background.
3. Inbreed F2 males and females. F3 progeny have mainly black eyes but some are expected to have the red eye phenotype (mainly females). At the same time, they are considered to have 75% BRA genomic background.
4. Backcross F3 red eye females with ‘BRA males. F4 progeny are all wild type and with 87.5 % BRA genomic background.
5. Backcross F4 females with ‘BRA’ males. F5 progeny are all wild type and with 93.625% BRA genomic background.
6. Inbreed F5 males and females. F6 progeny have mainly black eyes but some are expected to have the red eye phenotype (mainly females). At the same time, they are considered to have 93.625 % BRA genomic background.
7. Backcross F6 red eye females with ‘BRA males. F7 progeny are all wild type and with 96.8% BRA genomic background.
8. Backcross F7 females with ‘BRA’ males. F8 progeny are all wild type and with 98.44% BRA genomic background.
9. Inbreed F8 males and females. F9 progeny have mainly black eyes but some are expected to have the red eye phenotype (mainly females). At the same time, they are considered to have 98.44 % BRA genomic background.
10. Two crosses are performed here:
a. Backcross F9 red eye females with ‘BRA males. F10a progeny are all wild type and with 99.21 % BRA genomic background.
b. At the same time, inbreed F9 red eye females and red eye males (red eye males are very rare). This red eye line (F10b) can be considered as having 98.44% BRA genomic background.
11. Cross F10b red eye females with F10a wild type males. This is a GSS that is considered to have a BRA genomic background of 98.82 %.

Note that the percentages shown in the protocol are calculated based random admixture. However, the extensive recombination desert will likely impede the effectiveness of this process.

**Figure S1.** Mapping the available genetic crosses data (Juneja et al. 2014) to the most recent AaegyL5 PacBio-based assembly (Matthews et al. 2018) shows little or no recombination in a ~110 Mb region surrounding the sex locus, which is at ~152 Mb on the homomorphic sex chromosome 1 in *Aedes aegypti*. This low recombination regions (LRRs) in *Ae. aegypti* in all three chromosomes were previously determined by mapping to an earlier version of the assembly ((Dudchenko et al. 2017); (Fontaine et al. 2017), GBE, 2017). Although we only showed the sex-determining chromosome 1, the other two chromosomes have similar patterns of vast regions of suppressed recombination.

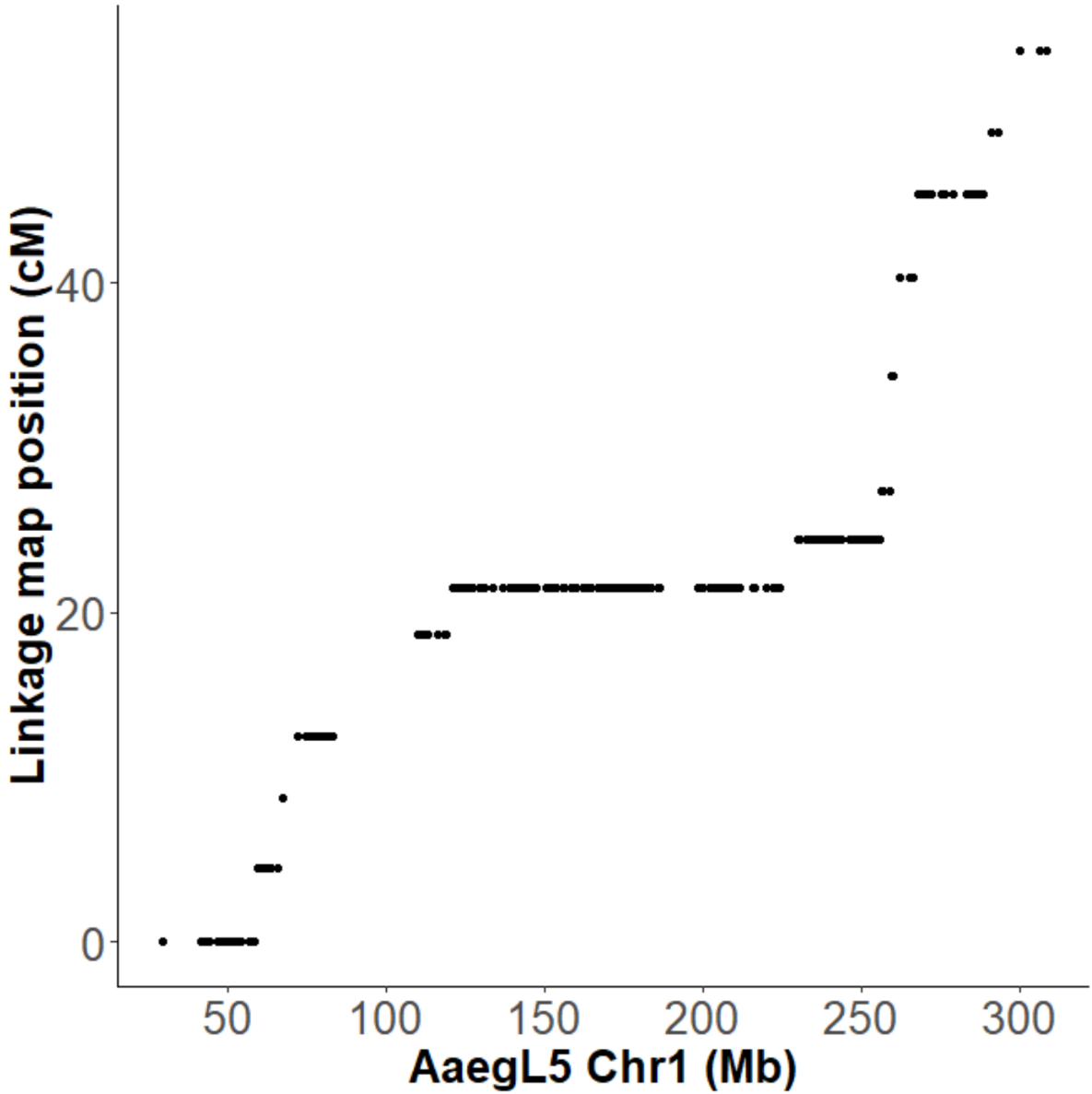

**Figure S2.** The donor plasmid used to obtain the P10 transgene insertion line. The piggyBac arms are not shown. PolyUb is a ubiquitous promoter (Anderson et al., 2010) that drive the expression of GFP. SV40 (yellow) polyadenylation signal is included in addition to an attB site.

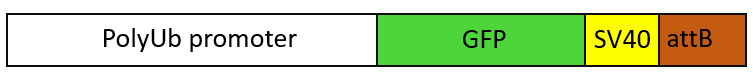

**Figure S3. An alignment between the sequences flanking the piggyBac-mediated insertion of one of the 4 ONT reads and the AegyL5 genome** The TTAA target site duplication caused by the piggyBac (P10) insertion are highlighted in blue. The precise location of the insertion is **224,678,447 on chromosome 1**. Note that single molecule sequencing (ONT) reads show >5% error rate, which is reflected in the less than 100% identity in the alignment.

```
Query  730        TTGCTCAACTGGCTGTACAGAAT-GCTTTACAGAAAAATTTAAATGTCTTATTTAATAAG  788
                  |||||||||||||||||| ||||  |||||||||||||||||||||||||||||||||||
Sbjct  224678891  TTGCTCAACTGGCTGTAC-GAATAACTTTACAGAAAAATTTAAATGTCTTATTTAATAAG  224678833

Query  789        TAAACGTTCAGTTAAGTATCAACGATTTTCGTTTGTTTTATATTAAGCACAACTCGTTTG  848
                  ||||||||||||||||||||||||||||||||||||||||||||  ||||||||||||||
Sbjct  224678832  TAAACGTTCAGTTAAGTATCAACGATTTTCGTTTGTTTTATATT-GGCACAACTCGTTTG  224678774

Query  849        GATGTTTT-CACCTGACTTTAAGTGTTATTTCCTGACTTTCCAAGTCAAA--TGGGATT-  904
                  |||||||| |||||||||||||||||||||||||||||||||||||||||  |||||||
Sbjct  224678773  GATGTTTTTCACCTGACTTTAAGTGTTATTTCCTGACTTTCCAAGTCAAAAATGGGATTC  224678714

Query  905        CCTGACGTTTCCAGTTTTT-CTTCTTTTTCGGTTAGTCGACATTCTAAAACAGCCTTGAG  963
                  ||||||||||||||||||| |||| |||||||||||||||||||||||||| ||||||||
Sbjct  224678713  CCTGACGTTTCCAGTTTTTTCTTC-TTTTCGGTTAGTCGACATTCTAAAACGGCCTTGAG  224678655

Query  964        AAATGCAAATTTGTTTT-CAAGCT-AAAACTTCACATATGAGTGGTTGTTCAAGATTGGA  1021
                  ||||||||||||||||| |||||| |||||||||||||||||||||||||||||||||||
Sbjct  224678654  AAATGCAAATTTGTTTTTCAAGCTAAAAACTTCACATATGAGTGGTTGTTCAAGATTGGA  224678595

Query  1022       GAAACTCTTTACCCTAACAGGGGGTGACCTTTAAGCTACCCTTTTGCACTGGTGCCTAAC  1081
                  |||||||||||||||||||||||||||||||||||||||||||||||| |||||||||||
Sbjct  224678594  GAAACTCTTTACCCTAACAGGGGGTGACCTTTAAGCTACCCTTTTGCA-TGGTGCCTAAC  224678536

Query  1082       TCATTAAGATCTCGTCAAACTAGTGGTTCCGCTATTACCTTTTCAAAAGATTTCGACAGC  1141
                  ||||||||||||||||||||||||||||||||||||||||||||||| ||||||||||||
Sbjct  224678535  TCATTAAGATCTCGTCAAACTAGTGGTTCCGCTATTACCTTTTCAAA-GATTTCGACAGC  224678477

Query  1142       CTAAGCAAGGGACAAGTTTTGCGACCAACTGCCTACCTCAAAGAGACAACATTTCAATGT  1201
                  ||||||||||||||||||||||||||||||||||||||||||||||||||||||||||||
Sbjct  224678476  CTAAGCAAGGGACAAGTTTTGCGACCAACTGCCTACCTCAAAGAGACAACATTTCAATGT  224678417

Query  1202       TGACTGACTAGCTACTCTTGTGATCGCAGTTCTTAA_P10_TTAAGCTGCATAACGCCGACCGGT  1261
                  ||||||||| ||||||||||||||||||||||||||         ||||||||||||||||||||
Sbjct  224678416  TGACTGACTGGCTACTCTTGTGATCGCAGTTCTTAA     ----GCTGCATAACGCCGACCGGT  224678361

Query  1262       TTTGTGGATTGCATTCTTGCAACTCTATCAGTCGGATTAACCATATATAGGTACCTGCCT  1321
                  || |||||||||||||||||||||||||| |||||||||||||||||  |||||||||||
Sbjct  224678360  TT-GTGGATTGCATTCTTGCAACTCTATCGGTCGGATTAACCATATA--GGTACCTGCCT  224678304

Query  1322       ACTTCACGGATACCTATAGCACGAAA-GTTCAAGACGGCTGGCAACAACGACCGAACAAC  1380
                  |||||||||||||||||||||||||| |||||||||||||||||||||||||||||||||
Sbjct  224678303  ACTTCACGGATACCTATAGCACGAAAAGTTCAAGACGGCTGGCAACAACGACCGAACAAC  224678244

Query  1381       GAAGCACTCTTTGAGACACTATGATAGTTTGTTACAGAGGGTTCATTCTCCAAACCAGTT  1440
                  ||||| |||||||||||||||||||||||||||||||||| |||||||||||||||||||
Sbjct  224678243  GAAGC-CTCTTTGAGACACTATGATAGTTTGTTACAGAGG-TTCATTCTCCAAACCAGTT  224678186

Query  1441       CATTCGTCGGTTCCTTATGGATTTGATCGTTTGGGTTTAAACCAGTTTGAATTATGCCGC  1500
                  |||||||||||||  |||||||||||||||||||||||||||||||||||||||||||
Sbjct  224678185  CATTCGTCGGTTC-ATATGGATTTGATCGTTTGGGTTTAAACCAGTTTGAATTATGCC-A  224678128

Query  1501       TAACTATCGAGATTGTGGTAGACTAGGCAGATGATAATTTGCATAGATATTTT--GGGGC  1558
                  |||||||||||||||||||||||||||||||||||||||||||||||||||||  |||||
Sbjct  224678127  TAACTATCGAGATTGTGGTAGACTAGGCAGATGATAATTTGCATAGATATTTTTGGGGGC  224678068

Query  1559       TTACCAATCAGCTTGCG--GGTGCCTATAAATCC-ATTTTATGGCGATA-GATGCGGATG  1614
                  |||||||||||||||||  ||||||||||||||| ||||||| |||||| ||| ||||||
Sbjct  224678067  TTACCAATCAGCTTGCGCGGGTGCCTATAAATCCCATTTTATAGCGATAAGATACGGATG  224678008

Query  1615       CCTTTCAATGTGTACTCCAGCAGAAATAGCTAATCTTTGCGTCCACCGAAATCAAATGCA  1674
                  ||||||||||||||||| | ||||||||||||||||||||||||||||||||||||||||
Sbjct  224678007  CCTTTCAATGTGTACTC-AACAGAAATAGCTAATCTTTGCGTCCACCGAAATCAAATGCA  224677949

Query  1675       CAAAA-CAAGCTTATCACATATTTGCGCTGGTGGCTGGG-TAATAAATTACAAATG---G  1729
                  ||||| ||||||||||||||||||||||||||||||||| ||||||||||||||||
Sbjct  224677948  CAAAAACAAGCTTATCACATATTTGCGCTGGTGGCTGGGGTAATAAATTACAAATGAAAC  224677889

Query  1730       CTTACCGTTCTTCGGTAATC-CTTGACAATGTTCACCACAACATTGAAGCGATCCTTTT-  1787
                  |||||||||||||||||||| |   ||||||||||||||||||||||||||||||||||
Sbjct  224677888  CTTACCGTTCTTCGGTAATCGCCAAACAATGTTCACCACAACATTGAAGCGATCCTTTTT  224677829

Query  1788       GGACATGGTTCGGTTCTGTTCTTCTTCGAGAGGATGTCGTTTTACTGATATTGTTT-ATC  1846
                  |||||||||||||||||||||||||||||||||||||||||||||||||||||||| |||
Sbjct  224677828  GGACATGGTTCGGTTCTGTTCTTCTTCGAGAGGATGTCGTTTTACTGATATTGTTTTATC  224677769

Query  1847       ATTCGTTAAGCATACCAACCAAAAAGAATGATAAGCTACTGTCTAGCCGTTTCTTAACAA  1906
                  ||||||||||||||||||||||   ||||||||||||||||||| |||||||||||||||
Sbjct  224677768  ATTCGTTAAGCATACCAACCAA---GAATGATAAGCTACTGTCTGGCCGTTTCTTAACAA  224677712

Query  1907       TGATGTGAAAAAATCAGTGGCCCGCGAAGGGGATAGGATAGGTAGAACAAGTCATACGCA  1966
                  ||||||||||||||||||||||||||||||||||||||||||||||||||||||||||||
Sbjct  224677711  TGATGTGAAAAAATCAGTGGCCCGCGAAGGGGATAGGATAGGTAGAACAAGTCATACGCA  224677652

Query  1967       CACAGCTACCTGGGTAGGCCAGCAAAACGATTGTCCTACCCATGATACTACAAAAACATA  2026
                  ||||||||||||||||||||||||||||||||||||||||||||||||||||||||||||
Sbjct  224677651  CACAGCTACCTGGGTAGGCCAGCAAAACGATTGTCCTACCCATGATACTACAAAAACATA  224677592

Query  2027       AAAACAATCTCTTTGCCTTTTAAGATTAAGGTCGTATTTCTCAATGAACATACGACAAAA  2086
                  ||||||||||        ||||||||||||||||||||||||||||||||||||||||||
Sbjct  224677591  AAAACAATCT--------TTTAAGATTAAGGTCGTATTTCTCAATGAACATACGACAAAA  224677540

Query  2087       GTCTGGA-AAACCACTTT-A-AATTCTTGCAATAATACAACGGAACATGCGTAATTTaaa  2143
                  ||||||| |||||||||| | |||||||||||||||||||||||||||||||||||||||
Sbjct  224677539  GTCTGGAGAAACCACTTTAAGAATTCTTGCAATAATACAACGGAACATGCGTAATTTAAA  224677480

Query  2144       aaaa--TAAGGAACGAACAGCATTCCAAA-GGTTCTCTAATTGATTTCATGTAAAAAATC  2200
                  ||||  ||||||||||||||||||||||| ||||||||||||||||||  ||||||||||
Sbjct  224677479  AAAAAATAAGGAACGAACAGCATTCCAAAAGGTTCTCTAATTGATTTC--GTAAAAAATC  224677422

Query  2201       CTAAGGAATTTCTACAGCCAGTTAGATAGATGTTCTGGAAAATAATTGAATCTAGGCATT  2260
                  ||||||||||||||||||||||||||||||||||||||||||||||||  ||||||||||
Sbjct  224677421  CTAAGGAATTTCTACAGCCAGTTAGATAGATGTTCTGGAAAATAATTG-GTCTAGGCATT  224677363

Query  2261       -CTGGTGGTATTTCTTGATAAATAACTGCATTATTCAAGGAAAAACTATTGGGAGATTCG  2319
                   ||||||||||||||||||||||||||||||||||||||||||| |||||||||||||||
Sbjct  224677362  CCTGGTGGTATTTCTTGATAAATAACTGCATTATTCAAGGAAAA-CTATTGGGAGATTCG  224677304

Query  2320       TAAAGAATCATTGGAAGAATTAATAGATTCACCCAAGACTGCAGTCTTGTAACATATATG  2379
                  ||||||||| ||||||||||||||||||||||||||||||||||||||||||||||||||
Sbjct  224677303  TAAAGAATCGTTGGAAGAATTAATAGATTCACCCAAGACTGCAGTCTTGTAACATATATG  224677244

Query  2380       TTACACACTGATGGCTCCCTTCTTGA---TAGAGCTCGTGCAGGAGTAACTATTCTCGTG  2436
                  ||||||  ||||||||||||||||||   |||||||||||||||||||  ||||||||||
Sbjct  224677243  TTACAC--TGATGGCTCCCTTCTTGAAGGTAGAGCTCGTGCAGGAGTA--TATTCTCGTG  224677188

Query  2437       AGCTAAAGGCTGAATCAGTTTTACTC-CTTGGTAGAAAC-ACACCGTTTTTCAGGCT-AA  2493
                  |||||| ||||||||||||||||||| ||||||||||||  |||||||||||||||| ||
Sbjct  224677187  AGCTAA-GGCTGAATCAGTTTTACTCACTTGGTAGAAACTGCACCGTTTTTCAGGCTGAA  224677129

Query  2494       ACATTTGCTCTTATGTG--GAATGCAATCAGCACTTCAACAGCAC--ATGTGGTAAAGTC  2549
                  |||||||||||||||||  ||||||||||||||||||||||||||  |  ||||||||||
Sbjct  224677128  ACATTTGCTCTTATGTGTGGAATGCAATCAGCACTTCAACAGCACGTAATTGGTAAAGTC  224677069

Query  2550       ATATACTTCTGTTCAGATAGTCCAGGCTGCTATAAAAGCTCTTGCTTCCCAGCCAACTCA  2609
                  |||||||||||||||||||||| |||||||||||||||||||||||||   |||||||||
Sbjct  224677068  ATATACTTCTGTTCAGATAGTC-AGGCTGCTATAAAAGCTCTTGCTTC--GGCCAACTCA  224677012

Query  2610       AGGTC-AAGCTTGTTGTCTTTATGTCGAACTCAAATTGAGGAACTAAATTCAGTCACTCA  2668
                  ||||| ||||||||||||   |||||||||||||||||||||||||||||||||||||||
Sbjct  224677011  AGGTCGAAGCTTGTTGTC-GCATGTCGAACTCAAATTGAGGAACTAAATTCAGTCACTCA  224676953

Query  2669       GGACATTCTGTT-GTG-TAA-CCGCTGCGGTGATAAGCTATAGGCTCTCTTTACGCTCAT  2725
                  ||||||||||||  || ||| |||||||||||||||||||||||||||||||||||||||
Sbjct  224676952  GGACATTCTGTTATTGCTAACCCGCTGCGGTGATAAGCTATAGGCTCTCTTTACGCTCAT  224676893

Query  2726       GCGTTTT-CGTTATTTGAACCCATTGGGGTATGGAGCTAAATACTCTTTTT-C-CTTATG  2782
                  ||||||| |||| |||||||||||||||||||||||||||||||||||||| | ||||||
Sbjct  224676892  GCGTTTTTCGTTGTTTGAACCCATTGGGGTATGGAGCTAAATACTCTTTTTTCGCTTATG  224676833
```

**Figure S4. F1 progeny from the Rexville female x RED male cross (A and B) and from the RED female x Rexville male cross (C and D).**

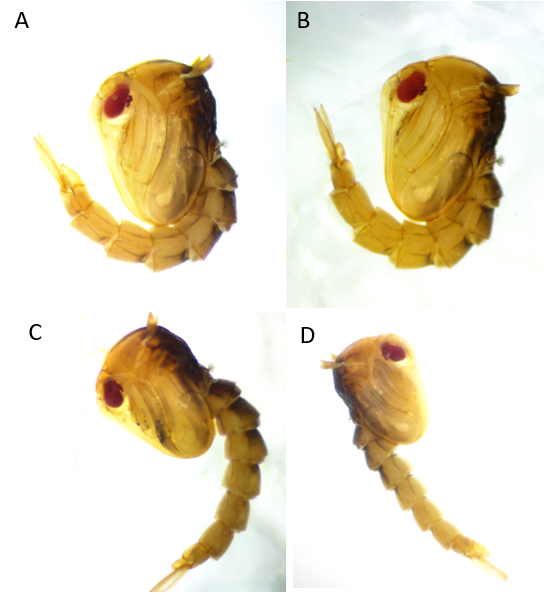

**Figure S5.** Sequence alignment of the Cardinal proteins (Car) from various mosquito species. Cq, *Culex quniquefasciatus*; Aalbo, *Aedes albopictus*; MRP, red-eyed males from the RED strain; Aaegy, *Aedes aegypti* wildtype; Aalbi, *Anopheles albimanus*; Astep, *Anopheles stephensi*; Agam, *Anopheles gambiae*.

```
Cq_car
Aalbo_Car
MRP_re_Illumina      -----------------------------------------------------MTDERTP	7
Aaegy_Car            -----------------------------------------------------MTDERTP	7
Aalbi_Car            ----------------------------------------MFFSASFVTDTAAMADERTP	20
Astep_Car            ------------------MVAI--ELSV----NRKVPTPFLLSRVVLLLLSMVMVDERTP	36
Agam_Car             ---------------------------------------------------MVMVDERTP	9
                                                                          *.*****

CLUSTAL O(1.2.4) multiple sequence alignment
Cq_car               -----------------------------------------------------MTDERTP	7
Aalbo_Car            MMGNVGYRMSYMRLPPRTILSIISSLAVSHLVVVERPGKSHTVLVLHEDLVPEMTDERTP	60
MRP_re_Illumina      -----------------------------------------------------MTDERTP	7
Aaegy_Car            -----------------------------------------------------MTDERTP	7
Aalbi_Car            ----------------------------------------MFFSASFVTDTAAMADERTP	20
Astep_Car            ------------------MVAI--ELSV----NRKVPTPFLLSRVVLLLLSMVMVDERTP	36
Agam_Car             ---------------------------------------------------MVMVDERTP	9
                                                                          *.*****

Cq_car               LTLGPPPPLDTD---SGSSYVHQHKSYETLRERQVKTFQCWICSAILGAFVIGIIISLSY	64
Aalbo_Car            LTSVPPSALDSE----SSSYVHQLKGHETLRERQVKNFQCWICTAIFGAFIIGIIISLSY	116
MRP_re_Illumina      LTSVPPSALDSE----SSSYVHQLKGHETLRERQVKNFQCWICTAILGAFVIGIIISLSY	63
Aaegy_Car            LTSVPPSALDSE----SSSYVHQLKGHETLRERQVKNFQCWICTAILGAFVIGIIISLSY	63
Aalbi_Car            LTTVPPSTPPFSAGPSSAGTVHHLKPHESLRERQVRTFQCWICSAILGAFVLAIVICISY	80
Astep_Car            LTSDLSGPLPLVSGPSG-N-VHHLKSHESVRERQVRTFQCWICSAIMGAFALAIVISISY	94
Agam_Car             LTSDLSGPLPLASGPSG-TAVHHLKSHESVRERQVRTFQCWICSAILGAFALAIVISISY	68
                     **              .   **: * :*::*****:.******:**:*** :.*:*.:**

Cq_car               IILGDARKGAANVTNTTESANFPDLFNLISFPLADESIPTWDGPDVSDEDAAAAISEGEK	124
Aalbo_Car            VIMGDAKRGASNSTNVTETANFPDLLNLISFPLADEPTPMWDAPNITEQDAADAIAEGEK	176
MRP_re_Illumina      IIMGDAKRGPSNSTNVTETANFPDLLNLISFPLADEPTPMWDAPNITEQDAADAIAEGEK	123
Aaegy_Car            IIMGDAKRGPSNSTNVTKTANFPDLLNLISFPLADEPTPIWDAPNITEQDAADAIAEGEK	123
Aalbi_Car            IIFGDATISPLDGSNT-TAADFPELLNLISFPLADEPVPEWNGTDIGEDEKMAAVAEGEK	139
Astep_Car            IIFGDATKPPLDGANA-TAADFPELLNLISFPLVDELPPQWNGTDVSEDAKAAAIAEGEK	153
Agam_Car             IIFGDATQPPLDGANT-TAADFPELLNLISFPLVDESSPEWNGTAVSDDAKAAAIADGEK	127
                     :*:***     : :*.  :*:**:*:*******.**  * *:.  : ::    *:::***

Cq_car               ALGDKELLEEALTAPPVNSPTYRHQRAVSTSLAARLVSKLGYVENKATKAIARKFNMNKR	184
Aalbo_Car            ALGDKELLEEALTAPPVQSATFRHQRAVSTTLAARLASKVGYVENKATNAIARKHNMKKR	236
MRP_re_Illumina      ALGDKELLEEALTAPPVQSATFRHQKAVSTTLAARLASKVGYVENKATNAIARKHNMKKR	183
Aaegy_Car            ALGDKELLEEALTAPPVQSATFRHQKAVSTTIAARLASKVGYVENKATNAIARKHNMKKR	183
Aalbi_Car            ALGDKELLEETLAPPPVNSPSFRHQKSVGATMAARLAAKVGFVEDRATKALVRKFGSIRR	199
Astep_Car            ALGDKELLEETLSSPPVNSPSFRHQKSVGATVAARLAAKVGFVEDRATKALVRKLDIRH-	212
Agam_Car             ALGDKELLEETLSSPPLNSPSFRHQKSVGATKAARLAAKVGFVEDRATQALVRRLDIRR-	186
                     **********:*: **::* ::***::*.:: ****.:*:*:**::**:*:.*: .  :

Cq_car               HRRWSIGKGLALKLPRVNETQVCDFNARYRTNNGTCNNRKQPYSYGVALIPFRRQLTPDY	244
Aalbo_Car            HRRWSIGKGPAIKMPHQNGTQQCDFNARYRTNNGTCNNKKNPHTYGVALIPFRRQLTPDY	296
MRP_re_Illumina      HRRWSIGKGPAIKVPHQNGTQQCDFNASYRNNNGTCNNKKNPHTYGVALIPFRRQLTPDY	243
Aaegy_Car            HRRWSIGKGPAIKVPHQNGTQQCDFNARYRTNNGTCNNKKNPHTYGVALIPFRRQLTPDY	243  deleted in the G1 male #3
Aalbi_Car            -QGKSIGRGPVINVPRVQHETQCDFNARYRTPNGTCNNKEHPFEYGVAMVPFRRQLNPDY	258
Astep_Car            --RGSVGRGPVMNLPRTHRHPQCDFNARYRSANGTCNNKERPYEYGVAMIPFRRQLNPDY	270
Agam_Car             --RGSIGRGPPMDLPRAHRQPRCDFNARYRTANGTCNSKERPYEYGVAMIPFRRQLNPDY	244
                         *:*:*  :.:*: :    ***** **. *****.::.*. ****::******.***

Cq_car               GDGVSSPRESVDRKALPSARQVSIELHRPSYHNDPNFSVMLAVWGQFLDHDITSTALNQG	304
Aalbo_Car            GDGVSSPRESVEGKELPSARQVSLQIHRPSYHNDPNFSVMLAVWGQFLDHDITSTALNQG	356
MRP_re_Illumina      GDGVSSPRESIEGKELPSARQVSLEIHRPSYHNDPNFSVMLAVWGQFLDHDITSTALNQG	303
Aaegy_Car            GDGVSSPRESIEGKELPSARQVSLQIHRPSYHNDPNFSVMLAVWGQFLDHDITSTALNQG	303
Aalbi_Car            GDGVSAPRVAIDGSELPSARHVSLDIHRPSYHNDPNFSVMLAVWGQFLDHDITSTALNQG	318
Astep_Car            GDGISAPRASVDGTELPSARQVSLDIHRPSYHSDPNFSVMLAVWGQFLDHDITSTALNQG	330
Agam_Car             GDGISAPRASVDGAELPSARQVSLEIHRPSYHNDPNFSVMLAVWGQFLDHDITSTALNQG	304
                     ***:*:** :::   *****:**:::******.***************************

Cq_car               VGGKAIECCDPGQPQHPECYPVKLGPGDPYFHEYNLTCMNFVRSIPASTGHLGPRQQLNQ	364
Aalbo_Car            VGGKAIECCDPGQPRHPECFPVPLGPGDPYFHDYNLTCMNFVRSIPAPTGHFGPRQQLNQ	416
MRP_re_Illumina      VGGKAIECCDPGQPRHPECFPVPLGPGDPYFHDYNLTCMNFVRSIPAPTGHFGPRQQLNQ	363
Aaegy_Car            VGGKAIECCDPGQPRHPECFPVPLGPGDPYFHDYNLTCMNFVRSIPAPTGHFGPRQQLNQ	363
Aalbi_Car            VNGKPIECCDPGQPQHPECFPVPIGPGDPYFHQYNVTCMNFVRSVPAPTGRFGARQQLNQ	378
Astep_Car            VDGKPIECCDPGQPQHPECFPVPLGPGDPYYHQYNVTCMNFVRSVPAPTGHFGPRQQLNQ	390
Agam_Car             VDGKPIECCDPGQPQHPECFPVPLGPGDPYYTQYNVTCMNFVRSVPAPTGHFGPRQQLNQ	364
                     *.** *********:****:** :******: :**:********:** **::* ******

Cq_car               ATAYIDGSVVYGSDDAKVKRLRSGIDGRLRMLTTPDNRELLPQSTDPNDGCNEASMNAAG	424
Aalbo_Car            ATAYIDGSVVYGSDDAKVKRLRTGQDGKLRMYVTPDNRELLPISTDPNDGCNEEAMNAAG	476
MRP_re_Illumina      ATAYIDGSVVYGSDDAKVKRLRSGKDAKLRMYVTPDNRELLPISTDPNDGCNEEAMNAVG	423
Aaegy_Car            ATAYIDGSVVYGSDDAKVKRLRSGKDGKLRMYVTPDNRELLPISTDPNDGCNEEAMNAVG	423
Aalbi_Car            ATAFIDGSVVYGSDEALMRSLRSGEGGRLRMLRTPDGRELLPVSTDPEDGCNEAEMNAAG	438
Astep_Car            ATAYIDGSVVYGSDEERMKKLRTGEGGRLRMLRTPDGRQLLPVSTDPLDGCNEQEMNAAG	450
Agam_Car             ATAFIDGSVVYGSDDERMGALRTGAGGQLRMLRTPDGRDLLPVSTDPLDGCNEQEMNAAG	424
                     ***:**********:  :  **:* ..:***  ***.*:*** **** *****  ***.*

Cq_car               KYCFESGDDRSNENLHLTSMHLIWARHHNNLTGELKKVNPEWDDERLFQEARRILAAQMQ	484
Aalbo_Car            KYCFESGDERANENLHLTSMHLIWARHHNNLTGELKKVNPDWDDERLFQEARRILAAQMQ	536
MRP_re_Illumina      KYCFESGDERANENLHLTSMHLIWARHHNNLTGELKKVNPDWDDERLFQEARRILAAQMQ	483
Aaegy_Car            KYCFESGDERANENLHLTSMHLIWARHHNNLTGELKKVNPDWDDERLFQEARRILAAQMQ	483
Aalbi_Car            KYCFESGDSRANENLHLTSMHLIWARHHNSLADGLAKVNPGWNDERLFQEARRILAAQMQ	498
Astep_Car            KYCFESGDTRANENLHLTSMHLIWARHHNSLADGLARVNPHWDDERLFQEARRILAAQMQ	510
Agam_Car             KYCFESGDARANENLHLTSMHLIWARHHNSLARGLARANPHWDDERLFQEARRILAAQMQ	484
                     ******** *:******************.*:  * :.** *:*****************

Cq_car               HITYSEFVPVIIGANNSDQMGISPTPDSD---RDTYNASVDASIANIFAAAAFRFAHTLL	541
Aalbo_Car            HITYGEFVPVIVGEDTAERMELAPNPESD---RDTYNVSVDPSVANVFAASAFRFAHTLL	593
MRP_re_Illumina      HITYGEFVPVIIGEDTAERMEISPNPESD---RDTYNVTVDPSVANVFAASAFRFAHTLL	540
Aaegy_Car            HITYGEFVPVIIGEDTAERMEISPNPESD---RDTYNVTVDPSVANVFAASAFRFAHTLL	540
Aalbi_Car            HITYSEFVPVIVGNETARRMGILPDPESG---RDTYNSTVDASIANVFAGAAFRFAHTLL	555
Astep_Car            HITYAEFVPVIVGNATAARMDLLPESTGR---DDTYNASVDASIANVFAGAAFRFAHTLL	567
Agam_Car             HITYAEFVPVIVGNETAGRMGLLPVSAGGEPAGDTYNATVDASIANVFAGAAFRFAHTLL	544
                     ****.******:*  .: :* : *   .     **** :** *:**:**.:*********

Cq_car               PTLMKQTRDPTSSASGIELHKMLFNPYSLYGSTGLDDAIGGAMSTPLGKYDQFFTTELTE	601
Aalbo_Car            PGLMKKTHDPTSSPSGIELHKMLFNPYSLYGKTGLDDAIGGAMTTPLGKYDQYFTTELTE	653
MRP_re_Illumina      PGLMKRTHDPTSSPSGIELHKMLFNPYSLYGKTGLDDAIGGAMSTSLGKYDQYFTTELTE	600
Aaegy_Car            PGLMKRTHDPTSSPSGIELHKMLFNPYSLYGKTGLDDAIGGAMSTPLGKYDQYFTTELTE	600
Aalbi_Car            PGLMKKTRNPAASSSGIELHKMLFNPYSLYAATGLDDALGGAISTALAKYDQYFSTELTE	615
Astep_Car            PGLMKKTRNPTSSSSGIELHRMLFNPYSLYAHDGLDNALGGAMSTSLAKYDQYFSTELTE	627
Agam_Car             PGLMKQTRNPAASASGIELHRMLFNPYSLYARDGLDNALGGAIGTALAKYDQYFSTELTE	604
                     * ***:*::*::* ******:*********.  ***:*:***: * *.****:*:*****

Cq_car               RLFEKSE-DLLHDRPCGLDLVSLNIQRGRDHGLPSYPHWRKHCRLPPVDTWAQMADAVDP	660
Aalbo_Car            HLFEKAQ-DLLHDRPCGLDLVSLNIQRGRDHGLPSYPHWRRHCRLPPVDTWDQLEKVVDA	712
MRP_re_Illumina      HLFEKAQ-DLLHDRPCGLDLVSLNIQRGRDHGLPSYPHWRRHCRLPPVDTWDQLEKVVDP	659
Aaegy_Car            HLFEKAQ-DLLHDRPCGLDLVSLNIQRGRDHGLPSYPHWRRHCRLPPVDTWEQLEKVVDP	659
Aalbi_Car            RLFEKADEHLLHNHPCGLDLVSLNIQRGRDHGLPAYPHWRRHCHLTPADTWDQLERIVDP	675
Astep_Car            KLFEKADEHLLHNHPCGLDLVSLNIQRGRDHGLPAYPRWRKHCHLTPADSWAELERIVDP	687
Agam_Car             RLFEKADEHLLHGQPCGLDLVSLNIQRGRDHGLPAYPRWRKHCHLTPADSWEELERIVDP	664
                     :****:: .***.:********************:**:**:**:* *.*:* ::   **

Cq_car               GSLEQMKKMYAEPENVDVYSGALSEPPVKGGVVGPLITCLLGDQFVRLKQGDSFWYERRR	720
Aalbo_Car            GSYQQMRKIYGEPENVDVYSGALSEPPVEGGVVGPLITCLLADQFLRLKQGDSFWYERRR	772
MRP_re_Illumina      GSYEQMRKIYGEPDNVDVYSGALSEPPVEGGVVGPLITCLLADQFLRLKQGDSFWYERRR	719
Aaegy_Car            GSYEQMRKIYGEPDNVDVYSGALSEPPVEGGVVGPLITCLLADQFLRLKQGDSFWYERRR	719
Aalbi_Car            ASFQQMKTIYHNPVNVDVYSGALSEPPVNGGIVGPLLTCLLADQFLRLKQGDSFWYERRQ	735
Astep_Car            ESFRQMKSIYRDPANVDVYSGALSEPPVKDGIVGPLLTCLLADQFLRLKQGDSFWYERRR	747
Agam_Car             ESYRQMRRIYREPANVDVYSGALSEAPVRDGIVGPLLTCLIGDQFLRLKQGDSFWYERRR	724
                      * .**: :* :* *********** **..*:****:***:.***:*************:

Cq_car               GPQRFTRDQLQQIYNTKLSSIICRNSDAITHSPVDLMRKVDRNQNPERPCSELDTFDFGA	780
Aalbo_Car            GPQRFTRDQLRQIYNTRLSSIICRNSDAITQSPVYLMRKVNREDNPELPCSQLDTFDFSV	832
MRP_re_Illumina      GPQRFTRDQLRQIYNTRLSSIICRNSDAITQSPVYLMRKVNREDNPELPCSELDTFDFSV	779
Aaegy_Car            GPQRFTRDQLRQIYNTRLSSIICRNSDAITQSPVYLMRKVNREDNPELPCSELDTFDFSV	779
Aalbi_Car            GVQRFTEEQLQQIYETKLSSIICRNSDHIEQSPVYLMKKSDPITNPETNCKELDTFDFNA	795
Astep_Car            GPQRFTEGQLQQIYDTKLSSIICRNSDNIEQSPVHLMKRTDSRTNPETDCKQLDTFDFEP	807
Agam_Car             GPQRFTEAQLQQIYNTKLSSIICRNSDHIEQSPVYLMKRTDSRTNPETDCKQLDTFDFEP	784
                     * ****. **:***:*:********** * :*** **:: :   ***  *.:******

Cq_car               FRERKKDL---TGRVKVATAKVGVLVIQTSSTSTEASVGTSATEPSTTST---TGAATTT	834
Aalbo_Car            FRDRKNAD---SRRVKLATAQVGVVVQPSSSTVVSTNSSTA--D---TTT---EGLTTTT	881
MRP_re_Illumina      FRDRKNAD---SRRVKLATSQVGVVIQPGATTGMSDGMNSSGVE---TST---EVFGTST	830
Aaegy_Car            FRDRKNAD---SRRVKLATSQVGVVIQPGATTGMSDGMNSSGME---TST---EVLGTTT	830
Aalbi_Car            FHEDAHH--WQNKKSKLATDQMKVLVMEPKQQPAVVSFTTVATPESA------------V	841
Astep_Car            FREDKDAEPQHTRSAKIATDRVKVLVMEPKSAGTTTTRTTIEPEMVERDK---ATDSTTM	864
Agam_Car             FREDAE-QPQRNRAAKIATDRMKVLVFEPKATGTTTEHAV-GAEMQDVEEGERQAASTTT	842
                     *::  .     .   *:** :: *::

Cq_car               TTTA-SVPAT-----------------	843
Aalbo_Car            TTTSTSMTSD--------NTLI-----	895
MRP_re_Illumina      S----TTTAN--------NTIV-----	840
Aaegy_Car            S----TTTAN--------NTIV-----	840
Aalbi_Car            VTTATPEPTTSISSGESAETTEGANGA	868
Astep_Car            ITTTESLPTTLSNT---------VSGV	882
Agam_Car             VATTTDATTTTTTTTT---TMKEAAGA	866
```

**Figure S6.** The black-eyed and male mosaic-eyed siblings of the mosaic-eyed females shown in Figure 6B.

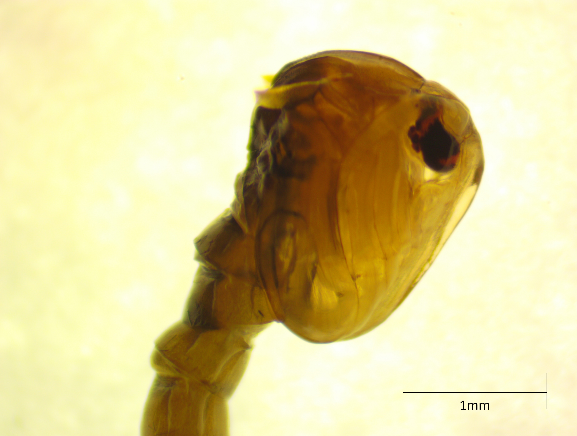

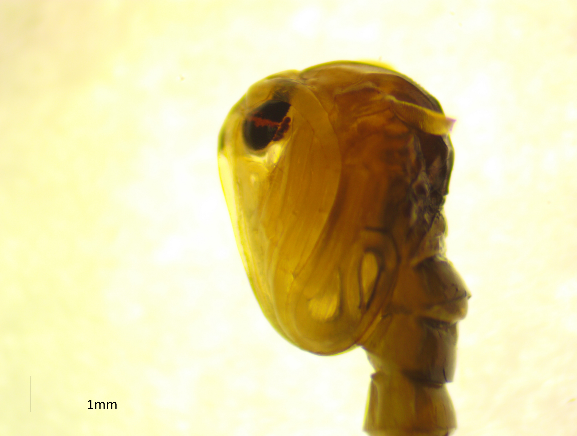

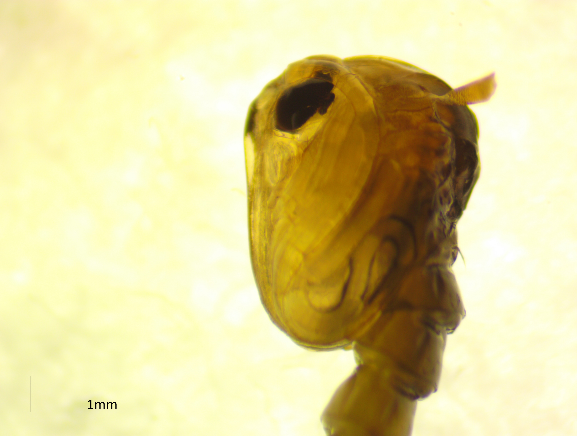

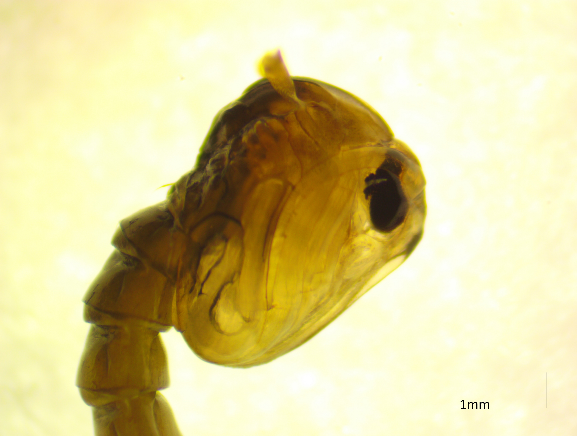

Mosaic ♂

Black-eye ♂

**Figure S7.** When black-eyed (+/cd-) G2 female mosquitoes crossed with RED (re/re) males, 1:1:1:1 ratio among four phenotypes is expected. The numbers are presented in Table S5.

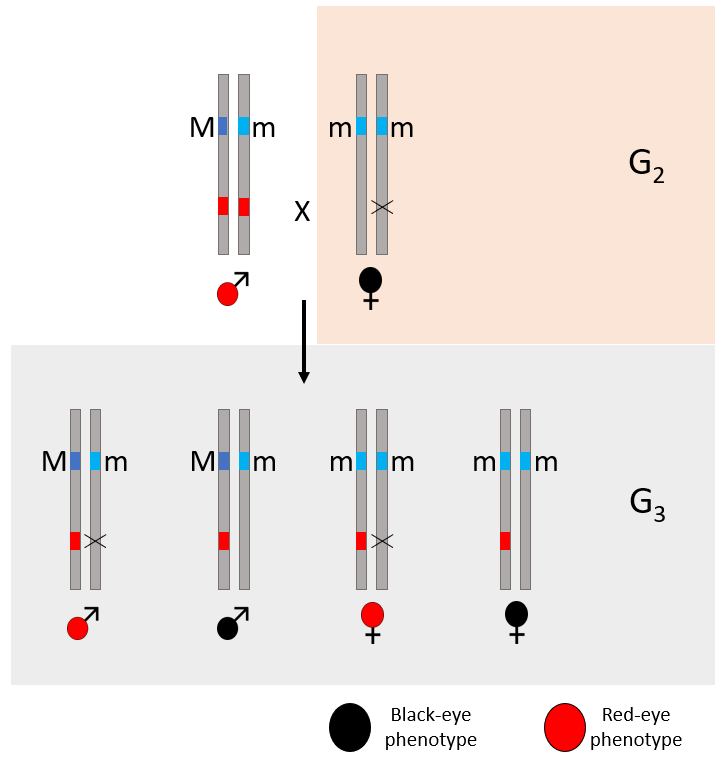

**Figure S8.** The yellow-eyed phenotype starts to appear in the compound eye in pre-pupal stage. From left to right, black-eyed control, yellow-eyed, and red-eyed G_3_ larvae siblings. The red ocelli adjacent to the yellow compound eye is circled.

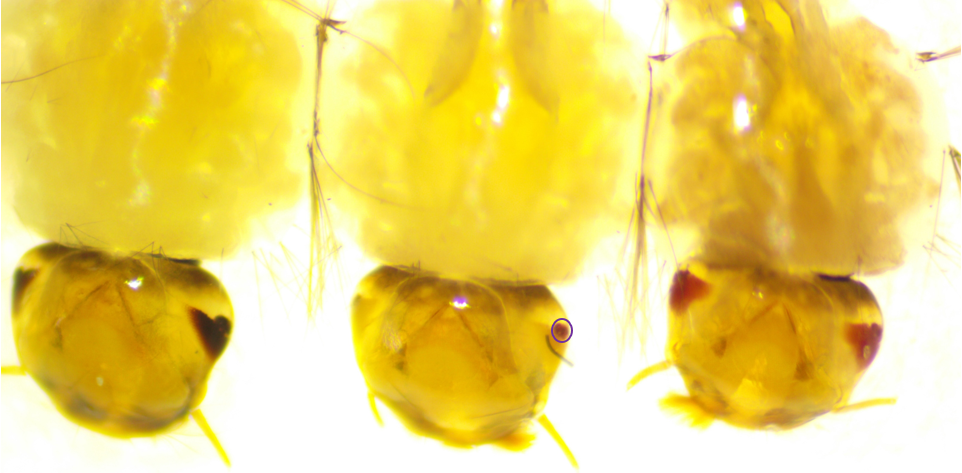

**Supplemental Table S1.** Mapping statistics of Illumina reads to the AaegyL5 genome assembly.

| **Sample name** | **Number of reads** | **Mapped rate** | **Estimate coverage (mapped)** |
| --- | --- | --- | --- |
| MPB | 501477681 | 99.55% | 62.66144089 |
| MPR | 516237494 | 99.52% | 64.48558045 |
| FNB | 511274734 | 99.57% | 63.89915887 |
| FNR | 305549189 | 99.59% | 38.19607736 |
| MNR | 469917491 | 98.06% | 57.83742803 |
| FPB | 495198337 | 97.89% | 60.84593701 |
| F7PAKF3BRA_F | 214553783 | 98.48% | 26.521593 |
| S2m | 228835353 | 98.03% | 28.15835454 |
| F8ThaiF9Mex_F | 224207234 | 97.11% | 27.32916263 |
| F8ThaiF9Mex_M | 223787419 | 98.44% | 27.65239873 |

Note that the coverage is calculated using the sum of the length of the three chromosomes.

**Supplemental Table S2.** The sgRNAs were designed to target specific regions within the predicted open reading frame of the LVP copy of *Cardinal*.

| Name | sgRNA synthesis primers with target sequence | Used in experiments |
| --- | --- | --- |
| sgRNA1_forward | GAAATTAATACGACTCACTATAGGTTTCCGGCGTCAACTAACTCGTTTTAGAGCTAGAAA | Somatic, Germline |
| sgRNA2_forward | GAAATTAATACGACTCACTATAGGCAATGCTACGCCATATGTGTGTTTTAGAGCTAGAAA | Somatic, Germline |
| sgRNA3_forward | GAAATTAATACGACTCACTATAGGCTAACTCCGGACTACGGTGAGTTTTAGAGCTAGAAA | Somatic |
| sgRNA4_forward | GAAATTAATACGACTCACTATAGGTTTTGCCGTACAGCGAGTACGTTTTAGAGCTAGAAA | Germline |
| sgRNA5_forward | GAAATTAATACGACTCACTATAGGCTGGTCGTATTTGCCGAGGGGTTTTAGAGCTAGAAA | Germline |
| sgRNA_common_reverse | AAAAGCACCGACTCGGTGCCACTTTTTCAAGTTGATAACGGACTAGCCTTATTTTAACTTGCTATTTCTAGCTCTAAAAC |  |

**Supplemental Table S3.** Germline Knockout Numbers

| G0 Embryos injected | ~300-360 |
| --- | --- |
| G0 Survival | 27 (12 male*, 15 female) |
| G1 red-eye ** | 11 (7 male, 4 female) |
| G1 Total ** | ~500 |

*We included sgRNAs targeting a M-locus gene *myo-sex* gene (Aryan et al., 2020) as a positive control as G_0_ males with myo-sex knocked out will be flightless. Indeed, six of the 12 G_0_ males are flightless. We focused our downstream analysis of cardinal knockout on G_0_ females as females do not have the M-locus.

** The G1 numbers are from G0 females mated with red-eyed (*re/re*) males.

**Supplemental Table S4.** Primers used for PCR amplification of sgRNA target regions and for Sanger validation

| Name | Amplification Region | Sequence | Orientation |
| --- | --- | --- | --- |
| Tu_2056 | Region 1 | 5’-AGAACGGTACCCAACAGTGC-3’ | Forward |
| Tu_2057 | Region 1 | 5’-TGACGGAGAAGTTAGGGTCG-3’ | Reverse |
| Tu_2063 | Region 2 | 5’-CCGGGCCTCATGAAGAGAAC-3’ | Forward |
| Tu_2064 | Region 2 | 5’-AGGAAACCAAATCCAGCCCG-3’ | Reverse |
| Tu_2065 | Region 2 | 5’-AAGAGAACCCACGACCCGA-3’ | Forward |
| Tu_2066 | Region 2 | 5’-GTCCGCGTTGGATGTTCAAG-3’ | Reverse |
| Tu_2071 | Region 1 | 5’-CAATGTGGGACGCTCCGAATA-3’ | Forward |
| Tu_2073 | Region 1 | 5’- GTGATGTCGTGATCTAAGAACT-3’ | Reverse |
| Tu_2075 | Region 2 | 5’- CCTCATGAAGAGAACCCACGA-3’ | Forward |
| Tu_2076 | Region 2 | 5’- ATCCGGGATCCACAACCTTCT-3’ | Reverse |

**Supplemental Table S5.** Inheritance of the stable germline knockout of *Cardinal*. Eye pigment phenotype counts are represented for the offspring of black-eye (+/cd-) G_2_ female mosquitoes crossed with RED (re/re) males. Each row shows the count of offspring from a single G_2_ female mother. *ne* indicates no eggs were produced. A 1:1:1:1 ratio is expected.

| Female | Red-eye male | Black-eye male | Red-eye female | Black-eye female |
| --- | --- | --- | --- | --- |
| 1.1 | 13 | 13 | 24 | 17 |
| 1.2 | *ne* | *ne* | *ne* | *ne* |
| 1.3 | 27 | 24 | 17 | 22 |
| 1.4 | 18 | 22 | 26 | 15 |
| 1.5 | 18 | 12 | 14 | 17 |
| 2.1 | 32 | 23 | 27 | 25 |
| 2.2 | 20 | 18 | 22 | 17 |
| 2.3 | 18 | 25 | 22 | 19 |
| 2.4 | *ne* | *ne* | *ne* | *ne* |
| 2.5 | 22 | 24 | 28 | 26 |
| 3.1 | 21 | 23 | 18 | 21 |
| 3.2 | 25 | 21 | 22 | 20 |
| 3.3 | 22 | 16 | 22 | 17 |
| 3.4 | 10 | 20 | 17 | 22 |
| 3.5 | 26 | 16 | 25 | 21 |
| Total | 285 | 270 | 308 | 276 |
| Percent Total (%) | 25.02 | 23.71 | 27.04 | 24.23 |

**Supplemental Table S6.** The numbers of the four phenotypes among the G_3_ siblings, examples of which are shown in Figure 7.

| Red-eyed male | Black-eyed male | Yellow-eyed female* | Black-eyed female |
| --- | --- | --- | --- |
| 17 | 10 | 17 | 11 |

* These yellow-eyed individuals turned red-eyed as they aged.

**Supplemental data: All are attached as separate files**

**1) The four Oxford Nanopore reads that contain the P10 transgene insertion**

**2) Trace files of Sanger sequencing**

**3) Original image files**
