## Supplementary material for "Marker-assisted mapping enables effective forward genetic analysis in the arboviral vector *Aedes aegypti*, a species with vast recombination deserts": Four ONT reads that contain P10

**The 4 ONT reads that contain the piggyBAC insertion**

>e677388e-9ab7-49f3-bf3f-d42041a58300

CTGTTGTGCTTCAGTTTCAGTTACGTATTGCTAGCACGCCTCACGGGGCTCAAGCGGCATTGGAGATGTCCTAAATGCACAGCGACGGATTCGCGCTATTTAGAAAAGAGAGGAAAAATATTTCAGAATGCATGCGTCAATTTTACGCAGACGCTATCTTTCTAGGGTTAAGCTGCATAACGCCGACCGGTTTGTGGATTGCATTCTTGC

AACTCTATCGGTCGGATTAACCATAGAGTACCTGCCTACTTCACGGATACCTGCTTTGAAAAGTTCAAGACGGCTGGCAACAACGACCGAACAACGAAGCCTCTTTGAGACACTATGATAGTTTGTTACAGAGGTTCATTCTCCAAACCAGTTCATTCGTCGGTTCATATGGATTTGATCGTTTGGGTTTTAAACCAGTTTGAATTATGC

ATAACTATCGAGATTGTGGTAGACTAGGCAGATGATAATTTGCATAGATATTTTGAGGGCTTACCAATCAGCTTGCGCGGGTGCCTATAAATCCCATTTAGCGATAAGATACGGATGCCTTTCAATGTGTACTCAACAGAAATAAGCTAATCTTTGCGTCCACCGAAATCAAATGCACAAAAACAAACATCACATATTTGCGCTGGTGGC

TGGGGTAATAAATTGCAAATGAAACCTTACCGTTCTTCGGTAATCGCCAAACAATGTTCACCACAGCATTGAAGCGATCCTTTTGGACATGGTTCGGTTCTGTTCTTCTTCGAGAGGATGTCGTTTACTGATATTGTTTATCATTCGTTAAACATACCAGCAAGAATGATAAGCTACTGTCTGGCCGTTTCTTAACAATGATGTGAAAAA

ATCAGTGGCCCGCGAAGGGGATGGGATAGATAGAACAAGTCATACGCACACAGCTACCTGGGTAGGCCAGCCAAAACGATTGTCCTACCCATGATACTACAAAAAACAAAAACAATCTTTTTTAAGATTAAGGTCGTATTTCTCAAATGACATACGACAAAGTCGGCTCACTTTAAAATTCTTGCAATAATACAACGGAACATGCGTAAT

TTAAAAAAAATAAGGAACGAACAGCATTCCAAAGGTTCTCCTGGAGTGATTTCGTAAAAATCCTAAAGGAATTTCTACAGCCAGTTAGATAGATGTTCTGAAAATAATTGGTCTAGGCATTCCTGGTGGTATTTCTTGATAAATAACTGCATTATTCAAAGGAAAACTATTGGGAGATTCGTAAAGAATCGTTGGAGAAAGAATTAATAG

ATTCACCCAAGACTTTGCAGTCTTGCTTAACATATATATTACACTGATGGCTCCTTCTTGAAGGTAGAGCTCGTGCAGAATATATTCTCGTGAGCTAGGCTGAATCAGTTTTACTCACTTGGTGAAACTGCACCGTTTTTCAGGCTGAAACATTTGCTCTTATGTGTGGAATGCAATCAACACTTCAACAGCACGTAATTGGTAAAGTCA

TATACTTCTGTTCAGATAGTCGAATGCTATAAAAGCTCTTACTTCGGCCAACTCAAGGTCGAAGCTTGTTGTCGCATGTCGAACTCAAATTGAGGAACTAAATTCAGTCACTCAGGACATTCTGTTATTGCTAACCCTGCGGTGATAAGCTATAGGCTCTCTTTACGCTCATGCGTTTTTCGTTGTTTGAACCCATTGGGGTATGGAGCT

AAATACTCTTTTTTTCGCTTATGCGATTATCCCTCTTCAAAGGGACCCCACTCCTATTTCATCTCATCTCCATTCTCTTTCCTCTCCCCATCTGATATATGATGAAATAGGCTCAAATAGCGATGGCACAAATCTCCCCAACTGATATGATCTGTTGATATCTGCTACCTTCTAGGACATCTGAAAACTATTTGAGAGAAATTACTAGGA

GTAATCGCTAGCAATGCTGGGAAAAATCTTTGGGCTTGTTGGATAAACCCTATAGAGATAATGTCTGAAAAAATAAATGAAGTAAAATCACCATGGGCAAATCGATAGTCTTAGCATTTTTTTTTTTTGAAATGCAAGTGTCGACTCAATTTTTAAAATTAAAATTCCTGACTTTCCCTTGTTTTCCAGTCCTTTTCCAGAATAATTCAG

ACCACTGAATTGGTTGGTTGCACAAAGACGAAATAAAGTCATATTAGTCGGGCTCAGACTACTCAGGTATCACTTCTAGTACTACATTTGGTCCGTTGGTACTACAAAAGTTACCCTAGTTTGACATATCGCAGTACACATTCACGATATTCTAGGCTTTGATCTAGGATACTCGACTTAAGGATTTTAACTGTGATTTTGTGACTTTCC

CTAGACTGCGAAAGCAACAAAATCACAGCTAGATTTCTTACTGTATTACTATTATTTCAAGCGCCATACATTTTTGAGCAACTACTAGGAACATGGGGGTATTTATTTCGTATTTGCTTAAACCGTGATAATTACAGTATGTACAGAAGATAAATATCTGAATATTTTTACATAATGTATAGTTATGTCCCCAACCTGCCAAACTATCAT

TCTACAAGTTCTTCCACACAGCATAGTTCCCACGCCAGGAGATTAGCGCCAGACAAAGCAACAACAACGGACATCCGATCCTGCTTGCGTGACAATTTGGATTCCCAGCTGACCTCAGCATCCCGAACGTGACGATCGCATGGGAAACATCTCGCCGTCATCGATGGTTCCCAGCTGACCGTAAGGCTAGGTAGGTTACTAGATTACTAA

AGCTAGGGAAAAACCGTCAGTTACAGTTTAGATCTAGGACAGAGGTTCTTTACAAAAGTGATTGTGTTTGACATTGTAAACCAAAAATAATACGTAAATAAAGTGATTGTGAAAGTTTGTTCCTTTTTTTTTACAAAGTGATGAACACTCTTTCAAAGTGTATAATTTAATGTTAGGGCGTATAAATAAACTGAGCAAAAGCTTCTACTT

TACTCAAAAACATCTGTTAGATAATTTTCCAGGGGCTTTTTTAAAGCTAGTATGATTAAACTGAGTACATTGATAATTTTTCATTTAGACGATCCATAAATGACGTAGCATTTTTTCACTGATTTTTGACACCCCTCCCCCTCGTAGCATTTAGTGCAAATGCTGACAGACCACTCCCCTGGGAAAATACGTAACATATCAAACAACCCC

CCTCCTTTTATTTTTCTTCAGCGATTGAGGCTTAAAAGGAACCATTAAAATCCAGAAATTCAAAAAAGTTGTTTATTGATATTAATTATTGGCTGAAACATCAGGATAACCAGGCGTATGATGGTTTGATTTACCGTAGCGCTCAAAAACCATTTGGAAAACAGTTTTAAATAAACTTATTTCCTGTTTAAACTATAATTGGTTCCCAGT

TTATACAGGTTAAATTTAATTTGTGGTACTTTTGCTGTTGTTAAAGTCTTAAACAAGTAGACAGCTAAAATGTAGAACAAAAGGTTATCATCTATTTCGGCTTGATAAAGCTAACAGACTCATTGCTGTTGTTGCAAGTGTTCAAGATATTTTATAATATATTAAACAAAAAAAATATTAGAGGATTTTTTTTAAATTATTTGATTATTA

ATTTTACAATATTTAATTAAGTGATTGGAATGGATCTTCGTCATCATTCTCTAAATTCAAATCCCTTCAGAATAATATTAGTAATATTCGGAATGTTTAAAACAGCGATTTATATACAATCATCTTTATTAATATTTCAATTAAACTCCATCATCGTTCCCCATACTGAAATATTCTGACACAACTCATAATGAAACAATTACTAATACA

AATGCTCTTTCGCTGAATTACCGAGTGATTTAGAAAACTGGATGATAGGATATTTTCGATAACACTCGAATTTGATATCCTCATTATTATTCTGAATTTCCATCAAGAACATTCATACACAATGCAAAAATAATGAAGATTACGTCATGTAAACTTCGTTTATGCGATCATAAACATGACGTCATGTAAATTTAAATCTGAAATCATGTA

AAAATGTATTTATATGTAATGTAATCACAGAGGATCCTGCTGGATTAAATGACATATAATGGAAGTTTACATGACATATCATGTAAATATCCATGATATTCGATACTCAATTATGTGCATTATATGTCTCAAGAGAGATTTGCACGTTTCGTTCGAACTGTGTATATCCAAAACAAATCTATGAGAGTTGACTGAAGTCTTCTACAACAT

TGAATAAATATCTTGTTTTACTATAATAACGTTAATATTTGCCTAAATAGGCACATTAAAATCTTTTGAAGAAATAAAAGCTCCAACAGAAAAATGAATCAAACTAAAACAGTTGATTCTATGTCTTCATTAACAAATGGAAAAACAATATGTTCAGTCGATAAAGCATTGATATTAACATATTCATCTTTTAATTTGTTTATTTATAAA

TCAAATAGTATAGACAGCAATATAATATATTATTCACATAACTTTTAAAAAGCTTCACAAAATCTTAAATTTAAAATTTATTAGAATTATTTTAATATTATCATATCAATGTTATGAGGCTGAATCATCATTTATAACCCAAGTTTAATAAGTTACACATAGAAAAACTGCTATGTTGAGATTACTATACTGTTAATTACTTTTACTTAC

ACTGCAAATAGTTCTTTCAGAAGTAAAATTAGCTACGCCAACTTACTAAATTAACATCAAATCAAATACACACACTTCAATGATTCAGCTTTCCTCTTGCACTTATATCAATTATAGTTTTGGATAATTACCACTTTCCGAAAACATTCACATCCTAGACTTCAAGTCATGCTCACAAATCTACTTACTTTTAACCGCAAGGAGAGGGGA

GGCACCGATACTACACACAGGGGGCTGGATACAATTGTTTTACAAATTGCAAGATAACTCCAGTTTGCCAACGACAATCAAGGCAGGCTTAGTGAAAATCGCTAGTTTTTCATTTGTTGTGTACCTTCGATCTTTCTGGACAAACATTGAAAAAGGGCCACAGTTTTCTATGCGTTTTAATTTTCAGTTTAATGAAAACTGTAAAGCCTC

ATTCAATACGTTACGTTAAGAAATTAAGAACAAACGATTGCTCTCGTGACGTTACTTTTGAATGTGCATTAATTGATTCTAATTCGATTTTTGCTACTTTTGATGAAATGCAAAAAATATTCTTGGAAACTAATTACGCGCTTTTATCATGTTTGTCCATTGTTTAAGATCAGGCTCACAGTGACAGTTCGTGACAGCAAAATCTCGGCT

CACTGTGACGTAGAGAATTTTGTATTGGTTTCTCCTCCCAGCTTTTAACTGCAATGAATAGTGTAGGTGCTAACCACCTTACGCAGGAGACCACGTATCAAGTTTACCAACGCGCACTTTGTTTTTTTACAATGTTTTTTTTTAGATAGATAAAGCAGCAACGAATCTTTTCAAATATGATGATATAGCAAAACAAAATGTTGAGTGAAC

ATTGCAAAATGTTGCAAACTAAATCAAATTTACTTTGAGTCAACTAAAATATAGTTTGAAGTTTGTAAATACAATAAAAATATAGTTGTCAACAACTACATGACATAGTTATTTTAAACAACAAATTGTCAAATCAACTATATTTATAGTTATGTCTAAATTAACGCCAAAATAAGTTTATTTGACCGCAAAATGATTAGGCTTTTCTGT

TTGTTCTCCGTGTAAATAAAGAAGTTAGATAAAATAAATGTTGTCAATTTTTCCAGCGCTACTGTTTTTACTGACATTGGCTAAAATGCTACGTCCACTAGTTGGAACCTCCTCAACAGCAACACATCGTCACAAATCATCAGAGAAGCCCCACCTTCCCCCTAAAATGCTACGTCATTATGGACGACCCCTTACAGCAGGAAGATTCGT

ATGCATAAAATGGTAAAACCTGATATTCTGAAAACTCTGAACCATCTACTAATCATACTCAGTTGAAATCTTCAAGCGTGAAAGTTAGATACCTTTACTAAACTATCTCAGAAATAATTGCTTAAATTTGAAATAAACAACTTTAAACTTCCGCTACACAGTGATATACTAAAATGTTTATGAGGCGCAAACTATAAAGATGCATCATGC

TCTTTTTGGCAAAGATCCCTCAAAATTGATCAGCTTTCTTTTCAGGTAAAATTCCCAGGTTTGCGAAGAACTTTATTACGAAAAATTGTAAAATAATACAAGTGATTTCCAACTTGGAATTCTAAAACGTCTCTAAAAAAAAAATTTGAATAATTTCATAAAAAAGTTTTAATAATTAATTAATTTATTTTCTTCAAATACTTCGAATAA

TTTTCCGATGATTCGTTACAGCTTATTTTGTTTCATTACACTTTTTTACTTTGGAAGATTATTGCAGTATTATGCATATGAAAAAAAACTAATTGAATCTTAATACTTCATTTCACAATGCAACATTCCTTCCTAGAAGATTTTCTGGATTCATAAAAAATCTTTCGAAGAATCATTAGAAAATGTTTAGGCGTTATTTCTTTGGTATGT

TCCGAAACAGGATACAAAAACAGAAGTAAACAGGTAGATTGCGGTGTAGGATTTGCTCTATAGCATGAACTGCTGGTATTGATGATCGAATCCATTATTAGATCGAAACGTTCGGTAAATGCTACGGTTTAAGTTATTTTAACGAAAAATCCGATTAAATCGTCAACAAAACTATGAAAGCGGTAATGATCCTGGCATAAACAGTTCTGA

AAAATACAGAATAAAATATGAGAAATCTCTATAAGAATCTCCAGTACAGAAATTTGAAGAGTTTATTCTAAAAAACTGGGCAGCACACTTGAGCAAACAGCAGAGTATCACTTGGACCAGCGGTTTTCATATCGTGCTCTGCGACGCATGGCAGTGCTCTGTATAACAATGCCTTTATGTGAAATCCAAACGGGGCTTCTTTTGACATTA

TTTCCACATTCTTTTAACTTTGAGGATTTGTAACTCTTAGAATTATGGATAGATGTTGATAATTTTGCCTATATTTGGTTGTATATGTACGTAACAAATGAGCAAAATATGAGGCAATCCATCAAGAAATAACAAAAATGCTTGAATAGCGAAAACAGTGTGTCCTTCAAGACAAAAAAGAGGGCTACGTTGGACATTTCACAATTTGCT

AATGAAATGATTTTTCCTTATAACTTTCAAACTGATTTAATAGATTGTCGTCCACTGTGATGTTATGGAACGTTTTCATTGATAAACTTAATGTAGAAAATTGCAAAGCTAGAATCCAATTACATATATAACAAATTTCAACAGAAACATACCATTCACTATTCTACTCCGAAGTTTGCAGAGTATGGAAACTTAAAATTTTCAACTTCC

TCTTAAATGGAACTATCAGTTACGTGCTCGATTTTCAAGTTGACATAAACGAACGACGTAATTACTAGGGATGAATGAGTTATGTACAAAACTCTTTTTTCTTACTGTTATGAACGATATGTTGAAGTCACATTAATACCGGTAACTAATTTAATACCAGTAAGTAATTTTCATTAAAAAGCAATCTATGAAGTAAGGATTTAACAAGAT

CGTAAAGAAGTAAGTTTGCTTTTGTAATATGCTTTTAGCTTCTTAGTAGTTTAGAGCTGGCTTAAGAGTAGTTTAATTTAGGCATTTGAAGGATGCAACTGTTCTATACTAAAATTCATGTAAATATGACGAAAAATAGTTTTATGGATTATATTTGTGAAATTTAGCATTAGTACGTATTGAAATATATGTGTTTATGTCTATTCCTTT

TACAAACATTTATTTTATTTTTTTTTATGTTGTAAAATACACAAGCAAAAACATTATAATAAAATAAAGTCGCAAAAATAAGACATTTGATCAATTATTTTAATAAAACAACAATTTTAAGCAAAACATGGGTAAAATTCAAACAATGTTTATATGTAGCCAATGTTTTTATACCTTATGATATTTATTCAGGCTTTTTGAAATATTTTC

TTGTTTATCCTGTTATTGTTGATGTTGTTCAAGAAAAAAAGCATATATTGAGATTTAATAAAAGTCTTTGAACACAAAATTGCTCAGATTAATATTTACGCTGCTAATAAATACAAAGAGGTGAGTGTCCAAAAGTAGCCCCTGGAATCAAAAGTAGCCCCGTCCGACGGTATATCAGGTTTTGCTTCGAAATACGCTTTTAATTATACC

AAACGAAGCCGATATTATATAATTTTCAATGGTAGTATAGTACCAGAACATTTGGTCAAGACCGCTGTTTTATACTCTAGATTTGTATTCTCAAAAACATAAAGCTGGTTTTAAAGCATCTTTATCTTATGATATATTCGATGCAATAGTGGATGACGAAGACAGTTTTTAGAGTGTTATACACGGTGTCAAAATCTCGATCATTATTGA

ACTTAGATGAACCTCGGATCTTTAAGCTTGAAAACATTCAGCATTTAGGCTAATATTGGCGCTAAATAGAAGTATTCTGTTAATTGAATGTTTTCGCATTGATTTTGTATGAGATTTGTGTATTGAAAGCTGTGAAAAATTGTAACCTTTTCATCTGCCGTTTCAAACCACGATTAAGTATTCGACGAAAAATTGAGGTATTCGACAGAA

AATTCAAATATTCCTACAAGCCACGAACTTTGAAATATATGCCCAAAAATTAAATGAGGAGTTAGAAGCAAAGGATGTAAACAGCTAGTTTTGAGGAAAATCGATTAACAATCAGTTTTTTTTCATTTTATACAAATTGTAAAAAAAACAAGGCAATCTTCCACAAATTATGTTTTTCTCTCTTTGACGGAAAAATCACTTAAAATAATG

TGCGAAAGCTTGGAAACAGTAAAATTATGTTTTTTAATATAAAGTTGCTCAATCAAACAAACCAAATCTTTTCGATTCGAATTAAATTTTAAATTTAAATCCAGAGACCTTAAATAAATTGACCTACATAGATCTTACTTCTTGCTGTGGTTTTGATCACCGTTCTCTTAATCGTTTCCCTTTAGTATTGTTCGGTGACAGGTAATGATA

TCCGATATATAAATTTTACATTTCAACCTACTTTTTATTACGGTCATCTTCAGAAAAAAATCATAGCGTTCAGTAAAATCTCGGCGGTATGCCAATTCTTACTTCATAAATTTTGTAGTAATTATAAAGTCGGTTTAAGCGTATGAAAAAATCGAGATATCAAACGGGAACAAATGCTCCGCGCAACAACTTGATTACCAAAGTGCTCCG

CAGAGCCAAAAGACTGAAAACCCCTGGATTAGATTATAGCTAAGCACATTTTAAAAATTTTAAATTCTAGAAAGTAATAAATAGTTTACTATGGATAATAGCGTTTTTCTCTTTTCTTTGCTATTTTGAAAGATTAATTACCCATTTCATCCACCAAATTTATGTGGTATTCAATGTAAACATAATGATTCTTACCTATTGTATTGCTGG

AGTTGCTCTAATCTCTAACTTTGCAGACAATCAACTCGTCTGATGTTGTAGTATAGTATTAGTGTTTTCCCTGACTTGTAGTGAATTTCCCTGACTTTCCCTGACATTCAAAATTGGATTCCCTGACATTATTATCTTATCTTTATTCCCTGACATTCCCTGACTTTCAGGTTTTCAGGTAGTCGACACCTGGAATGAAAAAAGGGAAAT

TAGATAATTTTTTCCAAGGAATCTCTGAAACAATTCTTAAGGTAGTTCATAAAGGTATCCAAACAAAGATTTATTTATATATGTTTAAGAAGTTCAGGTTGTATTTCATCTAGAATTTTCTAAAACAAAATTGACTGTTTAACGATAAACTTGCGACGATCGACAACGCCACCATATTTCATATAACAGGCTATTAGAACTAACTTAGGT

GATGATTTGGAGGTTATTTGGCAAGTTGAGTTAAAATCACCTCCAAACACTAGCGATTTTGCGGTGGGATGTTGACGTTTGATCACGAATAGGGCAATATTCAAAACTAAAATGGCAGCAGACTAAAGGCTCCAAGGGAAATTCCTCGGATGCATTTATGGATGGACGTCTAAAAAAGATCATTAGTTAATAAAGAAATTTTGTTTTGCG

AGATTCTCAAATGATTCTGACGAGGAGGTCTATTTTATAAAACATTGAATATCTGTAGATTATTTATAACTGTGAGGCGCACTTAATGGGTTATAAATTATGAATTGGAACTTACCATTTTTCGGCAACCCCGAATAACGTTCACCGCCCGCATTGCGGCGATCCTCTATCGACATGATTCGGTTCGATTAGTTCTCGTCTTTGTGATGT

CTGTTTTGGTGTCTCTCTTTGCGCCACCAAACAGCCAACGCAACCAGCAGCTGATTTCACGCTGATTTCGTCGTCGCGGTTTTGTTGTTGAGAAACTGGCTCAAATAATAAACGCCGAAAAGCCAAAGGTCTTCACGGACAACGCGTCCAGTTGTAGTGGATAGATAGATGCTACAGCCTATAGAGAGGTCTCCGTATCTCCCTCTAACT

TTCGACGGCTAATAATGCGGCGACGTCCGCAGGTTATTACGTAAAGGAACGCGTTTCGCTTGGCCGTTCGATCGGTGGGCGTTTACTTCTCTAACAGAGCTTCCCTCCGGGCGGTGCTGGATAGCTGCCTGTTTTGTGGCGTTCCACGCTTTGAATCCGATGCAAAGGTATAATTATATGTGCATATTAAAAACTGTCGACGGCAATGAT

TGCACCAGCGATGACCTGCGTGTCGATTAAGAAAGAACCCCGTCAGTCGTAGACCCAATTCCTCAGGATCACCTTTCCTGCCCTCGGGAAACAAAGCACTGTCTATTCCTCTTATTTACCGGGATCTTCTTTTGTCACCAGCAGCAGATAATCGAGCAGCTTTATTGCTTTATTGACATTTCACGCTCGGGTACTCCTTCTTCCAGTGCT

TCTGTCCCTTATTATTTCGATTCAGCTTCTTCACTCGCGGAACGACAAACGATTACTCGCGAATGACGAAAAGCAAACTGCCAACCGCAGAATTTCGCGCACACATCGAACCAGAAGCGCCCAAGACAAACAACAAATTTCCTGCAGCGTAATACCTATTCTGAACTTCGGCCACTAGAACCTGTGTACACTTTTTGTCTACAGAGCACT

TCCAAAGCGAAACACAAAACTACTCATTCATCGAGGTGGCAATTTGGCTCAGCTCTTGTCGATGATGACCTTTTGTTCTTCGTCAGAATAATTGAGGAACAACGTCGAGTCGAGAGCAATGCATGCCGTCCGGGTGAGGCCACAAAGATTAACTTCATGAAGCGCAACACAAGACAAAAAAAAGATTCTGGTGGCTTCGAATCTTCACGA

AATCTTCCTTCTGCACTTTCAACGCGCACAAGAACATCCACACACGACAAACGGCTGAAGAAAACTGACGGACAGAAAAATGACAGAACGGTTCGACGGACCGAATAGCGCTTTGGGTGAACAAAAATGTTGAGGGTTCCGACCAGTTCATACCGGGAATCGCTCTAATTTATTTTGTTATTTTTAGTATTTAGTATTTATTTCATTTTA

AACTTAAAGCACTGACAAACAGACGTCACACTCTCATGTTGTCCATCGACCACCTTTTTAACAGCCGATTCAAAAATATGGTAGGTGGCCAATCCGCCACCAGCGCTAGCATCGTTTTGTTCGCGTTTGACGTTTACACACTACCGCCATCTGGTGGCCTGTCGGCCAAACACGCTATTTTTAGCATTGGGCGTACATGCGTATGACTAT

GATTTTGATCAAAATTTGTTCTAAGTGTTACGTCTGTTTGTCTATTAACAAAATAAATCTGTCGGTTATACAACAACAATTGATATTGCGACTTTAAGAAAATAACAACTATACGGAATAAAAGTTAAAAAAGATTTGAATGTCATGTAACATAAGGTTAATTTAACTGATAATATAATTTGTGCACTGAAAGCGCAGCTTTGGATGCGC

CAGTGCCACATACACAGATTTATCTGTAATTACACCGATTTTTTTCTGTTTTTGCCTACAGATATCTGTGATCACAGATCACAGATTTTTGAATAAAAAAGATTTTTAAGATTTTAAAATATTTCACGAGTAAATAAAGATATAAAGAAATACCATTACTCTGTTTTATTTTTCTCAAAAAATTTAACTATTTAAATACTATATACTGCT

TCGCGACTAAAATGGGTGAACCAACGCGATGCGAAGTTTGATTTTGACCAACTTCGCCGCGTCGCAACTCGATTCTCAATAGTGCGCATAATGCACACGGTGGAGGGCATAGATGCGCACTATAACAATGACTCGCGACACGGCGAAATTGGTCAAAACCCAACACGTTGGTTGACATTTTGGTCGCGAAGCAATATACGTTGCAAAAGG

GACTTCTCGATCTAAACACATACGTTTTGGATTTTGACACAGATTTTTAACAAGATTTTTTGGAGGTTGACCTAAGATAAAAAAGATTTTCAGCCGAAATCACAGATGAAGATTCGAAAAAATGTGGCAACCTGAATGCGCCTATCGTTGGTAGCTGTCTGGTCACTAGTCCCAGCTAGTCTAAACAGAAAATTTGTTGCCCAGTCCCCA

ACTGGATCATATCGCTTCTGCCCAGGCAGACGGACCCGAAGTTTCTTGTCGTTACGAATAAAGATGGATTGGATTGTTTTTCTTCTGTTTCAGCCACAAGGCCAGGTTACAGATGCGGAAGCTTGGATTGAAAGCATTTTGTTAACTGTAAACTGGTATTCAGAGGAGCAGATGATTTTTAGATTACATAAATTTGCATCCTTGTGCTTT

TCAAGTAGCGCCTGAGTACTAGCGACCTCCTGTAATCGATACAATCATATTATCATTCACATATATATATATATATATATATATATATATATATATATATATATATATATATATATATATATATATATATTTGTTTTATTCATTCGAGAAAGCAACCAACAAATCCCCTCAGCATCACACAAAAGTTCTGCCATTAGTTCTGATTTAATT

TTCCCCGATTTCCGATTCCTTGGCGTTTTGTCCATGGATTTACATTTTCTTTTAGAGATGATCTATCCGATAAGAGCGATTGGGTACATCATAAAGATTCCGATATATCTGAAGAGAAAACTCTCGATTAATTATGTATGGCTGTTTCATACCTAGACAAAAAAACACTCACCTGCATCTAAACAATGTTCCGACGAAGCCTTTAATCTC

TTGTACTACACAATTAGTCCAGCTATTGTTTTCTTTAAAAGCGATTATTTATAAAGGGAGATTTTGAGCCGAAATATTTACAAAGCATAATTGGGTCCTAAAATTTTTAATGAAACCTCGTTTTATTCAATAACACGAAAGAGAATCTTACTGCAACTACTTTAGCAATTTTTCTCGCTCAAATAACGGCTTTATCATGTTTTAAACTTT

AATTTAAAAATTGGAGTCCATAAATGAACCTTGACACTTAAGATCATGTTTGACGTTCAGCAATCGACAAAAACACCACAGGGGTTTTAGTTTTTAACACTGGGGTTGTTCCTATATGACATTTCGGAAGGACACGGAAAACAAATACACCCAAAATTTGAGTTTAAGCCAATATATAAAAAAATGTTTTTGGAGTTCAAGGCAACGGAA

AACATTAGAAAATTGAGTAAACATGTGTTATTAGCCTAAACTTAAGCGTTTGGCACTAAAATTGGGACATGGCTTTAGGGACCCTATTCCTTCAAACAAAACATGACTGGGTGCTTTTCTGTGCCGGATTCGAAGTGAGTGCTGCGGCTGCAGGCTGGAGTCGGAAACGAACATCATTATATGGTTCTTCTAAGGTTCAGCAAATTCTGC

GCATTATTGACCATTTCCGAATAATAAATGGACATCACTTAAGGCCGCACGGGACGTCATCATAATCACCCTATCCATTTGAAAAGCAAATCCCAAGAACCACTCCATATATCGACGTGAAAAATGTATCACTATATATGAGTGCACAACCCAATAAATTTTCAGTTCCATCGGTTCGCCAAACTCGAATTTTCCTCCCCAAAGTTTTGA

TGATAATTGGTTAGAGTGAGACAAAGATAGAAATAACGCACGGTCTCCCATGTCACCTTATAATAAATATTGTGAAAATAATAACTACAACTTTGATGCTGTTTCATCATACCTTAGACCACACATGGAATGTACAACAATAGGTGACAGAATCAGCATTGATTGCAATACGGTTGAATCATTCCGGCTACAATAAATTTCTATGCGGGA

AAAGAGCGATTTTCTGACGACGTCACTGTTCAACCAATAAAACTTTTCCCGGGAAGCAGAATGGACCTATGCACTAGTTCATTATCTGACGTTTTAGCGGTACCGTGTTATCTATGTGACCATGAAACCAATAGGCCAGGAGAAGTGGATGAACTTGTTATTCGTTTTCTGTCAATTGGGTTGTTTAGCGATGACAAAGTTACAGTTTTC

GCGGCAGTCGAAAGATCACATTTTTTTAAAGTATAAACACAACTTTATTGCTTCAGTATCAGAATTTTGACATTATAAATGATTAATGAAATTTCAATACGTAGACTCAATGACATTTCAATTTACATAATGGCAGCTCGTCCGTAGTAAAATTGCCGAGGGGTGATTCAAAGCGATTTCCATACTAACTTCAAACGTGTTTAAAAATGG

TTCAGGGCACGGAAAATTATGAAACTTTGGATTCACACTCCATTTTTAGCAATAGAATCAGAATACGTAAAAAACTGGATACCCCTAAACAAACCAATTGGGTTGTTTAGTGAAAAAAATTCACTGTTTGTTCCAGCTTGATCAGAAAAACTCATCTTTCTAAGTATAATACGATTTTATTGCACTTCAATATCTCAATGTTGATATTTA

AATGATTTAAAAGATTTCCAATCCGTCAAATGTGACATTTCAATTTACACAATGACAGTTTGTCCAGTAAAATTTGCCGAGGGAGTGATTCAAAAGTGATTTCCATACAAATTTCAAATGTGTTTTAAAAATAGTTCCAGGTAACGGAAAATTGTTAAACTTTGGATTCTAGTTTAATTTGACGTAGATTCAGAATATGTAAGAAAATTG

ATACCCCTAAACAAGCCAATTCTCATACAAAATCTTCTTTTTCTCTGACATAAATTCACTCTAGGTGAACCACTTCCCCCTGGCCTATGATTTGGCGCTCAAACGTCAGGTTAATGAACTCATGCATAGGCCAGGGAAGTGGTTCCTAAATGGAGGCCTCATCCAGAAAAAATGGAGGCCCTGAAATTGTTTCAAGAAAAAAATGAATAT

TAAGTTCATCCACTTCTCCTGGCCTATGGTCACATAAATAACACGGCACCGCTTAAACGTCAACAGTGAACGCGTGCATTGATCCATGGACCTATGCACGGGTTCACTGTTGACGTTTTAGCGATGCCATATTTAGGTAGCCATGGAAACTAGTGAATTCGGCACCGCTTAAACGTCAAATTAGTGAACTCATGCATTGGTCCATGGATC

GATGCACGAGTTCACTATCTATTTGTTGAGCGGTATATTACGTGACCATGGCAACGAGTGAATTCAGCACCGCTCAAACGTCAAATTAGTGAACACGTGCATTGGTCCATGGATCAATGCACGCGTTCCCTCGTTGACGTTTAAGCGGTACCGTGTGACCACGACCAGTGAATTCGGCCGCCCAAACGTCAAATTCGTAGAACTCGTGCA

TTGGTCCATGGGTTCATACCACCCTTCTTGATCATTCCAATAATCCGGGGATGTAACCCCTGCCAAGAGTTCATCGTGCTTCACATTAATCTTCACGGCGGTCGTTCTCGCGTCGTCGTCGTCGGCAAAATGTCAAAATGGCATTGTTTAACTTTCTGCGCGTAGTTGTACACTGTGCGATCCAGTTAATACAAATTTTTAGATTTTTAG

TGTAAATTTAGTTAGAGATTCTATAAAAGCCAGATAATTCGGTGCCACTATGCCTACCATCTCAGCACGAGGCGGCTGTTGGCCATAGTGGACGATATTGAAATCATTTCCAAGTAAGTGGCTTTCATAAAATTAGCTATTGGAGTAGTTTGTGTTCAAAAGATTTTGTTACTAGGGAACTAATCGAGAACACCATCGCTCCGAAACATC

AGAAAAATGTCAGCGCCGATCACGGCCAGCTGGGTGGAACTGGTATCCAAAGACAAAGAACTGAAGGCCACCCTGCAGCTGGCGGCAGAACAAAGCCGGCATCGAGAAGAAGACGGATTGCGGGAACAGGTCAAGGAGCAAGATGAAATCAACCAGCTGCGAAACAGCTGAAGGAAGCCGAGCATTCCAGCGTCCATATTCAGGGGCGCG

AAGCTGAGCAGTATTAACAAAAGCCGTCAAGCGACCAGTCTCTTCAGAGGAGCTGATAAAAGTTCGCTCATCGGATCCAGTGCTTCAGATGCAATCTGCGCCCGTTAACGTGGCAACAAGGTGACCTGCAGGCCTTACCCGACGGACATCGAGATGGGGTTGGGATTCCTGGGCAAATCGGACGGCCTCAACATCCAACGGGCTGGCTTG

CCCGAATCAGAACAACCTGAACGAGATGCAGCAGATGCTGACCGGGGCGGGAGCGGGAAGCGGAGTGGCCGACATTCCGGCCTCTACCTGAGAACCAGTTCGCGTGGCATCCGTCCGGCGAGTTGCACGCATGACGATGGGGCAGGGCGGGATCGGTTTCACTGGATACCCGGTCGCACAAAGATGCCTCAAGGATGATGTGGAAGTGAT

GTCCACGGACAGTTTCCAGTTTCAGCTCGAAATTGATTCGCAGTGAAGTGCTGTACTCGAAATGGCTGTTTCGGTGAGTATTTTTTAAATATTTAATAAAGTCAATGCTATAACGTAGAAAAAAAATGATTCATTGTGAAAAGTCTTCTTCCTTCAGGCATTAAGTTTCTACTACTGCGGCAAGCACAAAGAAGTCGAGGGGGATATCGA

AATTTTCCAAATGCAGAAAACATCCGGAACCCCCGCGTTGGCCCATCTATTCTTCTGGTACCGGTACTTAAAGTGTAGTTAAATTAAGAGTTATTAACGGAAACATGATTTTATATAGTGAGAACACGTATCAATAAATTGAAAATTGAATTAAATTTGTAATCATTTCTATGAAACACGTAAGCTTGAAATATATCGTGAGTATATCGC

AAATTGTTGACCGTTATGGTTTAAACAGTAATTAAACACGTGATACCACAAGGTCGATAATTCAGACCATTGTCGTGTGTTCTCTATAATGTTCACCAATCAATCATGTACTGAAACTAATGTTAGATATTTAAACACTCTATGGGTTTCAAATTACTATTCTGAATTCTGATAAAATTTAATGTGTTTGTCTTCACGATTACTGCAATG

TTGTTGAAATTACTTTTGCCAGGGTGGATGAAATTTACTATCATTTGCAGTCAACAGCACTATTGTCCGGTATTAGAGATAAATGATACAAGATCGTTAATTTCTAATGAACAATGTTTTCATTTTACCATGTGCATGGTAGTTTAAACCACAAAATCGTGGTAATAATGTCTCTATCCTCTTGTCTGTTTTCTCCGTCAGCTTATGAAG

GATTAACACCACAGGGTGTCGACTACCTGGAAAAACCTGGAAAGTCAGAAAGAATGTCAGGAGTGATTTTGATCTGGAAAGTCAGGGAAAGTCAGGGAGAATTTCACAAGGGTGGGAAAAATATTCAACTTTAAACAAGTTGATTAAGGAGAAAAGCAACTTTTTATGCATATGAGTAAGCTGTACTTCTCTATCTATACTTATTCTTAG

AATCCAAAATTTTCACAAATGTAAAGCATCAACAGTGATCTCTGAACAATTTCTTCCAAAGTTAGCCGTAATTAAACTTTATTAATATTTGATTCTAAAATTTCTAGATCAAATTATTATCTTCTCTTCAAAATCGCGGAAGAAAAATGGAATCCACTTCATCCAAATCATGCCAGTCTTTAAGTTTGCCCAAAATATCCAAGTTAAACT

GAGTAAAATTAAATATAGAGAACAATAAATGAAGTTTCTCGTATTTAGGTGAAGATGAAAACCGAATCTCAAAATTTTCAGAGCACAAAGCTGCAAGACCAGGATGTTTCCTGAATGTTTATCATGTCGTTTCTGGAAACAATATAGATAATACCGTAATCATAAGTTCTTCAATATTGATATGTCAATTTTTTCGATTTTTCAATTTCA

ATTTTCAGGATTTAGGAAAATATCGTCAATCAAGCATTACTTGAGGTTTTATTCAAGGGCTTACTTACAGATATATTCAGTTTGTTTTTTGTTCTAAGAATATACAGGAAGTTCTCCAAATAATATGAATCATTATCCAAGATATCTTTGTAAATCCATGAAATCCATGAAAGCAGAAATTTCAGAATTCAATTTGAAGGAACTTGCTTG

ATAATCTTTTGATAGTTTGTGAGATTACTTGAGGTTAGTCGGACATCGTCCGTAGATGAGCTAGATCGAGCTCAAGCCGGTCGAAATTTAAATCTTTTTTTAATTGTTTTGTAAAGAACAATATGGTATATCCGTTCTCGTGTCGCGTTCTGTAAGTGTACATGAAATTCTTGTATTACTTCAAACACCCGAAAAAAGTTCATAGAAGAT

ACTCATTGAATACTCATTCTTAGTGGAGTTAAATCGAGGCTTGTTGGAGGCACGTCACATTAAAACGAACATCTGAAAACATATATTAGCACTTGTTTTAATTATGAATCGTTTGACCAACCAGCCGAGAGGAAGTTAACAACACTGAAAAGCCAAACATTACATATAATTTGCAATTGGATTAGATGGACAAATTGATGTGGAGATTTG

CGAAAAAGTTACGTCGACTCGATCTCTAGTTGGGGCGTTACCACTACGCCATGAGAGGACTCATGAACGCAGAAGTTAACCTGAATTCGATTTCAGCTCAATAATCACGTGGTCCTTTTTCGCAAAGTGCACCTCTTTTGGAAGAAATTAGATGCCCATCCAAACACAACACTTTCTATATATCCGTACTTCAGCCGAGCGCAATGTTTT

TAGGTATAGGAATGGCACACTACACTAGCCAGCAACGCTGCGCTGGCTGAGGTTTCTATTGTGTGGGCTTCCGATGGTCGACGGTATTAGCGCATTATTCAATTCCAAATTCATTCCGTTAAAAAATATTAAAGCTTATTTCTTAGGAAAAACGGTTGCTTGGATTTATAAAAAATCACCGGAATTCAAAATTAACTAAAATAAATTACC

GTCAAGCAGGGCTTCTTTTGACATTATTTCCACATTCTTTTAACTTTGAGGATTTAGCAACTCTTAGAATTATGGATAGATGTTGATAATTTTTTGCCTATATTTAAGTTGTATATGTACACGTAACAAATGTGCAAAATATGAGGCAAATCCATCAATAAATAACAAAAATGCTTGAACAGCGAAAAAAGTGTGTCCTTCAAGACAAAA

AAAAGGGGCTATTTTGGACATTTTCACAATTTGCTAATGAAATGATTTTCCTTATAACTTTTTAAACTGATTCAAAAGATTGGCAATCGTCCACTGTGATGTTATGGAACGTTTTCATTGATAAACCTAATGTAGAACATTGCAAAGCTAGAATCAATTACATATATAACAAATTTCAACGAAACATGTCATTCACTATTCTCAGTTTGC

AGTATGAAACTTAAATTTTCAACTTCCTCTTAAATGGAACTATCAGTTACGAATACTCGATTTTCAAGTTGACATAAACGAACGACGTAACTACTAGGTACTGCAGTAAATGAGTTACATTACAAAAACTCTTTTTTTTTCTATTCACTTCAATATATGAACGATATGTTGAAGGATCACATTAATACCGATAACTAATTTAATACAGTA

AGTTATTTTCATTAAAACGCAATCTATGAGTAAAAAAGTTTAGCAAGATCGTAAAGTAAGTTTGCTTTTGTAATATGCTTTTAGCTTCTTAGCAATTTAGAGCTGGCTTAAGAGTAGTTTAATTTAGGCATTTGAAGGATGCAACTGTTCTGTACTGCAAATTCATGTAAAATATGACGAAAAATAGTTTTTAGAGATTATGTTGTGAAT

TTGGCATTAGTACGTATTGAAATATATGTGTTTTATGTCTATTCACTTTTACAAACATTTATTTTATTTTTTTTTATGTTGTAAAATACACAAACCAAAACATTATAATAAAATTAAAGTCGCAAAAATAACACCTTTGATCAATTATTTTAATAAAACAACAATTTTAAACAAAACAAATCACAAGATTTTCATGAAACATGACGATTC

GATAGTTTTGTGATGCTCCCAATAAGTTTCCAAGACATGGGTAAAATTCAAACAATGTTTGTATAGCCAATGTTTTATACCTTGTGAATATTTATTCGGGCTTTTTTTGAAATGTTTTCTTGTTTATCCTGTTATTGTTGATGTTGTTCAAGAAAAAATCATGCCTAGTGGTAGAGCATATATTGGTTAATAAAAGTGCTGAAGACACAA

AATAGCTCAGATTAATATTTACGCTGCTATTAAATACAAAGGTGGTGTGTCCAGTAGCCCTGGTACGATCCCGACGGTACCTGTAAATAATCATGGTGAAGAAGAAATTTAAGAAAAAATGAACACATTTTTGGAAAAGTGTCTATGTGATCCCCTAAACATTTTTTGTAGTATTTCTGTAATATTCCCATCGTTCTAAATTAATTTCTA

GGAAAAAAAATCATGATAAAAAATTCTGTGGGTGAATTCCGGTGAACAATTTCTTGAGAAGTACCTTCATGCCAACAAAGCATTCTTTCTTCCAAAAATGCGATGCCAGGATTCGCCTACTAGGGTATATTTATAAAATTCTTAAACTAGTTTAAAAATCTTTGGACTAGTTTATGGGCTAACGTAAGAGTTCATAAAACATAATTCTTT

GTGATATTAACATCATATATTTGAATAAAACATTTGAGAGGTTCTGGGAGAAATATCATAAGAGTTCCTGGAGAAAAGTGTTAGAGGTTACTTCGAAAAATATATTTGAGAATTCAAATTAAGATATCTGAGAATTTTTGTTCAATTCCTTGATGCATTTTCAAAGAAATTTTTTGGCAATGGTGTAATTTTACCTGAAGAAGAAGTAGC

CAAATTTCAGAAGAAAGTCCTTCCAATAAAAAAGTTTAGTAATAATCTTAATAGAGTGACGTCTTATGTATGAAAAATTGTGTAATCAGGTTGCGGAAGGTGTCAAAAATTGTCTTTTATGCATGAAACTTGTGAAATTATAAGGGTAGTTCAATATCCAAATACATCGTTTTCTACAATTTCAGCTTTTTACTTGAAGATTTCAATATG

GTCAGTAGATGACTCAGAATTAGCAACTTCTACTCAGCTTTTACCGGTTTATTAACCTTCCAGTCGTCGCGTGGTTTGCCACCGTCAGAACCACCACGCTGCTGTTGTGAATAAATAGCGAGCTTTTTCCACACTAGTGCTCAGTTTCATCATATCAACTTGAACAAAACTATCATAAAAATTATTGCTCAGTTTATTCATACGACCCAA

AACATAAACTTTACCAAACAAACAAATCAACATGGTCTGGCATTTTTCTGGGAAAATGACTGGAAAATCAGGAAAGTCAGAAATTTTATTTTTTCAAAATTGGGTCGACACCCTGCCAGCACGGACATAAAATCTATTATTTTAAATAATTTTAGCGTAATATCAAACAGTTTCATCATAATACATATATATCCATGACGCTATTGATCC

AGATTTGGTAAGAAATGTTCAAATTTAAACACAAAACAAACAATTCATGCTCCACAACTTGTTTTCATGTTGCAAGATCACATCGATGTAAACTTCCAGAGCCCGATTAAGTGATTCAAAATAACATTAATGTATTCTATTGTTATTGGTGAGTATTCTATCAATCATCGATTTGTTAGAAATTCAAATAGGTGGAAATAACAGGGTGTC

TACTACATGGAAAAAACCTAGAAAACCTGTAATTATCAGGGAATTTTATTCAACCTGGAAAAAAACCTGGAATACTCAGGGGAATTTGGCTACACTCAGGGAAATTATTTCAGAAACAATAGTGCATGGTAGCATGTATCGTTTTTGTGGAACAACATCTTTCAATCAAAAAATTTTGGCTGCGCCGCGCATTTAGTCTTTTTTAACGAT

TTTTTATTTAACAACTACCGAAAAGCTAAATATTACATATAATTTGCATTCAATTGGATTAGATGGACAAATTGATGTGAAGATTTGCGAAAAAGTTACACGTCTTCTCAGTGAGAATCAGACTCACGACTCCCCCGATCTCTAGTTGGGGCGCGTTACCACTACGCCATGAGAGGACTCATGAACGCAGAAGTTAACCTGAATTCGATT

TCAGCTCAATAATTACGTGGTCCTTTTTTCGCAAAGTGCCTCTTTCGGAAAATTAGATGCCATCAAACACAACGCTTTCTATATATATATCAAAGGTGGTTAGATGGAAAAAGATAACCGACTCCGGCCTTTTCTAGGCGACACGAACAGCGCAGCTCTGGTGAGTCAAAGCCAACTAAAACATAATAATGTTCTCCTAGTGGCCACCAG

ATGTCCAGAAATGTTCGTTTATGACATTTACGCTTAAAAAAGATACACAGCCACACGGGTTCATTCGATATTCCCAATTATCTCTCAATTCCACATAAAAAGTAAGAAAAAAAAGCAGCACAGGAGTGGTTGCAGCATTACACAAATTTTGTGTTGGTTTAGAAACCAGATTATTGTGTTGGTCGTACTCAATTCTGTGTATCTTTCTTT

TTAAGCGTGTTTGTTTTGATTTCTTCCTCTTGCTCATGAACGCAGAAGTTAACCTGGTCGATTTCAGCTCAATAATTACGTGATCCTTTTTCGCAAGTACACCTCTTTCAGGAAGAATTAGATGCCCATCCAAACACCTTGCTTTCTATATATATCAAAGATGGTTAGATGGAAAAAGATAACCGACTCCACGGCCTTTTCTAGGCATTT

TGAACAGCGCAGCTCTGGTGATCAAGCCAACTAAAACATAATGTTCTCCTAGTGGCCGCAGATGTCGAAATGTTCGTTTATGACATTTACACTTAAAAAGATACACAGCCACACGGGTTCGATGTTGGTGTCTCTCTTGATTCCATATGTAAAGTAAGAAACCAGCACATGGTGTGTGATGATGATGATGTGATGATGATGATGTGTGTG

TGTGTGTGTGTGTGTGTGTGTGTGTGTGTGTGTGTGTGTTGATGATGATGTGTGTGTGTGATGTGTGATGTGTGTGTGTGTGTGTGTGTGTGTGTGTGTGTGTGTGTGTGTGTGTGTGTGTGTGTGTGTC

>a52f6291-2d70-49b2-9048-f0bd9b812ad0

ATCAGTGGCCTTGCGTTCAGTTACGTATTGCTATATAGTTGATTTGACAATTTGTTGTTTAAAATAACTATGTCTGTAGTTGTTGACAACTATATTTTTATTGTATTTACAAACTTCAAGCCTCTTTAGTTGACTCCAAAGTAAAATTTGATTTACACATAATATTTTGCAATGTTCACTCAACATTTTGTTTTGCTATATCATCATGTT

TGAAAGATTCGTTGCTGCTTTGTCTATCTAAAAAACATTAAAAAAACAAAGTGCGTTAGTGAAACTTTTGATACGTGGTCTCCTGCGTGCCAAGGTGGTTAGCACCTACACTATTCATTGCAAGTTAAAAGCTGGGAGGAGAAACCAATGCAAAATTTCTCTACGTCACAGTGAGCCGAGATTTGCTGTCACGAACTGTCACTGTAGGCC

CGGATCTTAAACAATGGACAAACGCAGTAAAAGCGCGTAATTAGCCCTAAGAATATTTTTGCATTTCATCAAAAGTAGCAAAAATCGAATTAAATCAATTAATGCATTCAAAGTAACGTCACGAGCACCGTTGTTCCTAGACCCAATGTAACGTATTGGAATGAGGCTCTTTTACAGTTTTCACAAAACTGAAAATTAAACACATAAACT

GTGGCCCTTTTTCAGCTATTTGTCCAGAAAAGATCGAAGGTACACAACAAATGAAAACTAACCCGATTTTTCACTAAGCCTGCCTTGATTGTCGTTGGCAAACTGGAGTTATCTTGCAATTGTAAAACAATTGTATCCAATTTCGTGTGTAGTATCGGTGCCTCCCTCTCCCAGGTTAAAAGTAAGTAGATTTGTGAGCATGACTTGAAG

TCTAGGGATGTGAATGTTTTCGGAAAGGTGAAGTATTATCCAAACTATAATTGATATAAGTGCAAGGAAAGCGCTGAATCATTGAAGTGTGTGTATTTGATTTGATGTTAATTTAGTGGCGTAAATTAATTTTACTTCGAAAAAGAACTATTTATTGCACGTAAAGTAAAAGTAATTAACAGTATAGTAATCTCAACATAAAGAAATTTT

TTCTGCGTGTAACTTATTAAACTTGAGTTATAAAATGATGATTCAGCCTCATAACATTGATATGATAATATTAAAATAATTCTAATAATTGCAAATTTAAGATTTTGTGAAGCTTTTTAAAAGTTATGTGAATAATATATTATATTGCTGTCTATACTATTTGATTTATAAATAAACAAATTAAAAAGTGAATATGTTAATATCAATGCT

TTATCGACTTGGAACATATTGTTTTTTCCATTTGTTAATATAGAATAGAATCAACCGTTTTTAGTTTTGATTCATTTTTCTGTTGGAGCTTTATTATTTCTTCAAAAGATTTTAATGTGTGCTCATTTAGGCAAATATTAACGTTATTATAGTAAAACAGATGTTTATTCCAATGTTGTAGAAAGATCTCCCTCTCCAGTCAGCTCCTTA

GATTTGTTTTGACTCTGCACAGTTCAGAACAAAACGTGTAAATTTCTGAGACATATAATGCACATAATTGGGTATCGAATATCATGGATATTTACATGATATGTCATGCTCAAACTTCCATTATATGTCATTTTAATCCAGCAAGGATCCTCTGTGATTACATTACATATAACACATTTTACATGATTTCAGATTTAAATTTACATGACG

TCATGTTTACATCGCATAAATGAAGTTTACATGACGTGTAATCTTCATTATTTTACTGTGTGAATACTCTTGATGGAACAACCAGAATAATAATGAGGACACTCAAATTCGAGTGTTATCAATATCCCTATCATCAGTTTTCTAAATCCTCAGTAATTCAGCGAAAGAGCATTTGTATTATATAATTGTTTCATTATGAGTTGTGTCAGA

ATATTTCAGTATGAGAACGATGATGAGAGTTTAATTAGAAATAATAATAAAGATGATTGTATACTCGTAACTGTGTTTAAACATTTCCGAAGATATTACTAATATTATTCTGAAAGGGATTTGAATTTAGAGAATGATGACGAAGATCATTCAATCACTTAATTAAATATTAAAATTAATAATCAAATAATTTAAAAAAAAATCCTCCCC

AATATTTTTTTTTGTTTAATATATTATAAAATATCTTGAACACTTGCAACAACAGCAATAGGTCTATTAGCTTTATCAAATGAAATAGATGATAACCTTTTGTTCTACATTTTAGCTGTCCTACTTATTTAAGACTTGTAACAACAGCAAAAGTACCACAAATTAAATTTAACCTGTATAAACTGAGACACCTAAATTATGTAGTTTAAA

CAGGAAATAAGTTTATTTAAACTGCTCCAATGGTTTTTGCCCGAGCGCTACGGTAAATCAAACTATCATACGCCTAGTTATCCTGATGTTTCAGCCAATAATTAATATCAATAAACAACTTTTTTTGAATTTCTGGATTTTAATGGTTCCTTTTAAGCCCAATCTAACTGAAGAAAAAGAAAAAATAAAAAGGGAGGAGTTGCTTGATAT

GCTACGTATTTTCCCAGAGCAGGTCTGTCAGCATTTGTGACTAAATGCTACGAGGGAGGGTGTCAAAATCAGTGAAAAATGCTACGTCATTTGGACGGCCCCTAAATAAAGAAAATTATCAATGTACTCAGTTTAATCATACTAGCTTTAAAAGCCCTGGAAAATTATCTAACAGATGTTTTGAGTAAAGTAGAAGCTTTTTGCTCGGTT

TATTTATACCTATTAAATTATACACTTTTGAAGAGTGTTCATCACTTTGTAAAAAAGACAAACTTTCACAATCACTTTTTATTTACGTATTATTTTGGTTTACAATGTCAAGCCACAATCACTTTTTTGTAAAACCTCTCTGTGTCCTGAATCTAAACTGCCTTTGACGGTTTCCCTAGCTTTAATTAATCCTAGTAACCTACCTAGCCC

TACAGTCAGCTGAGACCATAAATGACGGCGAGATGTTTCCCATGCGATCGTCACGTTCCGGGATGCTGAAGTCAGCTAAGGATCAATTGTCGTACGGATCGGATGTCCGTTGTTGTTGTTTTGTCTGGCGCTAACTCTGGCGTGGGAGCTATGCTGTGTGGAAGAACTTGGCATAGAGATGATAGTTTGGCAGGTTGGGTTATAACTATA

CGATATGTAAAATATTCAGATGTTATCTTCTCGTACATACTGTATTATCACGGTTTAAACCAAATACAAATAAATACCCCCATGTTCCTAGTAGTTGCTCAAAAAATGTATGGCGCTTGAAATAATAGTAATACAGTAAGAAAATCTAGCTGTGATTTTTGTTGCTTTCGCAGTCTAGGGAAAGAGTCACAAAATCACAGTTAAAATCAC

TCAGTCAGTATCCTAGATCAAAGTCCTAGAATATCGTGAAATGTGTACTGATAAAACTAGGGTAACTTTTGTAGTATATGTGGACAAAATGTAGTACTAGAAGTGATACCTGGAGTAGTCTGAGCCCGACTAATATGACTTATTTCTAGATCGCTTTGGTGCAAATCAACCAATTCAGTGGTCTGGAATTATTCTGGAAAAGGACTGGAA

ACAAGGGAAAGTCAGGGGAATTTTAATTTTAAGTGGAGTCAACCTGCATTTCCAAAAAAAAAAAAAAAAATGCTAAGACTATCTGACGGCTCCATACAGGTTTTACTTCATTTATTTTTCCAGACATTATCTCTAGAGGTTTATCCAACAAGTTTCTCCAATTTTCCCAGCATTGCTAGCATTATTCTAGTAATTTCTCTCAAATAGTTT

TCCAGATGTCCTAGGAGGTAGCAGATATCAACCGAATCATATCAGTTGGAGATTTGTGCCCATCGCCATATTTGAGCCTATTTCATCATATATCAGATGGGAGAGGAAAGAGAATGGAAAGATGAGATGAAATAGGGTGAAGGTCCGCCAGAAGAGGATAATCGCATAAACGAAAAAAGAGTATTTTAGCTCCATACCCCAATGGGTTCA

AACAACGAAAACGCATGAGGCATAGAGAGCCTATAACATCACCGCAGCTGAGTTAACCAATAACAAGTCCTGAGTGACTGAATTTAGTTCCTCAATTTGAGTTCGACATGCGACAACAAGCTTCGACCTTGAGTTGTGAAACAAGAGCTTTATAGCAGCCCTGACTATCTGAACAGAAGTATATGACTTTACCAATTACGTGCTGTTGAA

GTGCTGATTGCATTCCGCACATAGAGCAAATGTTTCAGCCTGAAAGCGGTGCAGTTTCACCAAGTGAGTAAAACTGATTCAGCCTTGGCTCACGAGAATATACTCTCCTGCACAGGCTCTGCAGCAAGAGGGAGCCATCAGTGTAACATATATGTTACAAGACTGCAGTCTTGGGTGAATCTAATAATTCTTCCAACGATTCTTGCAATC

TCCCAATAGTTTTCCTTGAATAAATGCAGTTATTTGTAAAGAAATACCACCAGGAATGCCTAGACCAATTATTTTCCAGAACATCTAATATCTAACTGGCTGTAGAAATTCCTTAGGATTTTTACAGAAATCAATTAGAGAACCTTTTGGAATGCTGTTATTCGTTCCTTATTTTTTAAATTACGCATGTTCCGTTGCTTATTTCTGCAA

AGAATTCTTAAAGTGGTTTCTCCAGACTTTTTGTCGTATGTGCATTGAGAAATACGACCTTAATCTTAAAGATTGTTTTTTATGTTTTTGTAGTATCATAGATGGGATAAATCGTTTTGCTGGCCTACCCAGGTAGCTGTATTGCGTATGACTTGTTCTACCTATCCTATCCCTTCATGGGCCACTGATTTTTCACATCATTGTTAAGAA

ACATTGAACCGAGTAGCTTATCATTCTTAGTTGGTATGCTTAACGAAGATGATAAAACAATATCCAGTAAAACGACATCACTCTCGAAAGAAAGACAGAACCGAACCATGTCCAAAGGATCGCTTCAATGTTGTGGTGAACATTGTTTGGCGATTACCGAACAGTAAGGAGTTTCATTTGTAATTACCCCAGCCACCAGCGCAAATATGT

GATAAGCTTGTTTTGTGCATTTGATTTCGGTGGACGCAAGGTGGCTATTTCCTGTTGGTACATTGAAAGGCATCGTATCTTATCTATATAAAATGGGATTTATAGACTGCGACTGATTGAGTAAGCCCAAAAGAAATATCTGTACAAATTATCATCTGCCTAGTCTACCACAATCTCGATAGTTATGGCATAAATTCAAACTGGTTTAAA

CCCAAACGATCAAATCCATAGACCGACAGATAAGACTGGTTTGGAGAATGAACCTCTGTAACAAACTATCGCAGTGTCCTCAAGAGGCTTCGTTGTTGTTCGGTCGTTGTTGCCAGCTCGTCTTGAACTTTTTCGTGCTATAGGTATCCGTAGAGTAGGCGGGTACCTGTATAGTTAATCCGACCGATAGAGTTGCAGAATGCAATCCAC

AAACCGGTCGGCGTTATGCAACAACCTAGAAGGAATGGTCTGCGTAAAAATTGACGCATGCATTCTTGAAATATTGCTCTCTCTTTCTAAATAGCGCGAATCCGTCGCTGTGCATTAGGACATCTCAGTCGCCGCTTGGGGCTCCCGTGGGGCGTGCTTGTCAATGCAGTAAGTGTCACTGATTTTGAACTATAACTTGACCGCGTGAGT

CAAAAATGACGCATGATTATCTTTGCAATGACTTTTAAGATTTAACTCATACGATAATTATATTGTTATTTCATGTTTCTACTTACGTGATAACTTATTATATATATATTTTCTTATTTATAGATATCATTGACTAATATATAATAAAATGGGTGGCTTAGTTCTTTAGACGATGAGCATATCCTCTCTGCTCTTCTGCAAAGCGATGAC

GAGCTTGTTGGTGAGATTCTACGACAGTATCGAAATATCCAGATCACGTACTTAAGTGAAGATGACGTCCAGGCAGTACAGAAGAAGCGTTTATAGATGAAATTTTATGAAGTGCAGCCAACGTCAAGCGGTAGTGAAATATTAGGCGACAAAATGTTATTGAACAACCAGGTTCTTCGTGAAAACAACAGAATCTTGACGCACCTGAGG

ACTATTAGAAGTGAGAATAAACTGTTGAATTAAGCTTCAAAGTCCTGAGGCGTAGCCGAGTCTGCACTGAACATTATTCAGGTCTGTCGACATGCCCGCCGTGACCGTCGAGGCCCGCTGACGCTGCCCCGCGTATCCGCGCACCCGCCGACGCCGTCGCGCGTCCCGTGCTCACCGTGACCACCGCGCCCAGCGGTTTCGAGGCGAGGG

CTTCCCGGTGCGCCCGCGCGTTCGCCGGGATCAACTACCGCCACCTCGACCCCGTTCATCGCAGTGGACCAGATGGGTGAGGTGGAGTACACGCCAGGGAGCCCAAGGGCACGCCCTGGCACCCGCACCGCTTCGAGACCATTGACCTACATCGTCGACTAAGATACATTGATGAGTTTGGACAAACCACAACTAGAATGCAGTGAAAAA

ATGCTTTATTTGTGAAATTTGTGATGCTATTGCTTTATTTGTAACCGTGCCGGCTGCAATAAACAAGTTAACAACAACAATTGCATTCATTTTATGTTTCAGGTTCAGGGGGAGGTGTGGGAGGTTTTTTAAAGCAAGTAAAACCTCTACAAATGTGTGCTGGCTGATTATGATCTAGCGGCCGCTTTACTTGTACAATCGTCCCATGCC

GAAGTGATCCCGGCGGCGGTCTGGGCTCCAGCGGAACCATGTGATCGCGCTTCTCGTTGGGGTCTTTGCTCAGGGCGGACTGGGTGCTCAGGTGGTTGTCGGGCAGCAGCACGGGGCCGTCGCCGATGGGGGTGTTCTGCTGGTAGTGGTCGGCGAGCTGCACGCTATATCCTCGATATTGTGGCGGATCTTGAAGTTCACCTTGATACC

CGTTCTTCTGCTTGTCGGCCATGATATAGACGTTGTGGCTGTTGTAGTTGTACTCCAGCTTGTGCTAAGGATGTTGCCGTCTCCCTTGAAGTCGATGCCCTTCAGCTCGATGCGGTTCACCAGAGTGTCGCCCTCGAACTTTCACCTCGGCGCGGGTCTTGTAGTTGCCGTCGTCCTTGAAGAAGATGGTGCGCTCCTGGACAGCCTCCG

GGCATGGCGGACTGAAGAAGTCGTGCTGCTTCGCTGTGGTCGGGGTAGCGGCTGAAGCGCACTGCGCGCCGTGGGTCAGAGTGGTCACGAGGTGGGCCAGGGGCACAGGCAGCTGCCGGTGGTGCAGATGAACTTCAGGAGGTCAGCTTGCCGTAGGTGGCATCGCCCTCGCCCTCGCCGGACACGCTGAACTTGGCGTTACGTCGCCGT

CCAGCTCGACCAGGATGGGCCACCACCCCGGTGAACAGCTCCTCGCCCTTGCTCACCATGGTTGGGAAATCTCTGTTGAGCCAGAAAAGAAACGAAACGCTTGAGTGGTGGTTGTGAAATGCAAACTCTCATTGATATTGATTCATTTGCCTTTGGCTTCGAGGCACGACACGACAGGTTTAAACTTGTTTTCGCTGTCTGCGTTTGCAG

TCGCGAGCCAAGTAGAAAAATATACACTTGAAGGTGATGACGTCACAACCAACGCCCTACTTTTAAGTGAAAATTAACGTTTTCGACTTTTGAACTACGTAGTTTTGAAATTGCGTATCTTCAAGTTTTCCCGATTTCCTCAAGGTTTTTCTCGATATGTGTTAATATTACCTTAATGGGTAATTACCATCAAAATATTTATTTTTAGAT

ATGTGACAGGAGCAAATACGTTATTCTTATTATTCTAGAAATTTAATTCAATTGGTAGCGATGATTCAACGAAATATGATTATCGCTGTGAATCTGATGGGTTTATCAATGATGATGAAACTGCGTTGCAAATTTTCACTAATCACTCAAAGCTCAATGGTCGCCATCTTAAGAAAATAGTTTGCTCATTCCAAAGATAAAGAAATCATT

CACCAAACTAGTTTTCGCTCATAGCTATAATTTCATCCAATTATTAATTTACCTACCTTCTTCAGAAGATTTCCCTTCACCGAAATGCACTTTCACCAATTTAATAATGGTAACTGTTCTCCTTTGAAATAAAGCTGTTCACTCTGAAATTTTCTCCTCTGGCTAATTGGATCACTCTTTTCACCTAGAGACTTCACTTCACTTGCACTA

TTTTTTTCTTACTGGGCCGCGTAATGTTCACTCACTAGAGCATATTCTGATTGAGCTAGTTTCGCCTTTGGGGGCGTTTATAGACACTGTCGTAGTGGTGGACTGTGCTACAAATTTCAGCAATACATTCATACATATACCTCTGTTCGGTTGGATGGCTCTATGTCGATCGAATATGGGTACCATCCCTGTCTGAATGGACCAGCAAAA

CTGATGCTTGTGTCTGTTAGTCGTTCATTCTGTTGCCTTGAACGATGCCAGTTCAACTACAGCAAGACGTCAATGTACCTACCCTTCGTGTATATGGCGCAGGAGGAGATGTGCAGATGCCTTTTTCGATAGAGAAAGGATTTCTACAATTTGATCGACAATTTCGGTAGCTAATCTTAGCAACTGAATGTAATCTGATAATAAACCAAA

TCAGAAAAATCAGTAATTTGGGAATTTCACTTGAATCATTCTAATTGCATCATTTGCTGTATTATTAATTTCAGTCAGCAAGTGACATCAACCCTTCTAAATCGATATACTTCTGGGAAACTTCTTTCTTGTCTGGCTCAGCTGGTGCCAAGGCAAATTATAAGTGAATTCAATGCAAACTACATGTAAAGATATTAACCATTGTAGAAC

CGTGCGATCAAACAAACGCAGAGATACCGGAGAACTGAAAAACAGTCACAACTCCAGGCCAATTAAGACATCGATGTTTGTTTTGACGGACCCTTACTCTCGTCATATAAACTGAAGCCAGCTAAGATGGTATACTTATTATCATCTTTGTATTAGCAGAGATGCTTCTATCAACGAAAGTACCAGTAAGCCGCAAATGGTTATGTATTA

TATCAAACTAAAGGCGGAGTGGACACGCTAGACCAAATGTGTTCTGTGATGACCTGCAGTAGGAAGACGAATAGGTGGCTATGGCATTGTTGTACGGAATGATAAACATGCCTGCATAAATTCTTTATTATATACAGCCATAATGTCAGTAGCAAAGGGAGAAAAGGTTCAAAGTCGCAAAAATTTATGAGAAACCTTTACATGAGCCTG

ACGTCATCGTTTTATGCCTGCGTAAGCGTTTGGAAGCTCCTACTTTGAAGAGATATTTGCGCGATAATATCTCTAATATTTCGCAAATGGGGTGCCTGGTACATCAGATGACGGTACTGAAGAGCCAGTAATGAAAAAACGACACTGTGCCTGCCCTCTAAAATAAGGCGAAAGGCAAAGTACATCGTGCAAAAAATGCCAAAAGTTATT

TGTCGAGAGCATAATGATATGTGCAAAGTTGTTTCCTGACTGACTAATAAGTATGTTGTTTCTATGCTATGCTTTTAAGTTAAACTAGTACTTGTTTTATAATATAACATGACTGTTTTTAAGTACAAAATAAGTTTATTTTGTAAAGAATGTTTAAAAGTTTTGTTGCGCTTTATAGAAGAAGAAATTTTGAGTTTTTTGTTTTTTAAT

AAATAAATAAACTTTAAATAAAATTGTTTATTTGAATTTATTATTAGTATGTAAGTGTAAATATAATAAAACTTAATATCTATTCAAATTAATAAATAAACCTCGATACAGACCGATAAACACATGCGTCAATTTGCATGATTATCTTCAACGTGCGTCACAATATGATTATCTTTCTAGGGTTAAGAACTGCGATCACAAAATGGCCAG

TCAGTCAACATTGAAATGTTGTCTGCTTTGAGTGAAGGCAGTTGGTCGCAAAACGTCCCTTGCTTGGGCTGATAATCTTTGAAAAGGTAATAATGGAACCATAGTTTGACGAAGATCACCAATGAGCGCACCATCATGCAAAGGGTAAGCTTAAAGTCACCCCTATTGGGGTAAAGAGTTTCTCCAATCTTGAACAACCACTCATATGTA

GAAGTTTTAACGAAAAACAAATTTGCATTTCTCAAGGCCGTTTTTAGAATGTCGACTAACCGAAGAAGAAAAACTGGAAACGTCAGGGGAATCCCATTTTTGACAGAAAGTCAGGGAAATAACACTTAAAATTGGGTGAAAAACATCCAAACGAGTTGTGCCAATATAAAACAAACGAAAATCGTTGATGCCAACTGAACGTTGCCAATA

AATAAGACATTTAAATTTTTCTGTAAAGTTATTCGGCGCAGCCAGTTGAGCAATACAAAAGATTGCAATCTTTAATGTCGAGTAATGGGCCTCGTGGCCCGTGCGGTTAGTGTCGTCCAGGCATTAAGCGCATCGTATCATGGGGTGTGGGTTCGTTCCCGCTTCTGCCTTTGAAACTTTTTTCGTCAGGAGGTCGCAAACCACTAGAGC

ATGCGTGTACGTTGTCTAGTGTTAAGTTACAGTCTGTGTGCAGCTAAATGGCTGAAGACGGTGTCCATGTCTTTTTTAAATATCTAACTTATCATTTCGACATACCTTTGAATAGGTAATTCGATATGTACGCCCTTATACGCCTACTTGTCCCATGTTCTATATGACAGCCTTATGGACCGCGGGACAAATATGTGTAGAACGGCAGTA

TTTTTTCACAGGTTTGCTTTGCCAGCGTTAGCGTTAGCGTTAGCGTTAGCGTTAGCGTAGTTTACATTATACTTCGTAGATTGGATACTAGCTATATTCAGTAGATGAAAAACCGAATTTGGTCACTATGCCATTAATTCCACTAGAGTTTTGTATCCTTTGACAGATACGCGTATTTCGACCTCAACAGATACGTCTTCAATTGTCGTG

TGCTAAACATTGCAGGTGCACGACACTGAAGCCGGCTGGTCAATATCTGTCAAGAGACAATTATGAATATAAATTAAATTTCTATTCTACTAAACAGCTCGAAGATTTATTAATATATTGGGTTTTTCTTTGATAAGCTCATCTGGGTGCTTTCCAAGATAAGGCTTGGGAGTTAATGCTATCCAATCTTAAACAAAGTACCAAAACCAT

GAATACTGCTATTCCCGGCCACGCCCATCTTGCGTAACTTGGGATAGGAAGGAAATGTTGATGTAACACTTGCCAATGAGAGGCCACCGACTCAACTGACACCCTCATAAGTGCTACAGGTTGGGAAGAGTTAGAGGAGGGTTTCAGGTCAGGATTCGCTGAAGCTAGCGATAGACACCAGATCATTGGATGTTTAATGAAACTTATTTA

ATCTCTTGAATACCTGACATTAATCAAAGAGAAACAATATGTTATTTACAGTATGAGAATGTTGTTTCTAATAAAAAATGATTAGATACGTGTTCAACGGTAGTAGTTTTCTGCTTTTTACCAGCAAAAACAGAATCGATCCTCAAAGGGTCATCAATTAGAAAGAATAAAACATAATTGACCGACTAAAGCGTGCGAATTGAAAAAAGC

CATATGAAAAAAATCTGGAACGAATAACGTGTATTGAATTTCTGACACGCGACGAAGGACCTATTTCGTTGTTGAAGAAAATTGCCCACGTCTTTTCACTACAATCTAATAGTAGAAAACGATTGAAACTGTCTAGCACTAACATATTTCAAATAACTGTGCACTTGAGACCCGAAAAAAAAAAAACTGACAGTGGTTCCAAAGCACAAC

CACCCCTTTCACAGGTTTGCTTTACTTATTGATATAAACTCATTGTAAAATTGACGACAGAGTTTCTTGTATTGTAGGATTAGAAATCTTTGAATGGGAAAATATATGTGCGGATTTTTCGTTTTAAGACGATTTGAGTTATGTTATTATTTGTTGTTCTAAAGACTGGCTTGAAAACCATGAGCAACAAATGAAAACTGTGGAATAATT

CTTGAAGGTTTAAATTCTATGCTAGTCGAACATTCTAGAAGACACCCTTAGAAGATCTTTGGAAGATACACCGGATGGTTAAAAAAAATCTCAAGAAAAGAATTCTATTAGTGGAATCACCGAAACAAGTCTATACGGAACCTGCACGAGAATCTCTGGAATAATTAAAGACAATTGACCAAATACTGGTTGAATTCTTCAAGAAATTTT

GAACAAGTCTTGATGGTGATTTCTCTACATATGGAATATACTGCTTGAACACAAAAATCTTATGCGCATCTAGAAAAAGGATGGAAGAATAATTGTAAATTTTCAGGACGATTTTGTTTTTTTCTGTGTTAACGAGATTTTTAGCCCTGACATCTCGGGACCAACGGCTGGCTTACTTCCTTCGAAGGAAGTTGTCCTATAACCTTTTGT

CATAGAAGTAACTATCTCTCTGAGGGATGGGATTCAGTCCATATCCTCTGACATAAGGGCATATGTGTTCTAATTTCTTACAGATCCATACTCCCAACTGGAGGACTTTGCTTCCGAAAAATGTCTGGTTGATTCTCAGAGTTAAGTAGAGGTTTAAAGAAATTTGAATTCCAGTCTAGAAAAAAATGGAAAATCTGAAATTTCCAACAG

AATATTTATGGAAAACCAATGAAAGAATAACCACATATTCTCTCTAACACTTTTTAATTTTAGATATTTTAGAAAACCAACTTATTAGACAATTTTATCAGTTTCAAGTCTGTTATTCAAGGATGCTGGAAAAGGAGATGGAATAAAGTTCTGTCATTGTTCTGCTTATTTCTCCTATTATTTATAACACTTTCAGTAACTATGTGTATC

AATACACCTCATCATCAGCTGGTGCCTTCTACTTTATTGGCCTCGAACTAAAATACTAAAGCCTTTCAGTGTCTATGAAAAGTCTTAAAGCATTCAGTTAAATTACAATCTGTTAATTTTCTATATTTTGGACCTGAAAAACCTGGAATTATCAGAATTTCGATAAAACCTGAAAAAAGCCAAAATTCTCAGGGAAATTTATTTGAGGCA

GTAATTCATTGTAAAATATGATGATATTTCAAGACAAAGGTTTTTCCGCGTCAAAATTCCCAAAAAATATTAACTGCGCTTTTGGGCTGTTCGGTGAGGATCTTTTGGGGTATGATATCTTATTGACTTCCCGACATGATTGAATTTCACGAATAATATTTGATCAATTAGCCGTTCACATATCTTCATGCACGTTTTAAAGTATTCAAC

GAAACTTAAAGCTAGAAAGGCAGTCTCGAGAAAATAATCCGGAATTCAAACAACTTTCCAATTAAAATGCCAAACTGTAAATTGCCAGTTTCTTCAGGGCTGAGTAGATGGTCTGTTTTTAGATGCTGGTCTTATTTTAATGCTGGTCTCTAAAATTATTTATTTGGTGAAACAGAAGGTCTCTAAGGTTTTTATTGTTTTATCCAAAAT

GGGTTCTAAAGTCTCTTTTTTCTATATTTTGTTCGCGATTTGATTAATTATGATGATAATGTAAATTATGCAGGCTCCAAAACCAGAGACTGTTACGGACACTAAACTATTTTCAAAAATTCCCTCTCAAACTTCTCGTAGAAGAAGCTTTTGGAATTATCGTGAGTGGTTTCTCCAGATATTTAATCTGGAATGCTCAGACTCCTACAC

GTATCTCTCTATGGTCTTCTAGAATTCTCTACCAATTCTTTCAGCGTTTTTTGGAAAGAAGAATATCTTGAGGCTACTCCCATAATTAAGCTCAAGAGGTGTCCAAGCACTTCAACATATACTCCAGTAATTTGCTCAGAAATTACTTTGAACTTTATACACTATCACCAGGATTTCACAATCAGATGTTTTTCCACTAATTCAGCGGGT

ACCAGTTGTTACTAGAGGGGTCTCCCAAAGATTTTATAATTTTTTCTCAAAAAAATTTAAGAGTTTTCTTTGTTAACTTTGAAAAAAATCTCAACAATTCCTCCAAATTCAGGCCTATGTTTTTGTCATTGAATTCCTCTACACTTTCTAGAATTTATCTAGAATTTCTCAATTATACTACAGTTCTTTGAAACATATCTTAAAAAGATC

CCGTGAAATCTTCTTCTTTCTGGCATTACGTCCCCACCTTAATGAGGCCTGCTTCTCAGCTTAGTGTTCTTATAGGCACTTGCACTGAGCTTACTATGCCAATGACCATTTTGCATTCGTATATCGTGCTTGGCGGAGCACGAAGATGCTCATGCCCTGGGAAGTCGAGAAAATTTCCGCTCGAAAAGATCCCTTGACCGGTGGGATTCG

ACCTCAGCAGTCTGCTGCTGAATAGATGCGCGTTACCGCTGCATCTGGGCCCCTCCGTGAATCTATTGGGAAGTTTATTTAAGGTTTCTACTAGTGCTTGCAGCATTTTCCAGAAAATCCTCCTACTCTCATATTTTAAGAGATTCCAGTCCGATGTTTTCCCAAAGAATTTCTTCAGAAAATGCTGCTTGATATTAAAGCCTTGATATT

GTTCATTTAAGAAAATGGACACGTGGACTTCTGAAACTTTATTTTCCAGACAAATTGATGAGTTCACCTAAAATACATTTCGTAATTTCGATCAAATTCACATACATTTCATCATTTATTAGACATTCCTGGAAATGCTCCTATTTCTCTTGAAACCTCCCCTTAAGGGTCTGCACAACCTTTTCGTTGCCGTAACTTTTCGTTGTGTGA

CCACAATTTTTCTTGACAAAATCATTAGACTTGTTCTTCCATAGCAATGCATGGCA

>fe307069-d616-41c4-8026-58873d6065ba

TCAGTAATACTTCGTTCCGATTTACGTATTGCTCTAGGGAAAGTCACAAAATCACAGTTAAAATCCTTGGTCAGGTATCCTAGATCCAAAGTCTAGAATATCGTGAAATGTGTACTGCGATATGTCAAACTAGGAGTAGCTTTTGTAGTATAACGGACCAAATGTAGTACTAAGTGATACCTGAGTAGTCTGAGCCCGACTAATATATTT

ATTTCGTCTTTGGGTACCAAATCAACCAATTCAGTGATTCTGGAATTATTCTGGAAAAGGACTGGAAAACAAGGAAAGTCAGGGAATTTTAATTTTAAAAATTGAGTCGACACCTGCATTTCAAAAAAAATACATAAGACTATCGACGGCTCCACACAGGATTTTACTTCATTTATTTTTCAGACATTATCTCTATAGGGTTTTTTTTTT

TTTTTTTTATCCAACAAGTTTCTCAAAGATTTTTCCAGCATTGCTAGCGATTGTTAATAATTCTCTCAAATAGTTTCAGATGTGTCTAGAAGGTGGCAGATATCCAACCAGATCAGTTGGGAGATTTGTGCCATCGCCATATTTGAGCCTATTTCATCATATATCAGATGGGAGAGGGAAGAGAGAATGGAAAGATGAGATGAAATGGAG

TAGAGTCCTTGAAGAGGGATAATCGCATAAACGAAAAAGAGTATTTAGCTCCATACCCCAATGGGTTCTAAACAACGAAAAACGCATGAGCGTAAAGAGAGCCTATAGCTTATCACCGCAGCAGGTTAGCAATAACAGAATGTCCTGAGTGACTGAATTTAGTTCCTCAATTTGAGTTCGACATGCGACAACAAGCTTCGACCTTGAGTG

TGAAAGCAAAACTTTTATAGCAGCCTGACTATCTGAACAGAAGTATATGACTTTACCAATTGCGTACTGTTGAAGTGCTGGTCCATTCACACATAAGAGCAAATGTTTCAGCCACAAACGGTGCAGTTTTCTACCAAGTGGTAAAGCAGTTCAGCCTTAGCTCACGAGAATATACTCCTGCACGAGCTCTACCTTCAGAAGGAGCCATCA

GTGTAACATATATGTTGTTACAAGACTGCAGTCTTAGGTGAATATTAATTCTTCCAACGGTTCTTTTACGAATCTCCCAATAGTTTTCAGCTTGAATAATGCAGTTATTGTTATCAGAAATACCACCAGGAATGCCTAGACCAATTATTTTTCCCAAATTTATCTATCTAACTGGCTGTAGAAATTCCTTGATTTTTTTACGAAATCAAT

TAGAACCTTTTGAATGCTGTTCGTTCCTTATTTTTTTAAATTGCATGTTCCGTTGTATTATTACTAAGAAATTCTTAAAGTGGTTTCTCCAGACTTTGTCGTATGTTCATTGAGAAATACGACCTTAATCTTAAAAGATTGTTTTATGTTTTTGTAGTATCATGGGTAGGACAATCGTTTTTGCTGGCCTACCCAGGTAGCTGTGTGCGT

ATGACTTGTTCTACCTATCCTATCCCCTTAAAGACCACTGATTTTTTCACATCATTGTTAAGAAACGGCCAGACAGTAATATCATTCTTGGTTGGTATGTAACGAATGATAAAACAATATCAGTAAAACGACATCCTCTCGAAGAACAGAACCGAACCATGTCCAAAGGATCGCTTCAATATTTGTGGTGAGCTGTTGAATTACCGAAGA

ACAGTAGGTTTCATTTGTAATTGCCCCAACCACCAGCGCAAATATGTGATAAGCTTGTTTTTGTGCATTTGATTTCGGTGGACATAAGTGCTTCTTTCTGTTAGTACACATTGAAAGGATCCGTATCTTATCGCTATAAAATGGGATTTATAGGCACCCGCGCAGACTGATTGGTAAGCCCCCCAAAAATATCTGTAAAATTATCATCTG

CCTAGTCTACCACAATCTCGATAAAGTTATGGCATAATTCAAACTGGTTTAAACCCAAACGATCAAATCCGCCTTGAACCGACGAATGAACTGGTTTGGAGAATGAACCTCTGTAACAAACTATCATAGTGTCTCAAAAGAGGCTTCACCGTTGTTCGGTCGTTATTTGCCAGCCGTCTTGAACTTTTTCGTGCTATAGGTATCCGTGGT

AGGCGGGTACCTATATGGTTAATCCGACCGATAGAGTTGCAAAGAATGCAATCCACAAGCCGGTCGGCGTTATGCCGAAAGCAACCTAGAAAGATAGTCTGCGTAAAATTGACATGCATTCTTGAAATATTGCTCTCTCTTTCTAAATGGCGCAAGATCCGTCGCTGTGCATTTAGGACATCTCAGTCGCCTTGGGCTCCCGTGAGGCGT

GCTTGTCAATGCGGTAAAGTGTCACTGATTTTGAACTATAACGACCGCGTGAGTCAAAATGACGCATGATTATCTTTTACGTGACTTTTAAAGATTTAACTCATACGATAATTATATTGTTATTTCTGTTCTGCCCGTGATAACTTATTATATATATTTTCTTGTTATAGATATCGTGACTAATATATAATAAAATGGGTAGTTCTTTAG

CGGACGATGAGCATATCCTCTCCTGCTCTTCTGCAAAGCGATGACGAGCTTGTTGGTAGGAATTCTGACGGTGAAATATCAGATCACGTAAGTGAAGATGACGTCCAGAGCGATACAGAAGAAGCGTTTATAGATGAGGTACATGAAGTGCAGCCAACGTCAAGCGGTAGTAGAAATATTAGACGAACAAAATGTTATTGAACAACCAGG

TTCTTCATTGGCTTCTAACAGAATCTTGACCTTGCCACAGAGGACTATTAGGGGTAAGAATAAACATTGTTGGTCCCAACTTCCAAAGTCCACGAGGCGTAGCCGGAGTCTGCCACTGGGCATTGTCAGATCTGTCGACATGCCGCCGGCGTCGAAACCCGCTGACGCTGCCCGCGTATCCGCACCCGCCGACGCCGTCGCACGTCCCGT

GCTCACCGTGACCACCGCCCAGCAGCGGTTTCGAAACGAGGGCTTCCCGGTGCGCCGCGCGTTCGCCGGGATCAACTACCGCCACCTCGACCCGTTCATCATGATGGACCAGATGGGTGGTGGAGTACGCGCCAGGGAGCCCAAAGGGGCACGCCTGGCACCACACCCGCGGCTTCGAGACCATGACCTACATCGTCGACTAAGATACAT

TGATGAGGTTGGACAAACCACAACTAGAATGCAGTAGAAAAAATGCTTTATTTGTGAAATTTGTGATGCTATTGCTTTATTGCTTAACCATTATAAGCTACAATAAACAAGTTAACAACAACAATTGCATTCATTTTATGTTTCAGGTTCAAGGGGAGGTGTGGGAGGTTTTTTTAAAGCAAATGAAACCTCTACAAATGTGGTATGGCT

GATTATGATCCTGGCAGCCGCTGCTTGGCTACAGCTCGTCCATGCCGAGAGTGATCCCGGCGGCGGTCACGAACTCCAGCAGGACCGTAATGATCGCGCTTCTCGTTGGGGTCTTTGTAGGGCGGACTGGGTGCTCAGGTAGTGGTTGTCGGGCAGCAGCACGGGGCCGTCGCCGATAAGGGGGTGTTCTGCTGGTAGTGGTCGGCGAGC

TGCACGCTGCGTCCTCGATGTTGTGGCGGATCTTGAAGTTTCACCTTGATGCCGTTCTTCTGCTTGTCGGCCCATGATATAGACGTTGTGGCTGTTGTAGTTGTACTCCAGCTTGTGCCCCAGGATGTTCTGCCGTCCTCGCTTGAGTCGATGCCCTTCAGCTCGTGCGGTTCACCAGGGTGTCGCCCTCGAACTTCACCTCGGCGCGGG

TCTTGTAGTTGCCGTCGTCCTTGAAGAAGATGGTGCGCTCCTGGACGTAACCTTCGGGCATGGCGGACTGAAGAATCGTGCTGCTTCATGTGGTCAGGGGTAGCGGCTGAAGCACTGCACGCCATGGAATCAGGAGGTGTGGTCACGAGGGTGGAGGGCCAGGACACGGCAGCTTGCCGGTGGTGCAGATGAACTTCAGGGTCAAGCTTG

CCGTAGGTGGCATCGCCCTCGCCTCGCCGGACACGCTGAACTTGTGGCCGTTTACGTCGCCGTCAGCTCGACCAGGATGGGCACCACCCCGGGTGAACAGCTCCTCGCCCTTGCTCACCATGGTTGAAATCTCTGTTGAGCAGAAAGAAACGAGGAAACGCTTGAGTAATTGGTTGTAGAAATGCAAACTCATTTGATATTGATTCATTG

CCTTTGGCTTCGAGCACATTTGACGGGTTTAAACTTGTTTTGCTTTGTCTGCGTTTGCAGTCGCAGGCCAGAGTGAAAATATACACTTGAAGGTGATGACGTCACAGCAACGCCCTACTTTAAATGAAAAATTAACTGTTTTCGACTTTGAACTACGTAGTTTTTGAAATTGCGTATCTTCAAGTTTTCCCCGGAATTTCCCTCAAAGGT

TTTTCTCGATATGTGTTAATATTACCTTAATGGGTAATTACCATCAAAATATTTATTTTAGTATGTGACGGAGCAAATACGTTATTCTTATTCTAGAAATTTAATTCAATTAGTAGCGATGATTCAACGAAATATGATTATCGCTGTGAATCACAATTAGGTTTATCAATGATGATGAAACTGCGTTGCAAATTTTCACTAATCCTCAAA

GCTCAATAGTCGCCATCTTGGAAATGGTTTGCTCATTCAAAGACAAAGAATCATTCACCAAACTAGTTTTCGCTCATAGCTATAATTTCATCAATTAATTTACCTACCTTCACTAGAAGATTCCCATCACCGAAATACTTTTCACCAATTTAATAGTAATTGTCCTTTGAATAAAGCTGTTCACTCTGAAATTTTCTCCTCTATGGATAC

TCTTTTCACTAGAGACTTCACTTCACTTGCACTGGCACTGCTTGGGCCGCGTAATGTTCACTCACTAGGAAACGTATTCGATTGAGCTGGTTTCGCCCCGCTGCCGGGGGCGTTTTATGAACTGTCGTAGTGGTGGTTGTACTTCTAGAAAATTTCAGCAATACATTCATACATATACCTCTGTTCGGTTGGATGGCTCTATATCCGATC

GAATATACGTTACCATCCTGTGTCTGAATGGACAGCAAAACGTGCTTGTGTCTGTTAGTCGTTCATTCTGTACAGCAGACGATGCAGTTCAACTTCTGGCAAAGACGTCAATGTACCTACCCTTCGTGTATATGGCATAGAAGATGTGCGAATGCCTTTTCGATAGAGATTTCCTGATTTGATCGACAATTTCGGTAGCTAATCTTAGCT

TCGAATGTAATCTGATAACCAAATCCAGAGAAAAATCAGTAATTTAGATTTCACTTGAATCATTCTAATTGCATCATTTGCTGTATTAGTGCAGTCAGCAAGTGACGTCAACCCTTCTAAATCGATATACTTCTGAGGAAGCTTTCTTTCTTGTCTGGCTCAGCTGGTGCCAAGGCAAATTATAATTGATTCAATGCACAAGCTACATGT

AAAGATATTAACCATTGTGGGAACCGTGCGATCAAACAAACGCAGAGATACCGGAAGTACGAAAAACAGTCGCTCCCAGGCCAGTGGGAACATCGATGTTTTGTTTTGACGGACCCCTTACTCGTCTCACTTATAAACCGAAGCCAGCTAAAGATGGTATACTTATTATCATCTTGTGATGAGGATGCTTCTATCAACAGAAAGTACCGG

TAAACCCGCAAATGGTTATGTATTATAATCAAACTAAAGAATGGACACGCTAGACCAAATGTGTTCTGTGATGACCTGCAGTAGGAAGACGAATAGGTGGCCTATGGCATTGTTGTACGGAATGATAAACATTGCCTGCATAAATTCTTTTATTATACAGCCATAATGTCAGTAGCAAGGGAGAAAGGTTCAAAGTCGCAAAAAAATTTA

TGAAAACCTTTACATGAGCCTGACGTCATCGTTTATGCGTAAGCGTTTGGAAGCTCCTACTTTGAAGAGATATTTGCGCGATAATATCTCTAATATTTGCCAAATGAAGTGCCTGGTACATCAGATGACAAGTACTGAAGAGCCAGTAATGAAAAAACATTACTTACTGTGCCTGCCCCTCTAAAATAAGGCGAAAGGCAAATGCATCGT

GCAAAATGCAAAAAGTTATTTGTCGAGGCATAATATTGATATGTGCCAAAGTTGTTTCTGACTGACTAATAAGTATAATTTGTTTCTATTATGTATAAAGTTAAGCTAATTACTTGACACATAATACAACATGACTGTTTTAAAGTACAAAATAAGTTTATTTTTTGTAAAAGAGAATCATTTAAAAAGTTTCAATTTACTTTATAGAGA

AGAATTTTGAGTTTTTGTTTTTTTTAATAAATAAATAAACATAAATAAATTGTTTGTTGAATTTATTATTAGTATGTAGTGTAAATATAATAAAACTTAATATCTATTCAAATTAATAATAAACCTCGATATATACAGACCGGTAAAACACATGCGTCAATTTTACACATGATTATCTTCTTGTACGTCACAATATGATTATCTTTCTAG

GGTTAAGAACTGCGATCACAAGAGTAGCCAGTCAGTCAACATTGAAATGTTGTCTCTTTGAAGACAGGCAGTTGGTCGCAAAACTTACATCCCCTTGCTTGAGCTGTCAAAATCTTTGAAAAGGTAATGGCAGGAACCACTAGTTTGACGAGATCTTAATGAGTTAGGCACCATGCAAAGGGTAAGCAAAGGTCACCCCTGTTAGAATTG

GGGGAATTTCTCCAATCTTGAACAACCACTCATGTGAAGTTTTTTAGCTTGAAAAACAAATTTGCATTTCTCAAGGCCGTTTTTGAATGTCATTAACCGAAAAGAAGAAAAAACTGGAAACGTCAGGAATCCCATTTTAATGGGGTGTGGGTTCGATTCCCTTCTGCCTTTGAAACTTTTCGTCAGGAATGTTTCTGCGGCTGTGCCACT

AGAGCATGCGTGTACGTTGTCTAGTGTTAAGTTACAGTCTGTGCAGCTAAATGGCTGAAGACGGTGTCCCGTGTCTTTTTAAATATCTAACTTATCATTTCGACATACCTTTGGATAGGTAATTCGATATATTGCCCTTATCGCCTACTTGTCCCATGTTCTATGTAAACCTTATGGACCGCGGGACAAATATGTGTAGAACAGCAGTAT

TTTCACAGGTTTGCTTTGCCAGCGTTGGCGTTAGCGTTAACGTTAGCGTTAGCGTAGTTGCAGTATACTTCGTAGATTGGATACTAGCTATATTCAGTAGATGAAAAAACCGAATTTAGTACTATACCCATTTAATTCCACTAGAGTTTGTATCCTTTGACCAGATACGCGTATTTTCGACCTCAGCTGTAAGGCCGTCTTCAGTGTCGT

GTACTAGACTGTACAGTACACGACACTGGGAACAACTGCCAATTAGTCGAAAATACGTATCATCAAAAACAATAGTGAAATAAATAATTATATTCTTTAAATCCGGGCCTCATCTACCAAATGATACTAAACAGCTCAATTTATTAGCTATATTCAGGTTTTTTTTCTTTGATAAGCTCATCTAGATTTGCTTTCAGGAACAAGGCTTGG

GAGTGTATCCAATCTTAAAGCAAAGTACCAAAACCACATGAATACTGCTATTTGGCCACGCCCATCTTTACCGTAACTGGGATAGGGAAAGGAAATGTTGATGTAACACTTGCCAATGAGGCCACCGACTCAGCGACACCCTCATAAGTGCGCTGCAGGTTGGAGGTTAGGAGGGTTTCAGGTCAGGATTCGCCTGAGAAGCTGGCGATA

GACACAGATCATTGGATGTTTAATGGCTTTTAATTTAATCTCTTGAATGCCGACAATCAAGAGAAGAAACAATATATTATTTACAGTATGAAGAATGTTGTTTCGCGTAAAAATGATTAGATACGTGTTCAACGGTAGTAGTTTTCTGCTTTTTTACCAGCAAAAACAGAATCGATCCCTCAAGGGTCATCAATTAGAAAGAATAAAAAC

ATAATTGACCGACTAAAGCGTGCGAATTGAAAAAGCCGCATACCGAAAAAAAAATCTGGAACGAATAACGTGTGGCCAATTCTACGACACGCGACGAAGAGGACCTATTTCGTTGTTGAAGAAAATTGCCACGTCTTTTTCACTTTCACAAATCTATTAGTGAAAACGATTGAAACTGTCGCACTATTTATGTTCAAATAACTGTGCGCG

GAACCGAAAAAAACGTGACAGTAGGTTCAAAAGCACAACCACCTTTCACAGGTTTGCTTTACTTATTGATATAAACTCATTGTAAAATTGACGAGCAGAGTTTCTTGTATTATGGGATTAGAAATCTTTAAAGATGGGAAAATATATATGCAGTTTTTCGTTTTAAGACGATTTGGATTATGTTGTATTTGTTGTTCTAAAGCTGCGGCG

AAAACCATGAGCAACAAATGGAAAACTTTATTGGAATAATTCTTGAGTTTAAATTCTATGCTGATCAGACATTCTAGAAGAACGCCCTTAGAAGTCTTTGGAAGATACACCGGATGAATTTAAAAAAATCTCCAGAAAGAATTCTGGTGGAATCCACCGAAACAAGTCTATGCAAGGAACCTGCACGAGAATCTGGAATAATTATTAGGG

ACATATTGACAAATACTGGTTTGAATTCTTCAGAAATTTTGAACAGAGAGTCTTGGTGATTTCTCCTACATAATAAGATATACTGCTTGAACACAAAAATCTTGTAACACATCCTAGAAAAAGGATGGAAGAATAATTTGTAAAATTTTCAGGACGATTTTGTTTTTTTTTTTTCTGTGTTAACGAGATTTTTTAGCCCTGGGCAAGTTC

ATCTCGGGACCAACGGCTGGCTACTTCCCTTCAGAAGAGTTGTCACTATAACCTTTTTTGTCATAAGTAACTATCTCCGGGGATGGGATTCGATCCCACTATCCTCGGCAATAAGAGCATGTGTTCTAATTCCAGCACCAGATCCGCCTCCCAACTGGAGGACTTCTTCCGAAAAATGTCTGGTTGAATTCCTCAGAGTTCAAATAGAAA

GTTTTAAAGAAATTTCGATTCAATTCTCTAGAAAAAAAATGAAAATCTGAAATTTCAACAAGAATGTTTATGGAAAACCAATGAAAAAGAATAATACATATTCTCTCAACACTTTTTAATTTTAGAATATTTTTTAGAAAAGAAACCAAACTTATTGAACAATTTTTGTCAGTTCAAGTCTGTTATTCAAGGATGCTGGAAAAGGAGATG

GAATAAAGTTTCGTAATTGTTCTGCCGGTTTCTCTGTATTTATAACACTTTCAGTAACTTCTAATGTGTATCAGCTACCTCATCATCGGCTTCAGTTGCCTTCTGCTTTGACTGGCCTCGAACTAAAATGCTAAGCCTTTCCAGTGTCTGAAAGTCTTAAAGCATTCAGATTAAATTACAATCGACAATTTTTTTTTCTATATTTTAGGA

CCTGAAAAACCTGGAATTATCAGGGAATTTCGTTAAACCTGAAAAAACCAGAATTCTCAGGAGAATTTATTTGAGGCAGTAATTCATTGTAAAATATGTTCAAGACAAAAAGGTTTTCGCGTCAAAATTCCCCAAAAATGTGCTGCTTTTGAGCTGTTCGGTGAGATCTTTTAAGAGATTATGATATCTTATTGACTTCCCGACATGAAG

TTAAGATTTCACGAATAATATTTGATCAGTAATAGATTCACATATCTTCATGCACGTTTTTAAAATTATTCAGAAACTTCAAAAGCTAGAAAACTGATCTCGAGAAAATAATCGGAATTCAAACAACTTTCAATTTAAAATGCCAAACTATGAAATTGCCAGTTCAGGGCTGTTAGTCAGTCTGTTTTTTAGATACTTGGTCTCTATTTT

AATGCTAGTCTCTAAAAATTATTTTATTTTGATGAAAGGTCTCTAAGGTTTTTATTGTTTTATCCAAAATGGGTTCTAAAGTCTCTTTTTTTTCTATATTTTGTTCATGATTTTGATTAATTATGATGATAATGTAAAATTATGCAGGCTCCAAAGAAACCAGGGAACTGTTACGAGACCTTTAAACTATTTTTCAAAATTCCCTCAAAC

TTCTCGTAGAAAAGCTTTGGAATTATCGTAGTGGTTTCTCCAGATATTTAATCTGGAGATACTTTCAGATACTCCTACACGTATCTCTCTAGCAATTCTTCTAGAATTTCCTCTACCAATTCTTTCAGCGTTTTGGAAAAGAATATCTTGAGAAATTTCTCCATGTTAACTCAAAGAAATTGTCAAAACTTCAAAAAGCATCTCCAGTAA

TTTGCTCAGAAATTACTGAACTTTACATACACTATCTTAGGATTTCTGTCAGATATTTTTCCACTAATTCAACCGAATTTTCCAGTTGTTACACTAGGAGGTCTCCCAAAGATTTTTATAATTTTTCTCAAAAAATTTAAGGGGATTTTTCTTTTTACGGTAACTGAAAAAATCTCAACAATTCCTCAAATTCGAGCCTATGTTTCATCA

TTGAATTCCTCTACACTTTTCTAGGAATTTATCTAGAATTTCATCGAAGAATTATTACTACAGTTCTTCCAAACATATCTTAAAAAAGATCCGTGAAATCTTCTTCTTTCTGGCATTACGTCCCCACTGACAGAGCCTGCTTCTCAGCTTAGTGTTCTTATAGGCACTTCCACTGAGAAGCTTACTATACCAATGACCATTTTGCATTCG

TATATCGTGTAGCAGGTACGAAGATACTCTATGCCCTGGGAAGTCAGAGAATTTCAACTCGAAAAGATCGCTGACCGGTGGGTTCGACCTCAGCTTAGTCATGCTGAATAGTGCGCGTTTACCGCTACAGCTATCTGGGCCCCTCCGTGAAATCTATTGAAGTTTATTTAAGGTTTCTACTAATTAACGCAGCATTTTCCAGAAAATCCT

CCTACTGCTTATTTAAGAATTCAGTCCGACTTTTTCCAAGGAATTTCTTCGAAAATGCTGCTTGATATTAAAGCCTTGATATTGTTCATTTAAGAAAATGGACACGTGGTTTTACAGGAAACTTTATTTTCGAACAAAATTGATGAGTTCACCTAAAATACATTTCGTAATTTCGATCAAATTCACATACATTTCATCATTTATTAGGCA

TTCCTGGGAGAATGCTCCTATTTTCTCTTAGAAACCTCCATAAAGGTCACAACCTTTTCGTTGCCATGCTTTTCGTTGTGCTGCTGACCACAATTTTCTTGACAAAATCAGTGGAGCTTGTTTCTCTTCCATCCTTCTTAAGATGTCGTTTGATTACACTGCGAATTACGTTTTGTCTCAAGCACTAAAATACCGCTATTTACTAAATTA

AGACTAGCTGATCCATTGCCGTTTTAAAATATCATGCTATATCTAGTTTTGAGATATTGACTGTTGAAATGCCAAATTAGACTATTTCGTATGCAAGTTTGCCAGCTTATGGCAATGTTTTCGCTTAAGTTGCCACAGGAATTCAAACTTCAGGTATTTAACAATGGTTATCAACAAGAAGTTTATAGTACTTTCGGTTTTCAAAAGTAT

GTTTTTATGATGGTTTCGAAAAGTATTGTATTTTATATAAGAGAAACGAGTTTTGTACGAAGAATGCAAGCATGTCCGGAAAATCGATTTCATCTAAGTACGATCGTGCAATTTCAACCCAAATTATATTTGAAATGAAAGTTGAAGTAATAGTTTGTGCTGGTGATTTTGAGTTACACATTTTTGTGATAAATCACACTTTAAATTTTA

TAAAAACTTCAATGACAACAGTGTTGCTTCTGGCTTTGGCGACCTATACAACTAAAAAGAATTGTGACGTGCTAGAACATTTCTGAGAACAACGTAGTGCAGTAATAAGAAATCTACTAGAAGACACATAATTTGATCCTTGAGACACGCAAATATGTCATTTTTATTTACGCTAGTATTTATTGCCCCGATATGTTTCAATGAGGGCGT

AGTTCTAGGTAATTTGATGGATTCGGTCATTCTACAAATTCGATTTAGTAGTAGCTGGGCTTATAGTGAGCCTATGCTGGGTCTGTAAAAACAATGGAGCATGGTATGCTTTCGGTAGATGGGCTTTAGAAGCCGTTTTTTAATGCATTCAGTTTCTGTAATTTTTTCGCCGTTGATTGAACCCGTCGTGAAATATGCAGTGTTTTTCAT

ACCGCAGAACAAATTACTTGCCATACCAAATCATTTTTTCTCAAATTTTTCTGTTCGGATGTATCGTACGTCGTCAGGAATATCTGTTTTTCATACAGCGATGAAAAATTCAATATATGGACGCTTCGCGATTTTAGCCAAAAAAAAACGTCCGAAAAACAAAATGTCGAATGCCAAAGCTTCGAAATACAAAATGTCGAATGCAGACGA

CCGAATGCCAAAACGTCGAAGCGAGAAAAAAAAATCAGAATGTCAATAAATGCTAAACGCCAGACAAATGCTAAATAAGTTTATTCGATTAACCCCTGTACTAGCAGCGTCATTTTTACAACAACAATATATTTAAATGATTTTTGATAATACAAGAAACAACATTATTTTTCTACCAATATTTTTAACAAGTTCTCAAAAGACTTTTCT

TCTCTTAGAATCTGCGTCGATTTCGATTATTGGTAATCTAGATCGAGAATACAACAAAATTCCTTGGGGAGCGTTGGCATGACCTGCTTCCTTTTTATTTTTTCTCTTGTTCATTAATTCCCAAGTAACCATGGAGCACTATATCATTGCTTATATAAAGCAGCTGCATTGCTTAAATAAGCCTATCATTAAAAAGTAGTTTAAAAAGTG

GAAAAAGGTCCTTGATATTGTCAGACTAATGCAACTAATGCTTATAAAGCAACCAAGCAATTCATTAAGTAATATTAGTGCATTGATACATTTCTCGCGTAACATTACTAAAGGACCTATGTACCAAATGAGAGATTCTCTCCTCTCTCGTTCTCTTTCGATTATAACAGTGGAATACTGAAGATTTTGGGAAGTTTTTTTCACTATAGA

TCAAGCCAATTTCCTTGACTAGCGTTTCATACACAAAAAACGCAACAGAAGAGGTTAATGTGACTCAGTTATTAATGAAGAGAAAGTAAACAAAGAGAGAGCCTCTCAATGTTAGATAGGTCCTTTAGTAATGTTCCATGAGATTTGTAATGTAGCGTTAATGCTCTATTGCCTGACAATCTTAAAATTTGTCGAATAAACAATGTCGCA

TCAGTTTTTGTTGATATGCAGTAGAATGTTGCAGCGTGCGAAGTAATTTATAATCGCTTGTGATTATGATCGATCGCTTTTGAAAACAATTTTTTGTTGAGGTAGAGAAAAAATAAACGATTTGGTGCAACCGTTTTCAAAAATGAGCATTTATGTTTGTGGTACCACCAAAGACGAAATAAAGACCTCTATGTTTCTTGCAGTTGACCA

GAATTTTTATGGCACATAATGAAATGAGAGCAAAATGTAGAATGTCGTGAAAGTGTACTTTTGCGATGCGGTACCAAGGGACCAAATGCAGTACTAAGTAGTACCAATCTGAACCGACTACCATGTCTCTATTCGTCTCTCTGGTACCCTCTGGCCAGCACGATCATGAAATCAAAAAAAAATCATCTAGTAATTTTAGGATTTCTCGAG

AACAACCTAACTAGGTTGCATGATTTATCAAAAAAGCTCTGATTTATGACCCATTTTTTTATAAATTGACTAAAACAAATAACGCCCAGAGAGATTTTATATAAGGAATGTCGGTCCCCCATAAACATTGAAATATTTTCCAAAACCAGCAATGATCAATATCGACAAATTCTGGAAATAGAAGAATTTTTTGAGAACTTGTGAAAAAAT

GAATTGCAAACTTATCGAGAAACAACAAAAAACTTATCGCGATTTGTATATTTTTGTTTGTGAAAAAATGGCTTCACCCCAATAACCTCTTTTTACGACCGTTATCGGGGTCGTAAAAAAATCGATCGTAAAAAAAATCAGGGTTATAATTATTTGTTGCAAAATGCAACGTCTAGATGCAGATACTGGTTCGAGATCTTCGGCAAAGTT

GAAGCATGGGTTGAGAACTTTCAAAATGGCCATTCAAGTTTCATATTTTTTTGGTAAAATATGGCAACACTGCTCGATTGTTATCAAAAGAGCCAATTTTATGGGGGTGTGATTTGCAAATACCAAAAAGTTTAAAGAATAACGATACCAATGTTCACAAAGTCGAAGGAAGTGTTGCGTGCCATACAATCAGCCAGAATCACAAATGTT

CATGTAGCTGGCAACACTGTTTAAGAAGAATAAAGTTAAGATCAAAACCATCAAACTTAAGCGGAACAGAAAAATATAATGTAGCAACATAGCAATACTCTTCAACTCTAATCCAGTTTTTCATGAAGTTCAATTTTCTTTCTTCGTAGCTGGCAACACTGCTTAAATTGAAATTGCCACCAGGCAGTACAGAGATGTGCATGCAACTAT

TGTTTTGCGGATATCCTCAGGATCCTGACCACCTAGAAAGATGGCGTCTTCGGCAAAGTTGTTCAGTAGCTCAAGGACTATCATTATAAACGCTATGAGGTTCAAAATGTTGCCGCCAGGCGGCATAGTGAGCATGAAACTTTTGTTTTGTGTGGATATCTCAGTTCTGGCCATTTAAGAAGGTGGCTGCTTTGGCAAGAAGGATTGTTC

CCGAAGTAGCTCATGGACTATCATTATAAACGCTATGGGGTTCAAAAATTTTGCCACCAGGCAACATGAGCTAGGCATGAATCTTTTGTTTGCTGATATCTCAGGATCCTGGCCATTTAGAAAGATGGCGTCTTCGGCTAAAGGTGTTTAGTAACAGGACTACCATTATTCTAGCCAAAATTCGAGATTTGCCATCAGGCGGCTTTAGTG

AGCATAGAATTTTGTTTTCCGGGTAGCAGGATCCACAATACCATTTAGAAGATGGCGTCTTCGGCAAAGTTGTTCAGTAGCTCAAGGACTACCATTATTCTAGCCAAGAAATTCCGAGATTTTGCCATCAGGCGGCGCTAGTGGTATATAATTTTGTTTTTCAGGTAGCTTAGGATCCTGACCCTTAGAAAGTTTGGCGTCTTCGAAGCC

AAAAGTTGTTCAGTAGCTCAAGGACTATCATTATAAACGCTATGAGGTTCAAAATTTTGCCACCAGGCGGCGCTAGTGAGCATAGTCTTTTTTGTTTTGCTGATATCTCAGGATCCTGGCCATTTAGAAAGATGGCGTGCTTCGGCAAAGTTGTTTAGTAGCTCAAGGACTACCATTATTCTAGCCAAAATTCAGTTTGCCATCAGGCGG

CGCTAGTGAGCATATAAATTTTTGTTTTCAGGTAGCTTGGGATCCTGACCAGCAGAAAGTTGGCGTCTTCGTAAAGGATTGTTCCAGTAGCTCCAAGGACTATCATTATAAACGCTATGAGGTTCAAAATTTTGCCACCAGGCGAGCGCTAGTGAACATGAATCTTTTGTTTTGCTGATATCTCAGGATCACCAACCCATTTTAGAAAGA

TGGCGTCTTCGGCAAAATTGTTTTAGTAGCTCAAGGACTACCATTATTCTAGCCAAAAGGGGTGAATTTTGCCATCAGGCGGCGCTAGTGAGCATATAATTTTGTTTTCAGGGTAGCTTAGGATCCTGACCACTTAGAAGGTTGGCGTCTTCGGCAAAGATTTGTTCAGTAGCTCAAGGACTATCATTATAAACGCTATGAGGTTCACAA

TTTTGCCACCAGGCGGCGCTAGTGAGCATAAATTTTTTGTTTTGCGGATATCTCAGGATCCCTGACGCTTAGAGAGATGGCGTCTTCGATAAAGTTGTTCAGTAGCTCATGGACTATTATTATAAACGCGAGGTTCAAAATGTTGCCACCAGCCAGCGCTAGTGAGCATGAAACTTTTGTTTGCCGGATATCTCAGGATCCTGGCCATTT

AGTGGCGTCTTCGGCAAGGGCGTTCAATGGCTCAAGGACTACCATTATTCTAGCCAAGAAATTCCGAGATTTTGCCATCAGCCGGCGCTAAAGTGAGAGCATAATTTTTGTTTTCCGGGTAGCTCAGGATCTGACCCACTTAGAAAGTTGGCGTCTTCGACAAAATTTGTTCAGTAGCTCAAGGACTATCATTATAACGCTATGAGGTTT

AAAGTTGCCGCAGGCGGCGCTGGTAAACATGAAACTTTTGTTTTGTGGATATCTTAGGATCCTGGCCACTTAAAAAGGCGCCTTCGGCAAAGTTGTTCAGTAGCTCAAGCGCTATCACTATAAGAGCTGATTCAAAACTCCGCCACCAGGCGCTAAGTGAGCATAGAACTTTGTTTTGCGGATATCTCAGGATCAGGTGAGGATATCGCA

TCTCAGAGGAATCTGAGATATCCGCAAAAACAAAATTTTCATGCTCTGCTGCCTGGTGGCAGGTACAATCTCGCCAGCTTTATAGTGATGGCGCGAGCTGAACAACATGCCGAACGCCATCTCTAAGCGGTCAGGATCTGAACACAAAACAAAGATCATCGCCACTAAGCGCCATCAAAATTTTGAACATCATGGCGTTTATAGTAGGTG

GCAGGCTTGCTTCTTTGCCCGGCTCATCTTCTCCTGAGATATCCGCAAAACAAAAATCATGCTCATAGCGCCGCCACCAGTAACAAGAGTACAGAACCATAGCGTTTATATGATAGTCCTGACCATCTTTCAAGTGATCAGGATCTGGGCTGTTCGCAAGGTAAAGTGCTCACTGGCACTACTTGACAAAATTTTGAACCTCATAGCGTT

TATAATGATGGTCGCCAGGCTATTAACGAAATTCTCCAACTCCCAAGTGTCAGGATCTGAACCTTCCGGAAAAGTAAAATTATGCTCACTGGCGCCGCCTGATGGCAAAATCTCGAATTTCTTGGCTAGAATAATGATGGTCAGCGAGCTATTCTTTGGTACCTCGTGGTCAGAATCCTGAAAATATCCGGAAACAAAAATCATGCTCAC

TAGCGCCGCCTGAGTAACAACATCGTAATCTTTTGGCGAATTATGATACTCGCCGAGCTGCTGGGCTAACGCCGAAACTCATCTCGTGGCCAGGATCGCAGACTATCGCAAAACAAAAGTTTCATGCTCGCTAGCACCGCGCAGTGGCAAGAACACGAACTCGCTGGCGTTAAATGAAGCAGTCGCGAGCTACTATTCTTTGCCAGCTCA

TCTCTAAGGTGGTCGAGATCTGAATATCAGCAAAACAAAAATCATCTTTCTGGCGCACCTGTTGGCAACATCATATCTGGCTAGAATTATGATACTCATGGGCTGAATTCTTTACCGAAAAGACGCCATCTCAAGTGACGGTCAGATATCGCAAAACAAAATTTCATGCTCACTAGCGCCTTCTGATGCTTTTAAATTGAACTCATAGCA

TTTAAAATGGTAGTCATCGAGCTGCTGAACAGCACCCAAATTTCATCTTCTAAATGTCCAGGATCGATATCGCAAAACAAAACTCTTGCGCCGCCTGTCTAAACCACAGACCTCGCCAGCGTTCTAATGGTCGCTGGCTGCGAACAACTTTGCCGAGCAAGTAACCCGGGATCTGAGATATCCGCAAACAAAAATTTCATGCTCCTGCGC

CGCAGTAACAAATGAACACTCGCTGGCGTTATAATGACAGTCGCTGAACTGCGAACAACACCGAAAACGCCATGGCGGGATCTGAATATCCGCAAAACAAAGTTTCGTAACCACTATGCGCCTGGTGGAAAATTTTGAACCTCATAGCGTTTAAAATGGTAGTCAGCCAGGCTACTGAACAACACGCCGAAAGACGCCATCTTTCTAAAT

GTGGGATCTGAGATATCCGCAAAACCAAATGTTTCATGCTCACTAGCGCTGCCTGGTGGCAAAATTTTTGAACCTCATAGGCGTTTATAATGATGGTCAGCCCAGGCTACTGAACAAGCTTTGCCCGAAGACGCCATCTTTCCTAAGTGGCCAGGATCCTGAGATATCCGCAAAACAAAGTTTCATGCTCACTAGCGCCGCCTAGTGGCA

AGAATTCCACAATTTCATGACCACCATATATTTAGCCCTTGAACTACTGAACAATTTTGCTGAACATCATGATCCTCTATATTGGTCTTATCAGATAATGATCAATTATAAAGCAGTCGTTAATCTGAATTTCGGTCGTAAAAAGTGGTCGGAAAAAAAGAACCCAATCGGTAAAAAACGGTCATAGAAGGGACAGGTGTACTGATAAGA

TTAATAACGAATGCGTAGAAAATTGATAAAAATGTAACATAATCAATAACAAACAATGTTTTGAATAGTTAAAGGCAGTAAAACACATTTTTCCGTGATACCAAAGACGAAATATAGACCTTCTATTCCTAGCAGTTATCCAAAATTGTATGGCACATTAGGTAATAGATCAAAATCTAAGAATGCTGTGAAAAAAGATTATTCTGCGAT

CTGTCAAACTTGGATACCGAGAATAGTGCCAAGAACCCTTTTCTGTATCCGACTACTCTTTCTTAATTTCGTCTTTGATGGTATCGTAATTTCAGCTGGTTTAAGTAATAATTAATATCCTGGGTAGGTGGGGCCTTCCTTCATATAGCTCATTTAGGAAGAAGTTTTCCAAGGATTTTAGAAAGTTGAAGATATACGGAATGTATTTAG

TACAACAAATGTTGTTATTTATTTATTTTATCATGTGCTACTAGACTGGCCTTTAAACAAAAAGTTGTAAACTCAACAGGGCACCCCCTAGATATGTGTCTTAGGAGTAAGGAAAAAACTCTCTCAAAATTTCAACTCATTTAGTTGCTCCCCAGCTAGCGCATTCAATTTGAAGTTTGTATGAGATATTCGTCTCAAATATATTGAAAA

TTGACACTATGTCACTGTTTCATTCCGTATACTGGTTAATCGCATACGTTTGAGCCCAGGTACAAATTATAGTTGATACCTAATGAACATAATTGCAGAAGGTTGTATCTGGATTTAAGCTCATTTTCATTAACCTTTCAGTTGTTGAAAGTTAGGTTTAGATCAGCACTTCCGTACCAATCACATTTTATGCATGTACCTTGTACCGCA

AGCTCGCCCAGCGTCATAGGTGGCTATGGTGCGCTTGAGTAGGTAACCGCAGCTAAGTTGAAGAAATACTAAGCAATATTACCGTCTACTTCGGATAGTTAAAACACCCACTTTCTCAATTTTATTCATCATACAACCTTCTACAATATTGTTCACTATGTTATCAAGTAGTGTTTTTTGACATTCTGGGTACAATTGAACGCGATCAAC

TTAATTTCCTGACCCAAAAGGATATGATGTGCTGTTTCCCATATGTTTGAACTGAAAATCATATTAACTTCAATTCAATTGCGCCAGCTCATGAATGACCAAATGAGCTGAAATTTTCAGGAAGTCTTCCTAACCCTAAGGAAAAATCACCAGGATGCCCCGTGAAACTATTCAACTTTATTTTTCTTCCATACTGAGCTGGGCGAGTCT

AATGTGCTACATACATATAACAGTTGTCAATTCAAAGCCATTCACGTCTTGCATTAGCTCGCAATAAAATAATTGTGTTATAAATCGAATTACTATCAACCAAGCTGCAGTTTTGAGTATTATTATAATCTGCTTGATGAACAGGTATTTTGTTTCCATTTATGTAAATTTCCATGGTCTAACTTGTCCAAGCATCTTTATGCAGGGTAA

AAGTGAGGCTAGAAAAACAACCTCAGAATCAAAACTTTTCCAACACGTTTTAAACTTTCCTAGTTATAGGGACTAGCACTAAGACAAAAAACAAATGGTTTTTCATCCAAAGAGATCAATCTCAAATTACGTAATTGTTTAAAGAGAGCATCCATTAGAAAAATTACTTTTCTATTCGTTATTTCTACCAAATTCAATAGCTGCTTCAAT

AGATAGTTATGTTCTAATTACGTCAATCATTAAAAGTATACGTGATACTTTTTAATTAATAATTTATAAAGAATGATATATTTACGAAAATGTTTTATTCCTCTCTACGCTTTTGAGGCATTTTCAGTTAATATTAAGTGCGATGTGACATACAATTAAATGTACGTTATCTTCCCTAAAAGCGTTACGTATTTATGGCTGAACCCTTAT

AAACATCAAATATAACTACTCAAAATTGAAATTTTATGATTTATTAATCCTATTGTGCAATGGTTCACTTAGTGATGTGGATTGACGCATGAAACTGGATGGATCACTTGCCCCCGTTGCTTGATTAACCGTCTAACTTTCTTTTAAGGTTTCCACAATTCGAAATTATTCGGTACATTCATGCAATATGTTGCCAAAATCTGAACGTCT

AGTGGGATTTGCAAAATATTGACACTGTAATTATATTGTACAATTTGTTTGAACTATCGTTTTTTTAGACCATACTTGCCCGAGGTACCTCATATGCATTATTATTGCAAGCTTTTATCCTTAGTCTTTCTCATATAGAAAATAATTATTGGATCACCTTGCGTTTCAAAATGAACAGTGCATTAATCAAGATAGTCAAACATCCTATCA

GCCAATCTTCCCATTGCCTTGTTTACCTTGGTTGTATTAGTCATAAATTACTCATTGAAACTTCTTATCTTCAAATCTCATTGCAATTTTGCAATCATTACGGTGTAGAGATCTCTTGATAATCGTTAGTTTTGTCTGCAAATTACGCTATTCTAAGTTCTTTGCCACAACTGCACAACGGCCTTTCCAGAAATATAAAGCACAATGTTC

ACTCCTGTTCCGAAGTTCAGGTCTTCGGCTGCTAGCTCAAACTATGATAGTTCAAGTAAATGCAAGTTCATACATTGTGTTCACATCGTGCATATGAACAATTACCAAGATTCACGGCCATATCAAAGCGGGTAAACTTGGGAATTTAAAATAGCCGTAGCGACAATCGCTCGAATGTCCATCAAGTCCATGCTGGAGGTGGTGGGCTCC

GAGATCTTAGGTCGGAAAAAGAATCTTTTTAGGTTGAAAATTTCTGGTTGCCGAGGCATAGCTTATTGTTGTACTTGCCACACATTATATGAATGCAAAAATTAACAAATGGCTAAAAACTGTTAAACCATTAAAGTGCTTTTAATTGAGAAGCAGGTTATGTGCTAGTTGGGATGTGATGTCGTATGAAGAACCTCACGCACAACACAA

TGTTCTTGATTGTTATTTGAACACCAACTATTATTGCCACGTAAACTCTGATAGTTGATAGTGAACAGCAAAGTATAGATAATTTAAGTTCAAATACCTCTTACCTTCAAGTTAGAGTTTGAATAGTAAATGCTAAAACAGTGGCCCATTAAAGCATGGAACAAATAACTTAAAAGATAGATGCTGCCCATTTCTACAGTAATACATAAC

ATATCTAGCGATCACTGACCAAGATAACCCAGTTATTAACATTATTGTTGAAAGCAGTCGTTGTTTGTAGTGGACGATCATCGTGTATCGTATGAACGAAGAGACAGTGGTTCTTATAATTATGCCGTCAGCTGTTACGTGCGACGCTTCGCTGGCGTTATTTACTAGCTAAGTAGAGGTACCGTAAAACGGGGTAATTTTATTAGTATT

TTTTGAGGAATCAAATCTGATTATTTCTACATTCTGTTTTGGAGATTTATGTATTATATGTTTAAAACAAGTACTACGTTATCATTATTATAAGATTCAGCTTCACTCAGCCAGTGGTTAGAGTCGCGGCTACAAAACAAAAAACCATGCTAGAAATTATCTGGGGTTTTAGAATCCCGGTCAGTCCAGGATCTTTTCGTAATGAAAGAG

AATTTCCTTGACTTCCCTGGGCATAAAGTATCATCAATACTTGCCACACGATATACAAATGCGAAAATGACAACTTTGGCAAAAGCTCTCCAGTTAATAACTGTGGGGAAGTGCTCATAAAGAACACTACGCTGAAAAGCAGTATCTGTCCCAGTAAGGACGTAAATGATAAGAGAAGAAGAAAGAAGACTACGTTATCAATTACAGATG

AAAGATTGCATAACGATTGATTTTGTATGTACATATATTTTGGTTTATAAAGTACTTGACAAAATATTTACAGCTTCTCCGATTTTCCATATTACAAAAATGGAAAATGTGTTTTAATGATCTCAGCTGCGTTGCTTGAATCTTTCAGCAATTTTTAGTCAAATAATCTGTGCATTTGCCTAAAGATTGAAATTAAAAAGTTGACCAGGA

GACTTTTGATTATTTTCATACTTTTGTCATCTGCAAACGCATACATAATTTTATCCGTCTAATCAGTGAGTTTGCTGAACGATTATATGCATGAATTGGCGTAAACTTACCAGATCACTGTCATAACAGCCTGGTATACCGCTAAACTATCAGTAAACACTAGGCTTAAAATCAGGTGTTTATTCCGGATAAGTTAATGTTGTGCTTTAT

GTCAATCCGTTTTCCTCTTTTTAATATTACCCATCACCGAAATTTACCTATAGTCATCTGTGTTTTAATCTATAAAAATGATGGAGATAAAGGCTAATAAAAGTGACCGCGGTGTACTATATCGAAGGTGACGTATTCCGAAAACTGACAACTACTTGACGACGTCATTCCTATCCATCTCGAACAAATTTGCTTGAACAAATGATCAAG

TGGTCCGGATGTATGGTACGATAAGAGCGACGTCACATGTTTACAACCAAATTTGATGTTTACACTATTAGGTAAAGAGCCTGCCTTGTTAATTTTTTATGCCGTGGGTAGAGACCTGAGAGAGAGAGGAGACATCTCACTGACAACTGGCTTGTTTCTTGGCTTCTTTGTTTTGTTTCTCATGTTTGGGTTGTGCACAATGGTTCGCAG

AGCACAAACTGGGTCAAGCCATTTTCACTATCCCATTTGTTATCAAAAGATCATAAAATAATAAAAAAAAATCAATAGCAATTTGGGTCCAATCGACAATATTCAAATATTCAACGTTTTTTATAATACGTGCTGAAAAAAAACCTGAATTTCAATCACATCGGTTTTGAACATTGCAGTTTTTCATGTTCTTTTTTTGCCCTTCAGATG

TGCGGTTTTCCAGTGACAGTTGATGACAGGTTCCTTTTGGGAAGGCTCTCGCAATCTAGAACAGCTGTCTTCTCTCTCTCAGATAGAGACTAACAATTTGTTTCATAATTTAAACATGTCAAATAGACTGGCCCTTAAGCAAAAAAGTTGAAAAAACTCAACAGGGCACCCCTTAGATATGTGCCTTAGGGTAAGGAAAAGCTCTCCCAA

TTCCAACTCATTTAGTTGCTCCCCAGCTGGCGCATTCAATTCAGAAGTTTGTATGGAATATTTGTTTCAAATATATTGAAAATTGACCCTATGCCTGTTTCGTTCCCATAGACCCCAAGTTCAATCGCATTCAATTGAGCCCCAGATTGACAAATTCATTAGTTGATACCCTAATGAACATAGTTGCAGAAGGTTGTATCTGGTTTAAGC

TCATTTTCATTAACCTTTCAATGGTTGAAGATTAGGCTCGAATCAGCACTCCCGTACAATCACATATATGCATGCCTTGTACCGCAACTCGCCCAGCGTCATGAGTGGCTATGATGCGCTTAGTGGCTACCACTCAGCTATGTTGAAAATACCAAGCATTATTGCAAATCTACTTCAGATAGTTAAAACACAAACTTTCTCAATTTAATC

ATCCTTCTTCTAAATATTGTTCGCTATGGTATCAAGTAGTGTTTTGACATTCTGGGTTCAATTGAACGCGATCAATAACTCTTGACCCAAATAGGAGAGATGATGTTCTGTTTCTCATATGTGTGACCTAAAATCCCATATTAACTTCCAATTCAATTGCGCCAGCTCACAAATGACAAATGAGCTGAAATTTTGAGCAAGTCTTCCTCT

AACCTAGAAATAATCCTGTGGTGCCCCGTGGAGATTATACAACTTATTTTTCTCCCCCTTTGAGCTGGGCCGATTTAAAATTGTCCAAGAAATTTCTTCATCGCAGCGATACAAACCAGAAATCCACCGCCTACTTCATCCAAAGAACAAAAATCATGGAAGCTTGATCGAGGTATTAATCTTGTCTAAAAACATTAATATGATCTCTAT

CAACAGTGCCTTGTCCCTAGCTTTTTAATTCCAAAATTCAGAATAAACAAATTGCTGTTTGTATGTCCGAACTGATGTATGCCCCTTCTGGTACTAAACCGAAGAAGAGCGCCTTTTCGCTTTTTACTACACAGTAAAAAATCAGCACGTTCGATTGAATAGATTCAATCACTTGACTCAATTCAAAGTACAAATCACCATGATTTGAAG

AGTTTCAACTTTAGTTTGCTCTGTCTCTAGAACTAGACTCTGAAGTGAAAAGTCGTTGATTTTGAACATCATGTCTTTGTATGAGAGCAAACATTGACAGCTCGTTGTAGAAAGTGATTGAGATCAATTCAATCGTTTCAACAGATGCAGTTAGTGTCGAGGTAAGACACTGGACTTGCCTCCGAGGATTGGTGGTTCATATCTGGGTCA

GGCGGATATGTTTTATTTTAACTTTTTTTATTTCAAAATCAAAACATGCAACGTTGAATTCAAGTGCTGTTACCTTGATTTGTCCAAGATTCTAACAATTAATTAAATCCAAGTGCAAAACTATTGATAGCATGGTGCCTATTTTTTACCGTGTAGTGCCTCTGCAAGTTAAAACTAATGCCTTGGAAGTAGTGCGCTGTATAGATCATC

AGATGATCTATACCAAGCGCCACTATATCCGACATTAAATTGCTACATTGCTAAATAAAAAGCGAAGCGCTATTCGGTATAGTGCTAGAAGGGGTATATTATTAGGTATTAGGTTCATTAGGTTCGTATTGGCGTGTACATCGATTATGCAGAACACGAGCAGTTCAGACAGCAGATCGATTTGAAAAGAAATGTGCACAATTTCTCCAG

AATGCCAACTTGAATATCTGTGCAGATTTTTCCATTACGGTTCCGAAGGAATAACAATAACGATTTTTATAATTCACATACCTAAACAAATTAAAGGAACAGCTCCATGATATCTTCAGCAACAAAGTTGTTGGTTTTGAGATGAACGAGTTTTCTGAAAGGAACAGATTTTTTGGTAGATTTTTTATTAATAGAAAGCTTTTGATTGTT

TTGTCAAACCTTAATTTCCGCTTTAAGTAAAATTGTTATTACTTTAAAATTTGCCCGATTGGATAAACTATTCAAAACATATGTTAAAAATTCAAGGTTTGTCTACAACATGCTTGTGGAACATTTATTCAAATCCGATCCATAAATAACTGGGATCTAGTTTGCAAAGTTGATCATTTTTGTAAAATCAAAAGTTTATAATGTTGGGCC

AGTGCACGACGAACAGTTCGCGAAACAGTACGGAAAACCGGAGAACGATTTCCCAATCAATTCAGTGCGAAAGTGAAAACATGCGTACAGTTTAACAGATTCAAATATAACTAATTTATTTATCTCAAATATGATCAAATGTGTAGTGCTTCAACAATTCAAGTCAACAATAGTATATGTAACCAAGTTTATATAAAATTTAGAGAGATT

TAGTTTGCCAATATTATTTGGATGTGGGCCGTCAGCGAATTCTGCGCCGACAAATGTATTGACCAATACTGTATAATCGGATTTTAAATTTTCAATATTTAAATATGACAATTTTGAATAGTTTCAAATGTTGTAGGTCCCGCATTATCAAAAGACACCCCGTTTCACAGTAACTAAACAGCAGCTGTTTGCATCGATCGATCAGTACGC

GCTAAGCACATGTTACCCATCCATGCCGATTCCGTCAAACGCGTCTACTTAGAAATACCATTTCGCTTGTAATATTCCAGCGCTCTTCAACGCCGGATGAGATACGTAGGGGTAGCATCATTGAAACCTATACGATTTTTTTTGCTCAGGTCGTCACCGTAATCATCAAAGCATTTTTTCGCGCCTTGAGTAGCCCTCCGTTCGATTTTG

AGATGACTTTTCAAAGAGCCACGACGCACATAGAAGACACCTTGGCAAGTGTGCAAAGCAGAGCTATTCAAGCACTCATGACACCGGCCGTTGCGGTTTTAATATTCTCATTGAAGCGGAAGAATTCAATCGACCTGCCTGCACGGCGGCGTGGTGCGAACCTTGGCTAGTGTTGCTGTTTAATATTCACGCAAAGAGTGGTTGCTGACT

TTGGAGGTGTTTGCAGCCGATGACCAGTGCCGTGCAGGGACAGAAGCGTGTTCAGAGAGAGAGGGGGGTGGCAGATTTATCTATTGCCTTATTGTTTTCTCTAAATTACTTGTTATGTTTCCTTTGGGTCGTTATACACATTCTTCTTCTTTTCTTCTTCTTAGCATTGCGTTCTCTTCGAATGCTTGCTCTTCAACTTTGGTGTTTATG

AGCATTATAACAGTCATGACCAGAGAACTTTCTTTGTCAAGTAGCCATTTTCGCATTCGTATATCGCAGTGATTCCCAAAGTAGGCGAAATCGTCCCACAAGGGGGCGATTTTTTTTCATGTCAAGGGGCCAAAATTTGTAAAATGGATTTTGGGGGCGAAAAAATCTGAAAGGGGGCGAAAGTGTGAGAAGCGAGGAACGATTCATAAC

ATTTAGAACCTCAATTTAATAGAGGATTAAACTGTAATAAACTGAAGATAAGTTGATTACATACTAAACTTTTCCCTCTAATTGTTTGTCTTAACATGCTCATTTATAATCACGATAGGCATCTCTTGGACAAACAATGACAATGATCAACTTCAGTTTTATAAGCATTCAATTTTAAAAGCTTTTTGTTCCTTAAGAATTATGAGTTTG

AACTGAAATTTTCTCAAAATACGGTGTATTCAGATCAGAGCTACCTATTTGTAGTCAAAAGTTTCATTAAAAGTCTTTCTTTTGAAGTCATTAAAAATCTTAGAAATTGTTTCGAAAACGAAAAGAAATAATTTGATGTTTTGTTGTAAGAACGCTTTGAATTCGACAGCTAATGTTACGATTATTGACATCCGGGTTATCACAGGCAAA

GTTTTTAACATAACTTGTTTTTACATATTGTTACGATAATTTCAATAATTACTTTTATTACATTTTTATTAACTAAATTCAACCAAACAGCAGTAAAAATCAAAGCATAGATCATGCAATTAAAAATTGATTTTTAATGAATATTGTAGTATCTCCAAAGGTACCAGAAAAAATCTGGATTTTTACGTAACATTGCAGTGAAAAGATAAT

TGGTTTTTTTACTAAATATTAGCATTATGAAACTGTCAAACATTTGTGCATATAATTCATGAAACTTTTTCTAGAAAATTCTTAAGATAATCGATAACTCAACGAAAATAACTGTTCACCAATCAAATTTAATAGATTCGAAATAGAATGTTAGCCAAATCTATTTAGTTCAGTAAAGTCTGGTCAACAGTCTGTAATGTACATATTTGC

TATTATAAATTACTGACAAGTTTCATAATTTAAAAAATTACCGTTTATTGAAATTTGAAAATATCGGATAGATTTGTTGATTCGAGAAAATCAATGAATACAAGTTGAGTGATTTTTGGTGTATAAAATGATTCAAAACCACATGTTTATTTTTCGCACCAATAGTTTTAAAAATTTGACCAATCTTCGAAACACTTAATATTAATCAAT

GTAATATATTGATCACATTTTTAGCAAAATAAAGCGTTCGATACAAATATTTCAAAAACATATCAGAGGCAATTCCAAAAATATATTGGCCTCAGGGGGCGAAAGTCAAAAAAGTTTGGGAAACACTGGTATATCTCGTGTAGTACGATACTCTATACGTTGAAGTCAAAGGAAATTTATATTTCGAAAGATTCTGGACCGACCGAGAAT

TAAGCCCAGCCTCATCAATGCGCGCCTCTTTGTAGCCGCCAGACGTTACCTCTAGGCTAAGAAGCCCCAATCAGCACCGTACTACGTTTTCTGATTCACCAAGCACAGTATTCGCATTGGTGCATTTTCTCCAAGCACACATTTCGTACCAAATGCTGCGCTATACGCTACGTAGATTATTGGGAAGCTCATTACAGGTGTCGGACGATC

GGCGGTACCAGTGGCCGGTAATTGGCCGCTAATAAAACTACGCGCCTCCTCAGCCTCACCTCTCTTCCACCTCTTCATCTGTATATTATAAATACGTCTAAATAATGCAACGTCGAATGAATAGCGTCGAGCCAACAGTCAGTTTGCCGGTGGATATGATATTTTGTACCAATGCACAGTGGGCAAAAACGGACCCGGAATCGCAACTTT

GGTAGCCTTTGGCTTATCCATATGAAAGAAAAACTACTAAACCCACTGATCCTGGGTAACTCAGGCACAGCAATCCTTGGACCACTTTAGGAAAACTAGATATGTGTGCTGCACACTGACAAAAAAATAATTGAAATCCGTTATGTATTCGCTGAGCGGTGAGAGCTTTCCGATTTTTTATTTCCTCAAGCCTATTACGATTTAAACGAT

AAATCATGCAAAAAACAATTCCATTTCGATTTTCCTTTAATCATGAGAAAAGCCATACTTTAAGGTATTTGGGGCAACTCAAAGTCATTAAATCATTTCTTTTTAGTCACCATCATCAACGCTCTGGCAAAAAGGGGCAAAATGGGGTAAAGTCAGGTAGCTTCGAAATCGGTACAAGACGAAACCATTACCAGTTTTATTCCTTGTGTG

AGAGTTTTTATATACAAACAAGTCATGATAATAAATAGTATTTGATCCTTCGACCAAGTCGCATGCTCCCCGCAGTTGAGGGGCATAATCAAACATGTCGCCCGCTGTTCGTGTACTAGGATCATTAAGTGGTGGTTTGATCTATTGATCGCGTCGCTTGCTCTTACGGGGAAAGAGTGATGAACATTCGTTCGGCGTACAATGCGTTAT

CCGAATAAAATTGCAGATACGGTTAGCGCCAACAGATATTGGTGACTTCCAGATCTTTATCTTACCCTACTAACTAATATCCTTTCCATGACAACTGTGGAGATGCAGAGGTATACACGGTCTAAATGACAACGGTTGTTTAACTAACATTCCTTCGCTTCCCCCGATGACCGTAAGGACGAGGCCAGCGCCATTATTGACGCTTTTTAC

ATTCTAAGCTCTCGATTTGTGCACATTGAAAATGTTAGCTAATTTCAAAGGCCCATTAATTGAATCTCTGTGCAAATTCGATTGTTTTTTGTCAATCTGGAGTAGCAACTAAACATTGTACGGTCATCTTATATATGCTCATGACAAACCAGCAGGTACATGATAATGTTTCAAGTTGCGATTTAGGGTCCGTTTTCGCCCGCTGTTCAA

TGGTTTTGAACGTCATCATCATTATTATCACACAACAACGTTGCCACTACTGTGACCTTGCGCCAACAGCACCTGTATCGAGATTAACGTCCGACTCCGGCATGAGTTGTACGAAGAGATGTTCACAGTTAATTAAATTAGAATGACTGGACGTAGTAATTACATCATCTTATCAGCTTTCAGTAACTCAATTAATCTCCTTTCTGTTTC

TATTTGTTTTCTCTACTTAATAGGTCCGTATCAGCCGAGCAATGATATGCTGCCCCAACTCTACTCGCTTTTCAAACAAGCGACCAAGGAAGATGCTGCGGAGCAGGAGGAGAAAGCCGGCTTTCTGGGATGTTGTCAGCAGGTTGGTACCAAACATACCGTACTCCAGCGTCTCTAATAATTGTTTAATCGATTTTTCATCCATTGCGT

TTGTTTTCCCAGGAGGCAAATGGGACGCGTGGAATCGGTTGGAGCCGAAATGCCAAAGGAAGTGGCCTATGCAGAAGTATGTCGATGAGTTAGAAAAAGGTGCGTATTCTGGTTGTAGACTTCATCAAAAATACTAAAGCTTTGATTTATACTACATGTGCAATGGACTAATTTCTGATTATCTAGCAATATCCTGATATCTATCAGTAA

TATGAGAAGCATTAATCTTCCTTAGTCCCCTATCCTGAGTTTAAATGCGGACTCCGAATGGACCATGTTTTGCCTAAAATTTCAGGTGCCTCAGTTCTTTAATGAAAAAAATATATAAAATTCTAATAGCTCGCAAAATCAACTGTGATAAAAATATTTACTGATTTAGGTTCTTCAGGCTTAAATGAAAGGATTTCCAAGAGATCCAGA

TTTTATGAATGTGGCCATCGGTTTTCTTCTACATCTGGAGAAAAGTACCTTTTGGAGATCCCTACTTTGACGACTATAACGCAGCCAATCCAAGTAATATGAACCGTTCATATAGGGTCTACCACAAAGGCCGAATCACAAAAGGCCAGATCATAGGCCAGATCACAAAGGCCAGAAAGCACAAAGGCCAAGAAACATACAAATGGCCAG

ACTCACAGAAGACCGAATCACAGAGGGCCGAATCACAGAAGGCCAGATCACAGTAGGCCAGATCACAGAAGGCCGAATCACAGAAGGCCAGATCACAAAAGGCCGAATCACAAAGAGCCAGATCACAAAAGGCCGAATCACAAAAGGCAGATCACAGAAGGCCAGCGAATCACAAAAGGCCAGATCACGGAAGGCCGAAATTTTTAAAAT

AAATTTTAGTGAAAGCCACAAACATTCAGCACATTTTTTCTACAATATTATGGACATTTTTACAGTAGGGTTAGTGGGACAAGACGCACAAATGACATTACTAGCTGAATAGAGTGTATGGTTAGTGTTTTTGGGTGTGTCAACTGGTCTATTCTTAGTTGACTTTTTAAAATAAATTCTATAATGGAAGCTTAAGTAGTCGCGCTGTTG

TATTTTGTACAACATAAGTGTAAAAACCTCCTTGAAAACTGGAAGGGGTATACAGTGCCTATTCGATTTGGCAACATGTCAAAAATTTCATGTTGCCAAAGTTAGTTTCGTTCAAAAATTTAATAATTGTTTTTTATTAAAACTTGTATCCCTTCATAATAATGACCACAAATTAAATTTCTTCTTCTTCTTCTTGGCATTACGTCCTCA

CAGGGACAAAACTATAGCTATCAGCTTGTTCAATGAGCAATTCCACAGTTGTTACAGGAGGCTTTATTTGCCAAAGTTGCCATTTTCGCATTCGTATATCGTGTGGCAGGTACGATGATACTTTATCTTAGGGGAAGTCAAGAAATTTCCATTACGAAAAGATCTGGACCGACCGGGAATCGAACAGACACCTTCAGCATGGCTTGCTTT

GTAGCCTGGACTCTAACCACTCGGCCTAAGAGGTTCGCTTCAAATTAAATTTACCTATTTTGAATATGGTAAGCTAAGAGGCCCACAGAAACATTAAGTTGACATGATTTGTGACTGACCATGATCCATTTTGTGTACATAGATTTCCCGTGTTGCCAGAATTCAAACCGAGTCAAAAATCAAACGGTTAAAAGTATTCTTTGTAAACTT

AGTATTCTTATGATCACTTCAAACACTTGTTGCCCTATCTGGCAATCATTCTCCCTTTCACTTACATTTGCGTAAAGACTATGTTGGCATTATACCGGGCAAACGCTTGCTGTCAAACTCGCTGTCGCTCATCGGACCTAAAATGGAAAAACATCCCACTCTGTTTGCATTATACCGGGCAAACGCCTGGATTTGCTATTTGCATGACAC

CGAGGCCGTGCTATACACAACCATCGGATATCCAGAACTTCGGCGGCCTTCTGCCGCCTCAGTACTATCCTCAATGTGGCTTCGCCGCATAGCAAGTTTACTGTTTACTGACATGACTGCATTTAACAAAGCATGGACTTGTTCTTTTCATATCCAAGCTTTATTTTGTGATTTGGCTTTGTGATTCGGCCTTGGCCTTGATTCGGCCTT

TTGGCAATTCGGCCTTTTTGTGATTCGGCCTTTTGTGATTCGGCCTTTTGTGATTCGGCCTTTTGCTTGTTCCGGCCTTCTGTAATTCGGCCTTCTGGCCAATTCAGCCTTTTGTGATTCGGCCTTCTGTGCATTCGGCCTTTGTGGTAAACTAGAAAATTATTCGGCGCGCAATTCGGCCTTCTGTGATTCGGCCTTTTGTGATTCGGC

CTTCTGTGATGTCCCTTCATATATGATATAGGTATTTACGTGGAATAACAAATGGACGGAGATAGATATGTGTTTTTTAATTCACATCGGTATCATTCAGTTCATAGATATTTTGCATGATAAGCTGCTGTCAAATAGAGAACATTTTTGTCTGGCCATTCCTGAAAGTTAGGTCAATTACGGTCTATGCTGAACTGATTTCTAGGATCG

AAAAAATACCCAATTTGGAAAGATTATTCTAGTCCACGAAACGAACAACCCGGGCTGGTAGCAGGTCTCATTTCAGAGACACGGTCACTAATTTCTTGCATTTCTCTAAGCAGTCTCTATTTTGATTGGAAGGTATACTAACTTTTTTAAGCAAAATGGTCTCTAAAAGCAGGATCCGTCATTGATTTCAGAATTTTCAGAAGAAAAAGC

AGTTTTCTTCTTCAAAATCAAGAAAATTTACAGGAAGAATGTATTTCTTTCATATTCTATTCTACAACAGAAACACAGTGAACACATTTTCATCGAAGATAATTGTAGTTTTGAACAGAAAACTGCTTTTAATTTCGATGATTACGATGACGGAGTGTTGCACGTTAAAGGTGATTATAAAATGAAGCCAAACTTTGTGAATTTTCAAGT

GCACAAGACTAGAGAATCTGACAACGGTTCGCGTGAAAATCAATCAAATTGCTTGCCTGCTGATGGTGATCAATGGATAAAAATTTCAACGCGGGCGCTGTTTGATTCTCGATAACGTAACTAACGAGGTTGGCTTCGTTCTATAATCACCTTAAGCCACTAACATAACAAAACCAATTCTGGTGCAGCCTCTAGAAACTTCAAAGAAAC

TTTTAGAAACTCTCACACGATAATCCTTCAGGATTTTTTTCTCAGATTACTCTAGGAAAGCCCCAGGAGACTATTCAGAGATTTCGTCTGGAACTCTTCCAGGTATTCCTCAGCACATCTCAGGGTATCAATAGATTTTTTCTGGATTTGGTTCTGAAAATCTTTCAGGATTTTTAAGGACATGTTTAAAGAGATTTCTCCAGAACTATC

TGGGGAGTTTCCAAATAGCCGTGGCAGCAAACGCGCAGCCCTATTCAACAAGACTATGCTGGGGGTTTGGGATTTGAATACAATTGGTCGAGATTATTTTCAGATTGGAAATTTTTCTAGACTACCAGGGCATAGGATTTGATGTTAAGCCTGTAGCCTGGTCAAATCACATAAACGATTTTGTGTGGAGCGCCATATTATCAAATGCAT

TCGCCTGAAATCTGACAAATTCAGTAAATGAAAAAAATTTCTTTTGTGAGAATCCTGACCATGGCTGCCATGGAAGTCGTGTCCAATATTTCGTGATAAATAGTCTATTCTATATAATCTATCAAAAATCCTGCATAAATCACTCCCAGAAGTTGGAGTTGCAAAATTAGTGATTCAATTGAAATCATTAAAAAATTTAGCATTTTTGGC

AATTCACTTTTGTCCGCTCTTCCTCATTTTGGACTTCTCATCTCAATAATAGTCTATACTAGCGCAGGGCATATTAAAGTACTATTCGATTAGCACTTGCATGATAAAAATGTTTTGTAATGTAAATTAGATTTATGTGTAGAACAATTTTACCTGTTTTTTACAAACTCGATGTGATCTAATCTATTTTAAATTAAAAATAAAGATTTT

AAACAATTCATAAGTTAAGAAGACTCTAAAAACATTATCGAGCTTAAAAATATAAGCTTCAAACTAAGCTGAGCTTTGGTATCTTCCTAACTTTACGTTGGTAGAATAGTCAGTGGGCCAAATTTGAGAGAATCTTCCTTTCCTCTGCGGGCTGCACAAAATGAGGTCCTTGGGACGTACGTTGGGCAGCCCTGGCATAAAGTATCTTCG

TGTTTGCCACGCGATATACGAATGCGATACTTAGCGACAGGAGAACAGATGAAGGAAGAAGAAGCTCAGAAAATACATTGTTATTCTGAATAATTTCCCAAATACTGCTTCAGATGGATTCCCGAGATTTTCCTGAATTCTTCCACAAATTTTTCAGTAGTTACTTTATAGCAGATCTTCTAAGGTTTCCTACAGGATTTCTGTCAAGTA

AATCACTCAAATGATCATTGCAGAGTTCTTAAAGAAATTTGTTTGGAGATTTTCAAAATCTTATAGATTCCTCAGAAGATCCTACACCGATTTTTTCAGTTATTATTCCATACATTTTAATATGGATTTCTTCGTGAATGCATCTAGAAATTGCTCAAAAACTTCTCCATAGATAATTTCCTTAGCTCCTAATAACATAAATCAAAGGAT

CCGTCACTCCAGAAATATTTCGAGAAATTTCTTCAACATTTCATCCCGTATTACTTCTGATTGATTTATAAAAAAAACAGTTTTGTAGGGTGACGAATATTAGTACACCTACCCTACTTGTCTTGGTTTTTGCAAAATCTTCTTCTTTTTTCTTCTCATGGCATTTCGTCCTCACTACAGGACAAAACCTGCTTCTCAGCTGGTAGTCCC

ATAATTAACCTCAGGGTCAATCTCTATAAGAAAATATGATAACAAAAATGGTGATGCTGTACATGACTAAATTATCTATTTTACGTTGCGTGATAATATGATATGAAGCATCTTGTAAATATAAATATCTCATGCTATATAAGATAAAATTTATTCTGTAACAAGAATTTTACCAAATGCTGTAACTTCTTCAAATCAATGATTATTTGG

AAACAAAAATCAGCACAAATTGACAGAAAGCTCTCCAGGAACCTTTAAGAATTATATCAGAACACTGAACTGGTGCTGGCTTGGATCTACTGCAAATGTAACCTAAAAGAGAGTGACAACCAGAAAATTAACGTTGAAGATGATTGCTGTTATGATTGATAAAACTTAATTCTTTATTTCGAATCATTCGGTTTTCAGGAATGAAATATA

ATTCAATGCACTGTATACTACCACATTCAAGCTTCTCTCTCTGTTAATAAAAGTGTAGAGATTATTCGATCTTGAGCCGTCCTGTAAGTAGAGTCAAACAATTCAAAATGTTTTATTAATTTCTATTTGAAAGTGTCTTTACAGTGAAGGTATATAATTAAATATTAAATTAAGTTATTACAATTCCCTAACCTTGTTCTGTTTCATGTT

ATATTGCTGTGTACGTTTGACGCAAATATCAAATACTTGGAAACACCAAATGCGGTAAGATTTTCAACAAGAAAGGCGAAGTCAAAGATTGGTTTTCTTTGCTAGGTTGACAGTGATATGGCTTTCACATGGTTTATAAGTATCTTTGATTACTTTTATTGTTACCGCATTTAAAGCTCTACTAGACTGGATTTGCGTCAAACGCAAACA

GCAATATTTAACGTGGTTGAAATCTAAATATCGAATAAAGAACACCGAACGCAACAATAATTTTTTAACAAAGAGAGCGAAGTCAACGGTTGATTTTCTTTGCTAGGTTGGCGGTGATATAGCTCATGGTTGCTAAGTATCCTTTTGATTTTGTGTTGCCGCATTTGAAGCCAATTAGAGTTAGATTGCGTCAAACGCGTACAGCAATAT

AATACATTAAATATGAATTATGATTCTTTATTTCTAAAGCTGATTTTCCGACAGTTGCATGCAGTTTGAAACTAAATAATTTTTAATCGACAAAAATCTCTGCATTAACCTTTACGTTACCGACTCCGACAAGGCATTGTCAAATAGCTGCCATTTCGTCAATTTCCATTCGATTTTTTGAAACTACCCTCAGTTTACTTTAAAGGTTTC

GTTATTTGTCATAAATAGAGATTGCTAAATTCGGAATAATGCCGGAGATATTCCGGAGTTGTACTGGGGTCACGATGGTGCCAAATTTGGGAAACGTGATTATTTTTTATTTATTTCAGCTACAACGCTTAAGTTTTTATGTAAAACGTGATACTACACTTTTTATATTAAATTTAGGGAGAGGATTATTAAAGTTGTGTTTGATCCTTC

GACCCCTTGACGCTTGCTCGGGGGCCGGCGTGAGGCGGGATCAAACATGTCGCACGTCGGTCGGTGAAAGTGAAAAAAGTGGCGTGATCGTTTAGTTGTGTTTGATCCTTCGACCTCGACGCTTGCTTCTGCGGGAGCCGGCGGGGCAGGATCAAACATGTCGCGCGTCGGTCGAGTGAAAGTGAAAAGTGGCATGATCGTTGGATTTGT

CGTGTTTGATCCTTCGACCTTGACCTTGCTCCTGGGGGCCGGCGATGAGGGGAGGGCAGGATCAAACATATCGCGCGATCGGTCGGGTAAAGTGAAAAAGTGGCGCGTTCGATTAGTTCTTTTGATCGCTTTTCCGAGATTATTGCTGCTCTGGAGCAGTGCAGCAAGAAAGCCCGAGCAGTCGTCAGTTGTTTTCCCTCGGGTCGTGCG

CCTATTTTCTCTGAGTGCTTTTATATAGCCGATCAAATGAGTGAAGCTGAAAGTGGATTATTTGCTGCTCAAGGATGACGGATCAATTGAGCTCGTTTCGTACTACTATTTCTAATCTATAGAAATATGTATGACTGCACCTTTAATCATTGCAACGTAAATATGATTTTTCAACAAAACTATGAAAGGTTCGTCACTGTGAGTGTCGAC

ATAAACTCTGATTCATAAAATTATTATGCTCTATTTTTTTTTTTTTCAAAACACCATTTTACCTTTATTTTGTAACTAAAAAAAAAAAACTTTAGATGATTCGTACTACAGTGTAGACTTGCGAAGGTTCACTTTATTCAACAAACTTTGAAAGGTTCAAATGCGTCAAGAGTGTCGGCATAATTTTCTCTACATAGATATTGTTTCATA

TCAAATGGAAAATAATATTTACATATGACATGTTATATCTCACCAATAATTATTTATCATGGTCTAATCGCTTATCAAAATTAGGATCTGTGACCTTGATCTCCCATTGGCAAACATCCTCCCAATAACTTCTTTATTGGAGATGCAGAGGCAAACACGGTCTCCAAATAACCAAAAGGTTACACTAACCTTCCTCCAATCCCACCTGAC

TGCAGGACGTGGCCGGCGCCGTTATTGACCATGTATAAATGAGGCACTGGATCATGCACACTGGAAGAAGATTATAGTAATCAGCCGATCTTCTAGTTGATTCTTTGTGCATTTTCACTTACTCAGTCAATCACGGAATGCTCGATGATATATTAATAGTCAGTCTAGCTAGCTAAGCTAAGCTAAGCTAAATTTAGGGAGGGATTATTA

GAGTATATGTTTAAAACTTCAAATTAAACTTTAACACTTCTTTGTGGGAATAATGGTCGGTGTTATCATTTCAACATAAATTTGGAATAAGTAGACCGTTTCAGACATCGTAGACAGTTCCACTTGAGGCATGATTTGAGGACATTTCGTTTTTTTTGTGAGCTTACCGTACGACGTATCAATCTTCACAAATTCAGAGAACTTAATAAC

TCAATAAAGAAGAAAACCCCCGGTAAAACAGATATGACCACTCAGATCTCAAGCTGGGTTCACTTAAGAGCTTCTTGAGGAACATCTCCATTTTATGTCCAAAAACATACCGTGCGACATCTCAATCTTAATGAATTTTAAATAACTTGTCAATAAATGAAAAATAACACACGATAAAAGATTTGGCCACTCCACAGCTTCTAGCTGGAG

TTCCATAGGGCTTCTTTGGGGATATTTCTGCTTTGTCTCCGAAATCTTTGCCCAAGGTGGCAAAAAAAACATGTCAATAAAGAAGAGAATCAACAGTACAACATGTGATGTGGCCACTCCATAGCTCAAGCACAGTTTCCACCTCGCTGGAGATATTTCTGTTTATATCCATAACAAACCGTGTGACGTGTCAATTTTCGTGAGTTCAGA

AAGAACATGTCAATTCATGAATTATGAACTAGGTATCAATTATTTAGGTCTCTCCCTGAAAGCACCTGGTACTGGTTACATAGGCCTCTTGAGGACATTTTTATGTTCAAAACATATTGTGCAACGTTTCAATCTACATGAATTCATTCGCGAAACATGTAGGGCTAATGAGAATAGACCCTGTGTTTTCCCTCCTGAAGCATCACGTAC

AGGTTCAAGGCTATTTTTATGACATTACCGTTATTTGAACAAAACATGATGTGTGCCACGGTAAAGCATAATGAAAATTCTGAAAGTATTCAAATAAATACCACCATAAAACTGAGTAAATGGCTCCCTCTTCACTGAGTACTTTTAAATGCTTGAGGCAGGATCTTTCACCAAGACTCAAGATGGTTCCACGATTTTGGAAAATCTGTT

ATGCAGATCTTCATAGATACACCAGGAATGCACTGCAGCGGATTTAAAACTCAGCAGAAGCTGAAAGAAAAGCATAACAAAGGAATCCACCAGAACTTTGTTTCGAACATCATCATAAATCCAACAAAAACCTAAAACGTTTTGGCTAGAATGTTAATCTATTGAAAACTTCGAAGAAAAATATTCTATTAATTACTCAGTTTTCTCAAA

TAAATTTGTTGATAAAAACTTGAAAGGAATTCGCTTTGGAAACTCCTCATGAATACATATTTACTGCCAATAATTGTTTTAACAGTATAAAAAGAACAACTTATTGCATGAGCATGAGCATGAGCATGAGCATAAATGACCGTACAATTCGTAGTTGCTACTCCGTGATTGACCAGAACAATCGAAGTTGCACAGGGATTTAGTGGATGG

GGGCTTGGGATTAGCGCCCCCACTGACTAAATGTGTGCACAAAATCAAAGAGCTCAAAACTTAAGGAGTCAATAACGGCATAAACCACGTCATGCAGTCATCCCAGGAAGGAAGGAATGTTGATTGGGCAACCCATTGGTTTGCAGAGACCGAAAAGAATGTGCCTGCATCTCCATAGTTGTCTCTGTGGAAAGGATATTGGGTTAGTGG

GATGGGTAAGGATCTGGGAGTCATCTTTAGTTGATGATGCGATCTTGATATATTCACGCCTGACCGGATTCGCGATATTATTTGATAAATGAATTGTACGAACAAATGTCCATCGTTCGCCCCCGCAAGAGCAAGCAACGCGACCAAGGATCAAACACTACTAGCACTAATCACGCGATCCGCGGCTGTTATATACTATCACGTGCCGCG

CAGTGCTGCTGATGATCCTGCCCGCATCACTCGCCCCCCACAGAGCAAGCAACGCGACCAAAGGATCAAACACTACTAACACTAATCACGCGATCCGCGACTCTTATACTATCGCGCCGCGCGAGCGTTGCCTCTTCGCATTGATCCTGTCCACATCACCCGCCCCGCAGGCAAAACAACGCGACCAAAGGGATCAAGCACTACTAACGC

TAATCACGTCGACTCTTATATACTATCATTGCCCAGCGCGAGCATAACACGATGATCTAATCCACATCACACCCGCCCCGCAGGCAAGCAACGCGACCAAGGATCAAACACTACTAACACTAATCCACGCGATCCACGACTCTTATATACTACCGTGCCGCGCGGAAACGCGCAACTCGATTGATCCTGTCCACATCACTCGCCCCCCGC

AGGGCAAGCAACGCGACCAAAGGATCAAACACTACTAGCACTAAATCCGCGCGATCACGGCTGTTATATATATCGTGCCGCGCGAGCATAACACGATTGATCCTGCCCGCATCCCTCCGCCCCCAAGAGCAAGCAACGCGACCAAAGGATCAACACTACTAACACTAATCACGCGATCCGCGACTCTTATACTATCGCGCCGCGCGAGCA

TGCCTCTTTGATGATCCTGTCCACATCACCCGCCCCCGCAGAAGCAAGCAACGCGACCAAAGGATCAAACACTACTAACACTAATCACGCGATCCACGACTCTTATATATACTATCATTGCCGCGCGAGCACAACACGATTGATCCTGTCCACATCACCGCCCCCCGCAGGCGACAAGCAACGCGACCAAAGGATAAACACTACTGCCGC

CAATCCGCGATCCGACTCTTATATACTACCGTGCCGCGCGAACGCTGCTCGATTGATCCTGTCACATCACTCGCCGCAGGCAAACAACGACCAAAGGATCAAACACTACTAACACTAATCACGCGATCACGGCACGGTATATGCTATCGTGCCGCGCAGGAAACATGCGCTGATTGATCCTGCCGCATCACTCGCCCCCCCAGAGCAAGC

AAGCGCGACCAAGGATCGAACACTACTAGCACTAATCACGCGATCCGCGACTCTTATATACTATCGCCGCGCGAGCAACACGATGATCCTGTCCACATCACGCCCCGCAGAAGCAAGCAACGCGACCAAGAGATCAAACACTACTAAGCACTAATCACGCGATCCGCGGCTGTTATATACTACCATTGCCGCTTTTGAGCATACTTTAAC

ACGATTGATCCCTGTCCACATCACCCCGCCCCACAGAGCAAGCAACGCGACCAAAGGATCAGAAACACTACTAACCCAAAAAAAAAAAAAAAAAAAAAAAAAAAAAAAAAAAAAAAAAAAAAAGAAAACTTATTGCCCGTTTTGAAAAATAAATTAGACCAATCGTACCGCTTATTCATGAATGGAGGAATTCCACGTGCAACAGAATAC

AAATTTTGCCTAGAATATGTTAAGTTTGATCAGAAATTAAACAATAATCTAGCTAGCAATTATTTAATTTGGAACCCATTCGAATTTCATCGATTACCCGCAAAGGTTTTCCTATAGATTAGTGTTCGGTCCATCGATTGTGTCGTTTGGGAGAGAGCGATGCAATAGAAATCAGGCGTATCCGGTTGCGCAGTGCGAAGAAAAAAGTGG

TGTAATCAGTCATAAATGTTTAGTCATCGATTTTGTGCTTGCTCCAGCTGGAGCGAGCGATGAACTGGAAACACCCACTAACCTATGTACTTTCCAAGACAATTGTGGAGATGCAAGTATATTCAGTCTTCGAACTCTCGATTCGTGCTCTTTGTGAGAATGGTTCGCTAATCCCAAGTACCTTTCATTAAATCTCTGTGCATCTTCGAC

TGCTCGGTCAATCACGGAGTGCAATTGCAAGCAGTCATCTACACTCTTAAAAATAATGAAAATTACGAGTGACGTAAACCTATTTGCAATTGAATATTTTGCAACTTAAACTAAAATATGACTTAAAATTTTGTTTCAATTGACGTAGTTTAACATTTCAATGTATAATAACCTACTTTTTACACCATGTTGAAAAGTAAACCACAACCT

TTGAATCTAGGTATTTCAAACCGTAAAATTTACATGAAGCTTATTTTCTGTTATTTAACACGTAATTTACATGATAATCAACGTCTGACGATAGCATGCAAACCAGCAAGATCTGTTTTTCCAACATAGCCGGCACAGAACGTCATGAGAGTTACTTCGTTCGCAATCGATAAATTGGTGATCAGTTTCATTTGCTGGGGAATTGGAAAA

AAAATGATTTCTGCAATTTGTGACGATGGAGTATGATGCATGATTATAAGCAGTGCGGTAGTGATCGCCTTGATGGGGGTGCTCGAATGCTAACAAAAACACAGTGCCTTTGTGAAATTATTTTAGCTCGGATTGCCACATTTTCAATGCTTTGGGTAGTTAAGTACTTTGAAAAATCTACCTAGTGATTGTCGCGACATAAAACACATG

ATTAAAGTGAAGCTATCTGTATTAAGAATGATTATATTGGACGGATTTCAAATGGATTGTGTGTTTCCCTATATGTAGCCAACTCTGTGAAAATGAAAATTTGTGTAGTGTTCTCAATTTCTATTCGATGGTCGAAGACATTGAGTGCGGTTCTGAAAAATCTCGCTGCTGGATAGTAAATATACGGTACAATATATTTACCCAACCTTA

TAGAGTCAAATGATGTGATTAGACGGTAAGTATTTCTATATTCTACGCTTATTCGCCTCAAGTACAAAATTGCAAAAGAAAACACACAGCATAAGTAGAAATCTACACTCTTCGAAAATTTCATGTAACTTTAAGTCTTTATAGATGCACATAAAAAGCGAAGCTTGACTCAAATTTACATCCTTTCTGAAAATTACATGAGGAACAACA

ACCCAAAAAAAAAAACACTTACACACGACATGACGTAAAATATATGAGGACTTACTTTTACGTTATGATGTAAACAAATCCTATAGATTTCACTTCATGCCGTGTGTTTTCTTTTGCAATTTTGTACTTTTCGGGGCGAATAAGCGCAGAATATAGAAACACTTACCCGTCTAATCACATCATTTGACTCTATAAGGTTGGGTAAATACC

TATATTGTACTGCATTTACTATCAGCAATGAGATTCTTTTTCAGAACCACTAATGTCTTCGACCATCGAATAGAAATTGAGAACACCACACAAATTTTCATTTCACAGAGTTGGCCACATATAGGAACACACAATCCATTTGAAATCCGTCCAATATAATCATTCTTAATACCAGATAACTTCACTTTAATCGTATGTTTTATGTCGCGA

CAATCACTAGGGGTAGATTTTCAGCAATTAACTACCCGTGTCATTGAAAATGTGGCACCTAAACTAGGAAATTTACCAAAGGCACTGTGTTTTTTGTTAGCATTAGACACCCCCATCAAGGCGATCACTACACCCTTATAATCATGTATCATACTCCATCGTCACAAATTGCAGAAATCATTTTATTTTCAATTCCCAGCAAATGAAAAC

TGATCACCAATTTATCGATTACCCGAACGAAGTAACTCGCGACGTCTTGTGCCGGCCATGTTTGAAACAGACTTGCTGGTTTGCATGCTATCGTCAGACAAGTTGATTATCATGTAAGTTACGTGTTAAATAACAGAAAAAATAAGCTTCATGTAAATTTTACGGCTTGAAATACCTAGATTCAAAGCGTTGGTTTACTTTTCAACATGG

TGTAAAAGAAGTAGGTTATTATACATTGAAATGTTAAACTACGTCAATTGAAACAAATTTTAAGTCATATTTTTAGTTTAGTTGCAAAATATTCAATTGCAAATAGGTTTACGTCACTCGTAATTTTTCATTATTTTAAGAGTGTATAGGGATTTGCTTTTACATCATAACGTAAAAGTAAGTCCTCATACTATATTTTACGTCATGTCG

TGTGTGTTTTTGGGTTGTTGGTCTCATGTAATTTTCAGAAGGATGTAAATTTGAGTCAACTATAACTTCTTTTATGTGCATCTAAAGACTTAAGTTACATGTAAATTTTCGAAAGAGTGTATGCTTGAGCTCATGCTCATTGTCCTGAGTTCTTACAAATTCTGCATGTATTTTTTTATTTTTTTTTTGAAGCTTGAAATTTAATACAGC

CAGTTCATTCGCCCCAAAGAGGCCCTCAGGAACCTGTGCCAGGGAGATCTGGAGTGGCCACATCTGTTAATACCAGGGTTTTCTTCTTTTATTGACATAATTTCTCCCAGAATTCCATGAAAATTGAGACGTCACACAGTATGTTTTGGACATGAAACGGAAATGTTCCCCAAACTGGCCCTAGGCGGAACCGGCTACCATGTTCGGAAC

AGTTCAATTATTCAGTCATGTTGAAATGATGACAATAGACCATTATGTTACATTTTTACATAAAAACTTCATTGTTGTAACTGCAATAAATATAAAAATAATCACATTTCCATATTTGGCACCTATGACCCCAGTACAATCACCAATATCTCGGCTTTATTCGAATTTAGCATTCTCCATTTATGACAAATAACTGAAATCTATTTAAAA

ATGTGAAAGCTGGCGGGTCGTTTCAAAAATCGAATTGGAAATTGACGAAATGGCAGCTATTTGACAATGCAACGTCGAGGCCGGTAACGTAAAGTTAACACTGACACTAGGTACTATATTATACTTTTTTTGACTTTACTACTGTAATCAAGCCACACCTAAACATACCACACTTTGGTTGTATAGCTCGAAAGAAATAGATCACTAGTT

TATAAATTTGGTGAGGCTCTCGTCGATCACACCACACGTGAAGTTTAAAAAACGTCGTCCAAAAATGTAAGAAACAGCCGATTATCACTCTTTCTTAGAAGTCCCTTCCCCAATAATCAGTGTGATTTTTCAAGGACAAGTAAATAGTCTTAGCTCTATTATGTGAATGCGATCTACTGAACACAAAAGTTTCCTTTGCTGAATGGCTAT

GTAAACAGTGTATAATCTCATATGGCTACAATTGGCGTATGGCCAAAACCGGTGCTGTACCCTTCATGCTATTTATACTAAACGTACTTTAAGCCAAATGTCTATTATGGCTAATGTATTAACGCCTAAAGTCCGTTATACACGGAGCCGCAAATCAACCCAACTTTGACTTCTTTTGACGCAATTCGGCTCTTCCGTGGGATAACCCAA

ATTTGAGTTAGCAATACCGTAACTTCA

>173ccd42-fdc8-4e1c-b262-0d6c006819e6

CTCGGTATGCCTTCGTTCAGTTGCATGTGCTATAGAAAAAACCGAATTTAGTACTATACCATTTAATTCCACTAGGTTGTATCCTTTGACAGATACGCGTATTTCGACCTCAACTGTAGGCCGTCTTCAGTGTCGTGTACTCGTACAGTCTAGTACACGACACTGAAGACGGCCTTACAGTTGAGGTCGAAATACGCGTATCTGTCAAGG

ATGCAAACTCTGGTGGAATCATCAGTATAATCTACTAAATTCGATTTCATCTACTGAATATAGCTGAAGTATCCAATCTACGAAGTATACCGTAACTACGCTAACGCTAACGCTAAACGCTAACGCTAACGCTAAGTAAAAAGAAGCAAACCTGTGAAAAAATACTGCCGTTCTACACATATTTGTCCCGCGGTCCATAAGGTTTACATA

GAACATGGGATTAAGTAGGCGTGCCGAGGGCAATATATCAATTACCTATCCAAGGTATGTCGAAATGATAAGTTAGATATTTAAAAAAGACACGGACACCGTCTTCAGCCATTTAGCTGCACAGACTGTAACAACACTAGACAACGTACGCATGCTCTGGTGGCACAGCCGAGAAGCATTCCTGACAGAGTTTCGAAAGAAGCGGGAATC

GAACCCACACCCCATGTATTTTGATGCGCTTAAATGCCTGACGACACTAACCGCACGGCCACGGGCCCGGTAAACATTAAAGATTGCAATCTTTTTGTATTGCTCAACTGGCTGTACAGAATGCTTTACAGAAAAATTTAAATGTCTTATTTAATAAGTAAACGTTCAGTTAAGTATCAACGATTTTCGTTTGTTTTATATTAAGCACAA

CTCGTTTGGATGTTTTCACCTGACTTTAAGTGTTATTTCCTGACTTTCCAAGTCAAATGGGATTCCTGACGTTTCCAGTTTTTCTTCTTTTTCGGTTAGTCGACATTCTAAAACAGCCTTGAGAAATGCAAATTTGTTTTCAAGCTAAAACTTCACATATGAGTGGTTGTTCAAGATTGGAGAAACTCTTTACCCTAACAGGGGGTGACC

TTTAAGCTACCCTTTTGCACTGGTGCCTAACTCATTAAGATCTCGTCAAACTAGTGGTTCCGCTATTACCTTTTCAAAAGATTTCGACAGCCTAAGCAAGGGACAAGTTTTGCGACCAACTGCCTACCTCAAAGAGACAACATTTCAATGTTGACTGACTAGCTACTCTTGTGATCGCAGTTCTTAACCCTAGAAAGATAATCATATTGT

GACGTACGTTAAAGATAATCATGCGTAAAATTGACATGTGTTTTATCAGTCTGTATATCGAGGTTTATTTATTAATTTGAATAGATATTAAGTTTTATTATATTTACACTTACATACTAATAATAAATTCAACAAACAATTTATTTATGTTTATTTATTTATTAAAAAAAACAAAAACTCAAAATTTCTTCTATAAAGTAACAAAACTTT

TAAACATTCTCTCTTTTACAAAATAAACTTATTTTGTACTTTAAAAACAGTCATGTTGTATTATAAAATAAGTAATTAGCTTAACTTATACATAATAGAAACAAATTATACTTATTAGTCAGTCGAAACAACTTTGGCACATATCAATAATATGCTCTCGACAAATAACTTTTTTTGCATTTTTGCACGATGCATTTGCCTTTCGCCTTA

TTTTAGAGGGGCAGTAAGTACAGTAAGTACGTTTTTTCATTACTGGCTCTTCAGTACTGTCATCTGATGTACTAGGCACTTCATTTGGCAAATGTTAGAGATATTATCGCGCAAATATCTCTTCAAAGTAGAACTTCAAACGCTTACACATAAACGATGACGTCAGGCTCATGTAAAGGTTTCTCATAAATTTTTGCGACTTTGAACCTT

TTCTCCTTGCTACTGACATTATGGCTGTATATAATAAAAGAATTTAGGCAATGTTTATCATTCCGTACAATAATGCCGCCGATACCTATTCGTCTTCCTACTGCAGGTCATCACAGAACACATTTGGTCTAGCGTGTCCACTCCGCCTTTAGTTTGATTATAATACATAACCATTTGCGGTTTACCAGTACTCGTTGATAAACATCCTCA

TCACAAGATGATAATAAGTATACCATCTTAGCTGGCTTCGGTTTATATGAGACGAGTAAGGGTCGTCAAACAAGCATGAAATGTTCCCCACTGGCCTGGAGCGACTGTTTTTCAGTACTTCCGGTATCTCGCGTTTGTTTACGATCTTTTGGTTCCACAATGGTTAATATCTTTACATGTAACTTGTGCATTGAATCCAATTATAATTTG

CCTTGGCACCAGCTGAACCAGACAAAGAAAGCTTCCCAGAAGTATATCGATTTAGAAGGGTTGACGTCACTTGCTGACTGCACTAATACAGCAAATGATGCAATTAGAATGATTCAAGTGAAATTCCCCCAAATTACTGATTTTCTCTGGTTTGGTTATCAAGATTACATTCGAAGCTAAGATTAGCTACCGAAATTATCATCGATCAAA

TCAAATCTTTCTCTATCGAAAAAGGCATTCGCACATCTTCCTCTCTATACCATATACACGAAGGGTAGGTACATTGACGTCTTTGCCAGAAGTTGAACTGCATCGTTCAAGGTACAGAATGAACGACTAACAGACACAAGCACGTTTTGCTGTCCATTCAGACACAGGGATGGTACCCATATTCGATCGATATAGAGCCATCCAACCGAA

CAGAGTATATGTATGAATGTATTGCTGAAATTTTCTAGAAGTACAACCACCACTGCGACAGTGTCTATAAAACGCCTGCAAAGGCAGAAACCAGCTCAATCGAATACGTTTCCTAGTGGAGTAGACATTACGCGGCCCAAGTAAGCAGTGCCAGTGCAAGTGAAGTGAAGTCTCTAGTGAAAAAGAGTAATCCAGTGTAGAGGAAGAAAA

TTTAGAGTGAACAAGCTTTATTCAAAGGACAATTACTATTAAATTGGTGAAAGTGCATTTCGGTGAAGGGGAATCTTCTAGTGAAATTAGGTAAATTAATTAGTGAAATTATAGCTATGAGCGAAAACTAGTTTGGTGAATGATTCCTTTGTCTTTGAATAGGCAAACTATTTTCCAAGATGGCGACTATTGAGCTTTAGTGATTAGTGA

AAATTTGCAGCACCAGTTTCATCATCATTAGTAAAACCCAATTGTGATTCACGGCGATAATCATATTTCGTTGAATCATCGCTACTAATTGAATTAAATTTCTAGAATAATAGAATAACGTATTTGCTCCGTCATATCTAAAATAAATATTTTGATAGTAATTACCCATTAAGGTAATATTAACATATCGAGAAAAACGAAATCGTAGAA

AACTTGAAGATACGCAATTTCAAACTACGTAGTTCAAGTCGAAAACAAGTTAATTTTTCGCAAAAGTAGGGCGTTGTTGTGACGTCATCACCTTCAAGTGTATATTTTCACTTGGCCTGCGACTGCAAACGCAGACAAAGCAAAACAAGAGTTTAAAACCTGTCGTGTCGTGCTCGAAGCCAAAGGCAATGAATCAATATCAAATGAGTT

TGCATTTCACAACCAATTACTCAAGCGTTTCCTCGTTTCTTTTCTGCTCAACAGAGATTTCCAAAAAAGCTGGTGAGCAAGGGCGAGGCTGTTCACCGGGGTGGGTGCCCATCCTGGTCGAGCTGGACGGCGACGTAAGCAGCCACAAGTTCCGGCGTGTCCGGCGAGGGCGAGGGCGATGCCACCTACGGCAAGCTGACCCTGAGTTCA

TCACCACCGGCAAGCTGCCCGTGCCCTGGCCCACCCTCGTGACCACCTGACCTACGGCGTGCAGTGCTTCAGCCGCTACCCCGACCATGAAACCTTTTGACTTCTTCAAGTCCGCCATGCCAGAAGGCTACGTCCAGGAGCGCACCATCTTCTTCAAGGACGGCAGCAGCTACAAGACCCGCGCCGAGGTGAAGTTCGAGGGCGACACCT

GGTGAACCGCATCAGGCTGAAGGGCATCGACTTCAAGGAGGACAGCAACATCCTGGGGCACAAGCTGGAGTACAACTACAACAACCACAACGTCTATATCATGGCCGACAAGCAGAAGAACGGCATCAAGGTGAACTTCAAGATCGCCGCAACATCGAGGACGGCAGCGTGCAGCTCGCCGACCACTACCAGCAGAACACCCCATCGGCG

ACGGCCCCGTGCTGCTGCCCGACAACCACTACCTGGGCACCCAGTCGCCCCTGAGCAAAGGCCCCAACGAGAAGCGCGATCACATGGTCCTGCTGGGTTCGTGACCGCCGCCGGGATCCTCTCGGCATGGACCCGAACTGTACAAGTAAAGCGGCCGCTAGATCATAATCAGCCATACCACATTTGTAGAGTTTTGCTTTAAAAAACCTC

CACACCTCCCCTGAACCTGAAACATAAAATGAATGCAATTATTTATTTAACTTGTTTGTGCAGCTTATAATGGTTACAAATAAAGCAATAGCATCACAAATTTCACAAATAAAGCATTTTTCACTGCATTCTAGTTGTGGTTTGTCCAGAAAGCTCATCAATGTATCTTAGTCGACGATGAGTCACGGTCTCGAAGCCGCGGTGCGGGTG

CCAGGGCGTGCCAGCCCAGGCTCCCGGGCGCGTACTCCACCTCACCCATCTGGTCCATCATGATGAACAGGTCGAGGTGGCGGTAGTGATCCCGGCGGGCAGCGGCGCACCGGAAGCCCTCGCCCTCGAAACCGCTGGGCGCGGTGGTGCAGTGAGCACGGGACGTGCGACGGCGTCAGCAGTGCGGATACGCGGGGCAGCGTCAGCGGG

TTCTCGACAGTCACGGCAGGCATGTCGACAAAGATCTGACAATGACAGTGCAGAGACTCGGCTACGCCTCATTGGACTTTGAAGTTGACCAACAATGTTTATTCTTACTAATAGTCCTCTGTAGCAAGGGTCAAGATTCTGTTGAAGCCAATGAAGAACCTGGTTGTTCAATAACATTTTGTTCGTCTAATGTTTCACTACCAGCCTGAC

GTTGGCTGCACTTCATGTGCCTCATCTATAAACGCTTCTTCTGTATCGCTCTGGACGTCATCTTCACTTACGTGATCTGATATTTCACTGTCAAGAATCCTCACCCAACAAGCTCGTCATCCTTTGCAGAAGAGCAGAGAGGATATGCTCATCGTCTAAAAGAACTACCCATTTTATTATATATTGATCACGATATCTATAACAAGAGAA

AATATATATATATAATAAGTTATCACATCAAAAGTAGAACATGAAATAACAATATAATTATCGTATGAGTTAAATCTTAAAAGTCACGTAAAAGATAATCATGCGTCATTTTTGACTCACGCAGTCGTTATAGTTCAAAATCAGTGACACTTACCGCATTGACAAGCACGCCTCACGGGAGCTCAAGCGGCGACTGAGATGTCCTAAATG

CACAGCGACGGATTCGCGCTATTTAGAAAGAGAGAGAGCAATATTTCAAGAATGCATGCGTCAATTTTACGCAGACTATCTTTACAAGGGTTAAGCTGCATAACGCCGACCGGTTTTGTGGATTGCATTCTTGCAACTCTATCAGTCGGATTAACCATATATAGGTACCTGCCTACTTCACGGATACCTATAGCACGAAAGTTCAAGACG

GCTGGCAACAACGACCGAACAACGAAGCACTCTTTGAGACACTATGATAGTTTGTTACAGAGGGTTCATTCTCCAAACCAGTTCATTCGTCGGTTCCTTATGGATTTGATCGTTTGGGTTTAAACCAGTTTGAATTATGCCGCTAACTATCGAGATTGTGGTAGACTAGGCAGATGATAATTTGCATAGATATTTTGGGGCTTACCAATC

AGCTTGCGGGTGCCTATAAATCCATTTTATGGCGATAGATGCGGATGCCTTTCAATGTGTACTCCAGCAGAAATAGCTAATCTTTGCGTCCACCGAAATCAAATGCACAAAACAAGCTTATCACATATTTGCGCTGGTGGCTGGGTAATAAATTACAAATGGCTTACCGTTCTTCGGTAATCCTTGACAATGTTCACCACAACATTGAAG

CGATCCTTTTGGACATGGTTCGGTTCTGTTCTTCTTCGAGAGGATGTCGTTTTACTGATATTGTTTATCATTCGTTAAGCATACCAACCAAAAAGAATGATAAGCTACTGTCTAGCCGTTTCTTAACAATGATGTGAAAAAATCAGTGGCCCGCGAAGGGGATAGGATAGGTAGAACAAGTCATACGCACACAGCTACCTGGGTAGGCCA

GCAAAACGATTGTCCTACCCATGATACTACAAAAACATAAAAACAATCTCTTTGCCTTTTAAGATTAAGGTCGTATTTCTCAATGAACATACGACAAAAGTCTGGAAAACCACTTTAAATTCTTGCAATAATACAACGGAACATGCGTAATTTAAAAAAATAAGGAACGAACAGCATTCCAAAGGTTCTCTAATTGATTTCATGTAAAAA

ATCCTAAGGAATTTCTACAGCCAGTTAGATAGATGTTCTGGAAAATAATTGAATCTAGGCATTCTGGTGGTATTTCTTGATAAATAACTGCATTATTCAAGGAAAAACTATTGGGAGATTCGTAAAGAATCATTGGAAGAATTAATAGATTCACCCAAGACTGCAGTCTTGTAACATATATGTTACACACTGATGGCTCCCTTCTTGATA

GAGCTCGTGCAGGAGTAACTATTCTCGTGAGCTAAAGGCTGAATCAGTTTTACTCCTTGGTAGAAACACACCGTTTTTCAGGCTAAACATTTGCTCTTATGTGGAATGCAATCAGCACTTCAACAGCACATGTGGTAAAGTCATATACTTCTGTTCAGATAGTCCAGGCTGCTATAAAAGCTCTTGCTTCCCAGCCAACTCAAGGTCAAG

CTTGTTGTCTTTATGTCGAACTCAAATTGAGGAACTAAATTCAGTCACTCAGGACATTCTGTTGTGTAACCGCTGCGGTGATAAGCTATAGGCTCTCTTTACGCTCATGCGTTTTCGTTATTTGAACCCATTGGGGTATGGAGCTAAATACTCTTTTTCCTTATGCGATTATCCCTCTTCAAAGGGACCCCACTCCCTATTTCATCTCAT

CTTTCCATTCTCTTTCCTCTCCCCATCTGATATATAGTGAAATAGGCTCAAATGTAGCGATGGCACAAATCTCCCAACTGATAACAGTCTGTTGATATCTGCTACCTTCTAGGACATCCCACAGAAAACTATTTGAGAGAGAAATTACTAGAATAATCGCGCTAGCAATGCTGGGAAAAATCTTTGGAGCTGTTGGATAAACCCTATAGA

GATAATGTCTGAAAAATAAATGAAGTAAAATCCTGTATGGAGCCGTCGATAGTGTAGCATTTTTTTTTTTTTTTGGAAATGCAAGTGTCGACTCAATTTTAAAATTAAAATTCCCTGACTTTCCCTTGTTTTCAGTCCTTTTCAGAATAATTCCAGACCACAGATTGGTTGATTTGCACCAAAGACGAAATAAAGTCATATTAGTCAGGC

TCAGACTACTCCAGGTATCACTTCTAGTACTACATTTGGTCCGTTGGTACTACAAAAAGTTACCTAGTTTGACATATCGCAGTACACATTTCACGATATTCTAGACTTTGATCTAGGATACTCGACTTAGGATTTTAACTGTGATTTTGTGACTTTCCCTAGACTGCAGCTTAACAAAATCACAACTAGATTTTCTTACTGTATTACTAT

TATTTCAAGCGCCATACATTTTTGAGCAACTACTAGGAACATGGGGTATTTATTTCGTATTTACTTAAACCGTGATAATTACAGTATGTACGAGAAGATAAATATCTGAATATTTTTACATAATGTATAGTTATGTCCAACCTGCCAAACTATCATTCTACAAGTTCTTCCACACAGCATAGTTCCCACGCCAGAGTTAGCGCCAGACAA

AACAACAACAACAACGGACATCGATCCTGTACGTGACGAATTGGATTCCCAGCTGACCTCAGCATCCCGAACGTGACGATCGCATAGGAAACATCTCGCCGTCATCGATGGTTTCCCAGCTGACCGTAGAGGAGCTAGGTAGGTTACTAGTTACTAAAGCCCAGGAAAAACCGTCGGTTACAGTTTAGATCTAGGACAAGAGAGGTTCTT

TACAAAAGTGATTGTGTTTGACATTGTAAACCAAAAATAATACGTAAATAAAAGTGATTGTGAAAGTTTGTTCCTTTTTACAAAGTGATGAACACTCTTTCAAAAGTGTATAATTTAATGTTATAGGGCGTATAAATAAACTGAGCAAAAAGCTTCTACTTTATAAAAACATCTGTTAGATAATTTTCCAGGGGCTTTTAAAAGCTAGTA

TGATTAAACTGAGTACATTGATAATTTTCATTTAGGGGCCGTCCATAAATGACGTAGCATTTCACTGATTTTGACACCCCCTCCCCCTCGTAGCATTTAGTCACAAATGCTGACAGACCCACTCCCCCTGGGAAAATACGTAGCATATCAAACAACCCCCCTCCCTTTTTATTTTTTTCTTCTTGATTGGGCTTAAAGGAACCATTAAAA

TCCAGAATTCCAAAAAAAGTTGTTTATTGATATTAATTATTGGCTGAAACATCAGGATAACCAGGCGTATGATAGTTTGATTTACCGTAGCGCTCAAAAACCATTTGGAAAACAGTTTAAATAAACTTATTTCCTGTTTAAACTACATAATTTAGGTTCCCAGTTTATACAGGTTAAATTTAATTTGTGGTACTTTTGCTGTTGTTAAAG

TCTTAAACAAGTAGACAGCTAAAATGTAGAACAAAAGGTTATCATCTATTTCGGCTTGATAAAGCTAACAGACTCATTGCTGTTGTTGCAAGTGTTCAAGATATTTTATAATATATTAAACAAAAAAAATATTAGGATTTTTTTTAAATTATTGATTATTAATTTTAATATTTAATTAAGTGATTGGAATGGATCTTCGTCATCATTCTC

TAAATTCAAATCCTTCAGAATAATATTAGTAATATTCGGAATGTTTAAAACACAACGGTTATATACAATCATCTTTATTAATATTTCAATTAAACTCCATCATCGTTCCCCATACTGAAATATTCTGACACAACTCACTAATGAAACAATTATATAATACAAATGCTCTTTCCTTCTGAATTACCAGTGATTTAGAAAACTGGATGATAG

ATATTTTCGATAACGCACTCGAATTTGATGTCCTCATTATTATTCTGGTTGTTCCATCAAGAGCATTCATACACAGTAAAAATAATGAGATTACACGTCATGTAAACTTCATTTTATGCGATGTAAACATGACGTCATGTAAATTTAAATCTGAAATCATGTAAAAAATGTGTTATATGTAATGTAATCACAGAGGATCCTGCTGGATTA

AATGACATATAATGGAAGTTTACATGACATATCATAATAAATATCCATGATATTCGATACTCCAATTATGTACATTATATGTCTCAGAAATTTACACGTTTCGTTCCGAACTGTATATCAAAACAAATCTATGAGTTGACTGAGAGGAGATCTTCTACAACATTGAATAAATATCTTGTTTTACTATAATAACGTTAATATTTGCCTAAA

TGAGCACATTAAAATCTTTTGAAGGAAACAGGCTAACAGAAAAATGAATCAAACTAAAACGGTTGATTCTATGTCTTGATAACAAATGGAAAAACAATATGTTCAAGTCGATAAAGCATTGATATTAACATATTCATCTTTTAATTTGTTTATTTATAAATCAAATAGTATGAACAGCAATATAATATATTATTCACATAACTTTTAAAA

AGCTTCACAAAATCTTAAATTTAAAATTTATTAGAATTATTTTAATATTATCATATCAATGTTATGAGGCTGAATCATCATTTATAACCCAAGTTTAATAAGTTACACGCAGAAAAATTTCTTTATGTTGAGATTACTATACTGTTAATTACTTTTACTTACGCACAAATAGTTCTTTCGAAGTAAAAATTAATTTACGCCAACACTAAA

TTAACATCAAATCAAATACACACACTTCAATGATTCAGCTTTCCTCTTGCACTTATATCAGTATAGTTTGGATAATACCACTTTCCGAAAACATTCACATCCCTAGACTTCAGTCAACAACTCACAAATCCTACTTACTTTTAACCTGGGGAGGGAGGCACCGATACACGAGTGGATACAATTGTTTACAAATTGCAAAGATAACTCCAG

TTTGCCAACGACAATCACAAGGCAGGCTTAGTGAAAATCGCTAGTTTTCATTTGTTGTACCTTCGATCTTTCTGGACAAACATTGAAAAAAGGGCCACAGTTTTTCTATGCGTTTAATTTTTCAGTTTTAATGAAAACTGTAAAAGAGCCTGTTCAATACGTTACATTAAATTAGAACAAACGGTGCTCTCATTGACGTTACTTTTGAAT

GTGCATTAATTGATTCTAATTCGATTTTGCTACTTTTGATGAAATGCAAAAAATATTCTTGGAAGCTAATTACGCGCTTATTGTTTGTCCATTTGTTTAAGATCCGGGCTCACAGTGACAGTTCGTGACAGCAAAATCTCCGGCTCACTGTGACGTAGAGAATTTTGTTGGTTTCTCCTCAGCTTTTAACTTGCAATGAATAGTGTAGGT

GCTCACCTTGCACGCAAGGAGACCACGTATCAGTTTACCAACGCACTTTGTTTTTACAATGTTTTTTAGATAGATAAAGCAGCAACGAATCTTTTCAAATATGATGATATAGCAAAACAAAATGTTGAGTGAACATTGCAAAATATTGCAAACTAAATCAAATTTACTTTAGTCAACTAAAATATAGTTTGAAGTTTGTAAATACAATAA

AAAATAGTTGTCAACAACTACATGACATAGTTATTTAAACAACAAATTCAATCAACTATATTTATAGTTATGTCTAAATTAACCTTAAAATATAGTTTATTTGACCGCAAAAATGATTACGAGCACTATATTTTGTTCTCCATGCCGATAAAAGTTAAGATAAAATAAATGTTGTCAATTTTCCAGCGCTACTGTTTTTTACCGACATTA

TTCAAAATGCTACGTCCACTAGTTTGGAACCCCTCCCTCCCCTTGTAACACATCGTCACAAATCATCAGAAGCCCCCACCTTCCCCCTAAAATGCTCGTCATTTATGGACGACCCACCCTGGCAAGGAAGATTCGTATGCATAAAATGGTAAAACCTGATATTCACAAAACTCTGAACCATCTATAATCATATAGTTAGAAATCTTCAAG

CGTGAAAGTTAGATACATACTAAACTATCTCAGAAATAGTGCTTAAATTTCAGAGAGATAAGCAAAGCTTTCTGACAACTTCCACGCTACCTGAGTAGATATACTAAAATGTTTTATGAGGCGCAAACTATAAGATGCATAATGCTCTTTTTAGTAAAAAAGGTCCTCAGAAATTCAGTCAGCTTTCTTTTCAGGTAAAATTCCCAGGTT

TTGCCGAAGAACTTCAATACGAAAATTGCTTCAAATAATACAAAGTGATTTCCAACTTGGAATTCTAAAACGTCTCTAAAAAATTTGAATAATTTCATAAAAAAAGTTTTAATAATTAATTAATTTATTTTTTTCTTCAAATACTTCGAATAATTTTTCCGATGATTCGTTACAGCTTATTTGTTTCATTACACTTTTTTTACTTTGGAG

GAATTATTGCAGTATTTATGCATATGAAAAAACTAATTGAATCTTAATACTTCATTTCACAATACAACATTCCTTCCTAGAGATTTTCTGGATTCATAAAAAAAAATCTTTCGAAGAATCATTAGAAAATGTTTAGGCGTTATTTCTTTGGTATGTCGAAACGGGATAAAAGCAAGTAAACAGGTGGGTGCGGTGTAGGATTTGCTCTAT

GGCATGAACCGCTGATGATGATCGAATCCATTGTTAGATCGAAACGTTCGGTAAATGCTACGGTTTTAAGTTATTTTAACCGAAAAATCCGATTAAATCGTCAACAAAACTATGAAAGCGGTAATGATCACAGCATAAACAGTTCTGGAAAAATACAGAATAAAATATGAGAAATCTCTATAAGAATCTCCAGAATTTCTGAGAAATTTG

AAGGATTTATTCTAAAAAACTGGGCCTTTGCTTGAGCAAGCAAGCAAGTATCCTTGGACCAGCGGTTTTCATATCGTGCTCTGCGACGCCTTGGCGATGCTCACGACAACATTAACAATGCCTTTATATATATAAATCAAACAGGGCTTCTTTTGACATTATTTCCACATTCTTTAACTTTGAGAGTTTGTAACTCTTAGAATTATGGAT

AGATGTTGATAATTTTGCCTATATTTAAGTTGTATATGTACGTAACAAATGAGCAAAATATGAGGCAAATCCATCAAGAAATAACAAAAAATGCTTGAATAGCGAAAACAGTGTGTCCTTCAAGACAAAAAAGGGGCTACGTTGGACATTTACTACAATTTGCTAATGAAATGATTTTCCTTATAACTTTCAAACTGATTTAATAGATTG

TCGTCCACTGTGATGTTATGGAACGTTTTCATTGATAAACTTAATGTAGAAAATTGCAAAGCTAGAATCAATTACACATATATAACAAATTTCAACGAAACATACCATTCACTATTCTCAGTTTGCAGTATGAAACTTAAATTTTCAACTTCCTCTTAAATGGAACTATAAAGTTACGAGTGCTCGATTTTCAAGTTGAAGCGCAAACGA

ACGACGCGTAATTACTAGGGATGAATGAGTTATGTACCAAAAACTCTTTTTTTTATTCTTACTGTTATATGAACGATATGTTGAGATCACATTAATACCGGTAGCAATTTAATACCAGTAAGTAATTTTCATTAAAAAGCAATCTATGAGTAAAGAGTTTAACAAGATCGTAAAGAAGTAAGTTTGCTTTTGTAATATGCTTTTTAAGCT

TCTTAGTAGTTTAGAGCTGGCTTAGAGTAGTTTAATTTAGGCATTTGAAGGATGCAACTGTTCTGTACTACAAATTCATGTAAAATATGACGAAAAATAGTTTTATGGATTATGTTGTGAATTTGGCATTAGTACGTATTGAAATATATGTGTTTATGTCTATTCGCTTTTACAAACATTTATTTTATTTTTTATGTTGTAAAATACACA

AACCAAAACATTATAATAAAATAAAGTCGCAAAAATAAAGACATTTGATCAATTATTTTAATAAAACAACAATTTAAATAAAACATGGGTAAAATTCAAACAATGTTTGTGTAGCCAATGTTTTATACCTTGTGAATATTTATTCGGGCTTTTTGAAATATTTTCTTGTTTATCCTGTTATTGTTGATGTTGTTTCAAAGAAAAAGGCAT

ATATTGGTTAATAAAAGTGCTGAAGACACAAAATTGCTCAGATTAATATTTACGCTGCTAATAAATACAAAAATTGGGTATCCCAAGTAGCCCCTGGAATCAAAGTAGCCCCGTCCGACGGTATATCAGGTTTTGCTTCGAAATACGCTTTTAATTATACCAAGCAAACCGATATTATATAATTTTTCAATAGTAGAAATTTATAGTACA

GAACATTTGGTCAAGACCGCTGTTTTATACTCTAGATTTGTATTCTCAAAAACATAAAGCTGGTTTTAAAGCATCTTTATCTTATGATATATTCGATGCAATAGTGGATGACGAAAGACAGTTTTTTAAGAGTGTTATACACACAGTGTCAAAATCTCGATCATTATTGAACTTAGATGAACCTCGGATGTTTTAACGAAACATTCAGCA

TTTGAGCTAATATTGTGTAAATAGAAGTATTCTGTTGGTGAATGTTTTCGCATTGATTTTGTATGAGATTTTGTGTATTGGCTGTGAAAAATTAGCTTTTCATCTGCCGTTTCAAGCCACGATTGAGGTATTCGACGAAAAATTGAGGTATTCGACAGAAAAATTCAAAATAATTTCTACAAACCACGAACTTTGAAATATATGCCCAAA

ATTAAATGAGGAGTTAGAAGCAAAGGATGTAAACAGCTAGTTTTGAGAAAATCGATTAAAACAATCAGTTTTTTTCATTTTATACAAATTGTAAAAAAAACAAGGCAATCTTCCACAAATTATGTTTTCTCTTTGACGGAAAAATCACTTAAAATAATGTGTGAAAGCTTGGAAACAGTAAAGTATGTTTTTAATATAAAAAAAGTTGCT

CAATCAAACAAACAAATCTTTTTCGATTCAGATTAAATTTTAAATTTAAAATCGAGATAATGAAACGATGACCTACATAGATCTTACTTCTTGCTGTGGTTTGATCACCGTTCTCTTAATCGTTTCCACTTTATTGGTATTGTTCGGTGACGGGTAATGATATCCGATATAAATTTTACGTTTCAACCTACTTTTTATTACGGTCATCTT

CAGAAAAAAATCCGCCGGCCGTTCGTCCGTCTCGGCAGTATACTTAATTCTTACTTCATAAATTTGTATTAATTATAAAAGTCGGTTTAAGCGTATGAAAAAATCAAGAGATATCAAACGGGAACAAATGCTCCATATAACAACTTGATTGCAAAAGTGCTCGCGAGCCAAAAGGCTGAAAACCCTGGATTGAATTATAACTAAGCACAT

TTTAAAAATTTTAAATTCTAGAAAGTAATAAATAGTTTACTATGGATAATAGCGTTTTCTCTTTTCTTTGCTATTTTGAAAGATTAATTACCCATTTCATCCACCAAATTTATGGTATTCAATGTAAACATAATGATTCTTACCTATTGTATTGCTGAGTTGCTCTAATCTCTAACTTTGCAGACCAATCAACTCGTCTGATGTTGTAGT

ATAGTGAAGTGATAGATATTTTTCCCTGACTTGTAGTGAATTTCCCTGACTTTCCCTGACATTCAAAATTATAGTTCCCTGACATTATTATCTTTATCTTTATTCCCTGACATTCCCTGACTTTCCAGGTTTTTCCAGGTAGTCGACACCCTGGAATGAAAAAAGAGGGGAATTAGATAATTTTTCAAGGAATCTCTGAAACAATTCTTA

GAGTAGTTCGCAAAGGTATCGAAACAAAGATTTATTTTATATGTTTTAAGAAGTTCCAGGTTGTATTTCATCAGAATTTTCTAAAACAAAATTGACTGTTTAACGATAAGCTTTGCGACGATCGACAACGCCACCATATTTCATATAACGGAGGCTATTAGAACTAACTTGGAGGTGATGATTTGGGTTATTTGGCAAGTTGGAGTTAAA

ATCACCTCAAACACTAGCACAAATTTGCGGTGGGATGTTGACGTTTGATCACAGATAGGGCAATATTCAAAACTAAAATGGCAGCAGACTAAAGGCTCCAAATTCCTCGGATGCATTTATGGATGGACGTCTAAAAAAAGATCATTAGTTAATAAAGAAAATTTTGTTTTTGCAGAGATTCTCAAATGAATTCTGACGAGGAGGTCTACT

TTTTATAAAACATTGAATATCTAGCAGATTATTTATAACTGTGAGGCGCACTTAATGAGTTATAAATTATGAATTGGAACTTACCATTTTTCGGCAACCCCGAATAACGTTCACCGCCGCATTGAAGCGATCCTCTATCGACATGATTCGGTTCGATTAGTTCTCGTCTTTGTGATGTCTGTTTTGGTGTCTCTCTTTTGCGCCACCAAA

CAGCCTTGCAACCAGCAAGCTGATTTCACGCTGATTTCGTCGTCGCGGTTTTTGTTGTTGAGAAACTGGCTCAAATAATAAACGCCGAAAAGCCAAAGGTCTTCACGGACAACGCGTCCAGTTGTAGTGGATAGATAGATGCTGGCCTATAGAGAAGGAGGTCTCTGTATCTCCCTCTAACTTTCGACGGCTAATAATGCGACACAGTCG

CAGGTTATTACGTAAAGACGCGTTTCGCTTGGCCGTTCGATCGGTGGAGCCGTTTTACTTCTCTAACAGAGCTTCCCTCAGGCGGTGCTGGATGGCTATGTCTGTTTGTGGCGTTCCACGCTTTGAATCCGGTAAAGGTATAATTATATATTATATTAAAAACTGTCGACGGCTTATTGATTGCACCAGCGATGACCTGCGTGTCGATTA

GAAAAGAGGGCCCCGTCAGTCGTAGACCCAATTCCTCAGGATCACCTTTCCCTGCCCTCCGAAACAAGCACTGTCTATTCCTCTTATTTACCAGGATCTTCTTTTGTCACCAGCAGCAGATAATCGAGCAGCTTTATTGCTTTATTGACATTCACGCTCAGGTACTCCTTCTTCAAGTGCTTCTGTCCCCTTATTTCCGATTCCAGCTTC

TTCACTCTCGCGGAACGACAAACGATTACTCGCATGAATGACGAAAAAGCAAACTGCCAGCCGCAGAATTTCGCGCACATCAGACCAGAAGCGCCAAGACAAACAACAAATTTCCTGCAGCGTAATACCTATTCTGTTGACCACTAAGAACCTGTGTACACTTTTGTCTACAGAGCACTTCCATGAAACACAAAACTACTCATTCATCGG

TGGCAATTTGGCTCAGCTCTTGTCACATTGATGACCACTTTTTGTTCTTCGTCAGAATAATGAGGAACAACGTCGAGTCGAGCAATGCATGCCGTCCGGGTGAGGCCACAAAAGATTAACTTCATGAAGCGCAACACAAGACAAAAAAAGATTCTGGTGGCTTCGAATCTTCACAAAAATCTTCGCTTCTGCACTTTCAACGCACAGAAC

ATCCACACACGACAAACGGCTGAAGAAAACTGACGGACAGAAAAATGACAGAACGGTTCGACGGACCGAATGAACGCTTTGGGTGAACAAAAAATGTTTGGGGTTCGACCAGTTTCATACCCGGGAATCGCTCTAATTTATTTGTTATTTTAGTATTTAGTATTTATTTCATTTAAGCTTAAAGCACTGACAAACAGACGTCACACTCTC

ATCATTGTCCATCGACCACCTTTTAACAGCCGAGTTCAAAAATATGGTAGGTGGCCAATCCGCCACCCGCAGCGCTAGCATCGTTTTGTTCGCGTTTGACGTTTACACACTACCGCCATCTGGTGGCTGTCGGCCAAACACGCTATTTATTTTTAGCATTGGGCACATGTTGTCATGACTATGATTTTGATCAATTTGTTCTAAGTGTTA

CGTCTGTTTGTCTGTGTGCTTAAAGTAAGTCTGTCGGTTATACAACAACAATTGATATTGCGACTTTAAGAAATAACAACTATACGGAATAAAAGTTAAAAAAGATTTGAATGTCATGTAACATAAAGGTTAATTTAACTGATAATATGGTGTGCACTGAAAGCGCGGCTTTGGATGCGCCCAGTGATACCACATACAGATTTATCTGTA

ATTACACCGATTTTTTCTGTTTTGCCTACAGATATCTGTGATGCGAATCACAGATTTTGAATAAAAAGATTTTTAAGATTTTTAGAATATTTCACGAGTAAATAAAGATATAAAGAAATGCCATTACTCTGTTTATTTTTTCTCAAAAATTTAACTATTTAAATACTATATACTGCTTCGCGACTAAAATGGGTGAACCAACGTGCGAAG

TTTGATTTTTCCCGACCAACTTCGCCGCGTCGCAACTCGATTCTCAATAGTGCGCATAATGCACACGGTGGAGAGCATAGATGCGCACTATAGCGAAAGTGACTCGCGACACGGCGAGGAAATTGGTCAAAACCCAGGCTGCCACGTTGGTTGACAATCATAATATCCCGTTACAAAGAGTTCTTCTCGATCTAAACACATACGTTTTTG

GATTTTTACGACACAGATTTTTAACAAAGATTTTGGAGGTTGACCTAAGATAAAAAAGATTTTTTCAGCCGAAATCACAGATGAGATTCGAAAAAAATGTGGCAACACTAAGATGCGCCTATCGTTGGTAGCTGTCTGGTCACACTAGTCCCAACTAGTCTAAGCAATTTGTTGCAGTCCCCAACTGGATCATATCCTTCTGCCCAGGCA

GACGGACCGAGACCTTGTCGTTACGAATAAAGATGGATTGGATTGTTTTCTTCTGTTTCAGCCGCAAGGCCGGAGTTACGGATGCGGAAAGCTTGGATTAGAAAGCATTTTGTTAACTGTAAACTGGTATTCCAGAGGAGCAGATGATTTTAGATTACATAAATTGCATCCTTGTGCTTTTCCAAGTAGCGCCTGAGTACTAGCGACCTC

CTGTAATCGATACAATCATATTATCATTCACATATATATATATATATATATATATATATATATATATATACTATATATATATATATATACATATATTTGTTTTATTCGATCGAGAGCTTAACCAACAAAATCCCCTCAACATCACAAAGTTCTGCCATTAGTTCTGATTTAATTTCCCCCGATTTCCGATTCCTTGTGCGCGTTTGTCCA

TGGATTTACATTTTTCTGAAGATGATCTATCCGATAAGAGCGATTGGGTACATCATAAAGATTCCGATATATCTGAAGAAAAACTCTCGATTAATTGTTATGTATGGCTGTTTCATACCTAGACAAAAGAAAAACACTCACCTGCATCTAAACAATGTTCCGACGAAGCCTTTAATCTCTTGTACTACACAATTAGTCCAGCTATTGTTT

TCTTTAAAAGAAAACTTGATTATTTATAAAGGAGATTTTGAGCCAGAAATATTTACAAAGCATAATTGGGTCCTAAAATTTTAATGAAACCTCGTTTTTATTCAATAACACGAAGAGAATCTTACTGCAACTTCTACTTTAATTTTTCTCGCTCAAATAACGGCTTTATCATGTTTAAACTTTAATTTAAATTGGGTCATAAATGAACCT

TGACATACTTAAAAGATCATGTTTTTGACGTTCAGCAAGTCGACAAAAACACCACAGGGGGTTTAGTTTTAACACTGGGGTTGTTCTATATGACATTTCGGAAGGTTACAGGAAAACAAAATACCCAAAATTTGAGTTTAAGCCAATATATAAAAAATGTTTTTGAGTTCAAACCAACGGAAAACATTGAAGAGTGAGTAAACATGTATT

TATTGGCCTAAACTTAAGCGTTTAACACTAAAATTGGGACATGGCTTTGAGACCCTATTCCTTCAAAACAAACATGACTGGGTGCTTTTCTGTACCCGGATTCAGAAGTGAGTGCTGCGGCTGCAAGACTCAGGTCGGGAAACGAACATCATTACTTATGGTTCTTCTAGGTTCAACAAATTCTGCGCATTGTGTGACCATTTCGAATAA

TAAATGGACATCACTTAAGGCCGCACGGGACGTCATCATAATCACCTATCCATTTGAAAAAAGCAAATCCAAGAACCACTCCATATATCGACGTGAAAAATGTCACTATATATGTAGTGCACAACCCAATAAATTTTTCAGTTCATCAGTTCACTTAAACTCGAGATTTTCCTCCCCAAAGTTTTTGATGATAATTAGTTTAGAGTGGAG

ACAAAAGATAGGAAAATAACACGGTCTCCCATGTCACCTTATAATAAATATTGTGAAAATAATAGCTTTGATGCTGTTTCATCATACCTTGTAGACCACACATGGAATGTACAACGAATAGGTGACAGAATCAGCATTGGTGCAATACGATTTGAATCATTCCGGCTACAATAAATTTCTATGCGGGAAAAGAGCGATTTTCTGACGACG

TCCACTGTTCAACCAACAAACTTTTTCGGGAAGCGAATGGACCTATGCACTAGTTCATTATCTGACGTTTAGCGGTACCGTGTTATCTAATAGCGACCATGGAAACCAATAGGCCAGGAGAAGTGGATAGACATTTATTCATTTTCTGTCAATTGGGTTGTTTAGCGATGACAAAGTTACAGTTTTCATAGCTTGATCGAAAGATCACAT

TTTTTTAAAGTATAACACAACTTTATTGCGCTTCAGTATCAAATTTTGACATTATAAATGATTAATGAAATTTCAATACGTAGACTCAATGACATTTCAATTTACATAATGGCAGCTCGTCCGTAGTAAAATTTGCCGAGGGTGATTCAAAGCGATTTCCATACTAACTAAACGTGTTTTAAAAATAGTTCCAGGGCACGAGAGAAAATT

ATGAAACTTTGGATTCACCTCCATTTTTAACATTAGAATCAGAATACGTAAAAACTGGATACCCTAAACAAAAACCAATTGGGTTGTTTAGTGAAAAAAATTCACTGTTGTTCGAAGCTTGATCGAAAGAGCTCATCTTTCTAAGTATAATACGATTTTATTACTTCGTATCTAATTTGATATTTTTAAATGATTTAAAAGATTTCAATC

CGTCAAATCAATGACATTTCAATTTACACAATGACAGTTTGTCAGTAAAATTTGCCGAGGGGTGATTCAAGTGATTTCCATACAAATTTCAAATGTGTTGTTTTAAAATAGTTCAGGTAACGGAAAATTGTTAAACTTTGGATTCTAGTTTAATTTTGACGTAGATTCAGAATATGTAAGAAAATTGATACCCTAAACAAGCCAATTCTC

ATACAAAATCTTCTTTTCTCTGACATAAATTCACTCTAGGTGAACCACTTCCCCCTGGCCTATGAATTTGGCCCCGCTCCAAACGTCAGGGACAATGAACTCATGCATAGGCCAGGGAAGTAGTTCACCAAGTAATTATCCAGAAAGAAAAAACAGGCTATATGAAGTGACTGAAAAATGAATGACAAGTTCATCCACTTCTCCTGGCCT

ATAATTCACATAAATAACACGGCACCGCTTAAGCGTCAACGAGTAGACGCGTGCATTGATCCATGGACCTGTGCGGTTCACTGTTGTTTGAATTTTAGCGATGCCATGTAGGTGGCCATGGAAACTAGTGAATTCGGCACCGCTTAAACGTCAAATTAGTGAACTCATGCATTGGTCCATGGATCGATGCACGAGTTCACTATCTAACGA

GCGATGCCGTGTCTTCACGTGACCATGGCAACGAGTGAATTGGCACCGCTCAAACGTCAAATTGAACACATTGCATTGGTCCATGGATCAATGCACGCGTTCGTTGACGTTAAGCGGTCGTGTTATCTTGACCATGGAAACCAGTGGTCGGCACCGCCCAAACGTCAAATTCGTGAACTCGTGCATTGGTCCATAGGTTCATACCACCCT

TCTTGATCATTCAATAATCGGGGATGTAACCCTGCCAAGAGTTCATCATTGCTTCACATTAATCTTCACGGCGGTCGTTCTCGCGTCGTCGTCGTCGGCAAAATGTCAAAAATGGCATTGTTTATAACTTTCTGCGCGTAGTTAGCACACTGTATTGCATCCAGTTAACAAAGTTTACAAAATTTCCATAGATTTTAGTGTAAATTTAGT

TAAATTCTATAAAAAGCCAGATAATTCGGTGCTATGTACCATCAGCACCAAAGACGGCTGTGGCCATAGTGGACGATATTGAAATCATTTCCAAGTAGGTGGCTTTCATAAATTAGCTAATGGTTTGTGTTCAAAAAGATTTTTTGAGTACTAGGAACTAATCGAGAACACCATCGCTCAACATCAGAAAATGTCAGCGCCGATCACGGC

CAGCTGGTGGAACTACTAAGGCCCATCCAAAAAGACAAAGAACTGAAGGCCACCCTGCAGCTGGCGGCGGAACAAGCCGGCATCGAGAAGAAGATGGACGGATTGCGGGAACAGGTCAAGGAACAGATGAAATCAACCAGCTGCAGAAACAGCTGGAAGCCGAGCATTCTGGCCACGTCCATGTTCAGGGCGCCAGTTGAGCAGTATTAA

CAAAGCCGTCAAGCGACCAGTCTCTCTTCAGAGGAGCTGATAAAGTTCGCTCATCGGATCAGTGCTTCGAATGCAATCTGCGCCCCGTTAACGTAACAACAGGGTGACCTGCGGAAGCCTTACCGACGGACATCAGAGATGCGGTTGGGATTCCTGGGCAAATCGGACCTCAACATCAGCGGGCACAACGCCCGAATCAGAACAACCTGA

ACGAGATGCAGCGGAATGCGGCCGGGGCGGGGCGGGAAGCCCAGGTGTATTCCCGGCTCTGCACAGGGGAACCGGTTCGCGTGGCATCCGTCCGGCGAGTTGCGCACATGACGATGGGCAGGGGCGGGATCGGTTTCACTGGATACCCGGTCGCACAGATGCCTCGCAGGATGATGTGGAAGTGATGTCCCCGGACAGTTCAGTTCAGCT

CAGTGATTCGCAGTGAGTGCTGTATACTCGAAAGATGGCTGTTTCAGTGAGTATTTTTTAAATATTTAATAAAGTCAATGCTATAACGTGAAAAAAATGATTCATTGTGAGAAAGTCTTCTTCCTTCAGGCATTAAGTTTCTACTACTGCAGCAAGCACAAAGAAGTCGAGGGGGATATCGAAATTTTCAAATACAGAAAACTGTCGAAC

CCCCGCGTTGTATCTATTCTTCTGGTACCGGTACTTCAAAAGTGTAGTTAAATTAAAGAGTTATTACTTGGAAACATGATTTTTATAGTGAGAACACATATCAATAAATTGAAAATTGAATTAGAATTTGTAATCATTTCTATGAAACACGTAAGCTTGAAATATATCGTGAGTATATCGCAAATTGTTGTCGTTATGGTTTAAACAGTA

ATTAAACCACGTGATACACAAAGGTCGATAATTCGAACCATTGTCTCGTGTGTTCTCTATAATGTTCGCAATCAATCATGTACTGAAACTAATGTTAGATGATACAACCACTCTATGGGTTTTCAAAATTACTATTCTAATTCTGATAAAATTTAATGTGTTTGTCTTCACGATTACTGCAATGTTGAAATTAGCATGCCAGGGGAATTG

GATGAAATTTACTATCATTTGCAGTCAACAACTATTATCCCGGTATTAGAGATAAATACAAGATCGTTAATTTCTAATGAACAATGTTTTCATTTTTTACCATGTGCATGATGGTTTAAACCACAAAATCGTGGTAATAATGTCTCTATCCTCTTGTCTGTTTTTCTCCGTCGAAGCTTATGGAGGATTAACACCACAGATTGTCGACTA

CCTGGAGAAAAACCTGGAAAGTCAGGGGATGTCAGAATTCGATTTTGATCTGAAGTCAGGAAAGTCAGGGAATTTCACTAGAGGTCAGGAAAATATTCAACTTTAAACAGTTGATTGGGAAAACGCTTTTTATGCATATGAGAGTAAGCTGTACTTTACTCTATCTATACTTGTTCTTAGAATCAAAATTTTCTGAAATGTAAAGCATCA

ACAGTGATCCTGAACAATTTCTTCCAAAGTTGGCGTAATTGGCTTTATTAATATTTAGTTCAAGTTTCCTAGATCAAATTATTATCTTCTCTTCAAAATCTTGAAGAAAAATGGAATCCCACTTCATCAAATCATGCCAGTCTTTACAAAGTTTGCCCAAAATATCAAGTTAAACTGATAAAATTAAATAGAGAACAATAAATGAAGTTT

CTCTCGTATTTAGGTCGAAGATGAAAACCGAATCTCAAAATTTTTCAGAGCACAAAGCTGCAAGACAGGATGTTTCCTGAATCTTTATCATGTCGTTTTGGAAACAATATAGATAATACCGTAATCGCAAAGTTCTTCAATATTGATATATGTCAATTTTTTCGATTTTTTCAATTTCAATTTTCAGGATTTTAGGAAAATATCGTCAAT

CAACATTACTTCAGGGTTTTATTCAGGAGCTTACTTACAGATATATTCAGTTTGTTTTTTGTTCTAAGAAGAATATGGAAGTTCTCAAATAATATGAATCATTATCCAAGATATCTTTGTAAATCCATGAAATCCATGAAAGCAGAAATTTCAAGAATTCAATTTGAAGGAACTTGCTTGATAATCTTTTGATAGTTTGGCTGAGGGTGC

CAGAGGTTAGTCGGACATCGTCCGTAGATGGGCTAGATCGAGCTCGAAGCCGGTCGAAATTTAAATCCTTTTTTAATTGTTTGTAAAGACATATGTGTTATAATCCCGTTCTCATTGTCGCGTTCTGTAAGTGTACATGAAATTCTTATGTTAGCACAAACACCCGAAAAAGTTCATAGAGATACTCATTGAATACTCATTCTTAGTGGA

AGGAGTTAAATCGAGGGCTTGTTGGAGGCACGTCACATTAAAACGAACATCTGAAAACATATATTAGCACTTGTTTTTTAATTATGAATCGTGGTTTACGGCCAACCAGCCGAGAGTTAACAACTACTGAAAAGCCAAACATTACATATAATTTACGAATTGGATTAGATGGACAAATTGATGTGTGGAGATTGCGAAAAATTTTACGTC

GACTCCCCGATCTCTAGTTGGGGCGCGTTACCACTACGCCATGAGAGGACTCATGAACGCAGAGTTAACCTGAATTCGATTTCAGCTCAATAATCACGTGGTCCTTTTTCGCAAAGTGCACCTCTTTGGAAGAATTAGATGCCCATCCAAACACAACACTTTCTATATATATCCAATGCCTAGCCGAGAGCGCAATGTTTTTGATATAGG

AATAGCACACTACACTGGCCAGCAGCTGCGCTGGCTGGGGTTTGTGTGTGGGCTTCCGATGGGTCGCGACGTTATTATTAGCGCATTATTCAATTCAGAATTCATTCCGTTAAAATATTAAGCTTATTTCTTAGGAAAACGGTTTGCTTGGATTTATAAAAATCACACCGGAATTCTAAAATTAACTAAAATAAATTACCGTCAAACAGG

GCTTCTTTTGACATTATTTCCACATTCTTTAACTTTTGAGGATTTGTAACTCTTAGAATTATGGATAGATGTTGATAATTTTGCCTATATTTAAGTTGTATATACGTAACAAATGTGCAAAATATGAGGCCAAATCCATCAATAAATAACAAAAATGCTTGAACAACGAAAAAAAGTGTGTCCTTCAAGACAAAAAAGGGAGCTACTGGA

CATTTTCACAATTTGCTAATGAAATAATTTTTCCTTATAACAAACTGATTCAAAGATTGTGTCGTCCACTGTGATGTTATGGAACGTTTTCATTGATAAACCTAATGTAGAACATTGCAAAGCTAGAATCAATTACACATATATAACAAATTTCAACAAACATGTCATTCACTATTTACTCAGTTTGCAGTGTAAACTTAAATTTTCAAC

TTCCTCTTAAATGGAACTATCAGTTACGAATGCTCGATTTTCAAGTTGACATAAACGAACGACGTAACTACTAGGTACTGCAATGATAAATGAGTTACGTACAAAAACTCTTTTTTTCTATTCTTACTATTATATGAACGATATGTTGAAGGATCACATTAGCCGATAACTAATTTAATACCAGTAAGTTATTTTCATTAAAAACGCAAT

CTATAGTAAAGTTTAGCAAGATCGTAAAAGTAAGTTTGCTTTTTTTGTAATACTTTTAGCTCTTAGCAATTTAGAGCTGGCTTAGAGTAGTTTAATTTAGGCATTTGAAGGATGCAACTGTTCTGTACTGCAAATTCATATATAAAAATATGACGAAAAATAGTTTTTTAGAGATTATATTTAATGAATTTGGCATTAATTCACAATATT

GAAATATATGTGTTTTATGTCTATTCACTTTTACAAACATTTATTTTATTTTTTATGTTGTAAAATACACAAGCAAAAACATTATAATAAAATTAAAGTCACTTAAAAATAACACCTTTGATCAATTATTTTAATAAAACAACAATTTTAAGCAAAACAAATCACAAGATTTTCATGAAACATGACGATTCGATAGTTTGTGATGCTCCC

AATAAGTTTCCAAGACATGGGTAAATTCAAACAATGTTGTATAGCCAATGTTTTATACCTTGTGAATATTTATTCGGAAGGGCTTTTGAAATGTTTTCTTGTTTATCCTGTTATTGTTGATGTTGTTCAAGAAAAAATCATGCCTAGTGTGACATATATTGGTTAATAAAGTGCTGAAGACAAAATAGCTCAGATTAATATTTACGCTGC

TATTAAATACAAAGGTGAGTGTCAAAGTAGCCCCTGGTACCGTCCGACAGTACCTGTAAATATTTATGTTGAAGAAATTTAAGAAAAAATAGACACATTTGGGAAAAAAGTATATGATCCCTAAACATTTTTTGTAGTATTTCTGTAATATTCCCATAATTCTAAATTAATTTCTAGAAAAATCATGACTAAAAAAATTCTGTAGGTGAA

TTCGGTGAACAATTTCTTCAAAGAAGAAATTTTCTTCCATTATTCACCAGCAAGAAAATTTCTTTCTTCAAAAAATACGTGCCGGGATTCGCCTACTAGAGGTGTATTTAGGAATTCTTAAACTAGTTTAAAATCTTTGGACTGATTTATGGAGGAGCTAACGTAAGAGTTCATAAAACATAATTCTTTGTGATATTAACGTCATATATT

TGGATAAAACATTTGAGGAGGTTCTGGGAGAAATATCATAGAGTTCCTGGAGAAAAGTACAGAAGGTTACTTCGAAAAATATATTTGAGAATTCAAATTGGAATATCTGAGAATTTTGTTCAATTCCTTGATGCATTTTCAAAAGAAATTTTTTTTAGCAATGGTGTAATTTTACCTGAAGAAGAAGTAGCCAATTCAGAAGAAGTCCTT

CCAATAAAAAAGTTTAGTAATAATCTTAATAGAGTGACGTCTTATGTATGAAAAATTGTGTAATCAGGTTGCGGAAGGTGTCAAATTCCATGCTTTTATGCATGAAACTTGTGAAGTATATCTGAGGTAGTTCAATATCCAAATACATCGTTTTCTACAATTTCAAAGAAACTTTTGCACTTGAAGATTTCAATATGAGGTCGATAGATG

ACTCAGAATTAACTTAGCTTCTACTCAGCTTTTTACCGGTTTATTAACCTTCCAGTCGTCGCGTGGTTTTGCCACCGTCAGAACCACCACGCTGCTGTTATTAATAAATAGCGAGCTTTTTCACTAGTGCCAGTTTCATCATATCAACTTGAACAAAACTATCATAAAATTATTGCTCAGTTTATTCATACGACCCAAAACATAAACTTT

ACCAAACAAACAAATCAACATGGTCTGCTTTTTCTGGAAAATGACTGGAAAATCAGGAAAGTCAGAATTTTATTTTCAAAATTGGGTCGACACCCTGCACCAGCGCGGACATAAAATCCTATTATTTTTAAATAATTTTAGCGTAATATCAAACAGTTTCATCATAATACATATATATCCATGACGCTATTGATCCAGATTTGGTAAGAA

ATGTTCATTAAAACACAAAACAAACAATTCGCTCTGGCTTGTTTTCATGTTGCAAGATCACATCGATGTAGCTTCCAGAGCCCGATTAAGTGGTCCAAAACAGCATTAATGTATTCTATTGTTATTGGTAATTATTCTATCAATCATCGATTTGTTAGAAATTCAAAATAGGTGGAAATAACAGGAGGTGTCTACTACATGGAAAAACTA

GAAAACCTGTAATTATCAGGGAATTTTATTCAACCTGGAAAAAACCTGGAATACTCAGGGGAATTTGGCTACACTCAGGAAATTATTTCCGAAACAATAGTGCATGGTAGCATGTGTCGTTTTGTGGAACAACATCTTTCCAATCAAAAATTTTTGGCTGCGCCGCTGTTGTTTAATCTTTTTAACGATTTTTATTTAACAACTACCGAA

AAGCTAAATATTACATATAATTTGCATTCCAATTGGATTAGATGGACAAATTGATGAAGATTTGCGAAAAGTTACACGTCTTCTCAGTGAGAATCGAACTCACGACTCTCCCGATCTCTAGTTGGGGCGCGTTACCACTACGCCATGAGGAGAGGACTCATGAACGCAGAAGTTAACTAATTCGATTTCAGCTCAATAATTACGTGGTCC

TTTTCGCAAAAGTGCACCTCTTTCGGAAGAATTAGATGCCCATCAAACACAACGCTTTCTATATATATCAAGATGGTTAGATGGAAAAAGATAACCGACTCCGGCCTTTTCTAGGCGACACGAACAGCGCAGCTCTGGTGAGTCAAAGCCAACTAAAACATAATGTTCTCCTAGTGGCCTGCAGATGTCTGAAATGTTCGTTTATGACAT

TTACACTTAAAAAAAGATACACAGCCACACGAGGTTCGTTCGATATTCCCAATTATCTCTCAATTCCATAAAAAGTAAGAAAAAAAAAAGAAAACCAGCACAGGGTGATTTTTTAGCATTACACAAATTTTGTGTTGGTTTAGAAAGAGCCAGATTGTGTGTTGGTCGTACTCAATTCTGTGTATCTTTCTTTTAAGCGTGTTTGTTTGA

TTTCTTCCTCTTGCTCATGAACACCAGAAGTTAACCTGAATTCTAGTTTAAGCTCAATAATTACGTGGTCCTTTTCGCAAGTGCCTCTTTCGGAAGAATTAGTGCCCATCCAAAACGCTTTCTATATATCCAAAGATGGTTAGATGGGAAAAAGATAACCGACTCCGGCCTTTTCTAGAGCGACACGAACAGCGCAGCTCTGGTGAGTCA

AAAGCCAACTAAAACATAATGTTCTCTAGTGGCCACCAGATGTCGAAATGTTCGTTTATAGCATTTACACTTAAAAAAGATACACACAGCCACGGGTTCGTTTCGATATTCCCAATTATCTCTCAATTCCATAAAGTAAAGAAAAAAAAAACCAGCACAGGAGTGATTTTTAGCATTACACAAATTTTGTGTTTTAGAAGAGCCAGATTG

TGTGTTGGTCGTACTCAATTCTGTGTATCTTTCTTTTAAGCGTGTTTGTTTTGATTTCTTCCTCTTGCTCATCGTTACACAATGATTTGCATTTAGCCGAAAATAAAGTTTACACGCTTTTCACTCAACAGATAGCAGTAGTGTTACTTGAATAGTGTGTCGTCGGCAAAATGTATGGAAAAATAAAGTCAGTCATCTTATTCCTTTTAA

TCATCTTCGATATATATCCAATGCCTAGCCCGAAGGCATTGTTTTTAGGTAGAGATAGCACACTACACTGGCCAGCAACTGCGCTGGCTGGGGGTTTTCCTGTGTGTGGGCTTCCAATGGGTCCGCGACGTCCTCAACAACGACCGGTTACGGAACGCCAAGGTCCGTTGCTCGATAAAGCCGTAAAACCACGATTCATAATTAAAAACA

AGTATGGCTGGTATAGTGTGCTATTCCTATACCTAAAAACAATGCGCTCTCGGGCTGAGCATTGGATATATATGAAAGCGTTGTGTTTGGATGGGCATCCCTAATTCTTCTGAAAGAGGTGCACTTTGCGAAAGGACCACGTGATTATTGAGCTGAAATCGAATTCAGGTTAACTTCTGCGTTCATGGAGTCCTCTCATGGCGTAGTGAT

AACGCGCCCCAACTAGAGATCGGGGAGTCGTGAGTTCGATTCTCACTGAGAAGACGTGTAACTTTTCGCAAATCTTCACATCCAATTTATCCATCTAATCCAATTGCAAATTATAATGTTTAGCTTTTCGGTAGTTGTTAAACTTCCACTTGGCTAGTTAGCCGTAAAACCACGATTCATAATTAAAAACGAGGATTTTTATTTATTTGC

CATTAACATGTTTGAACAAAAGCAAAACTTAAGATTGACAAGTTGGTAAAGACAAGTACAGTTTTAGTTGGGTGTCATTCCTGAGCATAGTTAAAACATCCATCGATAAAAAAATCGTTTGTTACAAAATAATAAACTTAAATAAATAAGGAAAAGAGAAAAAATACAAAAATAGTATAGTTTGAAGGTTAGCAGATTTGTTTATAGTCT

CGGTCACTATTTTAATGCAAGTCTATTTTTGGTTGGTCTCCTTAAGTGTCTATTTTAATCAAAATGGTCGCTAAACTCACATTTTCAATCTGCTTGCAGTGGATCACCAGTGCCAAATTAGAGACTCAGAGAGAAACCTACAAAGACTGTAACTTTGATTATTCCACGAGTTCTTAATAATATTTTTCTCAGATACCGAGAATGCTTCAG

GAATTCTAGTAATTTTTCCATCGTTTTTTCCGGAATACTTCCAGGAATGCTAGAAAAATTTCTTCAGATTTTGTCCAGAAAATCTAGAGCACTCCACCGCTGCCACAAAGGTTTAAAAAACCTGAAGTAATTTTCACAGAGATTATCCTGAAAAAAATCTAGCGATTATCACGATGTCCTCAGTTGTTTTCCCACTGATTCTTTATTATT

TCTAGAAATTTCGCACAGATCTATGAGTTCTTCTAAGGTTTATTCCAGAGTTTGACATTTTTCAAGAAACTGTATAATAATTCCTACGCCATTTTTTCGTTTATTCTAAACCTTTTCATAAGAATCCTTTTTTTAATTTATTCAGGATTACTCTCAGGAATTGCTTTAAAAATTCTTCAGGTATTGTTCGATGAAATCAAACTTATAAAC

TTTTCAAGAATTCCTCGATAAATTTGACATAGGATTCCTTATTATATTTCTCAGAAATTCTCCGCATTTCTTCAGCATTCATTAAAAAAAATGTTCAGAAAAAAATATTTTAATTTTTTAAATGAAAATTCAAGTCTCAATACAAGTGGCGGGAAGGATAAAAACCAGCACTTGTTGTGTCACTTCTGTTAAAAACTACGCAAATTAACG

ATGTACTACATAAAAACGCTAAATATGATGCTCACAGTGCGATCAGTTCTTTCACGAAACTGAACAAGGAAAATTTTTCAATACGAATTACATCACAATAAAGGGCACTAGTCAGCGATTAAAAACAAGAACAATCAAAATTTGTGGTGATAACTTTGGAAGAAAATCACCACAATTATTTTAGCGACAAGATTTTGCAATTTAAGGTAC

AGGAAATCGTAAACATTATTATAATTTTGAAAAACTTTCGCCACCTAACCTCACTTTTGATCACCAATTCGGTTTTGCGTTTCGATGACTCCGGCTCTCCAACTGGAGACGCGATTTGATATATGATATGGGCTGGAGTGGTTGGTTGCCGTCTTGATTTTACGGCTCATATAAGAATTATGGAATGTTCATACAAAAAGCGTTACGAGG

GGGAGGTGGGTGTGAAAAATTGCCATTTTCGGTGGTTTGAAATATGTGAATGAAACTTTGTTGAAACGCCTCATAAAATGCCATATGTCGTTTTGGTTCAGTGTTTCAAGTCCACAGTTGCCCATGCAGCAAATTCCAACTGATGTGAACATTACTGTGAATCGAGTCCAGGTGGATCGTAGCGTCATTTTCGTTTGATCCATCTTAACA

CCGACGGCGGCGCGCGCGAAGTAGAAAATTTGGGTCAGTTTTGTTTTACAGTATTTCCGCGGTTTCTTTTCTCGCAATCTCAAAAACAGAAAAACCTCTTTATTTTTACATCAATAGTTACCTGTTAAAACAAAAGCCTGAAACAATAATTGGTGATGGTTCGTTGATTGTGCTGCACCAAAGTTTATTTCAAGAAAAGCATCAACGATG

GGAATCACCGGTTTGGTCAATTCCTGGAGAAAGCTTCCAGCCGATGTCACCTGCGAGCTCCGGGGTCAATGCGTGGCCATCGACAGCTACTGCTGGTTGCACAAAGGGCCTTCGCCTGTGCGGACAAACTGGCCCGTGGGGAGCCAACCGATGTCCATCCAGTACTGCCTGAAGTAATGTAAATATGATGTTGCTTTCGCACGACATCAA

ACCGATTCTGGTGTTCGACGGAAGGCACCTTCCGGCCAAAGGCGATGACCGAAGCCAAACGGCGCGAGTCAGGATAATTCAGAAACCCCGGGCTGCTGAATTGCTGCGGGTGGTAAGACGGAAGAGGCCAAGAGCTACCTGCGGAGGTGCGTAGACGTACCCACGAGAGATGGCTTTGCAGTTGATCCAGGAATGTCGGCGGAGGAATGT

GGACTGCGTTGTGGCGCCCTACGAAGCGGACGCCCAGTTGGCCTATCTAAATCGGAAGGGCATTGCGCAGGCTGTGATTACGGAGGACTCGGATCTGATGCTGTTCGGATGTAGCAAGGTGAGTTATAAATACGTAGAGAGGAGACTGGGGAGACTGGGCAGACTTGATTCCAGGGTATCCATCCACATGAGTCTAGCCTCCCCGAAGTG

TTAGATACGAGTTCATCTATTGTTTGGATTTTTCAAATACTGTTTCAGTATTCAGTATAATTCATGTAGAGCACAGTCGTCATGCTGTTATATTTTTGTACAACACTGGTGGGTGAAACCTCGCTTGTAGTACACAGCATCAGCGTGGTGGTTTTGACGGTGACAAACCGCGCGACGACTAAAAGGTTAATCTTGTGTTTGAATTTCTTA

ACATTTTGTCTTCAGCATTTATAGGGGAGGTGCACCATCAAAAGACCTTTGGTTACTGCAATACATCAAACGAAAGCTTTCAATCCATGCTCAGCAGGAAAAATAATAAAACTTACCAGAAACTTTGTTTTGTATTAAAAATGGGGGTGGCAAATACTGCACTATTCGCACCAGTATTTGCCACGATTTAATTCGGTTCCTGGATTCGCC

ACCTACGTTGATTTCTTATGGAGGTGGCGAATCAAAATTTCATTATTCTGAGAATTTTTGCGGTTTCAGAATGTTTCCGTATCATCTTAAAACAATTTAGCAAACACTAAAACAATTGGCAACCATATATGTCGTAAAACGGTATAAGCCGGGGTAGCGGAACAAGCTGGTACATAGCGAATACCGGTATCTTCCCTACTTGTAGGGACC

TCCCATGTTATTATTGGCAGACGGCCAACAAATGGACTGTAAATGTACGAACGGACAGGATAAGCGGGACTTCGAACGACATAATATATTATAGGAATTGGAGAGGATCTTGACGACATACGAACAAATTATTGATGAAACTTGAAATCAGAACATAACGAGTGTTATAAGGGGCCGTCCATAAATGACGTAGCATTTTTTCGCTGATTT

TACCCTCCCCCCCTCGTAGCATTTCGTCACAAATACTGGTACTCCCCTGGAAAATACGTATATCGAGCACCCCCCTCATATTTTTATTTGTTTTCCTTAGCGACTGGAGTTAAAACCAAAAATTGGAATTGAAGTTCATTACAAAGTTCGATATTATTACTGACTGAAATGAAAGGATAATCTATTCCTAACAGTTTAATATACAGTAGC

GCTCAAAAGTAATTTGGAAAAGTTTTTAAATAAACATTTCCTTGGTTTTTGGTGTCAGTTTCTTATAGACTAAATAATATTAATATTGCTGTTCTTACCATAAAAACAAGTTTACTATAAAAAATGTGAAAAAAATTGATTAGATTCAGTGAAGCTCGTCACTGTCGAGTTAGGGTCCAATATATTTTATAATATATTGAACAAAATTTC

GTTATATTAGAGGATTGTTTTTTTTAATTATTCGTTGTTAGAAATAGATTTCAATGAAGCTGATCAAGAAAATATACTAATTTGAAATGAATCATCGTCATTATTATCAATTTTAAATCATTTTAGCAAACAGTGGCCAAATACATAAAGATTACAGTTGAAATATTTGGAATGTTTAGAAAATTTTAGCGATTATTTACAAATACAAAT

ACAAATAATTAATATTTCAGTTGATTTTCATTATCGTTTCCATAATGAAATAATCTTACACAACCTATAATGAAACCGTTAAAAAATTTTCACATTACCGAGTTATTAACAAAATTTAATGATAAGGGCCATTTTATTCCAACACTCGAATCATCCCTCGTAATATTCAGGCTATTCCATCAAGAGCATTGATATATCAAAACAAATGAT

GAGTTGATTGGGTGGATATCTTTTGCAACATTGAAGGCATAATTCACGGAATAAATATCTTGCTTTAAATGTAATATTTGCCAAAGAAAAGCATATTTTATTTTCTTGAAGAAATAAAGCCCCAACAGAAAATATGGATAAAACTAAAACAGCTGACTCTATGTCATGTTAACAAATGGCAAAAAAAAAAATGTACTCAAGTAGTTAGAG

CATTGGTTTTAATCAATTCATTGAACTTATTTTGTTTATTTATAAATTTATTAATTTGAACACGAATATAATATAATATTCACATAATTATTTAAAGCCTCAAAACATCTATTTGATTGCATTTTATCAAGAAAATCTAAAGTTCCATAGATAATTTTGCAATCTTGTCATATGAGGTTTTTAGACAACTTAGGATTTATAAGTAAAAAA

AAAGAAGTTTCATAAAAAATAATTGTATGCGATTTTTTTCAGCGCCTTTCTTTTTACGGTCTTGATTCAAAATGCTACGTCCACTAGCTAGGACCCCTCCCTCCCCTCGTCACACTTCGTCACAAATTCGTGATACCCCCTCCCCCCTAAAAGCTACGTCATTTATGAACGACTCCTAATGTCCTTCCTCTCCTCTATTAAACTTTTCTA

TTTATGCTTGACTCCCTACGATCTATTATTTTCTAGATTACTGTGATTGATCCTAATTCATTTAAAACATTATTTTCGTTAAATCTAATTATATTTGATGAGCAATCGGAACATTCAAGTCAATTTTGGAATACCGTACTGGATTGCTGTACATCTTTTACTTCTGACATGATTTTTCGGTTGCAGCTAATCTTAACATGCTGTTGATTT

TTGCATTAATTTACTTGTTAACTTTTCAATAGCACTTTGTAAACAAATCTTATCCGACTTTTATTTTTTAAATTTCAATATATTTTATTATTCTCATTAAATGTAGTTTAACATAAATTTTGCAATGATATTTGCTCAGATTACACAATTTCATAGTATCAATAACATGCTTGAAGTCAACGTTTAAATTATAGAAAGTTTTTTAATTTT

CTTAGAGGAACAAATTTTGTTCCAGCTTCGTGAATCAAAGTAGTTATTCCTCAATGCAATAAATTTTGCTACAAATTAGTGAAAATATCATGCTTCTAGACACATTATTTCAGCAATTTATCATTATCATTTATGTTAATAAAGAAGTATTGTTAATGTTAATTAATTAATGTGTTAATGTTAATTAAATGACACCATGCAAGATCCATC

GATAATTTCAAAAGATCATAACCTCGCTCAAATGGTTTTTATCTATCTAAGCAGTTTTACAACGATATCGAGTTATGACAGGTTGATTGTATAACATCAAATTCTAAGTCGTATGTTGCTTTACATCCATTCTGATGCCGTTTTACTTGCCTCTGGGACTCACCAGCCCACTCGGAGGTTGGATGTTTAATATTTGAAACGTTTAGTCTG

TTTGGGAGGCCATGTGTTCTATTCGTTTGAGTCAGATGAGTCCACTGAAAGACTCCTCTCCCTTGGTTGTCGACCGCGATGGGAAAGCATAATCCATTAGCCATATGATGATGGAAACCAAGGCGTAACCTGTAGTGGTGGTATCACATTTTGGCATTTTTTTTTGCTTAAAGCCGCGTCATAATTCATATGGCATAGACGCGACTAATA

TCTCGGATGTGATCAACGGACATTCTGCGTCATTGACAGAATTGATGCCCATTTCAAATAATTAATCTATTCGAAGTGGACATTAATACTATTGGTGACGACCAGTTTGCCTTTTGCTAATCAGAATGATCCGTAGCAGCCACTCTTACTATTAGCTAACTGAACATCGTTATCATGTACGTACTTTAGTGCTTCCAAGAGCCTAAATTC

GTGATCGAAAATGTTTTGCACCAGAAAAGATGGTTGTAGAGGTATGGTGTCTTCAGAGAGGATTGTTCAGATGTTGAAGCCCTTCTTAAGAAGACCATTTTTGTAATGAATCCACATAGCAGTGAGATAGATTTTCACTTTTTTTGCAAAGTGGAGATACAACAATGGTGTCTTCGGCAAAGTTGTAGATAATTTCACTACATGAAAATT

TGCTGAAGACACTTTAGCTCTATCTATTAAAAAACAAAGAGATATGGGTCATTTTTTGTGAATGACCCCTTAAAACTAGTTTTTCGATATATCTCTTCTTTCTGGAGTATTTTTTGAGATTTTCACATGTTCTACAAAGTTGTTTTAATTGATAAAATACGCATTTCTTTACTGAGACACAAAATGTATCTCTTATATTTTCGGAGTTAT

ATGATATTTATGTTGAAAATCGACTACTGCTTCCTCTAACATTTACAGGGGCAAATTGGAGCAACATTTTTTAGCTTACATATAAAGTAACTTAATAGCTATCAAATCCATGGTAACTGACACGTGGCACTGCGTTCCTTGACTACAAAATATGTCCGTAAAATGTTGAAAAATTAAAGTATGCCAGTTTAATGCCGTGTATTTTAAATA

CGCCATCTTTCATCAGGTCTAGGGTCATTGGCATAAGTTCATCTGGCATCAAGCCGTTTGTCATAAAGACATTTGGCATAACGGTCATTTGGCATAATGGTCATTTAACATAAACACATTTATAAAGAACAGGTATATTTTGAAGAAAAAGATTTTATTCTAGTTATGCGAAATGACCGTTATGACAAATAGCCATTATTTCAATTGGGG

TTATGCCGGCAAATATATACCAAATGGCTTTTTAATAGATGAACCGTTCCCGACCTTTCATGGCTGGTGGAAAATTATCAGGTCTTCTAGAATATGATATATTCGGTTATACATAGGAGAAAGTTTCGACCGTGATTGTTTAAATTATTTTTGTTCTATGGCTGTATCGGTCATGTGACTCAAGGTAAAGGGAGTCTATAGAAGCAAATG

ATGAAACGAATAGGTGCATAGGCGTCACCAAGGGGCGACAAGATTGAAATTTAATTAGGTGGTGAGTGGGCAAGCCATTTTAAAAGGGTAGTAAGCATCAGAGCGTCTGTATTTTGGAAAGGGTTGGCTCAAAGCTTTTAATGACTTTGCTACCAACACAGGCAAAATACAGGAGGCTTCAATGTGCTCTTATACTGACGGCAATCAACC

GAACAGTTCTATCGATCGATGTCTGATATTGAGATTGACACGCAGTCAGAGAATTTGTCTCTGTAATGTGCTCTTCCGCCAGTCACCACATGATCCAGTTAGAACTGCATTTTTGCTTATTAGCCAAATTTCATATACATAGGAGAAGGTTTCGACCGTGATTATTTTAAATTTTTTTTTTTTGACTGATACAGCCATAGATCTTTACTG

ACTTTAAATGGCTTGCTCAATTTTCACCACCTAATTAAATTTCAATCTTGTCGCTCCTTGCGACGCCTATACTTCTATTCGTTTTCCATCATTTGCTTCCTATAGACTCCTTTGCCTTTGGGTCACATGATCGATACAGCCATAGAACAAAAATAATTTATATTCGGTTATGACTGTAAAAGTAATGATCTCTCAAATACTAAATATTTT

TTGAATATTGTTAACGTGAAATAGCAAGTTATCTTAAGTAAATATTGCATAAACTTTTTAAGACACAAATCTGGCGAAGTAGGCATCGAACAACAGTGATTTTATTTTAGCTTTGTGCATTAGTAGAAAGCATAAAATAGAATATAATCTTGTTCATTAAATATTTGTGCAGTTAAGTGCTTTTGAATACGGACAGAGATATGACATAAG

TCGGTCATTGAGATGACCATCTCTAGATTCCGATATGTTTCACTTTAAGCACAGTGGATAATCAATTCGCCTAAGACACATATCATCCTTATGAACACTTTTGTAATAATATTTGTTAAAAAGATAGTCTTTTATGGAGAAAGAAATTGTTGAGAGAGAGAGGATCAAAGGAAATAATCATAGGCAAACATATTGAGATGTGGATAGTCG

ACTGTATTAACGCACAGAAATATATAAAGGTCTTATTTTAAATTTTCTTAAAAATACAAATAACAGCGAAATTATAGCTAGTTCATGGTAACAAATGATAGTCCGTTCGAAAGCCATTCAGTTCGTATGTCACGCCACTTTAGTTTTATTTATTAAGATGGATAGATTTATAAGATTATGTTCTTGTAAATGCAAAAGAGCAATACCAAA

ATAAAGAAAACATTATTATATTCTTTATTATTGTTATTATTATTATCATTTATATTATTATTATTTTAAGCTTAAATAGGATGACTTCCGTTTTAAGGCAAATAAATTACTAACATTATTCTATGTGCTTCATACTACGTTCAGTATAGGCTGATGTTACACGTAATAAAGGCTATTTCTTAAATTCATACATCTAGTTTTCACTGAAAT

GAAATTATGACGTTTGGTATCAAGATATCCTATCTTGAGCAATATCAGCATAGCCAAACCGAATATATCATTGCGTATTAAAAGACCTCAAAATTGTCCTCAGCCCTGAAAATGGCGTATTTAAAAATAAGCAGTATTAAAAGCTGCACAATTTTAATTTTTCAATATTTTACAGAACATATTTTTGTAGGCAAGGAACGCAGTGCCACG

TGTCAGTTACCATGGTTTGATAGCTATTGAGTTACTCTTTTATATGCAAGCTAAAAATGTTTAATTTGCCCCTGTAAATGTTATAGGAAGGTGAAATAGTCGATTTTTCGACATAAAATATTATATAACTCGAAAACATAAGAGATACATTTTTTTATGTTTCTAAAAACGTATTTATCGAATTAAAACAACTTTGTAGAACATGTGAAA

AATCTCAAAATACCCAGAAAGAGATATATCGAAAAAACTAGTTTAAAGGGTCGTTCACAAAAATTACCCATATCTCTTAGAATTCAATAGATAGAGCTAAAGTGTCTTCAACAAATTTTATGTAGTGAAATTATCTACAACTTTGCCAGAAGACACCATTATTCTATCTCCTCTTTTGCAAAAAGTGAAAATCTATCTCACTGCTAGGTG

GATTCACCACAAAATAATTCTTTCTTAAGAAGGGCTTCAACATCGAACAACTTCTCTGAAGACACCATACAATAACTTTTCTAGATTTGCAAAAGCAATTTTCGATCACGAATTTAGGCCCTTGGAGCACTGTGCGACGTAAGGGAATATTACTCTGAGAAAATGACTAGTAGATTCACTGAGTAGAAATAAGTGGGGACAATCTTATTT

TAATATGGTTTTACATTGCCATAAAAAGACAAACCTTCCTGGCCACGATCACAGGGTTTGACGGTGCCAAAGTAGAAGGTGCACCATAATATCCCGTCTGTGAAACATATCACCTTCCCAATACTTTGGCACACCTCTAACGGTTCATGATTTCATGATCAAGAAACGACTGCATTCAGGCGGAGGATCTGTTAATATGTTTCGGCATAT

TATCGGGTCCTCTTCGAACGGAGAGGACCGCCAGGGCTCATTGGTTACTTATAAACCAGTGCTGGTTGTTGATGGTTCAGTTCGTTTACCACGAGCTGGCATCCCTTTGTACTAATCAGTAATGGTTGTTAAGTTTCGAAACTAACGATGTGATCAATCGTTTATAATGCAACATAAGAATGGGTGTTTGGTGGAACTGCTTGTAACACT

TGTTATTAAATTTAGTGATTGTGTTGGTGCTAATGGTGTGTGTATAATACAAAATATATGGTGATGAAAATTTAAAATGAAAATTGGAAGTGATTCAGTAGTTTTTGTTAGAAATATAATAATGATGATGTGTATAATGATGTGAACTCTACTAATTTAATAAAGAACCATAATACATTTGCAATCATTTCTCTAAATATAAATATTATT

GCTAGTTCTTCAAACATGCATGATAAAAGTGTTAATAATGACTTCGAGTGCAGAAAAATGGATCAGAAGTGATTTTGTTTACTACTGTAACTTTGTTGACAGTGCTAAATTGTAATTTGAATAGTAATGACTCTACAGCAACAAAATAATGAAACATTTTAATTGAATATCTTTTAATTACCCAAGTAATGCTAACAGATGGTAAATGCT

AATAACAATGAATTTATAATAGAAGAAATGGTGCTTGATGTGAAAAGTGATATAATGGAAAGTATATCTGAGTGTATTACGAAGTTGATGAGTGATAATCAATTGAAGCCGTGATAGTGTCGAAAATAAAAGTGACGATAGTAGGTGATGAAATGAAATAGGGGAATGAGGACCTTCTAATAAAACTATTTCGTTCTTCACTTTAACCAC

TCTAGTAGCATTTCAGGCACATCATAGTTTGATAGTAGTCTAAAGTACTAGACTGCAAAACTACGGTAAGGGCTGATTGCTGCTAGGGTACGCAGTGACCATTTGAGCAACATTGCTTAGGAACGTCTGTGTGATGAACGCAATAGAGGCTAAAGCAAATGCTGGGAAGGAGCTTAGGGCAGATTAAGAGTGCCAGTACTACCCCTATCG

CTACAAACAAAAAAAATACATTCCATAAAAGACAAACCTCTTTTGTAACAACGGTATAAATAATCTAATTCCAGCTTCATTTTCGCAAAATTTACAAGTAGAAAGTAGGCGTTGTGAAGCATTGCGAAAAGTGAATCTTGTAGAGAGACTGTTATAGAAGCCTATAGGTAACTTTACCCTTAAATATTCACTATAAAAATGACTATGGAA

AGTAAGACGCTGAGATAGATCATTAGGGAGTACGCCTGGATTTCGAGAATTGAACAGACAAGGAAGAAGTGACTTTTTTCTTTAAGGATATACGAACATGTGACCAATACTCGCGTTCAGTATCATAAGGGCGTAACTGCAATGATCGTTTCTTCATCGATTTTCTCTTTAGCTAATTACTTCGACGGGTGACAGATTTTCGGCTGCTAT

TTTCGATTCAGTAGAAAGAACTCATCGTACTTAATCGTGCTGCGTCGTAAAATATTTCGAAAATTGATTAATTTTCTTGCTTATCGAAAGAGAAAATCGACGAAGAGAAATCGATCACTGCAGGCTTACGCCTGTATGATACTCAACGCGAATAAACTACTTTAATAACAAAAAATGAAATTAACTTGAACTACTACTAAATAGCCTAAT

TTTATTTAAGACTTGACGACAATTGACATTTAAGGACCTAATCTTACGTAAAGACATGAGTAATCAGAATAGCACTTAAGAATTACAATTTATTCAAAACCATTTCGACTAGACAGTATTAGTAATGACACTACTACAATACTATTTCAAACAAATTCTGATGGAGCTGCTGATTTTCACAATGCTTCCCTTTCTCAGAAACAAGGAATA

TGGGTATTTTTCAAAAAACTATCACGTCACAATACTAATAAGGCAAGGCTTCGTGGCCTTGCGGTTAGCAACGTCCAGTCGTCTAGACGTATGTTGCCATGTAGTGGGTTCGATTCCACTTCAGTCGGTGAAAACTTTTCGTCAAACGAAAATTCATCACTGGGCTACTGGGAGTGTTTCAAAGTGTCTTTTAAACAGTGTTCATGTGGG

CACCATAGCCTAGTCCGTCTGATATTGGTCTGGTAGAGGAAACTTGGCAAAAGCCAACCGAGGTTCAGCCTTGACAAGTTAAGCTGTCAGGCAGCTCCAACTGAATAACATACAAAAAAAATCTGGGCGAGATATTTATATAATTTCTGAAGGTAGTAAATTCCGACAACAGAGTAAATTTCAGACATATCTCTGCTAATTAAGGGAGGG

AGAGGATAGGCTCAAACGTAACAACTCATACAATATAGGGTAACCAATATATTTGGACCTCTATATATTTTGGACCCCCTGGGCATTTTCCTAGAAATTTCCCAGTTTAAATCGAAAGACCAACTCAATTCATGCCATTTGGAAGGATTGGACTATTCCGCACTTGCATCAACCGTTTTAACATAGAAAATGTGTTTATTTTTATCTGAA

ATTAATTTAATTTTATCCTTCATCCATGCAACCGTTACAACACGTTACACACATAAATTGGAAGCCGTCAGAATTTTCAAATGTAAACAAAAACTTTCCAAGTTTCTGAACTAAAAGCGTAGAGTTCCTTCGGGTTTCCATTGTCAATCGATTGGTTAGACTATGGCAAGCATATTATACAACCGGTGCCGTGAGCTTTGTCGCCAAATG

ATAAAAAATGCGGGGTCCAAAATATATACTTTGAGGTCCAAAAATATATTGAATTGTCCAAAATAATAAAAACACTGCTTCACAGATTCAATTATTATTTTCGATGTTTTCTTGGATCCTACATGACCGTTTGGGCATTTTATCTCGTAAGAGTTTTCTCCACATTTTAGCATGTTATTTGACTTAGTTACGCTGTGCAATTAATTAATT

GGGACAAAAACTAAAATCACATCATAGGGGGTCCCAAAAATACGACCGTTACCCTAATATTTCTCATAAATAATCACACATTCAACAACTAAAATTATTAAAACAAAAATAATCTTACAGGTGCTGTTCAAGAAACAAGATCTGACCAGGACCGGGTTGATGATCGAAGCGGAGAAGTTATTACCTGGCCATGGAGAACATGGAGAAAAG

TACACATTCGATAAGTTTCGTTACATGTGCATCCTTTCGAATTATGTGACTATCTGGAATCGCTCCCAGGAATCGGATTGGCCAAGGCAAAGTTTGTCTTGACGACGGAAGATACGGACATCCGCAGGGCTCTGTCGAAAATTCTGCCTATCTGAACATGAGACAGCTGGAAGTGTCCGAACAGTACAAGGAGGAATTCATGAAAGCGGA

TGCTACGCTGAAGCACATGGTAGTTTTCGATCCAGTAGGGCAGAATAAGCGCGACTCAACGAACCTTCAGATATGGGAACCGATCCAGATCTCTGTTGTAATTCTGGGAAACTTCCTAGACGACAAACAGCTTTGGAGTTGGCTCTGGGGAACCTGAATCCCTTTTCCATGGAAGCGATTGGACAACTGGCATCCGGACAGTCCGGACTG

CCAGCGACGAAGGTCAAACCTCCAACTGGAATCGAACAGCCGTGACTAAGCATCCCAGCATTTGGAGAGAGGTTCCTATGAGGCGTAGATTAAGAAGCACACCTCGAAATCCACTGAGGGAATAAGGTGCGGAATAATGTTCGATCTGGAGAGAAGATCGATATCTAATTCGAAAGTTGAAGAAGAAGGGACAAAGGGAGTCAATTGACG

ACGTACTGCAGTCATGCGGAATTCTACCGAGGGATGAACAGCCTCCCACCGAAACGGCTTTGTCTAACTACAATGACCTTAGCAAGAAAGACCAATCGACGTGGATGGGATACGGGAAAATAAAGAGTCACTCCCAGGAGGAATCCTTTTCATGTAAGTAACATTGCAAAAAATGTTCCGGGTCTATTAACCTTCCTTTGTCGGTGTTCT

GTGGTCCAATCTGCATATGTGCACGGCGACGTTCCTCATTTCATAATTGTCCCATGTTTTAAGAAATTTCGTGAAGTTATCGATTTGAATGTATGGTCAATCTGACCCCATGACACGAGCAATGTAACTTTTTGGCACAGACAGAAAGTTAGATTTATTCCTTTTCGGGTTAAATCCATAGCCCCGGTCCCAATCAGGAACAATCACCGG

ACTCGAACACGCTGAAATCTCCCATAAAAACTCACCTCGGAAATAGCTCCCTCCTGCGCCAGATAAGCCCGGTGAAGAAGATCAGTACGATTCAGACCCACCATGCAAGTGCGATCCAAGACGGAAAAGCAGTCGCTGACGGACCGTAATGGCGACGGCCAGAAGGTCATCAGTCGATTCTTTTCGGTTTCATCGTCCTCGCAAGTGACG

ACTACCGTTAGTAATCCTCCGGAATCGTCTTCCGCCAATGAAAGCAGGCCGCAGTTCAGGATCCGTACCAGGAGAGCAAGAGTCCCACTCAAACGGCAAACATATACCTACTCGCCGGAAGCCAGAGGAAAAGACTCCCAAAAGCAGCGACCTTCGCTGCATAGTGAAGATAACTGTTCTCCAGCTCCTCCCGGTGAAGGGAGAACGTAG

TAGAGAAACTGGACAAAGTGATTCGTGGAGGAACCCGAGGAAGTTAAGCTCATCACAGAAGGAAAACCAATCGGAGCAAGATGAGAGTCCACATCTCGATTGTCCTTGTTCAGCAAAAAGGAACCGATGCGATTGAGTGCTGAGAAAGTTGATATTACTGAAGAGAGGACGTTATAGAATTAAAGATGACGAAGAACAGGATTCCGGTCA

AGATTTGTCTGTTGAATCCGAGTAGCGACTATTATTGGACGAACTAGCAGCGGAAGTAGTCAGAAGAAAGTCACCTGTCGACGGGTAGGACTCTCCAGAACAAAAATCTGAGGGCGTTGGACCAAACCAGAGCAAGCTTTCGATGTTCGGGTTCCAGAAGAAGTAAGAGTACCCATATCTCTTATAGTTCTAGTACACAGATAACCTAAC

ATTATTTAAAATCTGTTACAGAGTGCAACTGGAAATGATGTGACTGAAACGTGACAATCACTGCCCGTGAATACTGCATTTATGAATTACGATTTAGCTTTAAGTAGAACCACTCGTATCCCAACCCAGGCTTGCGTTTGGATAGTTGAAAATTTGTGTTAGTAAACTCGTTATTGTTTCAGTTTTTTATATGCGCTGATTTATATGTAA

CGTTGAACTATGAACTACGAAAATCCCTTTTCTTCGTTTTCACTTTGTTCACCAAGATGTTTCTCCGCTCGTTCTATTGGTTTTCTTCGCTCATATTTGGGGGCACAATCATCACCAGACCGAAAGCTGCAACAACCGAAATGATCGCCACGAATTCCTTCTCCAAACCGGAAACACCTAAAAGTTAAAAATAATCAAGGCCTTCTATAA

TGAAGGACACGGTAGCCACCTGTCCAAGTAAATTATTCGTCTCAAAACCCGCTACCGGGTCTTAATGGATCATTAAAATAACCAAAACTGCCGCTATACAAAAGCATAGATAGCGCCACCGTAGCCTTGTGTGTTTGACAAGAACAGCAATAATGTCACAATGTTAATTCCCATATACAGTAGCGCCTTTGTTTTGTTGCGGTGAGCACT

GACAAAAACTACCTTCAATGATCATTTTACAGCATCCAATTTCGTTGCTTAACGATCAACCAAAACAAATCGTCCTGAGTATACAAGTCATCCGTCGCAACCAGCTGTCATACAAAGAGGAGAATGAAAGGAATATCGATCGTCGAAGGGTTTGTAGAAGCCTTTGTTTCACAGGGGCCAATATGTTCGTGTATGCTCTGCTTGACACGA

AAAGGAGGATTGCACGCCGCCTATACGTCAAGCGCAATTAACTTTGGAGATATCCCCAGTTCGTCGACAGCGATACAAAAGCCCAAAGGGGATTTAAGTTTATCTTAGCGTTCAACCATCTTCGGGACACCCTTCTATAAAAGGAGAACGAGCAACGAATATCGTTCTCTTTTTTTGCCATAAACACCATCTCGGCACCATCACGAGAAG

TGGATGGATCGTAGGACACTCTGAAAATTCCTTCGAAGTGAATTTGAATTATCCAAGTATTAATTGAACTAGTGTAAAATAAATTTAGTTGAATTAGCGAAGTGAAATAACGAGAATTTTAATAAATGTTTTGTGTTTAACGTCAGAGTGGCTAAAGTGAACTCTTGTGCCTGAATCAAGCTCTACAATTTTCACAAATTCTACTCAGCT

GAACTAGTTCCGTTGCTACTCATGTGCTCTCGACATATGGAGATATATAGAGACGAATGACCGCCAGGTTAAAGTCCCCATTCACACAACTGTTGAGGGAAATACAAAACAGAGATATTTAATTTATTTGGTTAACATCTAAACAGATAACACTGAATCAACAATTTCACGCCACAATGCACCCGATTCTGTTTTTGCACGGGGATGCGG

CGTACCGTGCAAAAAAGTTTCAGTTCAAAATTTCCAAAAACCGTGTAAAAAAGTAACACCAGTTCTCGAAGTTTCATGCAAAAAATAAGGTTTTGGCGGAAAAAAATGTATGGAAACTTTTTGCACGGCCGTGTAAAAAAATCCGTGCAAAAACAGAATCGGGTGTACTCGGTTCGTGAATCGCATCACTTCATCCTCAGTTTGCACGCT

CGCCAAATCGATGCGCACTTGATCCGCCCGTCTGGCTCGCTGCGCTCCACGCCTTCTTGTACCAGCGGGTCCGAAGCGAACACAATCTTTGCAGGGTTGCTGTCCGGCATTCTTGCAACATGCCCTGCCCATCGTATCCTTCCAGCTTTGGCCAACTTCTGGATACTGGGCTCGCCGTAGAGTTGAGCGAGCTCGTGGTTCATTCTTCGC

CGCCACACAACGTTTTCTGCACACCGCCAGATCGTCCTTAGCACTCGGCGTTCAACGGGAGCTCCAAGTGCTTGCAGATGCACTCTCGATTTTCGTTCACGTCTCATGCCCGTAGAGTACTATCGGCCTTATGAGCGTTTGTACATGGTACATATGAAGCGGGTGTGAATCTTTTTGACCGTAGTTTCTTGTAGAGCCACTGATGATGCG

CCTCCGTATTTCACGGCTAACGTTATTGTCGGCCGTCAGCGAGAATCAGGTAGACGAACTCGTCGACTACCTCGAATATATCCCCGTCTATCGTAACACTGCTACCTAGGTGAGCCGCAATAGCGTTCGGCCTCACCAGCCAACATGTACTGTCTTGGCCGCATTCACCACCAATCAACCTTTGCTGCCTCGCGTTTCAGGCGGGTGTAG

GTCTGCCACCTTTTCAAATGTTTTTATGAAACAATGTCCATATCATCCGCAAGCAAACAAATTAACTAGATCTCGTAAAATCGTTCCCCGGCTGTTAAGCCCAGCTCTCCGCATAACACAATCTAGCGCAATATTGAACAATAGGCACGAAAGTCCATCACCTTGTCGTTGTCCCTGGCAGGATCCAAACAAACTGGAATGTTCATACAA

ACCTTCCGCACACAATTTTACACACCTTCCATCGTCGCTCTTATCAGTCTCGTGAGATTCCGGGAAGCTGTTCTCGTCCATGATTTTCCATAGCTTTACCGCGGTCGATACTATCGTATGCCGCCTTGAAATCAATGAAAAGGTGATGCGTTGGTCTAATTGTTCACCACATTTTTGGAGATTTGCCGTACACTAAGATCTGGTCCGTTG

TCGACCGGCCGTCCTTGAAAACCAGCTTGATAACTTCTCACGAACTCGTTTACTACTGGTGACATAGGACAGAAGATGATCTGGGATAATACTTTGCAGGCCGCATTTAGAATGGTGATCGGTCAAAGTTCTCACAATCTAACTTGTCGCCTTTCTTGTAGATGATGCATATAACCCTTCCTCATCCGGTAGCTGTTCTGTTTCCCCAGA

TTGTGCCTATCAGCCGGTGCAGACAAATGGCCAGCCTCTCCGGGCCCATCTTTATGAGTTCAGCGCCGATACCATCCTTATCAGCAGCTTTGTTGTTCTTGAGCTGGTGAATGGCGTCCTTAACATCCCTCAAAATTAGGGGCTGATTGGTTTCCATCGTCCGCAGTACTGACGAAGGCAGCATTTCCTCCGTTGTCCCGTTCTTCATTA

CCTGTGCTCTCAGCGCCATTCAGGTGTTCGTCAGAGTGCTGCTTCCCCACCTTGTGATCCGCCTCACGCTCGTCCGTCAAAATGCTCCCATTCTTGCATCCCTGCATACCTCGGCTCGCGGCACGAAGCCGTTGCGGGATGCGTTGAGCTTCTGATAGAACTTACGTGTTCCTTGAGACTGGCACAGCTGTTTCATCTCCTCACTCCGTC

TCCTCCAGGCGGCTTTTTTCTCCCGAAAGAGGCGGGTCTGCTGTTGCCGTTTCCGTCTATAACGTTCCTCGTTCTGCCGGGTCCCTTGCTGCAACATGACCGCCGCGCATTCTTCTCCTTCAAAAACCTCCTGGCACTCCTCGTCGAACCAATCGTTCCGTCGACTCCGTCCCACATGCCGACGGGCTCTCAGCTGCATTGGTGGTGGCT

GCTTTCACTGTTCTCCAGTAGTCCTCATCCAACTCACACTCTTCCGGTAAAAGCAGCCTCAGGTGCTGTGCGTACGCAGTTGTGACATCCGGTTGCTTAAGCCGCTTTTAGGTCATGCCACGGCGGCCGTTGGTACCGTACGTTGTTGACGACGGAGAGTTTTTGGCGCAGTTTAACCATCACCAGATGAACCGCCACGATAGGTCCTGA

CGTCGATAATGTCGGGTACCGTCCATCAATCAGAACGTGGTCGATTTGTGATTCTGTCTGCGGTGATCTCAGGGTATCGGTATGGAGGGCTGTGCTGGAGGTAGGTGCTACGGTATATTCTTGGAGGCAGCCATCAATTATTCGTAAGCCGTTTTTCGTTCGTCAGCCGGTGGGCGCTGAATTTTCCAACAGTCAGTCTGAACTCCTCCT

CTTGTGATCTGAGCGTTTAGATCTCCTATGATGATATTTTGACGCGTCGTGGCTTGGGCGGCTGTCGTACTCGCGTTCGAGCTGAGCGTAAAAAGCGACTTTGACATCATCAATTGCTTCCTGAATGATGACTGTGCACGTTTATTATGCTGAGTTGTAGAACCGGCCTTTGAGAACGCCTCAACTTGCACATTCGATCAATCACCCTCA

TTCTTGTTTACGGCGTATCCAACTTTCGATCGAATCCTATTCGTTCGAATCCGGCCACTTGATTTTGCTGGCGTAAACGAATGCAAACGGTTTTCGATCGAAAGAGCATTCGAACGTAAACAAGAATAGAAATTGAATGATCGGCCACCACCCGCGTGATCGCCTCTGCATATCACCTATCACTATAAAAACTGTTCCCAGATCGATTGT

GCTACTTTTCCCCGAATGTCGGTTCCCCGAACGCCAGTTCCA
