## Supplementary figures and images for "Marker-assisted mapping enables effective forward genetic analysis in the arboviral vector *Aedes aegypti*, a species with vast recombination deserts"

### G0_Black-eye_male_left_30x.jpg

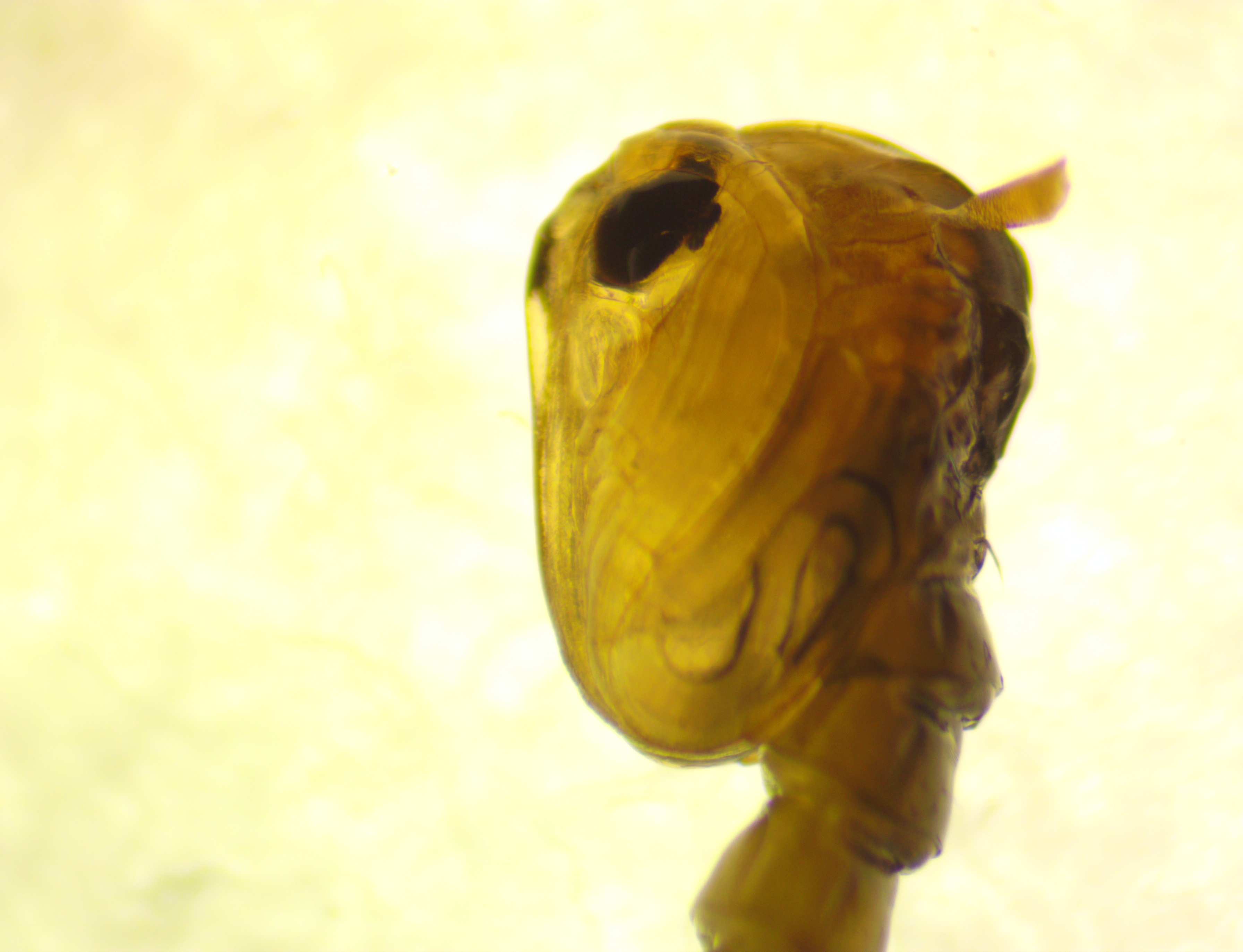

### G0_Black-eye_male_right_30x.jpg

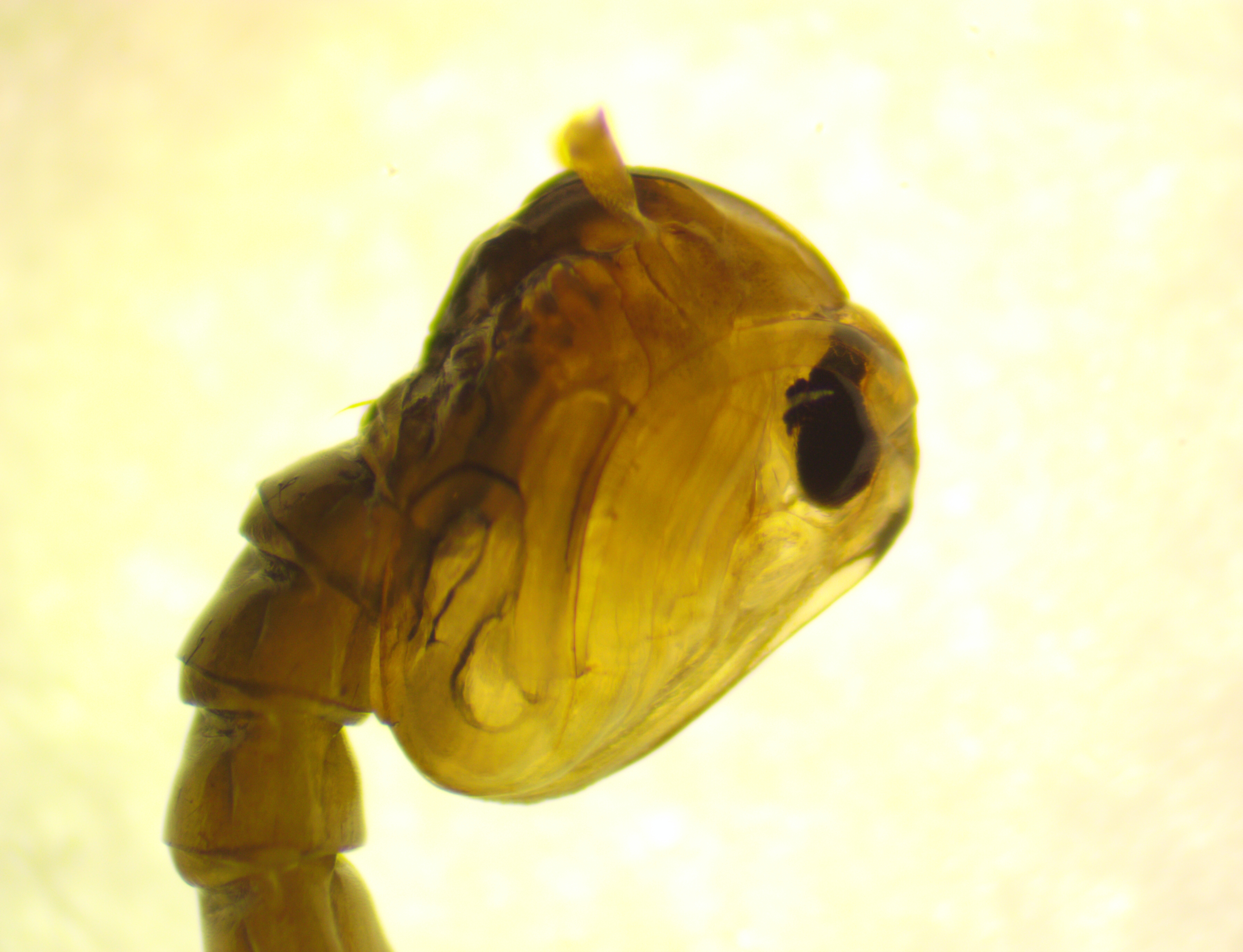

### G0_mosaic_female_left_24x.jpg

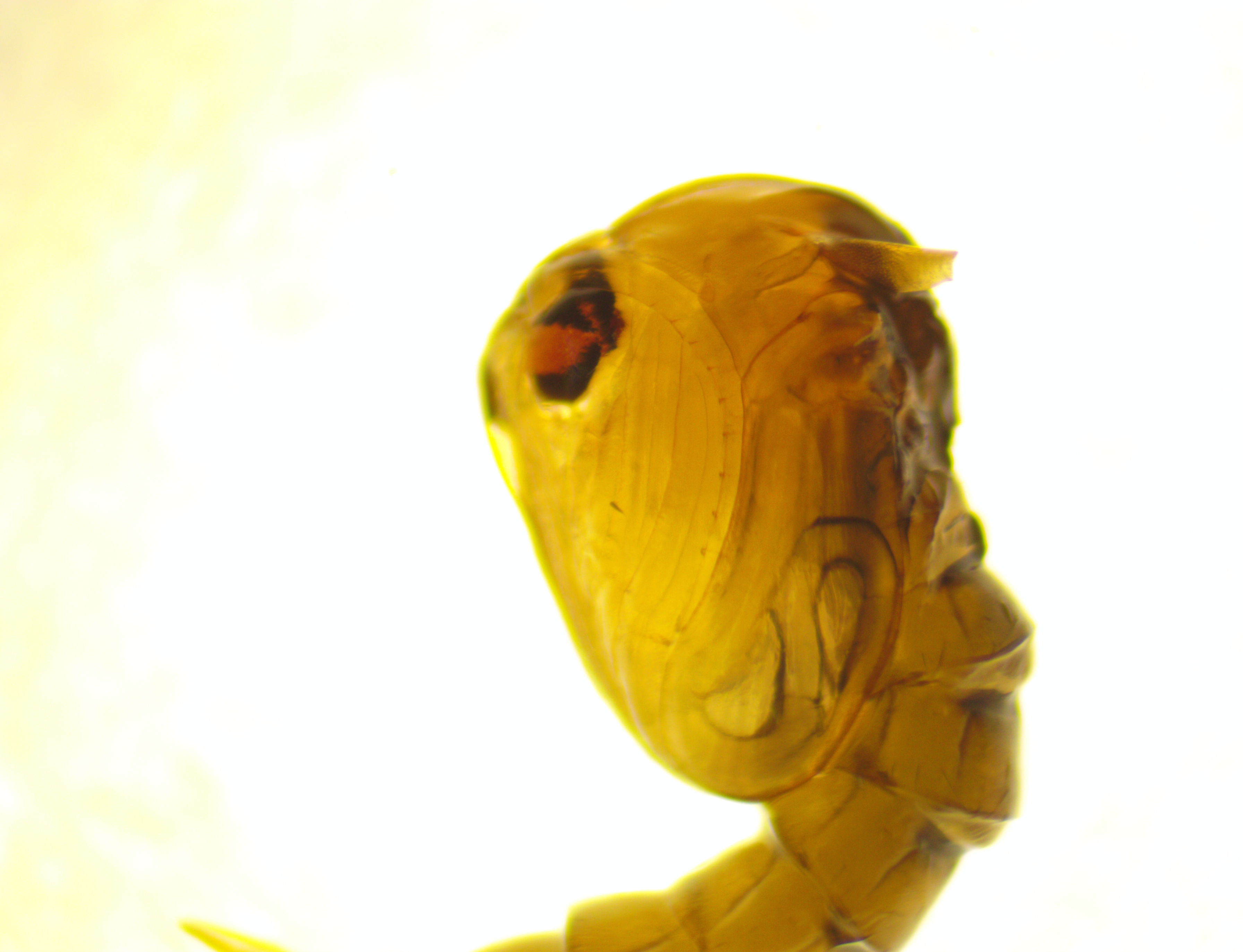

### G0_mosaic_female_right_24x.jpg

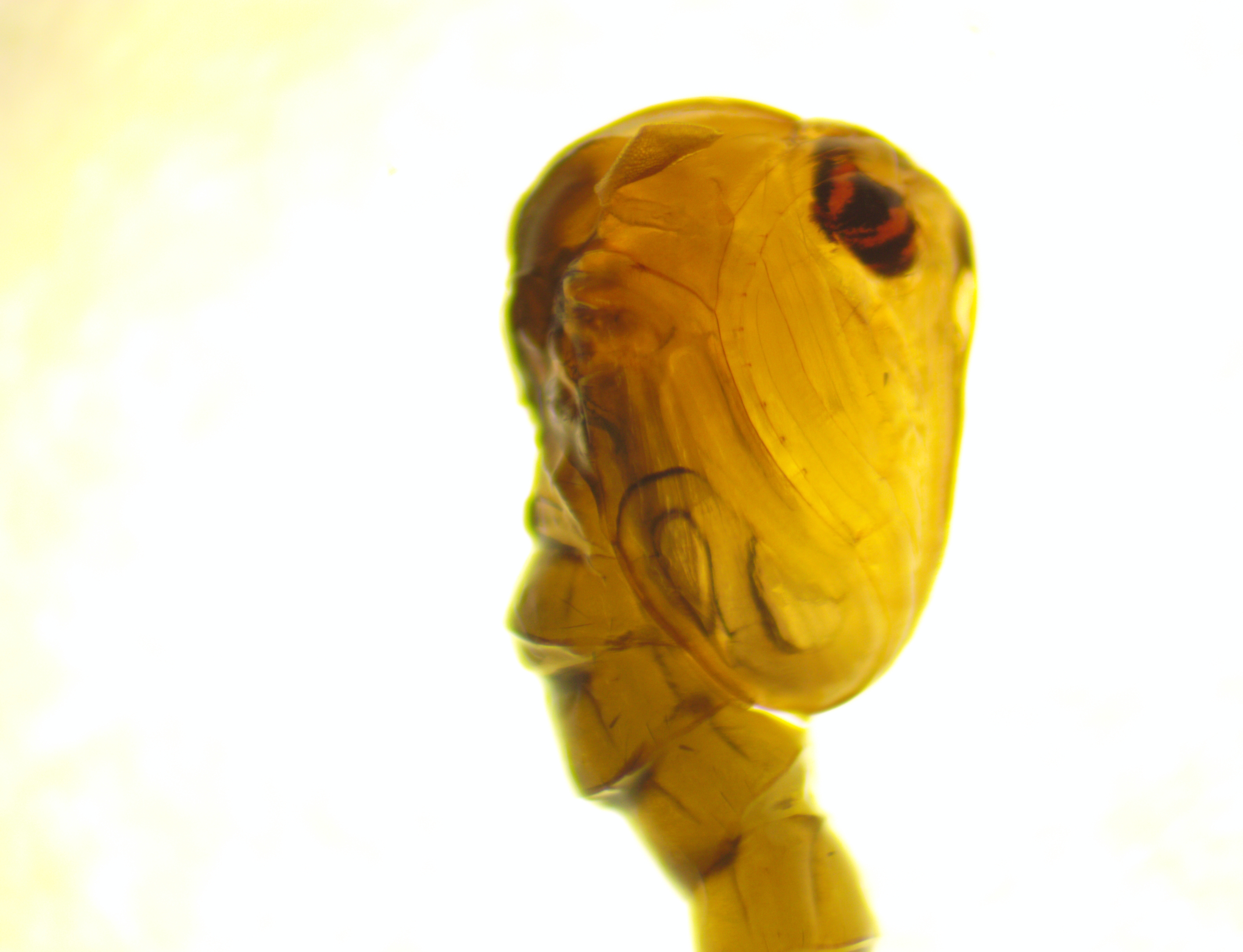

### G0_mosaic_male_left_30x.jpg

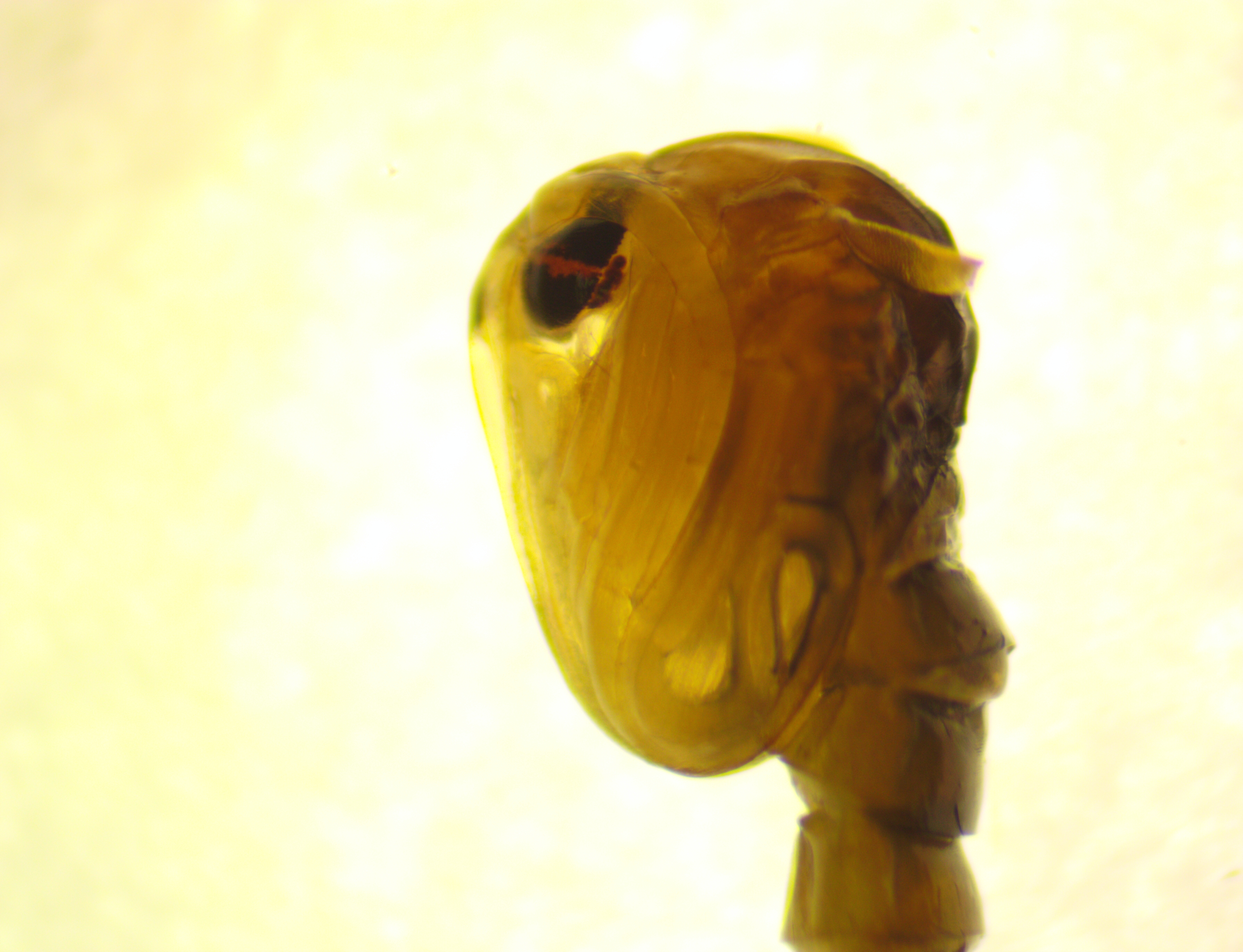

### G0_mosaic_male_right_30x.jpg

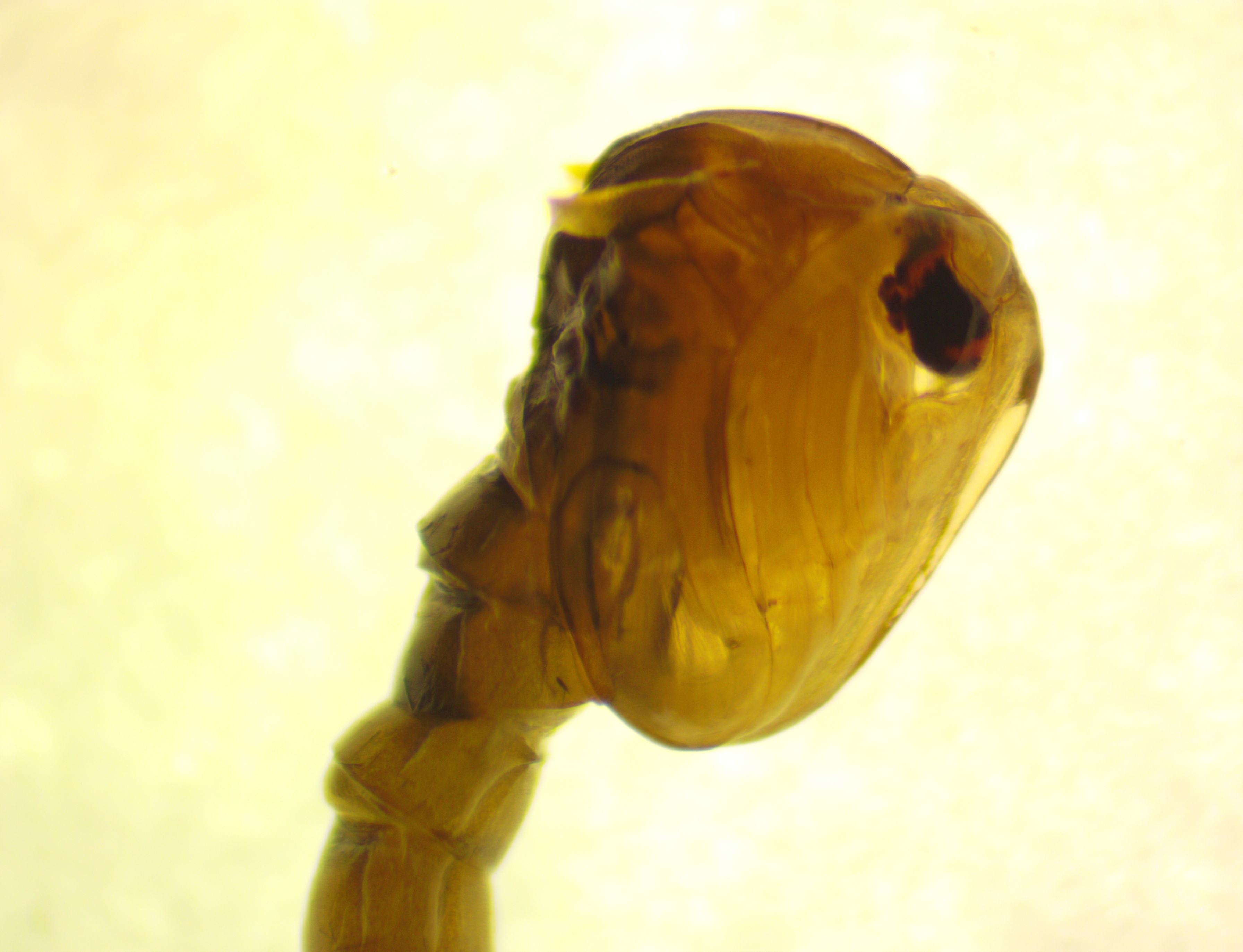

### G1_black-eyed_and_red-eyed_male.jpg

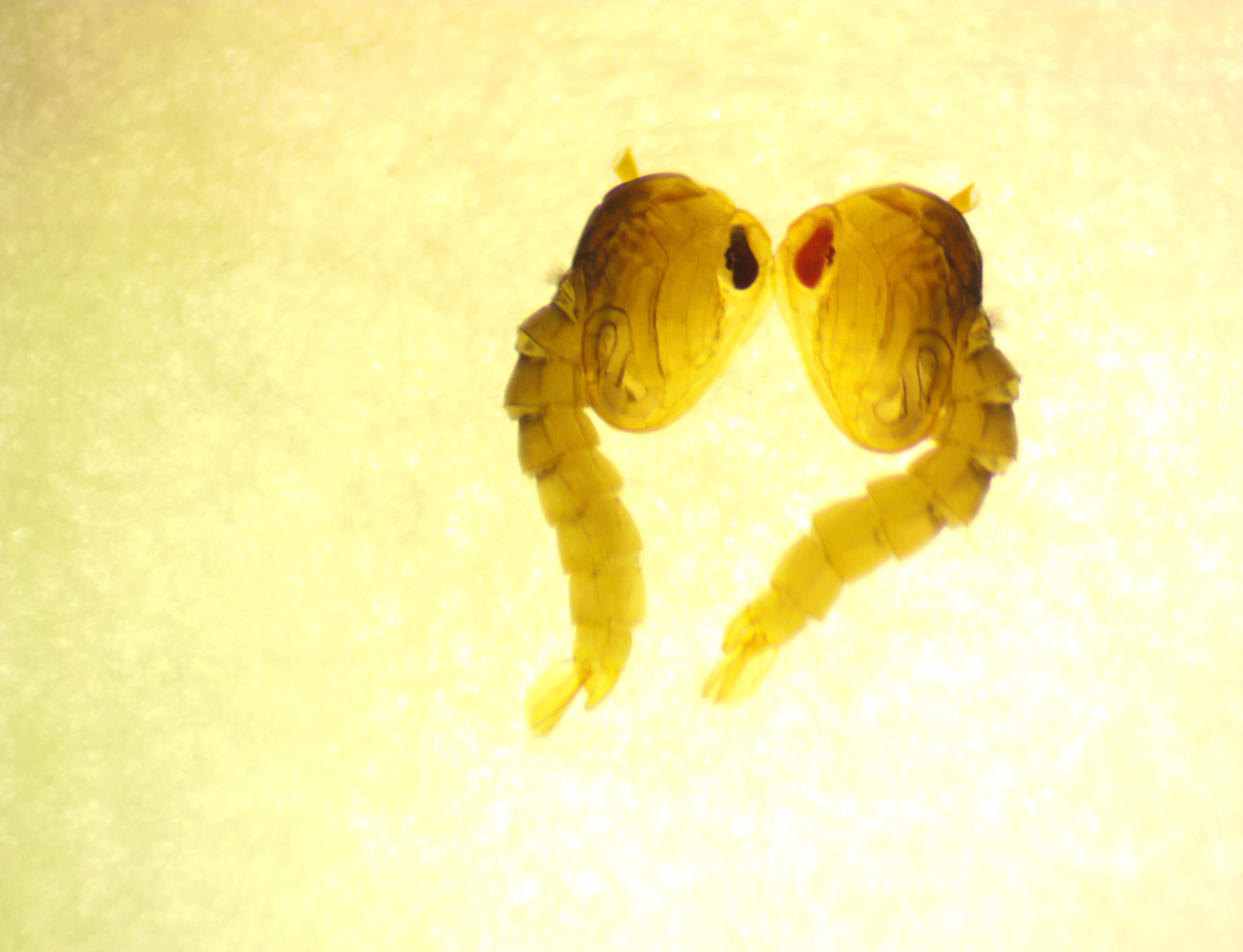

### G3_backcross_black-eye_female_adult_21x.jpg

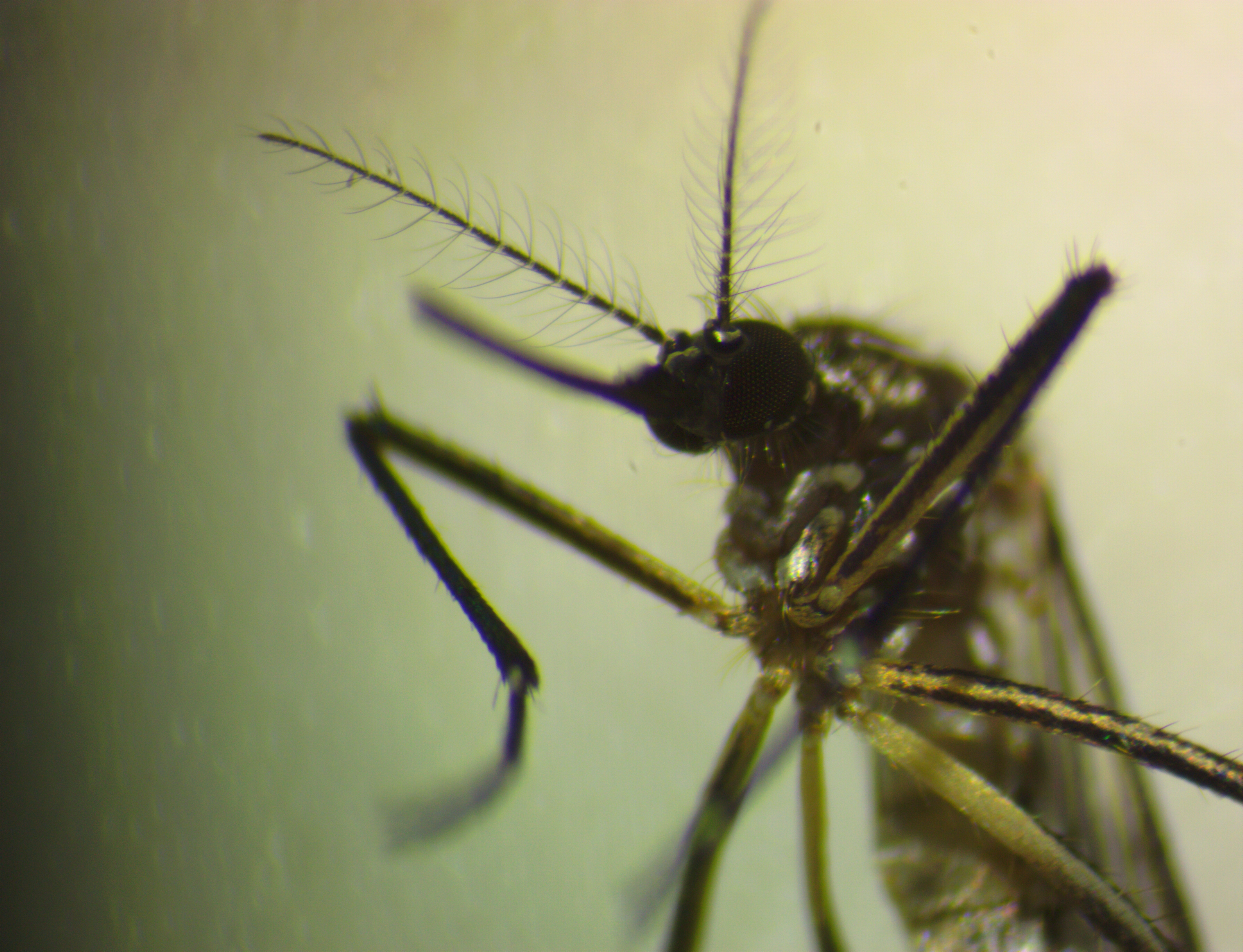

### G3_Backcross_pre-pupal_larva_3_phenotypes_29x.jpg

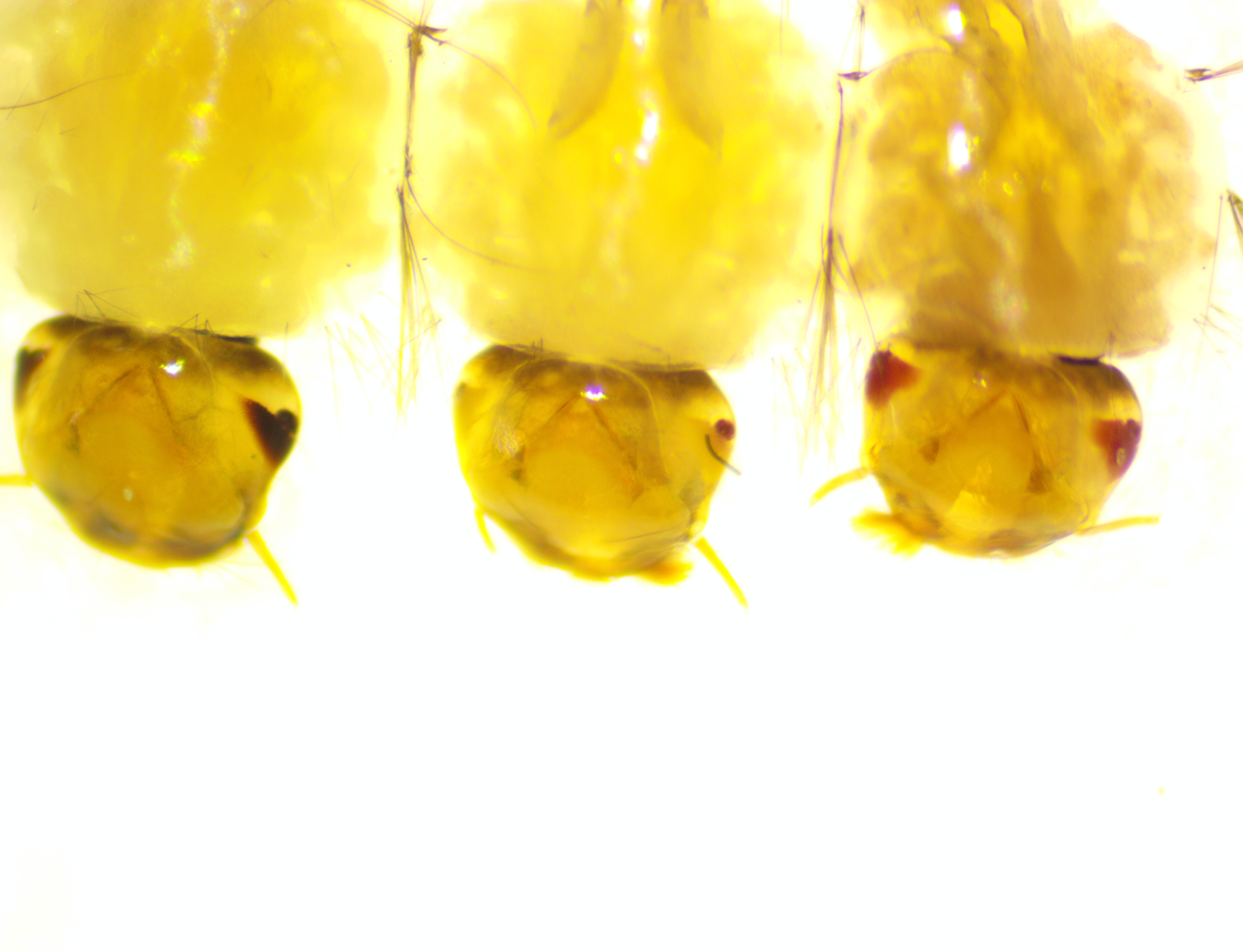

### G3_backcross_pupae_4_phenotypes_15x.jpg

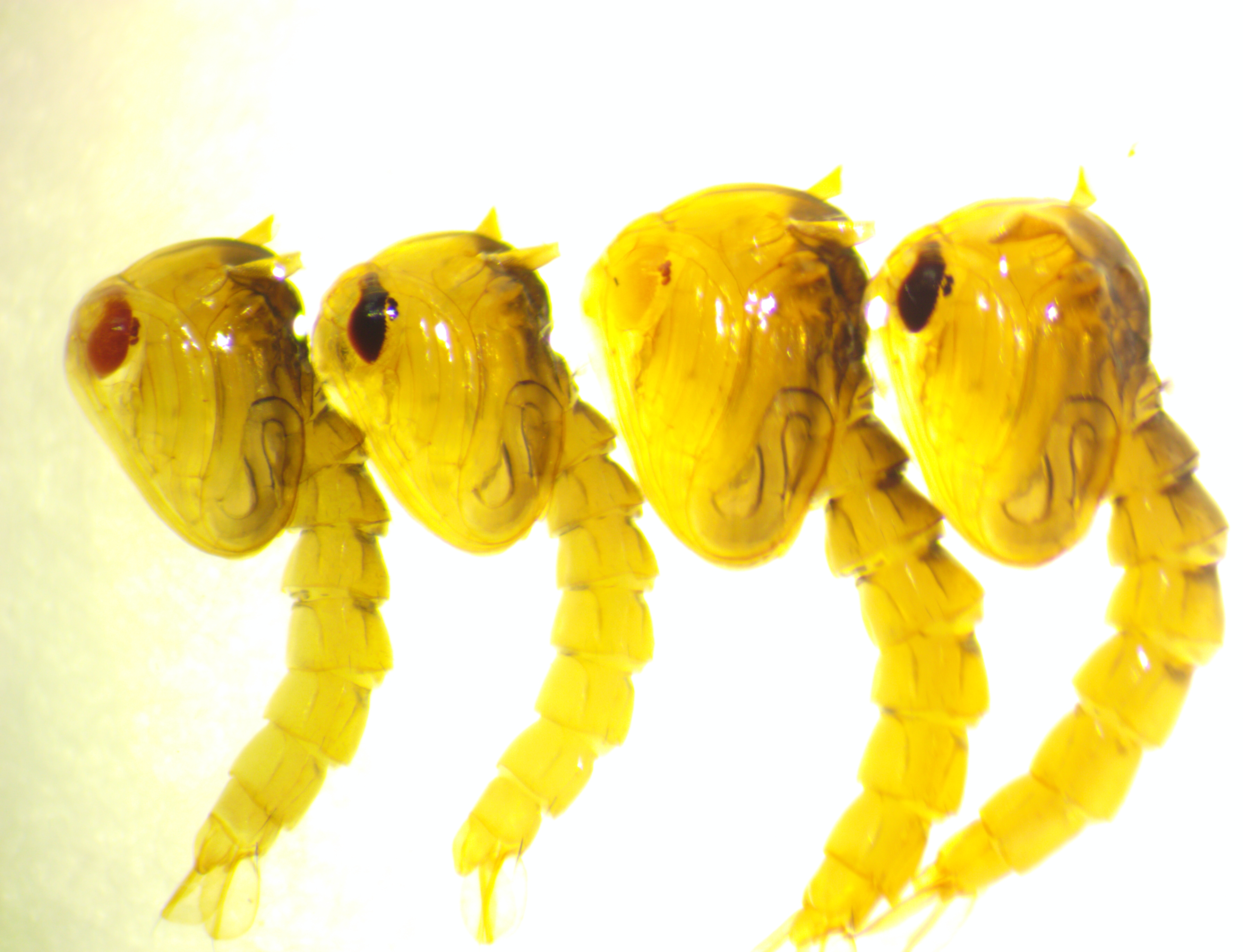

### G3_backcross_red-eye_male_adult_20x.jpg

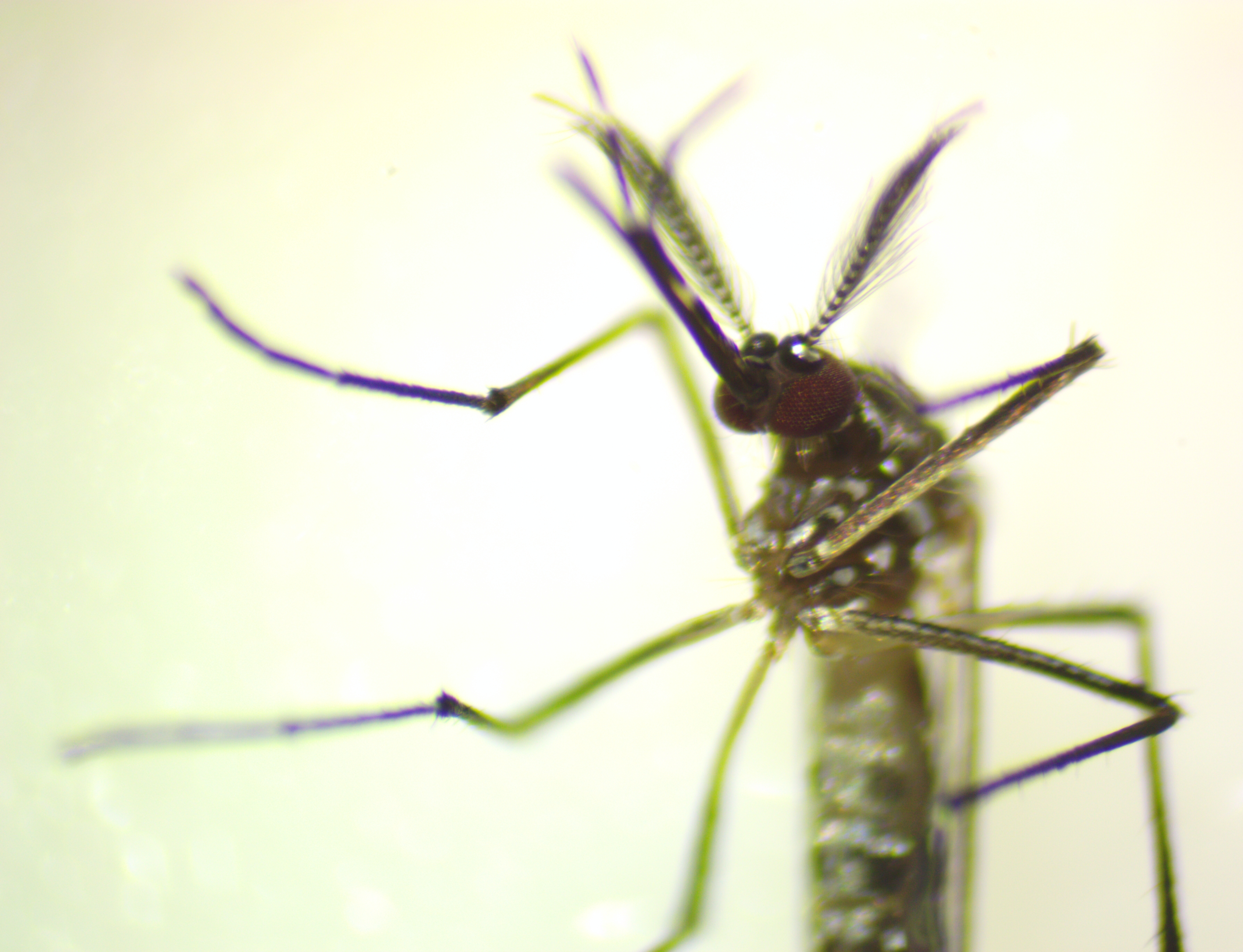

### G3_backcross_Yellow-eye_1_day_pupa_40x.jpg

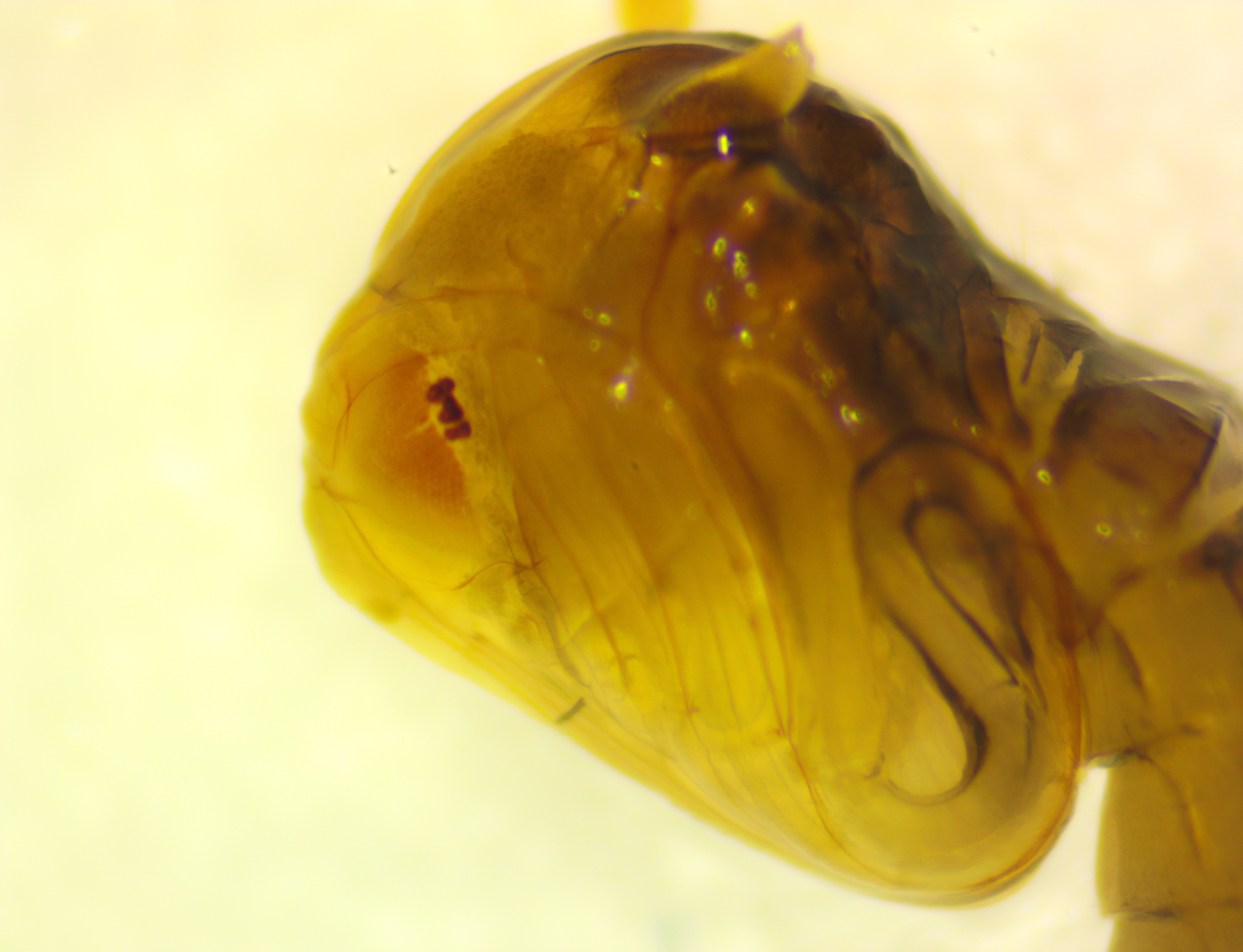

### G3_backcross_Yellow-eye_2_day_pupa_40x.jpg

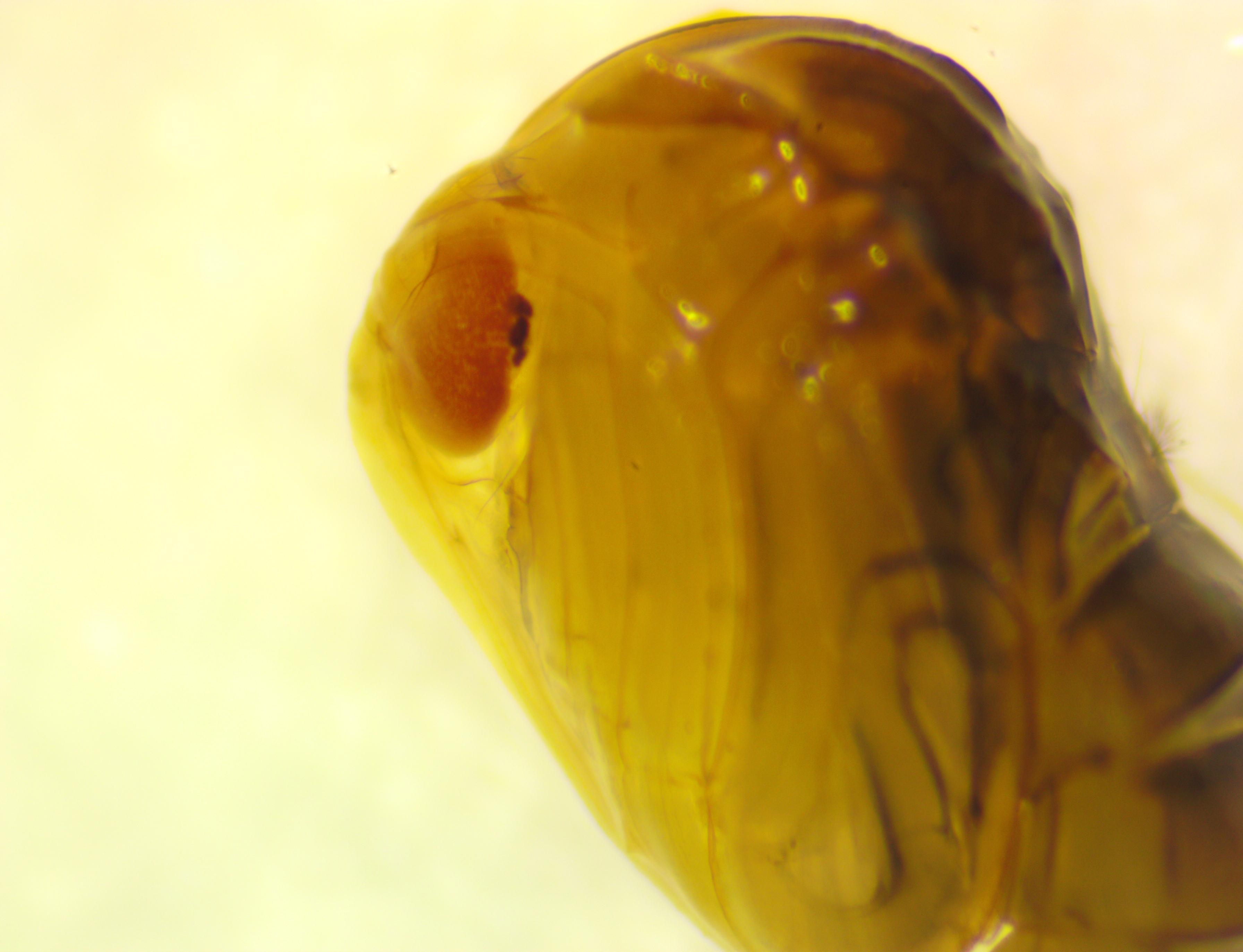

### G3_backcross_Yellow-eye_3_day_pupa_40x.jpg

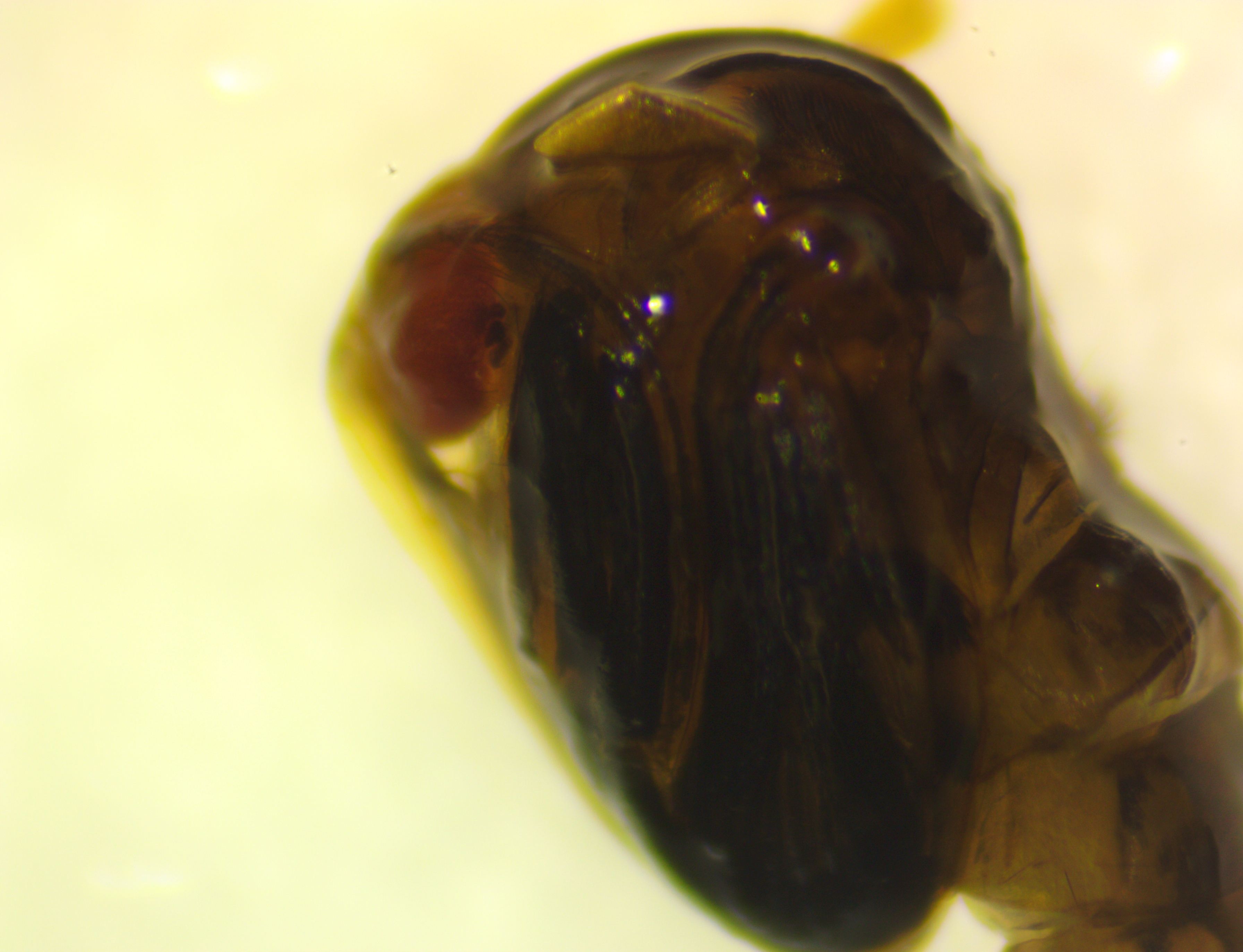

### G3_backcross_yellow-eye_female_adult_20x.jpg

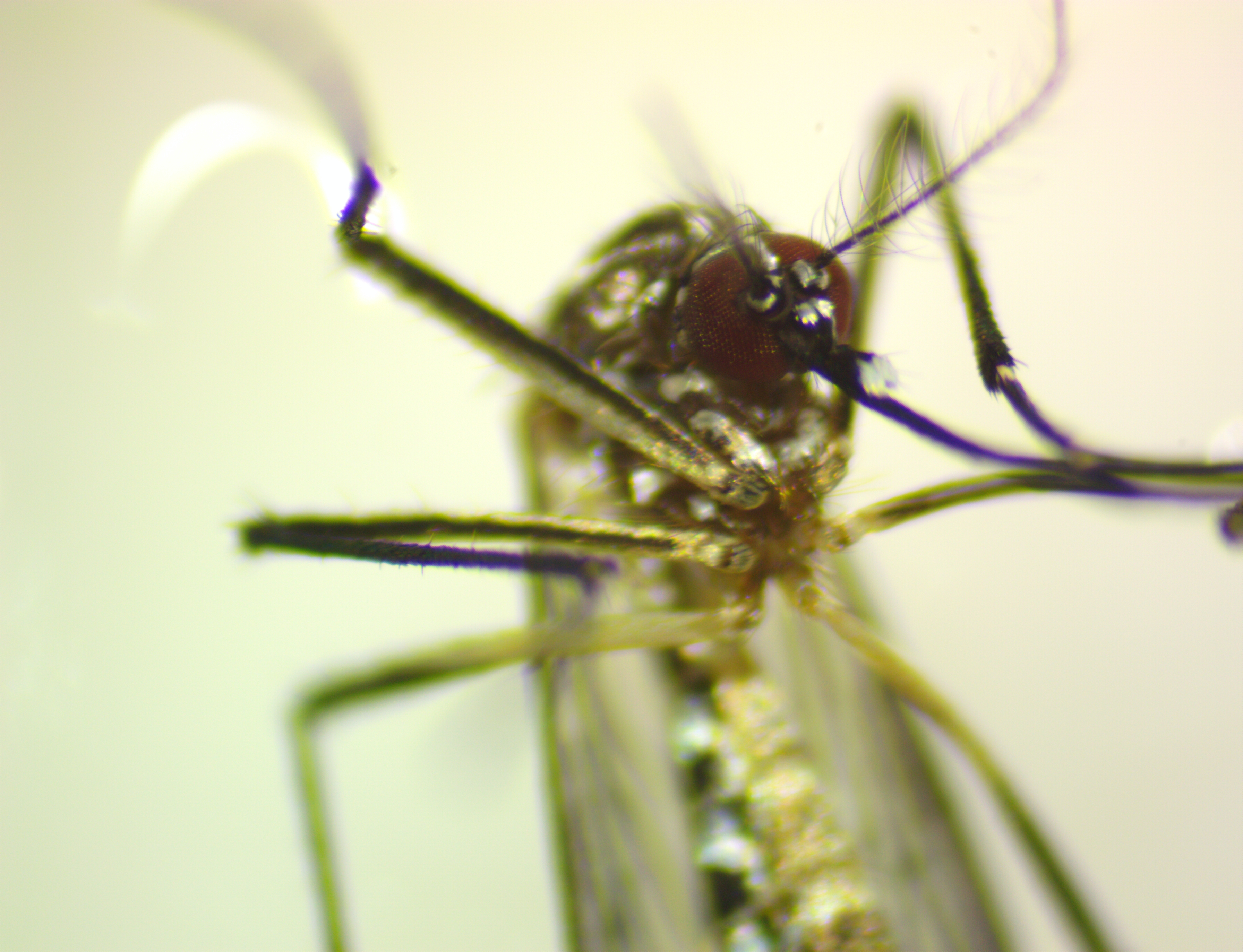

### Scale_15x.jpg

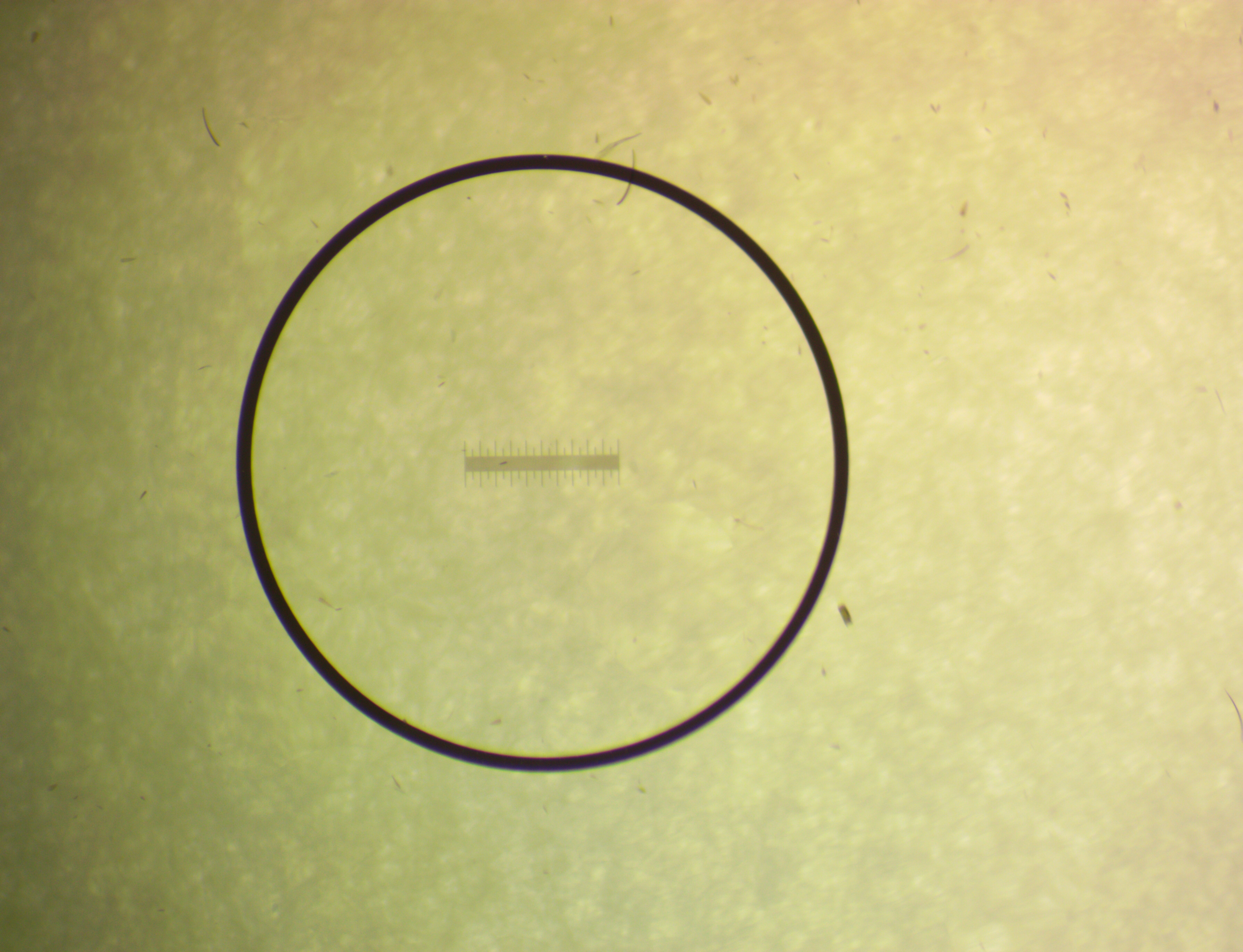

### Scale_20x.jpg

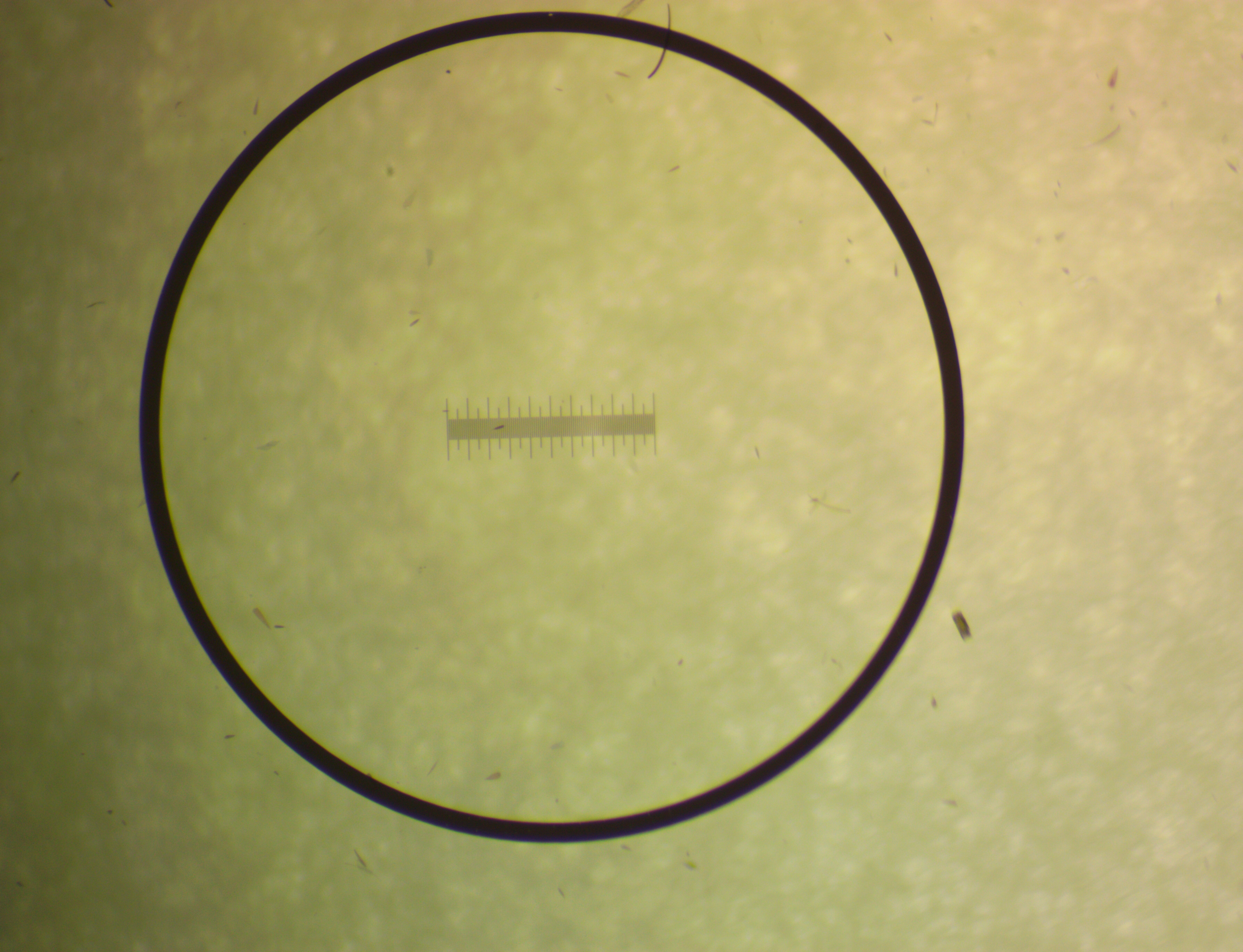

### Scale_21x.jpg

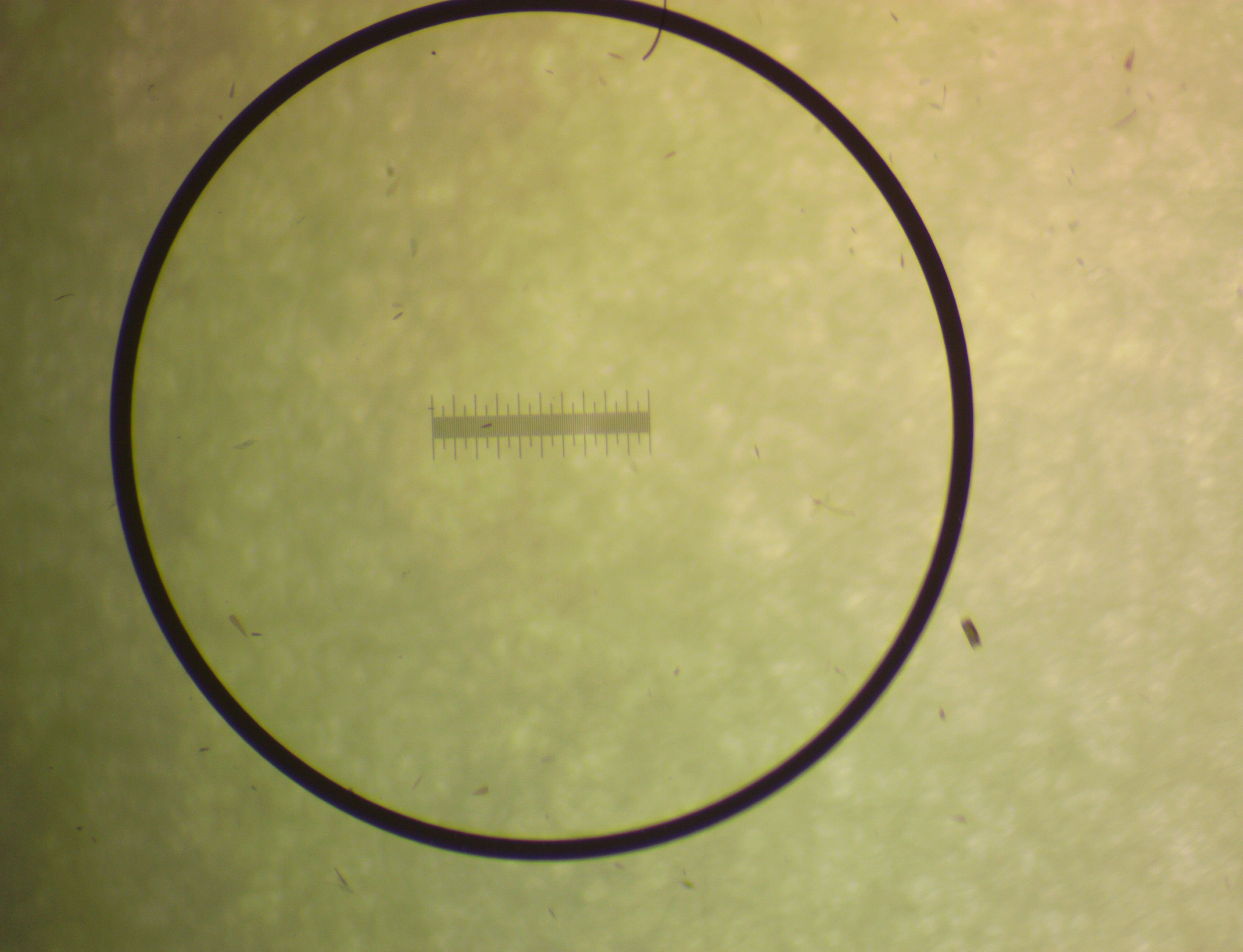

### Scale_24x.jpg

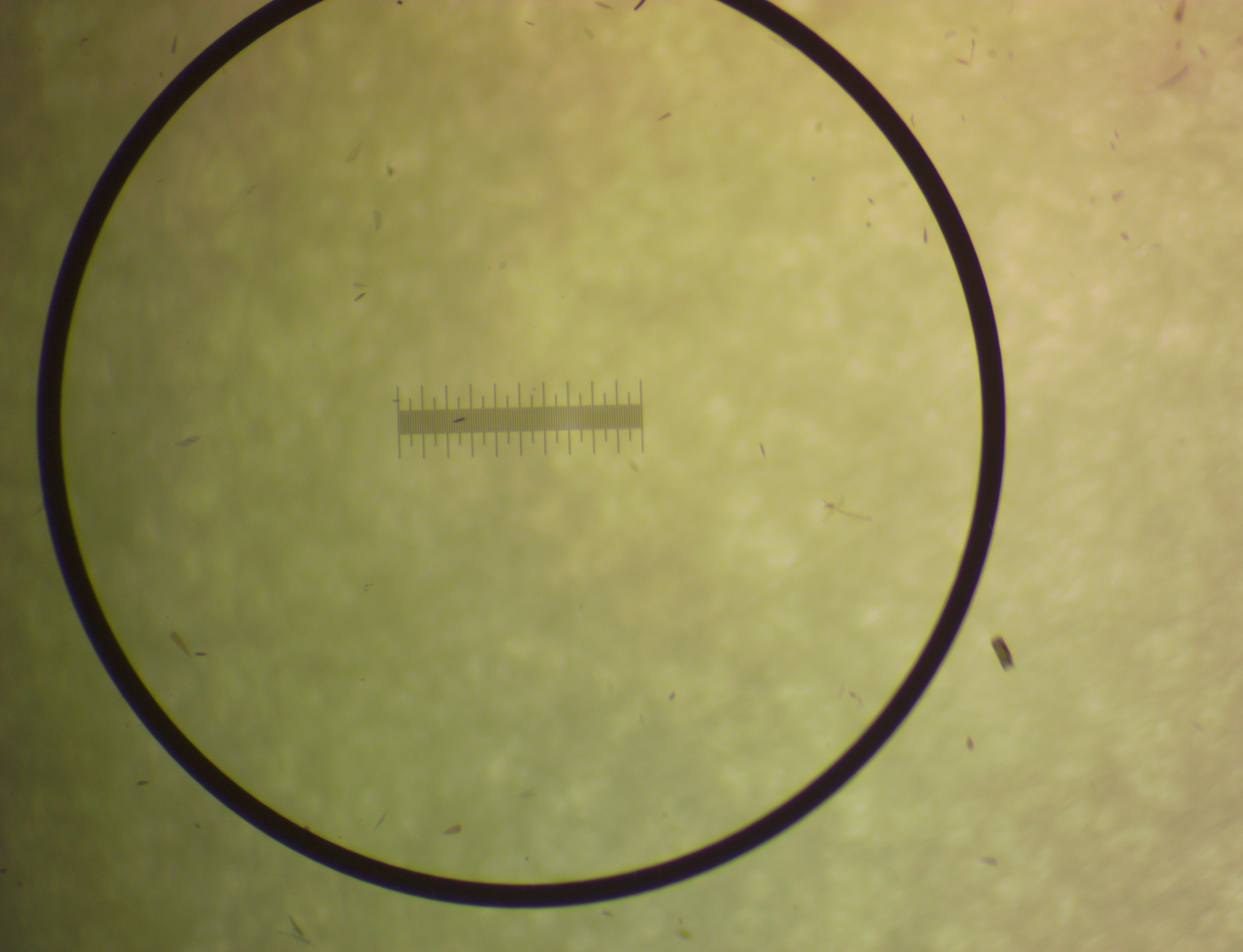

### Scale_29x.jpg

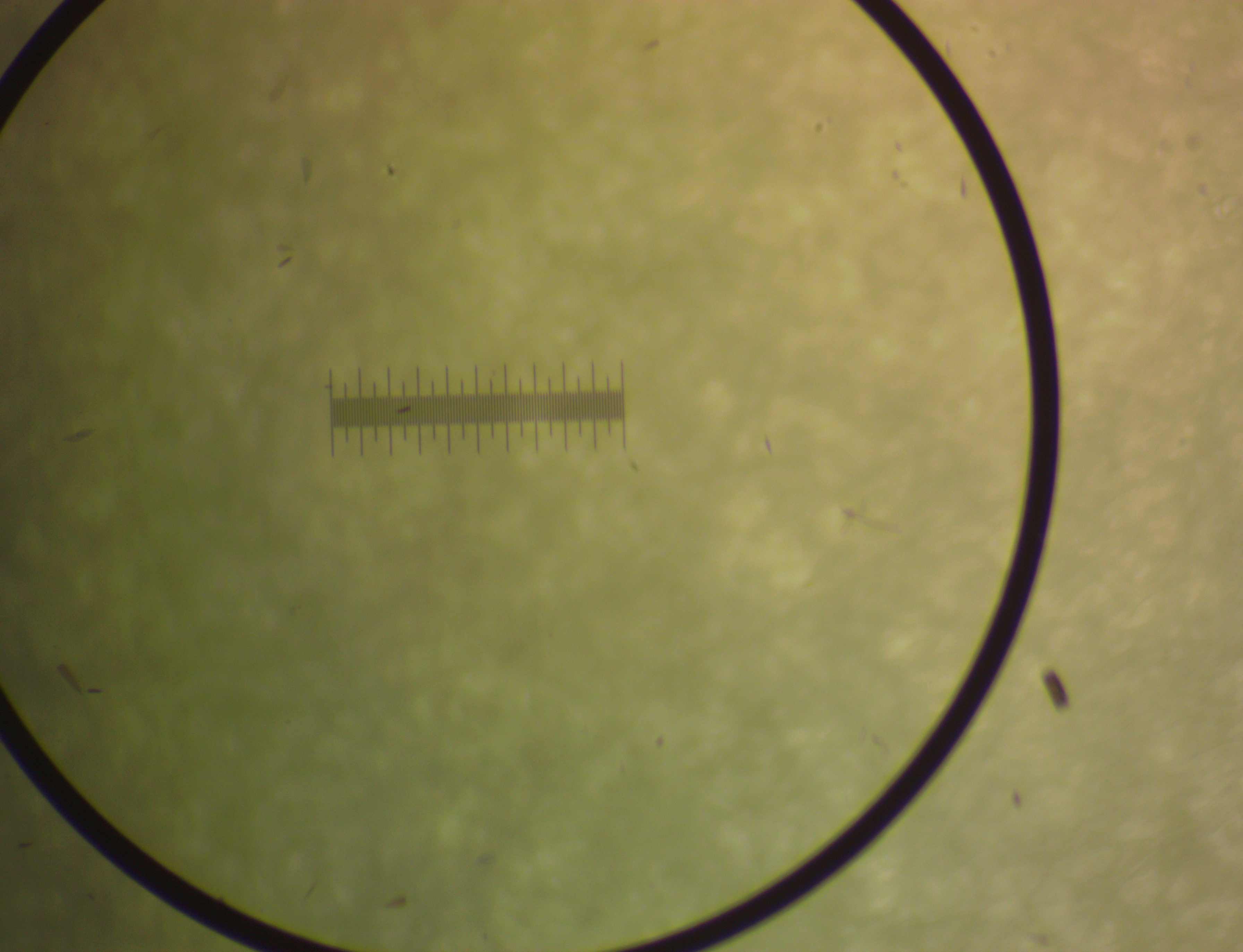

### Scale_30x.jpg

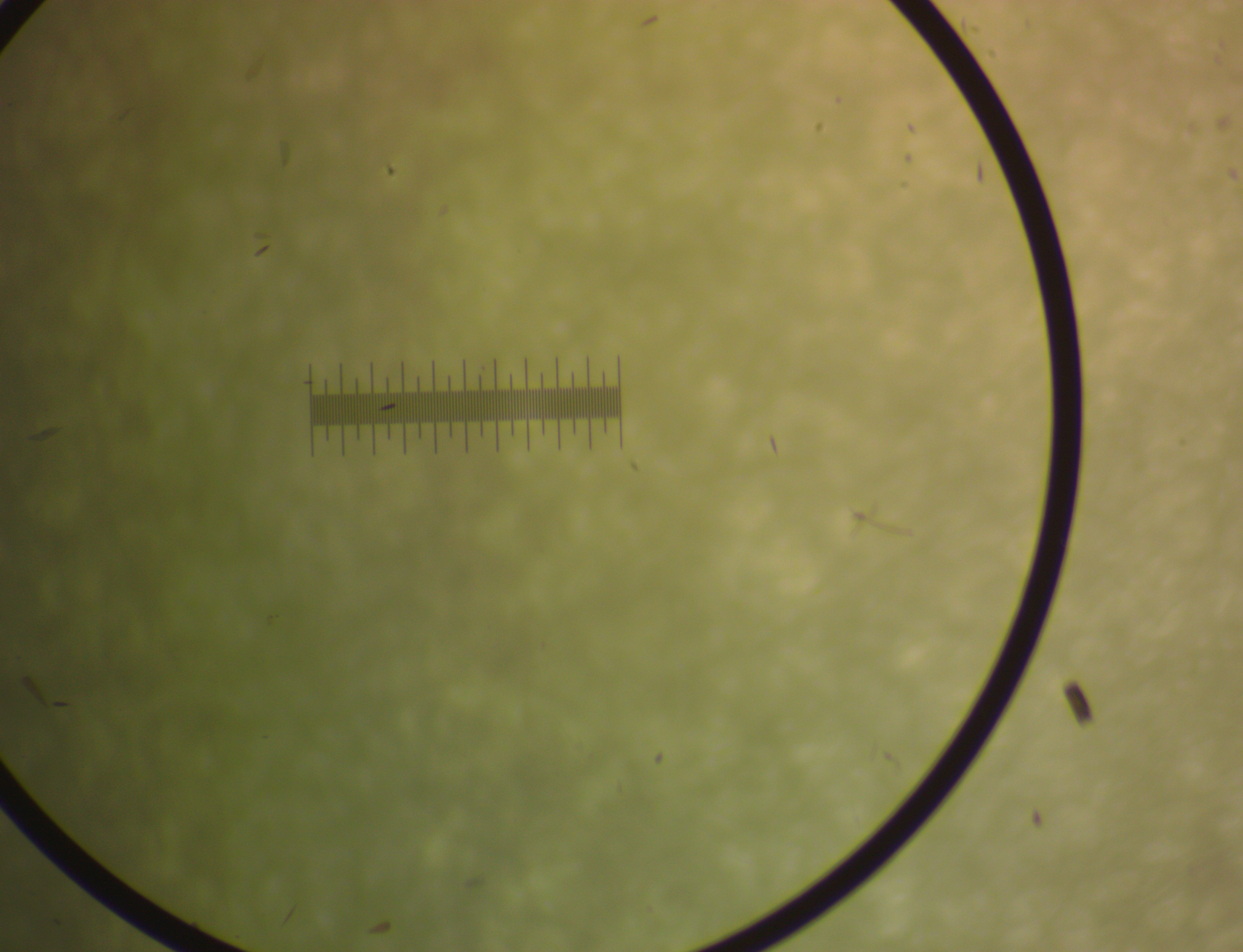

### Scale_40x.jpg

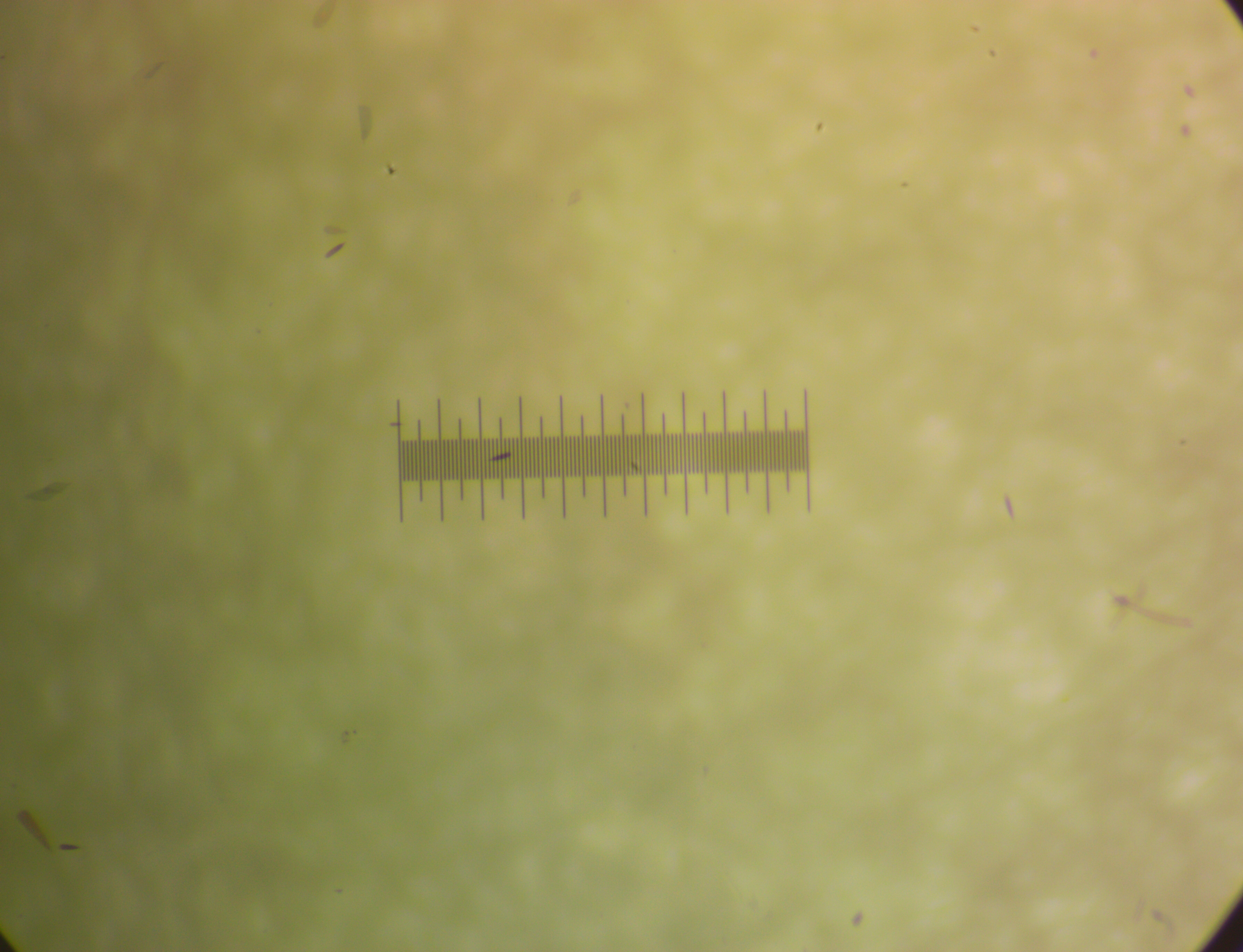
